# The 4^th^ GPCR Dock: assessment of blind predictions for GPCR-ligand complexes in the era of AlphaFold

**DOI:** 10.1101/2025.04.18.647407

**Authors:** Rezvan Chitsazi, Yiran Wu, GPCR Dock 2021 participants, Raymond C. Stevens, Suwen Zhao, Irina Kufareva

## Abstract

The GPCR Dock competitions are a series of community-wide assessments of computational structural modeling and ligand docking for G protein-coupled receptors, a central class of drug targets in the human proteome. The assessments are designed to provide an unbiased overview of progress and to pinpoint areas in need of refinement, thus shaping and directing the development of computational modeling methodologies for GPCRs. In the footsteps of 2008, 2010, and 2013 assessments, the 4^th^ round (GPCR Dock 2021) featured five diverse and challenging prediction targets and coincided with the emergence of AlphaFold, the revolutionary Artificial Intelligence (AI) technology for protein structure prediction from amino acid sequences. This report summarizes the assessment results and challenges in the context of the convergent evolution of computational and experimental structure determination techniques for GPCRs. We demonstrate that thanks to the breakthroughs in AI-powered modeling, the accuracy of modern computational models of GPCR complexes with peptides can not only approach but also exceed that of low-resolution experimental structures. However, our results highlight the unwavering need for high-resolution GPCR structure determination, especially with small molecule chemicals, and for the concurrent application of physics-based and expert-guided modeling methods.

## Introduction

With ∼800 members in the human proteome, the G protein-coupled receptor (GPCR) superfamily is a crucial class of cell membrane receptors and the center of the druggable proteome [1,2]. GPCRs specifically bind and respond to diverse endogenous agonists spanning a range of modalities, from photons, protons, and ions to peptides and proteins, and, canonically, activate intracellular signaling pathways by coupling to heterotrimeric G proteins and β-arrestins [3,4]. Highly regulated, dynamic, hydrophobic proteins, GPCRs have historically been challenging targets for structure determination [5–8]. The cryo-EM resolution revolution [9–11] led to an explosion in the number of available experimental structures of GPCRs, mostly in the active form and with intracellular effectors, and mostly with protein and peptide agonists. However, high-resolution structures with small molecules, especially antagonists and inverse agonists, continue to be challenging and scarce. Considering the importance of atomic-resolution 3D structures for the discovery and optimization of GPCR-targeted therapeutics [12], computational prediction of GPCR-ligand complex geometries plays an indispensable role in unraveling the signaling complexities and the therapeutic promise of GPCRs [13,14].

The GPCR Dock competitions [15–17] have been established as means of unbiased assessment of the state-of-the-art in computational modeling and docking for the GPCR superfamily. Conceptually similar to CASP [18], GPCR Dock capitalizes on recently solved and yet-unpublished experimental structures of receptor-ligand complex to challenge the community with blind prediction of complex geometries from amino-acid sequences (and a 2D structure for small-molecule ligands). The predictions are subsequently evaluated in terms of their similarity to the experimental ‘answers’ to identify most predictive modeling approaches and identify the areas in need of development. Because of the importance of GPCRs and the difficulty of modeling their structures, the performance in GPCR Dock assessments is a significant indicator of the advancement of protein modeling field as a whole.

In its 4^th^ iteration, the GPCR Dock 2021 competition found itself at the confluence of a dynamic evolution in both computational modeling and experimental structure determination. Learning from past challenges in the 2008 [15], 2010 [16], and 2013 [17] assessments, focusing on adenosine A_2A_, dopamine D_3_, chemokine CXCR4, 5HT_1B_, 5HT_2B_, and SMO receptors, GPCR Dock 2021 embarked on a new chapter underscored by innovation and strides in GPCR research. By primarily relying on homology modeling, the prior assessments highlighted challenges associated with robust target-template sequence alignments, precise loop geometry predictions, and diverse binding pocket identifications. By contrast, in 2021, the emergence of AlphaFold2 [19,20], a pioneering deep learning system for protein structure prediction, marked a potential breakthrough in addressing these challenges [21]. Still, despite its tremendous promise in enhancing structural predictions, AlphaFold2 also faced some limitations including those specific and applicable to GPCRs [13,22]. Against the backdrop of AlphaFold’s capabilities and challenges, the GPCR Dock 2021 competition was poised to leverage these advancements alongside experimental insights and computational innovations, to illuminate a transformative path for GPCR structural modeling and docking.

## Results

### GPCR Dock 2021 targets presented a diverse set of modeling challenges

The GPCR modeling and docking assessment 2021 was performed for five separate GPCR-ligand complexes: Apelin receptor bound to a small-molecule agonist cmpd6 (APJ/cmpd6), orphan receptor GPR139 bound to a small-molecule agonist JNJ-63533054 (GPR139/JNJ-63533054), κ-Opioid receptor bound to agonist peptide dynorphin (OPRK/dynorphin), Neuropeptide Y Receptor Y1 bound to its endogenous agonist neuropeptide Y (NPY1R/NPY), and Neuromedin U Receptor 2 bound to agonist peptide neuromedin U-25 (NMUR2/NMU25) (**Fig. 1A**). All five complexes featured active receptors and intracellularly bound heterotrimeric G proteins; however, predictions for receptor interactions with the G proteins were not evaluated as part of the assessment.

**Figure 1.**
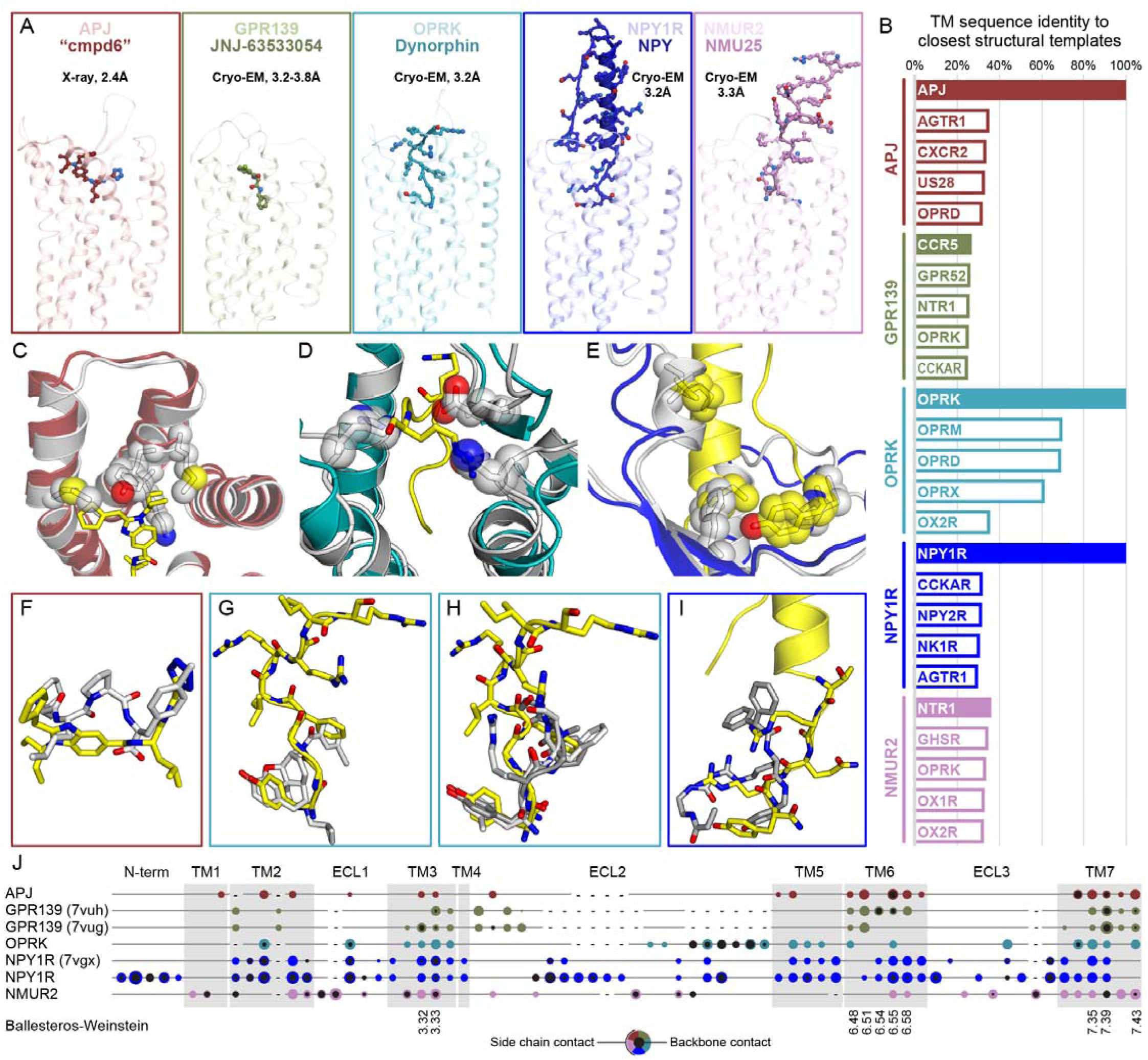
**GPCR Dock 2021 target challenges.** (**A**) From left to right: X-ray structure of Apelin receptor (APJ) bound with biased agonist Compound 6 (Cmpd6), Cryo-EM structure of GPR139 bound with agonist JNJ-63533054, Cryo-EM structure of κ-Opioid receptor (OPRK) bound with peptide Dynorphin, Cryo-EM structure of Neuropeptide Y receptor 1 bound with Neuropeptide Y (NPY), and Cryo-EM structure of Neuromedin U receptor 2 bound with Neuromedin U-25 peptide (NMU25). (**B**) TM domain sequence identity between the GPCR Dock 2021 assessment targets and the closest structural templates available in the PDB at the time of the assessment. (**C-E**) Superposition of the target APJ, OPRK, and NPY1R structures onto the relevant structure(s) of the same receptors available in PDB at the time of the assessment illustrates the need for modeling induced fit: (**C**) APJ21 (red) vs PDB 6KNM (gray), **(D)** PDB 7Y1F (teal) vs PDB 6B73 (gray), **(E)** PDB 7X9A (blue) vs PDB 5ZBQ (gray). (**F-I**) Structural overlay of the ligands in the assessment targets and ligands in the the relevant structure(s) of the same receptors available in PDB at the time of the assessment (**F**) Cmpd6 (yellow) and AMG3054 (aa 14-17), (**G**) Dynorphin (yellow) and MP1104, (**H**) Dynorphin (yellow), KGCHM07, and DAMGO, (**I**) NPY (yellow) and UR-MK299. (**J**) Key ligand-receptor contacts for the five assessment targets projected onto their sequence alignment.

These target complexes presented a diverse set of modeling challenges. Experimental structures with high sequence homology to the target (which can greatly aid the modeling efforts) were only available in the Protein Data Bank (PDB) for APJ, OPRK, and NPY1R; for GPR139 and NMUR2, only distant homologous structures existed (**Fig. 1B**). Even for receptors with available structures, participants had to predict large ligand-induced conformational changes in the receptor (induced fit) (**Fig. 1C-E**). Three of five targets required predictions of conformations for large flexible peptides and the remaining two for previously structurally uncharacterized small molecules. For some targets, the task was additionally complicated by insufficient or narrow ligand structure-activity relationships (SAR) and/or by limited available receptor binding pocket mutagenesis, with details provided below.

**APJ/Cmpd6.** There were two X-ray structures available for APJ, one bound to an unrelated agonist peptide (AMG3054, PDB 5VBL [23]) and another to a single domain antibody (JN241-9, PDB 6KNM [24]). Compound 6 (Cmpd6) is a benzimidazole carboxylic acid amide derivative from a series patented by Sanofi [25] and a (most likely G protein-biased) agonist of APJ [26,27]. The crystalized constrained peptide AMG3054 had features similar to Cmpd6 but their spatial arrangement in the pocket was quite different (**Fig. 1F**), which might have misguided attempts of 3D-pharmacophore-assisted ligand docking. There was extensive SAR data available for benzimidazole carboxylate amides (including Cmpd6 [25]) and a number of other APJ agonist chemotypes (pyrazoles [28,29], hydroxypyridinones [30,31]). Mutagenesis studies for APJ are limited [23,32,33] and none is related to the Sanofi compound series.

**GPR139/JNJ-63533054.** At the time of the assessment, there were no available experimental structures for this receptor, and the closest homologous structure (CCR5) had only 27% sequence identity in the TM domain (**Fig. 1B**). Actively pursued as a target in neuropsychiatric disorders [34], GPR139 has a number of known small molecule agonists [35–40], with one, TAK-041 (Zelatriazin), in a Phase 2 clinical trial (NCT03319953 [41]). The [3-chloranyl-N-[2-oxidanylidene-2-[[(1S)-1-phenylethyl]amino]ethyl]benzamide] JNJ-63533054 [36] is an orally available, brain-penetrant [42] GPR139 agonist developed by JnJ. Extensive SAR data is available for this and other GPR139 agonist series [35,36,43,44]. The agonist binding site on GPR139 has been mapped through extensive mutagenesis [43,45,46], providing hints regarding the compound binding geometry for this receptor.

**OPRK/dynorphin A (1-13).** For this receptor, multiple structures bound with agonists (e.g. MP1104, PDB ID 6B73 [47]) and antagonists (e.g. JDTic, PDB ID 4DJH [48]) existed in the PDB at the time of the assessment, but peptide-bound structures were not available. This said, structures with bound peptides were available for OPRD and OPRM, the two closest homologs of OPRK (PDB IDs 6PT2 [49], 6DD[EF] [50], 4RW[AD] [51]). These structures provided good constraints for dynorphin residues 1-4 but not for the rest of the peptide (**Fig. 1G-H**); in fact, their conformations were not open enough to accommodate the full-length dynorphin peptide (**Fig. 1D**) or recapitulate its interactions with OPRK ECLs (**Fig. 1J**). An extraordinarily potent endogenous OPRK peptide agonist, dynorphin A has been studied for decades; both the roles of individual peptide residues [52–56] in OPRK binding and activation and the roles of OPRK pocket residues in interactions with dynorphin, have been extensively explored by mutagenesis [48,57–59].

**NPY1R/NPY.** A 36 aa peptide, NPY was structurally characterized by NMR in 1995 [60] and shown to feature a C-terminal α-helix stapled to the unstructured N-terminus with a disulfide bond. For the receptor, two X-ray structures existed in the PDB at the time of the assessment, both solved with small molecule antagonists (PDB 5ZBQ, 5ZBH) [61]. One of the molecules, UR-MK299 (PDB 5ZBQ), was chemically quite similar to the ‘business end’ (C-terminus) of NPY and also shared a similar spatial arrangement in the binding pocket (**Fig. 1I**). However, the binding of the full NPY peptide promoted an opening of the binding pocket and substantial rearrangements in the receptor extracellular loops, necessitating prediction of induced fit (**Fig. 1E**) and extensive peptide contacts with receptor ECLs (**Fig. 1J**). With ∼31% TM sequence identity, the cholecystokinin receptor type A (CCKAR) represented a suitable homology modeling template (**Fig. 1B**); what is more important, CCKAR complexes with CCKN (PDB IDs 7MBX, 7EZH) [62,63] provided a highly structurally similar template for both the receptor in an active agonist-bound conformation, and the agonist peptide with an amidated C-terminus in the receptor’s pocket. Additionally, extensive mutagenesis studies on both the receptor [64–68] and the peptide [69–75] have been performed and could be used to guide peptide placement.

**NMUR2/NMU25.** A member of the “neuromedin” family of peptide regulators of smooth muscle contraction, Neuromedin U (NMU) was first identified in 1985 [76] in two biologically active forms, one containing 25aa (NMU25) and another only the C-terminal 8aa (NMU8). Subsequent studies detailing the synthesis, SAR, and biological properties [77–80] revealed the requirement for C-terminal amidation and an almost strict conservation in the C-terminal 8 amino-acids across species [78], suggesting role of the C-terminus as a key signaling modality. NMUR2 is one of two receptors for NMU25 [81,82], discovered in 1998 [83] and expressed in the paraventricular nucleus of hypothalamus and elsewhere in the CNS. There were no experimental structures for NMUR2 at the time of assessment and the closest structurally characterized homolog, the Neurotensin receptor 1 (NTSR1) had only ∼35% sequence identity in the TM domain (**Fig. 1B**). Extensive SAR data for the NMU peptide [77,78,84,85], but not for the receptor, was available to guide peptide docking.

In addition to the listed experimental structure of target and homologous receptors, models of all five receptors, in *apo* form, have also been predicted by the AlphaFold Monomer V2.0 pipeline and were available publicly in AlphaFold Protein Structure Database [86] at the time of the competition.

In summary, the GPCR dock 2021 targets presented a wide range of modeling challenges, including small-molecule and peptide complexes, X-ray and cryo-EM structures, varying levels of homology to previously solved GPCR structures, and complexity of ligand-induced conformational changes. However, considerable ligand SAR information was available for all five targets, and receptor mutagenesis for three out of five targets, which could assist and guide model selection.

### Structural variation across experimental ‘answers’

An important aspect of the GPCR Dock 2021 assessment was the availability, after the model submission deadline, of multiple experimental ‘answers’ for four out of five studied complexes (all except APJ/cmpd6, **Supp. Data 1**), with substantial structural variation among them. For APJ, no published structures still exist with cmpd6; however, a cryo-EM and an X-ray structure with a close analog from the same series, the Sanofi compound CMF-019 [25,26], were published in 2024 [87,88] (**Table 1**), similarly featuring differences from the ShanghaiTech APJ/cmpd6 X-ray structure. This made us consider the following questions: Why are the ligand poses in these experimental structures so different? Are any of the experimental structures ‘better’ or more ‘correct’ than others? And what is the ultimately ‘correct’ geometry of the complexes that the computational community should strive to predict?

**Table 1.** “Gold-standard” and other experimental structures for small-molecules targets in GPCR Dock 2021.

| PDB | Receptor | Ligand | Fusions and IC effector(s) | Method | Resolution | Release date | Reference |
| --- | --- | --- | --- | --- | --- | --- | --- |
| APJ21 | APJ | Cmpd-6 | Rubredoxin @ ICL3 | X-ray | 2.40 Å | n/a | Supp. Data 1 |
| 8XZI | APJ | CMF-019 | Gaiβγ + scFv16 | Cryo-EM | 2.70 Å | 2024-03-20 | [87] |
| 8S4D | APJ | CMF-019 | bRIL @ ICL3 | X-ray | 2.58 Å | 2024-12-04 | [88] |
| 7VUG | GPR139 | JNJ-63533054 | Gaiβγ + scFv16 | Cryo-EM | 3.20 Å | 2021-12-29 | [89] |
| 7VUH | GPR139 | JNJ-63533054 | miniGas/q (apo) + Gβγ + Nb35 | Cryo-EM | 3.22 Å | 2021-12-29 | [89] |
| 7VUI | GPR139 | JNJ-63533054 | miniGas/q (GTP-bound) + Gβγ + Nb35 | Cryo-EM | 3.30 Å | 2021-12-29 | [89] |
| 7VUJ | GPR139 | JNJ-63533054 | miniGas/q (GDP-bound) + Gβγ + Nb35 | Cryo-EM | 3.80 Å | 2021-12-29 | [89] |

The APJ complex with Cmpd6 was the most striking example of experimental structure variation. Cmpd6 and CMF-019, with 37 and 32 non-hydrogen atoms, respectively, share a substructure of 29 atoms and all key pharmacophores described in [25]; yet the root mean square deviation for the maximum common substructure (MCS) of the two compounds after optimal superimposition of the receptor TM domains is as high 9.9Å (**Supp.** Fig. 1A). Arguably, the structures were solved by different methods (X-ray vs Cryo-EM, **Table 1**), using different receptor constructs (ICL3-rubredoxin fusion for the Cmpd6-bound structure, ICL3-bRIL fusion for the CMF-019 complex X-ray PDB 8S4D [88], WT receptor for the CMF-019 complex cryo-EM PDB 8XZI [87]), and in different conditions (PDB 8XZI features a heterotrimeric G protein while the two X-ray structures do not), offering reasons for the observed differences; however, the complete flip in the compound bindings pose (**Supp.** Fig. 1B) was still unexpected and hardly rationalizable.

In the case of GPR139, GPCR Dock 2021 capitalized on the structures from the cryo-EM study [89] that were solved shortly prior but still unpublished at the time of the assessment. As many as 4 structures were solved, all featuring the same receptor and the same agonist but varying in intracellular binding partners/effectors and resolution (**Table 1**). Importantly, the poses of the JNJ-63533054 compound also varied between these structures; moreover, the highest resolution structure, the Gi-coupled PDB 7VUG, features two additional alternative conformers (P1 and P2) for JNJ-63533054 in the binding pocket. The observed compound poses differ from one another by overall vertical shifts of up to 1.8Å and an orientation flip of the extracellularly facing chlorobenzene ring, leading to pairwise compound root-mean-square deviations (RMSDs) of 0.4Å to 2.8Å (**Supp.** Fig. 1C-D), following optimal superimposition of the receptors. Proximal to the chlorobenzene ring, the GPR139 residue W170^ECL2^ is lowered into the binding pocket by >3Å in PDB 7VUG, compared to the second highest resolution structure, PDB 7VUH, and the remaining structures. Differences in the rotameric states among other pocket residues similarly lead to distinct contacts with the ligand, e.g. the formation of a strong hydrogen bond with N271^7.39^ in PDB 7VUG and 7VUI but not in the other two structures. Due to the uncertainty in predicting the orientation of the chlorobenzene ring and part of its attached linker, this part might have been more challenging to predict (**Supp.** Fig. 1D).

For the OPRK complex with Dynorphin A (1-13), the cryo-EM structure solved by Fei Xu’s group at ShanghaiTech formed the basis for the assessment and was later published as PDB 7Y1F [90]. Another cryo-EM structure solved by the teams of Bryan Roth (UNC) and H. Eric Xu (ShanghaiTech) was released on 2022-12-14 (PDB 8F7W [91]), after the GPCR Dock 2021 model submission deadline and while this paper was in progress. The latter structure featured a marginally higher resolution (3.19 Å for PDB 8F7W vs 3.20 Å for PDB 7Y1F), and one fewer resolved residue for the peptide (aa 1-8 for PDB 8F7W vs aa 1-9 for PDB 7Y1F, **Table. 2**). Notably, the conformations of the resolved N-terminal parts of the Dynorphin A peptide are quite distinct (backbone RMSD of 2.87Å), and substantial differences in peptide-receptor pocket residues interactions are observed as well (**Supp.** Fig. 1E-F).

For the NPY1R/NPY complex, similarly, one cryo-EM structure (PDB 7X9A [92]) was solved by the Qiang Zhao/Beili Wu collaboration at ShanghaiTech, before the assessment (but not released until later) and another structure (PDB 7VGX [93]) by Hee-Jung Choi group at Seoul National University; both structures were published in 2022. The structures have the same resolutions, with the common ordered region of the peptide consisting of residues 21-36 (**Table. 2**). The TM helices of the receptor are systematically shorter, by 1-1.5 Å, in PDB 7X9A than in PDB 7VGX (e.g. the distance between the backbone oxygens of R254^6.26^ and T284^6.56^ is 44.08 Å in PDB 7X9A but 45.58 Å in PDB 7VGX), making the optimal superimposition of the TM domains challenging. The structured parts of the NPY peptide are well aligned (heavy-atom RMSD of 2.0Å following receptor superimposition, **Supp.** Fig. 1G), although there is a slight upward shift of the peptide in PDB 7VGX compared to 7X9A (**Supp.** Fig. 1H).

Finally, for NMUR2/NMU25, one cryo-EM structure was solved at ShanghaiTech by the Zhao/Wu team (PDB 7XK8 [94]) and another by the Xu/Jiang collaboration (PDB 7W55 [95]). Both structures were released in 2022. The latter structure had a substantially higher resolution (2.80 Å for PDB 7W55 vs 3.30 Å for PDB 7XK8 or its early pre-release refinement, ‘pre-7XK8’, **Supp. Data 2**). The shared ordered region of the NMU25 peptide in both structures encompassed residues 18-25. Residues 21-25 of the NMU25 peptide in both structures are well aligned (after receptor superposition); however, Phe19 and Leu20 face in the opposite directions in the two peptides, resulting in the backbone RMSD of 2.65 Å and all-heavy-atom RMSD of as high as 4.03 Å (**Supp.** Fig. 1I-J). In the 7W55 geometry, the peptide features an intramolecular T-stacking between the side chains of Phe19 and Phe21 and a favorable backbone hydrogen bonding to the receptor N306^7.32^. Conversely, the orientation of Phe19 in PDB 7XK8 leads to an outward movement of TM6, as a result the binding pocket is tighter around TM5-TM6 in 7W55 compared to 7XK8.

Considering this plethora of experimental structures (**Tables 1-2**), we ran all model analyses against each. However, in the remaining parts of the manuscript below, we primarily focus on model comparisons to the highest resolution structure of each complex: the unpublished ShanghaiTech X-ray structure for APJ-Cmpd6, PDB 7VUG for GPR139/JNJ-63533054, PDB 8F7W for OPRK/Dynorphin, PDB 7X9A for NPY1R/NPY, and PDB 7W55 for NMUR2/NMU25. For the purpose of the discussion only, we refer to these structures as our “gold-standard” answers. Model analyses against other structures are presented in Supplementary Materials.

**Table 2.** “Gold-standard” and other experimental structures for peptide targets in GPCR Dock 2021.

| PDB | Recept or | Ligand (used in structure determination) | Ordered part of the ligand | IC effector(s) | Resolutio n | Release date | Reference |
| --- | --- | --- | --- | --- | --- | --- | --- |
| 7Y1F | OPRK | Dynorphin A (1-13) | aa 1-9 | Gaiβγ + scFv16 | 3.20 Å | 2023-05-24 | [90] |
| 8F7W | OPRK | Dynorphin A (1-13) | aa 1-8 | Gaiβγ + scFv16 | 3.19 Å | 2022-12-14 | [91] |
| 7X9A | NPY1R | Neuropeptide Y (1-36) | aa 1-36 | Gaiβγ | 3.20 Å | 2022-05-18 | [92] |
| 7VGX | NPY1R | Neuropeptide Y (1-36) | aa 1-5,20-36 | Gaiβγ + scFv16 | 3.20 Å | 2022-02-23 | [93] |
| pre-7XK8 | NMUR2 | Neuromedin-U-25 (1-25) | aa 1-25 | Gaiβγ | 3.30 Å | n/a | Supp. Data 2 |
| 7XK8 | NMUR2 * | Neuromedin-U-25 (1-25) | aa 1-25 | Gaiβγ | 3.30 Å | 2023-02-22 | [94] |
| 7W55 | NMUR2 | Neuromedin-U-25 (1-25) | aa 18-25 | Gaqβγ + scFv16 | 2.80 Å | 2022-04-20 | [95] |

\* Final published refinement of the 2021 ShanghaiTech structure

For each of the three peptide-bound targets, their respective peptides in the structures had a ‘core’ part binding deep in the receptor pocket and a more flexible part facing towards the solvent on the extracellular side. The OPRK structures were solved with Dynorphin A (1-13); however, its C-terminal part was completely unresolved in both structures and residue Arg9 in PDB 7Y1F had only superficial contacts with the receptor, prompting us to focus on predictions for aa 1-8 only. We also ran a secondary set of analyses for the distal N-terminal part, aa 1-5, which effectively form another OPRK agonist, Leu-enkephalin. For NPY, the outward-facing parts had very few contacts with the receptor in PDB 7X9A and were not modeled at all in PDB 7VGX, so we focused our attention on predictions for the peptide core (aa 21-36). For NMUR2, similarly, the outward-facing (N-terminal) part had only sparse contacts with the receptor in PDB 7XK8 and were fully disordered in PDB 7W55, so we chose to use the core (18-25 aa) for the assessment.

In conclusion, the dynamic nature of the target GPCR-ligand complexes, and possibly the limited resolution of the available experimental structures, created substantial uncertainties in what would otherwise be considered the “correct” structure of each complex, and confounded the assessment of the computational models. It might also have presented challenges for the computational community that strives to accurately predict these dynamic conformations.

### 45 groups from 14 countries submitted 904 models for the assessment

Forty-five groups (defined as institution/PI-associated teams with non-redundant modeling methodologies (**Supp. Table 1**) from 14 countries (**Supp.** Fig. 2A) participated in GPCR Dock 2021 and submitted a total of 904 models (**Supp. Data 3**). Among them, 18 models with no ligand, no receptor, and severe clashes (sterically unfeasible complexes) were considered uninterpretable (**Supp. Table 2**). As a result, 206, 198, 175, 155, and 152 models were assessed for APJ, GPR139, OPRK, NPY1R, and NMUR2, respectively (**Supp.** Fig. 2B).

The models were assessed by several criteria. The primary criteria for this assessment included ligand position, assessed as the RMSD on its non-hydrogen atoms following optimal superimposition of the receptors, and the shared atomic contacts between the ligand and the receptor (**Supp.** Fig. 3). These two criteria were converted into a single score representing the likelihood of observing similar deviations among experimentally solved high-resolution X-ray co-crystal structures (the so-called model ‘‘correctness’’ [16,17]) and used for final model ranking. Results are presented in (**Supp. Table 3**). For peptide targets, DockQ score [96], which takes the CAPRI measures [97–99] and formulates them into a single score for the protein-docking quality assessment, was included in this round of GPCR Dock for a comparison; we compared it to the GPCR Dock score (a.k.a. “correctness”) to assess the degree of their agreement across the model set. Finally, we also monitored the accuracy of transmembrane (TM) bundle structure prediction, as well as the structure of the extracellular domains and loops. Altogether, this process provided a thorough assessment of model quality through rigorous and relevant criteria, highlighting global effort and varied approaches in aligning models with high-resolution experimental data.

**Table 3.** GPCR Dock 2021 models predicted with AlphaFold2.

| Target | Submitted models* | Models with methods | Initial models with AF | AF (apo) | AF (complex) | Other/ unknown ** | % models with methods*** | % all models **** |
| --- | --- | --- | --- | --- | --- | --- | --- | --- |
| APJ/Cmpd6 | 201 | 150 | 61 | 61 | - | 89 | 40.7 | 30.3 |
| GPR139/JNJ | 198 | 155 | 106 | 106 | - | 49 | 68.4 | 53.5 |
| OPRK/Dynorphin | 170 | 128 | 70 | 61 | 9 | 42 | 54.7 | 41.2 |
| NPY1R/NPY | 155 | 119 | 98 | 36 | 62 | 21 | 82.4 | 63.2 |
| NMUR2/NMU25 | 152 | 115 | 97 | 31 | 66 | 18 | 84.3 | 63.8 |

\* Submitted models after removing group ID 8067
\*\* Other approaches/Unknown from the submitted methods = Models with Methods - Initial Models with AF
\*\*\* Initial Models with AF / Submitted Models

### Small-molecule targets: ligand predictions for GPR139 but not APJ approach experimental accuracy

**Fig. 2A-B** shows the plots of ligand RMSD vs. the fraction of correctly predicted contacts for the models of APJ/Cmpd6 and GPR139/JNJ-63533054 complexes when compared against the ShanghaiTech APJ/Cmpd6 X-ray structure and against PDB ID 7VUG, respectively. As evident from this plot, the models spanned a wide range of accuracies and for most, the compound RMSD from the experimental answer was as high as 5-10 Å, with less than 20% and 40% of the receptor-ligand contacts predicted correctly for APJ and GPR139, respectively. However, for a small number of models, their RMSD/contact combination placed them closer to the top left corner of the plot and to the shaded area which represents the distribution of the corresponding parameters among experimental high-resolution structures of complexes of identical composition in the PDB [16]. Depending on their percentile within that distribution, the models were ranked by “correctness”, measured in percent of the experimental pair distribution. The correctness for the best-ranking model of the APJ/Cmpd6 complex, **SDU-Yang** (model APJ-1912-0005) was 0.61%, with ligand RMSD of 3.47Å following the optimal superimposition of receptor TM domains, and 31.3% correctly predicted contacts (**Fig. 2C**). Interestingly, when compared to the ligand maximum common substructure in the CMF-019 complex (PDB 8XZI [87]), the best model, **KIST-Park** (model APJ-8824-0001) reached the correctness of 5% with ligand MCS RMSD as low as 1.18Å and 51.2% correctly predicted contacts, and at least 5 more models had correctness above 1% (**Supp.** Fig. 4A-B). For the most correct (according to the comparison with PDB ID 7VUG) model of the GPR139/JNJ-63533054 complex, **UW-DiMiao** (model GPR139-2717-0003), the correctness was 2.50%, with 2.05Å ligand RMSD and 47.80% of correctly predicted contacts (**Fig. 2D**). Same model also scored best when compared against PDB 7VUH or the alternative ligand conformers in PDB 7VUG (**Supp.** Fig. 4C-F)

**Figure 2.**
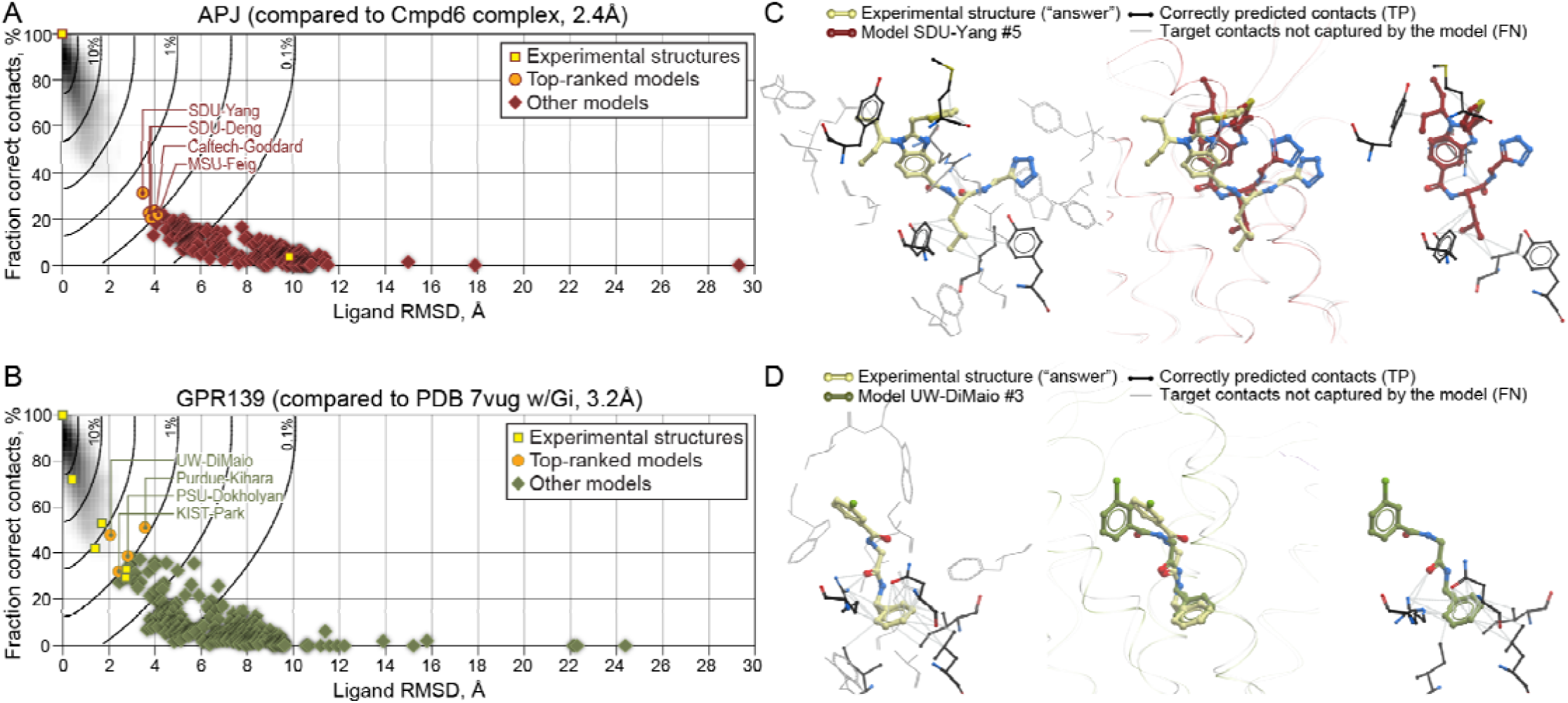
**Small-molecule targets: ligand predictions for GPR139 but not APJ approach experimental accuracy.** (**A-B**) Prediction of ligand pose (measured as heavy-atom RMSD) and ligand-pocket contacts by the APJ/Cmpd6 models when compared against APJ/Cmpd6 X-ray structure (**A**) and by GPR139/JNJ models when compared to PDB vug (**B**). The shaded background represents the distribution of the plot parameters observed across high-resolution X-ray structures in the PDB [16,17]. Solid black curves represent isolines of the model “correctness” used for model sc ring and ranking. (**C-D**) Comparison of the ligand predictions in the highest-correctness submitted models to the experimental a swer for APJ (**C**) and GPR139 (**D**). Left panels illustrate the target ligand-pocket contacts that are correctly predicted (bla k sticks) or not captured (gray wires) by the model, middle panels the overlay of compound poses following the optimal superimposition of receptor TM domains, and right panels the correctly predicted pocket contacts with the modeled ligands.

While these correctness percentiles (5%, 2.50%, and especially 0.61%) may seem low, it is important to realize that they were measured in the context of the *high-resolution X-ray structure* distribution (shaded areas in **Fig. 2A-B**). The recent progress in cryo-EM structure determination flooded the PDB with lower resolution structures of highly dynamic complexes such as those of activated GPCRs with their ligands. To place findings in this context, we thus projected onto the same scatter plots the calculated RMSD and % contacts values when the experimental structures of our targets are compared to one another. Notably, we found that a few best-ranking models of the GPR139/JNJ complex (orange circles in **Fig. 2B**) were closer to the experimental answer in PDB ID 7VUG than other, lower resolution experimental structures of the same complex (yellow squares in **Fig. 2B**) were to the same PDB. Therefore, we concluded that the precision of the top-ranking models was well within the experimental error observed in modern cryo-EM GPCR-ligand complex structures. Needless to say, the difference between the two APJ structures with the Sanofi analogs (Cmpd6 and CMF-019) was far outside of any expected experimental uncertainty (**Supp.** Fig. 4A).

It is also important to note that the close-to-experimental correctness was only achieved by a few submitted models. Considering that model building methods and practices, and researcher’s expertise, remain key factors of success, we looked closely into the modeling methods used by the submitters of the winning models.

According to their submitted methods (**Supp. Data 4**), the **SDU-Yang** team started with AlphaFold V2.0 [20] to predict the APJ model using a customized MSA and applying AlphaFold default settings. Then, they submitted the predicted model to the COACH-D server [100] to predict the ligand binding site and initial APJ/Cmpd-6 complex. The Rosetta Ligand Docking Protocol was applied as the final step to dock the ligand into the receptor predicted by AlphaFold. They selected the five submitted models by taking into account the energy scores, consistency with the predicted binding site, and steric clashes (**Supp. Data 4**). The best predicted model (APJ-1912-0005) is in a reasonable agreement with the APJ/Cmpd6 X-ray (**Fig. 2C**); however, the benzimidazole core of the ligand and its tetrazole ring (surrounded by TM7, TM1, and TM2) are slightly misaligned with the answer, which results in the divergence of di-ethyl chains (surrounded by TM5 and TM6) in the binding pocket. Consequently, this best model is still at 3.47 Å ligand RMSD and 31.3% contact prediction accuracy (“correctness” of 0.61%), relative to the APJ/Cmpd6 X-ray.

For their top CMF-019-like prediction (model APJ-8824-0001), **KIST-Park** built the apo receptor model using AlphaFold2 and selected the prediction with the highest pLDDT for subsequent docking with Rosetta GALigandDock. They picked models 1-3 by visual inspection and models 4-5 based on the rank predicted by DeepAccNet-ligand score [101]. In addition to the top-ranking APJ-8824-0001, one more model from their submission (APJ-8824-0003) had a correctness >1% (1.23%) compared to the MCS with CMF-019 in PDB 8XZI. This group also generated the 4^th^ most accurate prediction for the GPR139/JNJ-63533054 complex based on the PDB 7VUG comparison, and the 2^nd^ most 7VUH-like: GPR139-8824-0005 (correctness 1.41%, ligand RMSD 2.08, 34.74% correctly predicted contacts), using the same methodology.

**UW-DiMiao** used AlphaFold [20], RoseTTAFold [102], and RosettaCM [103] in their approach to predict the structures of the apo GPR139 from its amino-acid sequence. In addition, this group built partial threaded models using RosettaCM [103] based on predicted homologs from HHpred [104]. Then, Rosetta GALigandDock [105] was used to dock the ligand into the predicted receptor structures. Complex geometries where ligand polar atoms were buried without forming hydrogen bonds were excluded, and binding affinities estimated at the docking step were considered. The use of prior SAR or mutagenesis studies is not mentioned. The final five models selected for submission included two top models from AlphaFold, the top model from RoseTTAFold, and two top models built by conventional homology (**Supp. Data 4**). Among them, only one of the two AF2-based models, GPR139-2717-0003, had considerable accuracy; the remaining models were not successful (correctness between 0.2% and 0.03%), highlighting the importance of strategic selection of diverse conformations. The most accurate model successfully reproduced the orientation of the ligand in the binding pocket, except the position of the chlorobenzene ring which is deviated from the experimental ligand (**Fig. 2D**). This group (UW-DiMiao) also submitted one of the most CMF-019-like predictions for the APJ/Cmpd-6 complex (APJ-2717-0005, correctness of 1.31%, MSC RMSD 2.64, 40% of correct contacts), similarly generated based based on an apo AF2 prediction (**Supp. Data 4**).

Similar to these examples, the authors of other top-ranking APJ and GPR139 models took advantage of AF2 for the prediction of receptor structures, with subsequent docking and pose selection for the small molecules in these models.

When comparing the best prediction accuracy achieved by the competitors for the two complexes, the maximum correctness for APJ/Cmpd6 (0.61%) is considerably lower than for GPR139/JNJ-63533054 (2.50%, **Supp.** Fig. 5A). However, the participants generated several APJ-Cmpd6 predictions very closely resembling the MCS of CMF-019 in PDB 8XZI (maximum correctness of 5%, **Supp.** Fig. 5A). This trend correlates with the availability of homologous structures in the PDB at the time of the assessment: two were available for APJ (PDB IDs 5VBL [23] and 6KNM [24]) and none for GPR139; the closest GPR139 homologs with available structures (CCR5, GPR52, NTR1, OPRK, CCKAR) had less than 30% sequence identity in the TM domain (**Fig. 1B**). This limitation could have affected the AF2 prediction accuracy for GPR139 since the model is trained on existing structures.

### Peptide targets: near-experimental accuracy for the core parts of neuropeptide Y (NPY1R) and neuromedin-25 (NMUR2), but not for dynorphin (OPRK)

**Fig. 3** summarizes the “correctness” (ligand RMSD vs % contacts) of the submitted OPRK/dynorphin, NPY1R-NPY, and NMUR2/NMU25 models when compared against our chosen “gold-standard” experimental structures, PDB 8F7W, 7X9A, and 7W55, respectively. The distribution of model correctness values when compared against other experimental structures is given in **Supp.** Fig. 5B.

**Figure 3.**
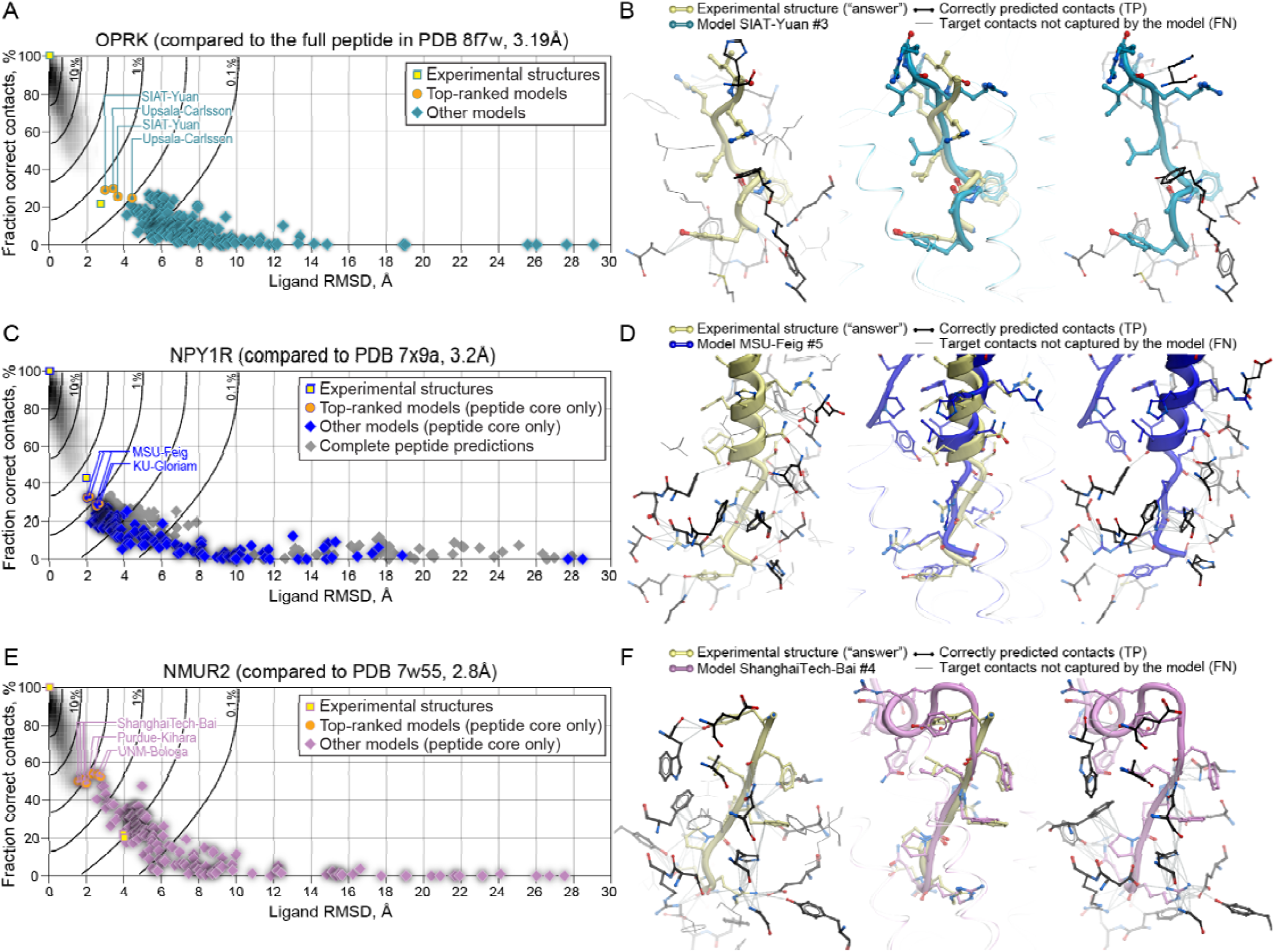
**Peptide targets: near-experimental accuracy for the core parts of neuropetide Y (NPY1R) and neuromedin-25 (NMUR2), but not for dynorphin (OPRK).** (**A**, **C**, **E**) Prediction of ligand pose (measured as heavy-atom RMSD) and ligand-pocket contacts by the OPRK/Dynorphin **(A)**, NPY1R/NPY **(C)**, and NMUR2/NMU25 **(E)** models when compared against OPRK/Dynorphin (PDB 8F7W, aa 1-8), NPY1R/NPY (PDB 7X9A, aa 21-36 ), and NMUR2/NMU25 (PDB 7W55, aa 18-25) Cryo-EM structures, respectivly. (**B**, **D**, **F**) Comparison of the ligand predictions in the highest-correctness submitted models to the experimental a swer for OPRK (**B**), NPY1R (**D**), and NMUR2 (**F**). Left panels illustrate the target ligand-pocket contacts that are correctly redicted (black sticks) or not captured (gray wires) by the model, middle panels the overlay of peptide poses following the optimal superimposition of receptor TM domains, and right panels the correctly predicted pocket contacts with the modeled ligands.

Top-ranking models for OPRK/dynorphin were generated by the groups **SIAT-Yuan** and **Uppsala-Carlsson**, and reached only 0.7% and 0.59% of experimental correctness, respectively (**Fig. 3A**): for OPRK-8407-0003, the heavy-atom RMSD for aa 1-8 was RMSD 2.96Å, with 28.55% correctly predicted contacts (**Fig. 3B**), and for OPRK-2736-0004, RMSD was 3.40Å, with 29.45% contacts. Both of these models were about as similar to PDB 8F7W as the other experimental structure of the same complex, PDB 7Y1F (RMSD 2.72Å, 21.71% contacts, **Fig. 3A**). Each of the groups submitted two models of comparable accuracy (**Fig. 3A**). Considering the structures of OPRK (bound to small molecule agonists and antagonists) and of closely homologous OPR[MD] (bound to morphinan and non-morphinan small molecules as well as peptides [PDB IDs 6DD[EF], 6PT2]) that were available in the PDB at the time of competition, we expected a large number of near-experimental accuracy OPRK/dynorphin complex models; however, none of the submitted models reached the correctness of 1%. The correctness, ligand RMSD, and fraction of correct contacts for the best-ranking model of OPRK (model 8321-0003) relative to PDB ID 7Y1F were 0.24%, 4.03Å, and 14.4%, respectively (**Supp.** Fig. 6A). When the analysis was focused on aa 1-5 of the dynorphin peptide only, (which effectively forms another OPRK agonist, Leu-enkephalin), the best models reached correctness of 2.88% (OPRK-8407-0003, RMSD 1.79Å, 47.3% contacts) and 1.06% (OPRK-8669-0003, RMSD 2.08Å, 28.34% contacts when compared against PDB 8F7W and PDB 7Y1F, respectively (**Supp.** Fig. 6B-C).

For the NPY1R/NPY complex, almost half of the submitted models have RMSD (for the heavy atoms of ligand ‘core’, aa 21-36) in the range of 2-4 Å **Fig. 3C**. For the best model, built by group **MSU-Feig**, the correctness reached 1.32% (NPY1R-3601-0005, RMSD 1.99Å, 32.12% correct contacts, **Fig. 3D**). The same model was also most similar to PDB 7VGX (**Supp.** Fig. 6D). It is worth mentioning that as many as 11 submitted models reached 1% or more of experimental accuracy when compared against PDB ID 7VGX (**Supp.** Fig. 6D).

Surprisingly, despite limited homology in the PDB at the time of the assessment (**Fig. 1B**), the highest accuracy among peptide targets was achieved for the NMUR2/NMU25 complex (**Fig. 3E**), relative to the resolved part (aa 18-25) of the peptide in our chosen “gold-standard” structure PDB 7W55 [95]. The closest model was generated by **ShanghaiTech-Bai** and had correctness of 3.91% (NMUR2-7750-0004, RMSD 1.5Å, 50.21% of correctly captured contacts, **Fig. 3F**); other accurate models were built by teams Purdue-Kihara and UNM-Bologa (**Fig. 3E**). By contrast to PDB 7W55, when the models were compared against PDB 7XK8 or its early refinement version (‘pre-7XK8’, **Supp. Data 2**), their correctness only reached 0.42%-0.48% at best (**Supp.** Fig. 6E-F).

The low success for OPRK/dynorphin (only 0.6-0.7% correctness when compared to the resolved part of the peptide in PDB 8F7W) may reflect the failure of AF2 on this complex. In fact, both groups that generated the most accurate predictions (**SIAT-Yuan** and **Uppsala-Carlsson**) built them using conventional homology modeling (MODELLER) and docking, and capitalized on the existing structures of OPRK with an epoxymorphinan agonist MP1104 (PDB 6B73 [47]) and of the homologous OPRM with DAMGO (PDB 6DDE [50]) (**Supp. Data 4**). **SIAT-Yuan** built 20,000 receptor models using multiple-template sequence alignment and build generation strategy, selected the best one by DOPE (Discrete Optimized Protein Energy) scoring, refined it with a 200 ns restrained all-atom MD simulation in GROMACS using CHARMM36m force field, to open up the binding pocket and thus enable peptide docking, and selected representative conformation from the clustered simulation frames. The initial docked peptide poses were generated by AutoDock CrankPep [106]. For complex model selection, they trained a Convolutional Neural Network (CNN) on all the existing GPCR-ligand structures with interaction distance, interaction types, and atom types as features; the obtained CNN was used as the final step to select the final models from the set of docked conformations (**Supp. Data 4**).

For their most accurate NPY1R/NPY complex model, NPY1R-3601-0005, **MSU-Feig** used a modified version of AlphaFold Multimer: the receptor was first modeled using the Multi-State GPCR modeling protocol [107] and then used as a template in AlphaFold Multimer, to bias the predictions towards the active state of the complex. The AlphaFold models with high confidence scores were then submitted by the group without any further refinements (**Supp. Data 4**).

The best ranking NMUR2/NMU25 model, NMUR2-7750-0004, was built by **ShanghaiTech-Bai** using AlphaFold2 Multimer, for co-folding of the receptor with the NMU25 peptide. The complexes were then optimized in the Prime module of Schrödinger suite 2021-3 followed by 100 ns MD simulations in a POPC membrane using S-OPLS force field. MD frames were clustered, the averaged conformations were further energetically refined and scored by MM-GBSA in Schrödinger. Model selection for the submission was guided by both MM-GBSA binding energies and visual inspection (**Supp. Data 4**).

To complement our assessment of receptor-peptide complexes using the ‘correctness’ metric, we also sought to evaluate them by the DockQ score [96]: a metric for evaluating the accuracy of protein-protein/protein-peptide predictions developed in the context of the CAPRI competition [98]. As expected, DockQ scores correlated well with our ‘correctness’ scores (**Supp.Fig. 7**), as the core equation of the DockQ metric similarly relies on LRMS (ligand RMSD) and Fnat (fraction of correct contacts). However, one of the main advantages of the ‘correctness’ metric is that besides ranking the models and assessing their accuracy, it puts them in the context of experimental high-resolution structures.

Altogether, the results of this section indicate that the participants of GPCR Dock 2021 were able to generate highly accurate predictions - approaching the accuracy of high-resolution X-ray structures -for NPY1R/NPY and especially for the resolved part of NMUR2/NMU25 (**Supp.** Fig. 5B). For OPRK/dynorphin, the correctness of best predictions was less than 1%; however, it reached 2.88% for the well-constrained aa 1-5 of the dynorphin peptide (**Supp.** Fig. 5B). In all cases, the best models were at least as close to the chosen answer as other experimental (cryo-EM) structures of the same complex, and in the case of NMUR2 they were considerably closer (**Fig. 3A, C, E**). The maximum prediction accuracy did not correlate with the availability of the homologous experimental structures (**Fig. 1B**) as the most structurally characterized receptor - OPRK - turned out to be the most challenging. Coincidentally, AlphaFold2-predicted models for the OPRK/dynorphin complex were at the time, and remain to this day, quite distinct from the experimental structure, explaining why the best predictions were derived using conventional homology modeling and docking. Conversely, the highest accuracy predictions were submitted for NMUR2 which did not have any prior experimental structures, and they were generated via complex co-folding with AlphaFold2 Multimer with subsequent energy-guided model refinement and selection. For the NPY1R/NPY complex, the most accurate models were directly from AlphaFold2 and have not undergone any refinement; consistent with the success of off-the-shelf AlphaFolding on this complex, a large number of reasonably accurate models were submitted (**Supp.** Fig. 5B - note the bi-modal distribution of correctness for NPY1R/NPY complex models).

### Improved accuracy of receptor structure prediction loosely correlates with ligand prediction success

Because accurate modeling of ligand geometries and interactions required the prediction of induced conformational rearrangements in the receptors, we next assessed how well these rearrangements were predicted in the submitted models. **Fig. 4** illustrates the distribution of RMSDs of TM regions and extracellular loops (ECL1, ECL2, and ECL3) of the target receptors across the models. For the majority of models, the TM domain RMSD lies within 1-3 Å from the experimental conformation, for all targets; however, the ECL2 backbone RMSD values show significant variation ranging between 0.5Å and 16 Å.

**Figure 4.**
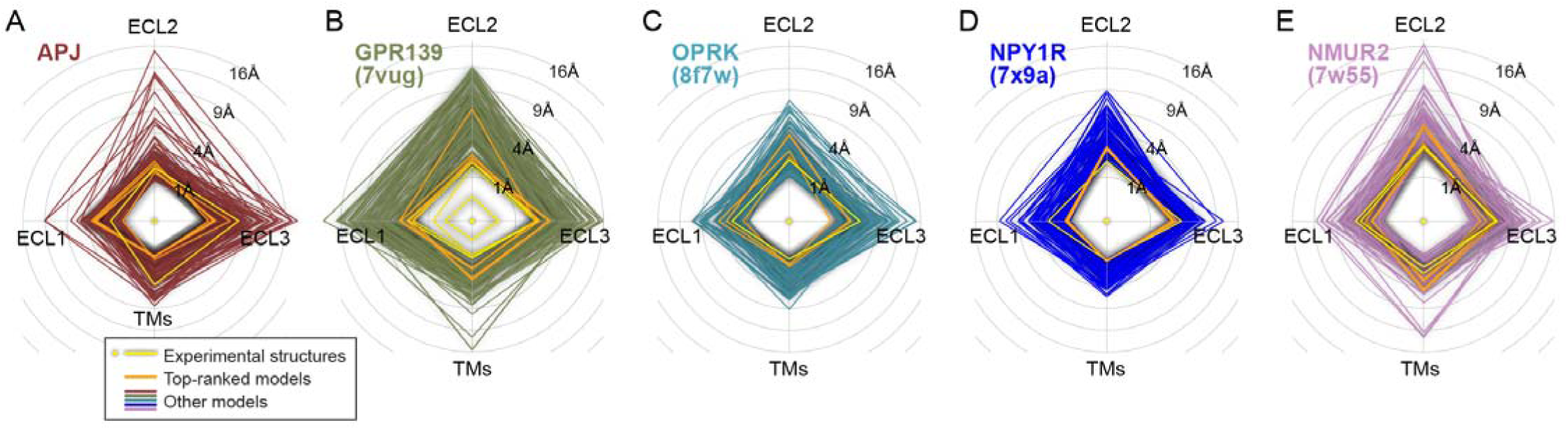
**Improved accuracy of receptor structure prediction loosely correlates with prediction success ligand** The backbone RMSD of model transmembrane domains (TMs) and extracellular loops (ECL1, ECL2, ECL3) from the selected “gold standard” experimental structures following the optimal superimposition of receptor TM domains. Other experimental structures and computational models are represented as quadrilaterals with a greater RMSD expressed as a larger radius on the corresponding axis of the radar plot.

For APJ, the majority of models (70%) had TM domain RMSD of less than 1Å relative to both the APJ/Cmpd6 complex X-ray and PDB 8XZI. However, ECL1 and ECL2 RMSD was within 1Å of the APJ/Cmpd6 complex X-ray for only 16.5% and 4.85% of the models, respectively, and ECL3 was not predicted within that accuracy by any of the models (**Fig. 4A**). Relative to PDB 8XZI, 21.4% of the models had ECL1 RMSD within 1Å, and none achieved this cutoff for ECL2 or ECL3 (**Supp.** Fig. 8A). The model with the most X-ray-like compound pose, **SDU-Yang** (model APJ-1912-0005), featured excellent ECL1 and ECL2 RMSDs of 0.51Å and 1.21Å, respectively, but 3Å RMSD for ECL3; whereas for other X-ray-like models, all three of ECLs had RMSDs of 1.5-2Å. Relative to PDB 8XZI, the ECL1 RMSD of the best models ranged from 0.57Å to 1.3Å, ECL2 RMSD from 1.4Å to 2Å (except APJ-2928-0004 with ECL2 RMSD of 3.98Å), and ECL3 RMSD from 1.1Å to 1.9Å (except APJ-8470-0005 with ECL3 RMSD of 3.26Å). Therefore, the most experiment-like ligand predictions were generally accurate on the receptor side. However, as expected, not all accurate receptor models translated into precise ligand predictions (**Fig. 4A, Supp.** Fig. 8A).

Consistent with the lack of homologous experimental structures in the PDB at the time of the assessment, GPR139 receptor models were generally less precise. Only ∼3% of submitted models had TM domain RMSD within 1Å of PDB 7VUG, and none had any of the three ECLs within this cutoff. However again, the loops were generally more accurate among the modes with the most precise ligand placement (**Fig. 4B, Supp.** Fig. 8B).

For OPRK, ∼1/3rd of the models had TM domain backbone RMSD within 1Å of PDB 8F7W. Two models had ECL1 RMSD none had ECL2/3 RMSD below 1Å, but many (56.57%, 38.29%, and 47.43%, respectively) had ECL1, 2, and 3 within 2Å of the experimental answer. Notably, the best peptide predictions are not most accurate in terms of the receptor TM domain or loops (**Fig. 4C**). The similarity of predicted models to PDB 7y1f is generally lower but the same trend holds (**Supp.** Fig. 8C).

For NPY1R, 17.42% of the submitted models had TM domain RMSD below 1Å. 26.45% of models had ECL1 RMSD below 1Å but none had ECL2 or ECL3 below not only 1Å but also 2Å (46.45% had ECL2 < 3Å and 59.35% had ECL3 < 3Å) (**Fig. 4D**). The models with the highest peptide placement accuracy are tightly aligned with each other in the TM/ECL RMSD space and close to the maximal TM domain and loop prediction accuracy (**Fig. 4D**). These trends also hold when the models are compared against PDB 7VGX (**Supp.** Fig. 8D) and may be reflective of the ordered helical nature of the peptide part that interacts with the loops.

Finally, for NMUR2, 12.5% of the models predicted TM domains and 4.6% predicted ECL1 with backbone RMSD < 1Å. No models had ECL2 RMSD of < 2Å but 15.79% had ECL3 RMSD within this cutoff (**Fig. 4D**). Unlike the NPY1R case, the most precise NMU peptide placement models had moderate ECL accuracy (e.g. 1.40Å, 3.20Å, and 1.76Å for ECL1, 2, and 3, respectively, in model NMUR2-7750-0004 by **ShanghaiTech-Bai, Fig. 4D**). However, the trend was restored when the models were compared to PDB 7XK8 (higher receptor accuracy is observed for most accurate peptide predictions, **Supp.** Fig. 8E).

Overall, the TM domain and loop predictions were tighter for the receptors that had experimental structures in the PDB at the time of the assessment (APJ, OPRK, and NPY1R) and more spread-out for the remaining two receptors (GPR139, NMUR2). Precise ligand placement generally required generally accurate prediction for the loops; however, this relationship was not strict and the reverse was not true (not all accurate receptor models featured precise ligand placement).

### AlphaFold2 played a vital role in GPCR Dock 2021 successes

According to the modeling methods submitted by GPCR Dock 2021 participants (methods were provided for 150 of 201 APJ/Cmpd6 models, 155 of 198 GPR139/JNJ models, 128 of 170 OPRK/dynorphin models, 119 of 155 NPY1R/NPY models, and 115 of 152 NMUR2/NMU25 models, **Supp. Table 4** and **Supp. Data 4**), most groups made use of the breakthrough AlphaFold2 structure prediction technology in their work. This was true for 40.7%, 68.4%, 54.7%, 82.4%, and 84.3% of the models submitted with accompanying methods for APJ, GPR139, OPRK, NPY1R, and NMUR2, respectively (**Table 3**). For small molecule targets (APJ and GPR139), the receptors could only be predicted in the *apo* form and required subsequent ligand docking. By contrast, for the peptide targets, the participants had two options: co-folding the receptor with the peptide or predicting the structure of the *apo* receptor with subsequent peptide docking. Interestingly, AlphaFold2 was extensively used even for the receptors that had experimental structures in the PDB at the time of the assessment (APJ, OPRK, NPY1R, **Table 3**).

For both small-molecule targets, the most accurate complexes were produced with the use of receptor models built by AlphaFold2 (**Fig. 5A**) and experimental structures (available for APJ) or conventional homology models. In fact for APJ, Cmpd6 docking to one of the existing PDB structures (reported as a method for 86 models) produced the best correctness of only 0.22% when compared to the APJ/Cmpd6 X-ray (**Fig. 5A**) (1.3% when compared to PDB 8XZI, **Supp.** Fig. 9). This correctness was exceeded by a model using the GEnSeMBLE methodology [108] (maximum correctness 0.35% when compared to the APJ/Cmpd6 X-ray, **Fig. 5A**, but only 0.02% when compared to PDB 8XZI, **Supp.** Fig. 9). Only 11 APJ models were reported to use conventional homology, and their best correctness was very low (0.04% when compared to the APJ/Cmpd6 X-ray, **Fig. 5A**, 0.2% when compared to PDB 8XZI, **Supp.** Fig. 9). In the absence of experimental structures, the conventional homology modeling approach was more popular for the GPR139 assessment (44 models); the best correctness achieved by this approach was 0.82% (**Fig. 5A**). However, AlphaFold2-generated models seem to have performed best for both small-molecule targets (**Fig. 5A**). This said, the use of best practices in subsequent compound docking / pose generation and pose scoring / selection was just as important as having the correct conformation of the receptor, and so was an expert assessment of the complexes, for example to evaluate the pose consistency with SAR or to exclude poses with buried polar atoms that do not make hydrogen bonds (**Supp. Data 4**).

**Figure 5.**
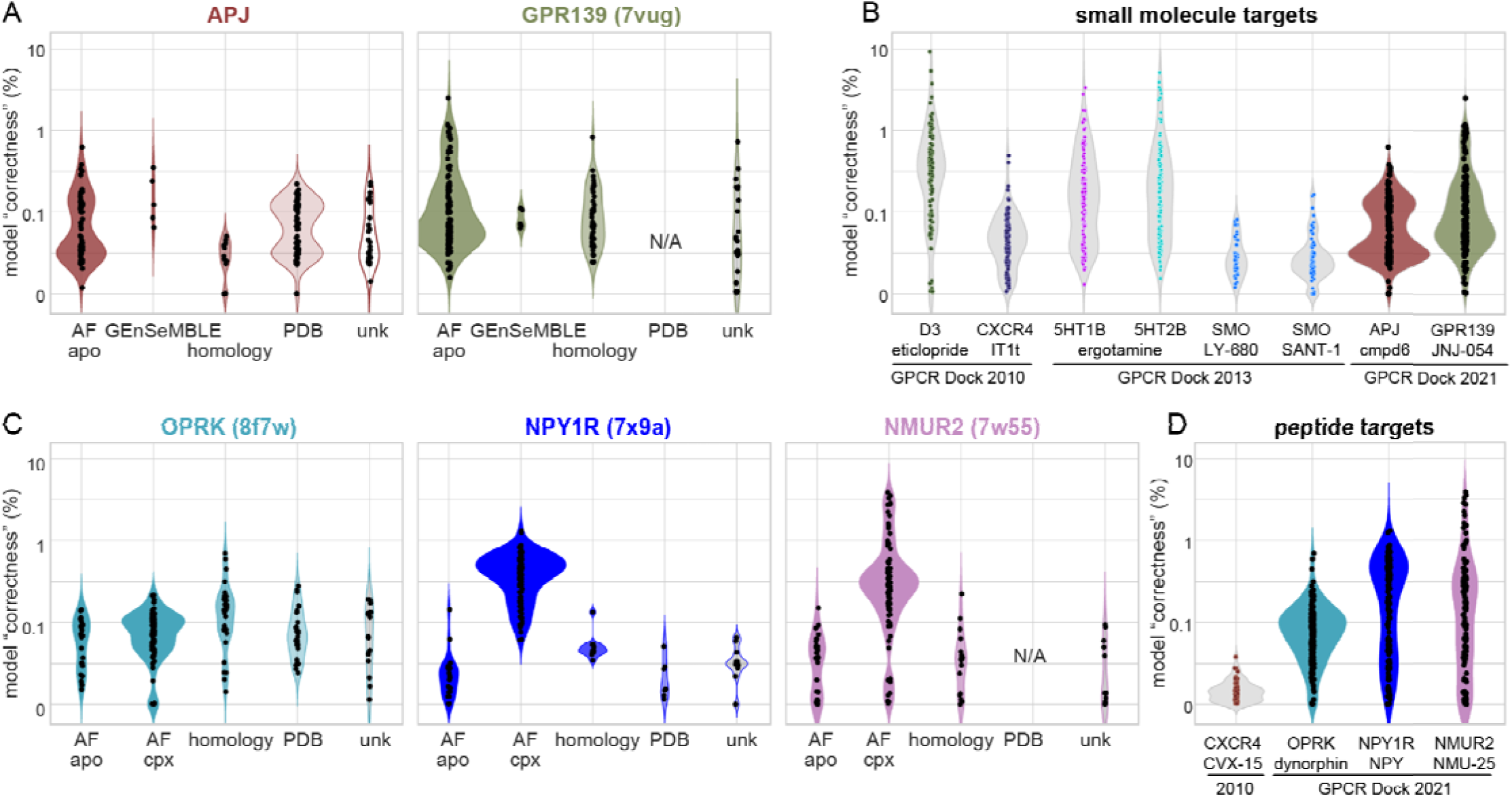
**AlphaFold2 played a vital role in GPCR Dock 2021 successes** (**A, C**) Distribution of model correctness, relative to the “gold standard” experimental structures, across different prediction methods employed by participants for small-molecule (**A**) and for peptide (**C**) target predictions. (**B, D**) Comparison of “correctness” distribution between GPCR Dock 2021 and previous GPCR Dock competitions (2010 and 2013) across small-molecule (**B**) and peptide (**D**) targets.

The use of the ‘correctness’ metric allowed us to directly compare the success levels in GPCR Dock 2021 to prior GPCR Dock assessments. Surprisingly, none of the small-molecule predictions reached the best accuracy observed in 2010 for the DRD3/eticlopride complex (**Fig. 5B**). The most CMF-019-like APJ prediction and the best GPR139/JNJ predictions had the level of similarity with the answer approaching that of best 2013 5HT1B/ergotamine and 5HT2B/ergotamine models in GPCR Dock 2013 (**Fig. 5B**).

For NPY1R and NMUR2, peptide docking to a homology model, to an apo AlphaFold2 model, or (in the case of NPY1R) to an experimental structure produced approximately the same levels of accuracy, well below 1% (**Fig. 5C**). The real improvement in both cases was achieved when AlphaFold was used in the Multimer mode for co-folding the receptor with the peptide (**Fig. 5C**). However, this approach did not work well for OPRK (**Fig. 5C**), possibly because the peptide was too short for building a reliable multiple sequence alignment (MSA). For the OPRK/dynorphin complex prediction, homology modeling appeared to have worked best, probably because of the available homologous templates (OPRM) with peptides. These examples thus illustrate the limitations of AlphaFold2 in application to not only small-molecule but also to peptide complexes. In any case, the prediction accuracy for all three peptide targets greatly exceeded that for the only peptide target in prior assessment: CXCR5/CVX15 in GPCR Dock 2010 (**Fig. 5D**).

It is worth pointing out that not all AF2 models had comparable accuracy. This emphasizes that even when the best tools are available, one often needs expert knowledge about (i) how to apply them and (ii) how to rank and score/rank the resulting models. In that sense, it would have been interesting to see the predictions of DeepMind scientists who were invited to participate but unfortunately declined. Finally, we note that several participating groups used AI beyond AlphaFold2, including the training and application of specialized neural networking for complex scoring and selection (e.g. **SIAT-Yuan** to derive their winning OPRK/dynorphin model, **Supp. Data 4**).

The release of the open-source code for AlphaFold3 [109,110] and other related diffusion-based models for protein co-folding with small molecules (NeuralPLexer [111], Chai-1 [112], Boltz-1 [113]) made it possible to assess this next generation methodology on the small-molecule complexes from GPCR Dock 2021. When challenged with predicting the APJ complex with Cmpd6, none of the four methods predicted the CMF-019-like geometry (similar to PDB 8xzi) and only Boltz-1 predicted the geometry similar to our X-ray structure of APJ/Cmpd6 (**Supp.** Fig. 10). For GPR139, GPR139/JNJ-63533054, only Boltz-1 predictions reached the accuracy of the best models in this assessment [12].

## Discussion

### Modeling challenges: do we still face the same challenges as in the past?

The 4^th^, 2021 GPCR Dock assessment was different from prior assessments in the series (2008 [15], 2010 [16], 2013 [17]) in several ways. First, it featured a larger number of targets: 5 target complexes of 5 receptors in GPCR Dock 2021 vs 1, 3, and 4 complexes of 1, 2, and 3 receptors, respectively, in GPCR Dock 2008, 2010, and 2013. Only two of five 2021 targets were complexes with small molecules; three were with peptides (vs only 1 peptide complex in prior assessments, that of CXCR4 with CVX15 in 2010 [16]). Peptide complexes have traditionally been considered most challenging and often intractable prediction targets, due to a large number of rotatable bonds that had to be sampled in docking, as well as the uncertainties of induced fit. For three GPCR Dock 2021 target receptors, experimental structures (in other complex compositions) existed in the PDB at the time of the assessment; however, for the remaining two receptors, only low-level homology structures were available. GPCRs exhibit significant conformational diversity in their orthosteric binding sites, consistent with complexities of molecular recognition within the GPCR family [6]. On the one hand, homologous structures may provide critical templates for modeling. On the other, by biasing modeling efforts, the available structures can have a detrimental effect on modeling success when the target complex is sufficiently conformationally distinct. In this context, GPCRs without existing structures presented a novel challenge where focus will be on developing and refining innovative modeling techniques. Without predefined structural templates, a wider range of computational approaches may be explored, potentially leading to more accurate predictions.

To counterbalance an overall increased difficulty of prediction tasks in GPCR Dock 2021, the state of the field was greatly advanced compared to earlier competitions. Many more GPCR structures from various families have been solved in both active and inactive states and available in the PDB, leading to much better general understanding of GPCR conformation and ligand interaction principles. Released only a few months before the assessment, AlphaFold2 became a go-to approach for capturing these principles in the models, with minimal human expert involvement. As a result, the challenges of predicting accurate receptor structures and complexes with larger peptides were largely solved. However, challenges for shorter peptides and small molecules remained, and induced fit was still a critical issue for these complexes.

### Experimental structures post-competition

Another key aspect of the 2021 assessment related to the multitude of experimental ‘answers’. In 2010 and 2013, such uncertainties were minimal. Even with the two chains of dopamine D3 receptor complex with eticlopride in 2010, and with the multiple structures of CXCR4 bound to IT1t, the differences were considered a consequence of experimental resolution, and the highest resolution structure was considered most trustworthy. The situation was drastically different in the present assessment, as experimental structures of identical (e.g. for OPRK [90,91]) or very similar (e.g. for NMUR2 [94,95]) composition complexes gave rise to quite distinct interaction geometries for the orthosteric ligand. Possible explanations for such variability include complex dynamics, resolution and experimental errors, or allosteric effects of the intracellular binding partners: for example, the presence vs absence of a nucleotide in the miniGαs/q [89] or the G protein-targeting antibody (as in the two NPY1R structures [92,93]) might have influenced the pose and interactions of the orthosteric agonist. Regardless of the reason, the multitude and diversity of structural forms of GPCR complexes of the same composition is increasingly being appreciated [114–116] and affects the definition of ‘success’ in structural modeling [117]. To account for the impact of changes in the experimental field, we assessed the submitted models in relation to *all* available experimental answers. At the same time, we chose to rely on the distribution of assessment parameters among the high-resolution X-ray structures of identical complexes as it existed in the PDB in 2013: this allowed us to quantify the improvements in model accuracy relative to prior assessments.

Among the multiple structures for each target complex, we chose one as the primary or ‘gold standard’ answer. For 4 out of 5 targets, this was the structure with a better resolution, even though for some targets, the differences were marginal (3.20 Å vs 3.22 Å for PDB 7VUG vs 7VUH for GPR139; 3.19 Å vs 3.20 Å for PDB 8F7W vs 7Y1F for OPRK). For NPY1R, the declared experimental resolution of the two structures, PDB 7X9A and 7VGX, was the same (3.20 Å) so we chose a more complete structure PDB 7X9A, as the ‘gold standard’.

It was surprising - and satisfying, - to see that best computational models generally more closely recapitulated higher resolution structures than their lower resolution alternatives. When compared to the higher-resolution ‘gold-standard’ structures of GPR139 (PDB 7VUG), OPRK (PDB 8F7W) and NMUR2 (PDB 7W55), the maximum ‘correctness’ achieved by computational models was 2.5%, 0.7% and 3.91%, whereas when compared to the alternative structures, the corresponding numbers were only 1.45%, 0.24% and 0.42%. The best models were approximately as similar (for GPR139 and OPRK) or even closer (for NPY1R) to the higher resolution structure than the alternative lower resolution structure(s). For NPY1R, the maximum model correctness was 1.32% when compared to the chosen ‘gold-standard’ structure (PDB 7X9A) but slightly higher than that for the alternative structure (PDB 7VGX, max correctness 1.44%); in both cases, the experimental structures were slightly more similar to each other than the models were to these structures. For APJ, due to the profound differences in the binding pose of congeners Cmpd6 and CMF-019 between the respective complex geometries in the 2.40 Å X-ray structure (this work) and 2.70 Å cryo-EM structure (PDB 8XZI, [87]), it is hard to conclusively relate the model success to structure resolution; however, we note that the best predictions were dramatically more similar to the latter (5% correctness) than to the former (0.61% correctness). Altogether, these observations suggest that for peptide complexes, predicted models can be as accurate as (or even more accurate than) low-resolution experimental structures. Similarly, for small-molecule complexes, best computationally generated models appear to be within the expected experimental error from at least one of the alternative complex geometries.

### Impact of AlphaFold on GPCR modeling

AlphaFold2 played a major role in the successes of GPCR Dock 2021 predictions, both for targets with (APJ, OPRK, NPY1R) and without (GPR139, NMUR2) close homologs in the PDB. This is consistent with other studies that demonstrated that AF2 prediction accuracy for GPCRs is only loosely related to the representation of homologous receptors in the training set [12,118]. The predictions for peptide complexes were greatly assisted by co-folding approaches. Interestingly, the OPRK/Dynorphin complex proved to be a challenging target for co-folding, likely due to lack of co-evolution of this short and extremely conserved peptide with the receptor. For small-molecule complexes, which required compound docking, two limitations of AF2 affected prediction accuracy: (i) inability to systematically and reproducibly predict receptor conformational ensembles and distinct functional states [107], and (ii) understandably lower prediction accuracy (and certainty) for receptor loops and flexible parts [22]. In this, the results of the GPCR Dock 2021 assessment for the small molecule complexes agree with other studies that compared AF2 to template based methods [13] or critically evaluated the ability of AF2 models to reproduce the poses of small-molecule compounds in docking [119] or to rank them in virtual screening [120].

### Looking forward - what is next

Since 2021, inspired by the raving success of AF2 for predicting structures of polypeptides, academia and industry raced to build and train a deep learning model that similarly predicts protein structures with non-peptidic ligands, including arbitrary small-molecule chemicals. These efforts culminated with the 2024 publication of AlphaFold3 (AF3, [109]) and other diffusion-based models for protein co-folding with small molecules: NeuralPLexer [111], Chai-1 [112], Boltz-1 [113], etc. Although the reported accuracy for these methods is excellent, they may still have limitations when it comes to predictions for complexes without homology and chemical similarity to the training set structures [121] or with multiple ‘correct’ complex geometries (**Supp.** Fig. 10 and [12]). Therefore, expert knowledge, incorporating information from compound SAR, and conventional model refinement remain important for generating accurate predictions even in the era of AlphaFold2/3.

In summary, our work demonstrated that the accuracy of computationally generated models of GPCR complexes with peptides can exceed that of low-resolution experimental structures. However, all the recent successes in AI structural modeling have not eliminated the need for experimental *high-resolution* GPCR structure determination [122], especially with small molecule chemicals - a much needed prerequisite for structure-based drug discovery, - and for conventional, physics-based and expert-guided computational modeling and refinement.

## Methods

### Data collection and filtering

The GPCR Dock 2021 participant registration and model submission system was implemented online using SmartSheets. Principal Investigators (PIs) were allowed to register for more than one group and submit models separately for each group as long as the methods used by the distinct groups were clearly different. Groups were invited to submit up to five models per target.

Originally, models were received from 49 groups. One group (group ID 8067) chose to withdraw at an early stage of the assessment. Upon receipt of their modeling methods, it appeared that groups HUST-Huang-3983 and HUST-Huang-8470 effectively operated as a single group; therefore, their submissions were merged and 5 models per target were selected for the assessment from the merged model set at random. By the same logic, submissions from group Tsinghua-Peng-6760 were merged with those from HeliXon-Luo-6867, and submissions from Galux-Won-4528 were merged with those from Seoul-Seok-8669. As a result, the number of participating distinct groups dropped to 45.

Group KIST-Park-8824 submitted 5 OPRK models of which 4 contained two peptides each, effectively making it 9 models. For this group and target, five models were selected for the assessment at random and the remaining four were disregarded.

Among the remaining models, 18 models were found to contain no ligand, no receptor, or severe steric conflicts, and were thus considered uninterpretable (**Supp. Table 2**). The final filtered set of models used for the assessment contained 206, 198, 175, 155, and 152 models for APJ, GPR139, OPRK, NPY1R, and NMUR2, respectively.

Model PDB files were standardized to ensure that each contains a single polypeptide chain representing the WT sequence of the target receptor. In several cases, this required deletion of fusion proteins, segments, or entire helices containing non-native aa sequence. The ligands were also standardized to ensure topological identity to the largest possible substructure of the target ligand. In selected cases, this involved deletion or renaming of ligand atoms provided by the participants. All non-receptor/non-ligand molecules were removed. The final set of cleaned-up, standardized, and filtered model PDB files is provided as **Supp. Data 3**.

### Analysis of receptor structure prediction accuracy

The protein molecule of each model was superimposed onto the backbone Cα, C, and N atoms of the TM domain (residue positions 1.30-1.60, 2.39-2.64, 3.22-3.54, 4.38-4.64, 5.35-5.64, 6.31-6.58, 7.32-7.55 in Ballesteros-Weinstein notation [123]) of the ‘answer’ structure using an adaptive algorithm attenuating the contribution of farthest deviating structure fragments [124,125]. The models were assessed by calculating the RMSDs of their TM Cα, C, and N atoms from the target structures. With the same TM superimposition, backbone RMSD values were calculated for the model extracellular segments defined as follows:

● **APJ:** NT = …-28, ECL1 = 89-99, ECL2 = 168-198, ECL3 = 271-288
● **GPR139:** NT = …-26, ECL1 = 87-98, ECL2 = 165-179, ECL3 = 251-265
● **OPRK:** NT = …-57, ECL1 = 119-128, ECL2 = 197-222, ECL3 = 297-310
● **NPY1R:** NT = …-38, ECL1 = 101-110, ECL2 = 176-207, ECL3 = 286-300
● **NMUR2:** NT = …-43, ECL1 = 106-116, ECL2 = 184-213, ECL3 = 291-307

Residues that were disordered in the ‘answer’ structures were omitted from the calculations.

### Analysis of ligand positions and conformations

Static (receptor-aligned) root-mean-square deviation (RMSD) was calculated between heavy (non-hydrogen) atoms of the ligands in each model and the corresponding ‘answer’ following the optimal superimposition of the TM helix backbones as described above. Chemical group symmetry was taken into account.

### Analysis of ligand-receptor atomic contacts similarity

Continuous distance-dependent contact ‘strengths’ were calculated for all receptor-ligand atom pairs as previously described [16,17,126]. Briefly, the ‘strength’ was assigned to 1 for *d* < *d_min_* = 3.23 Å, 0 for *d* > *d_max_* = 4.63 Å, and continuously decreased from 1 to 0 as a linear function of *d* for *d_min_* < *d* < *d_max_*. Ligand-pocket atom pairs separated by distance *d* < *d_min_* were assigned steric clash penalties calculated as *d_min_*/*d* – 1; if the sum of all penalties in the model exceeded 3,000, the complex model was classified as sterically impossible.

To assess the similarity of contact strengths between a model and an experimental ‘answer’, each containing an *n*-atom protein and an *m*-atom ligand, two matrices were computed: *C_A_* (*n*×*m*) for the ‘answer’ and *C_M_* (*n*×*m*) for the tested model (**Supp.** Fig. 3). A contact similarity matrix, *C_A_*_⋂*M*_ (*n*×*m*), was then generated by using the formula *C_A_*_⋂*M*_[*i*,*j*] = *min*(*C_A_*[*i*,*j*], *C_M_*[*i*,*j*]). Model contact recall and precision were quantified as *recall* = |*C_A_*_⋂*M*_| / |*C_A_*| and *precision* = |*C_A_*_⋂*M*_| / |*C_M_*|, respectively, recall and precision were averaged to produce contact prediction accuracy. To accommodate ligand symmetry, the enumeration of topologically equivalent atom permutations within the ligand was executed, while protein side-chain symmetry was factored in by treating symmetric atoms as indistinguishable [16,17].

### Quantification of model ‘correctness’ in the context of high-resolution experimental structure pairs

To estimate the ‘correctness’ of each model in comparison with the selected experimental answer, its ligand RMSD and contact prediction accuracy were interpolated in the context of the expected ligand RMSD and contact variation among high-resolution experimental structures of protein-ligand complexes of identical composition. The background set (34,317 pairs of PDB complexes representing 1390 proteins) and the equation for this calculation were exactly taken from [17] to ensure that the results are directly comparable to prior assessments.

### End-to-end modeling of APJ/Cmpd6 complexes

Molecular models of the APJ complex with Cmpd6 were predicted using NeuralPLexer [111], Chai1 [112], Boltz-1 [113], and AlphaFold3 [109]. For NeuralPLexer, the published GutHub version 1.0 [127] and model checkpoints from [128] were used; 16 predictions were generated with --chunk-size=2, --num-steps=40, and -- sampler=langevin_simulated_annealing, on an NVIDIA GeForce RTX 3060 Ti GPU. For Chai-1, the server implementation was used ([129], accessed Nov 13, 2024); 10 predictions were generated by using 2 seeds and 5 models per seed. For Boltz-1, the published GitHub version [130] was used, with the default Boltz-1 model checkpoint, 3 recycling steps, 200 sampling steps, default --step_scale (1.638), and --use_msa_server (MMSeqs2); 15 predictions were generated by using 3 seeds and 5 models per seed on an NVIDIA GeForce RTX 3090 GPU. For AlphaFold3, the published GitHub version 3.0.1 was used with default parameters; the database preset was configured to the full database mode and multiple sequence alignments were constructed using Jackhmmer version 3.3; 15 predictions were generated by using 3 seeds and 5 models per seed on an NVIDIA GeForce RTX 3090 GPU.

### Plotting and molecular visualizations

Molecular graphics were created in PyMol (Schrodinger Inc., **Fig. 1C-I**) or ICM 3.9-3b (Molsoft LLC, San Diego, CA, all other molecular graphics). Bar graphs and scatter plots were created in ICM (**Fig. 1B**, **Fig. 2A-B**, **Fig. 3A, C, E**, and corresponding supplemental figures), python/matplotlib (**Fig. 4**, **Supp.** Fig. 2, **Supp.** Fig. 3, **Supp.** Fig. 7, **Supp.** Fig. 8), or R/ggplot2 (**Fig. 5**, **Supp.** Fig. 5, **Supp.** Fig. 9).

## Acknowledgements

The authors are grateful to Dr. Mike Hanson for help with the model submission system, to Drs. Fei Xu, Zhi-Jie Liu, Tian Hua, Beili Wu, and Qiang Zhao (iHuman institute at ShanghaiTech University, Shanghai, China) for providing the coordinates of the target complex structures ahead of publication, and to the members of the Kufareva group (UC San Diego) for valuable discussions and critical review of the work. This study was partially supported by NIH grants R21 AI149369, R21 AI156662, R01 AI161880, and R01 GM136202 (to I.K.).

## Author contributions

The participants of GPCR Dock 2021 (**Supp. Table 1**) provided the computational models and the description of corresponding modeling methods. R.C., Y. W., and I.K. performed model analysis. R.C.S. and S.Z. coordinated experimental structure access and model collection. I.K. coordinated model analysis. R.C. and I.K wrote the manuscript with contributions from all authors.

## Data availability statement

All data in support of the manuscript findings is included in the manuscript and its supplementary materials. Except for APJ21 (the X-ray structure of the APJ/Cmpd-6 complex, **Table 1** and **Supp. Data 1**) and pre-7XK8 (the preliminary refinement of the NMUR2/Neuromedin-U-25 complex structure, **Table 1** and **Supp. Data 2**), the structures of the experimental ‘answers’ are available in the PDB under accession codes listed in **Table 1**).

