## Supplemental Materials for "The 4^th^ GPCR Dock: assessment of blind predictions for GPCR-ligand complexes in the era of AlphaFold"

### Supplemental Figures and Legends

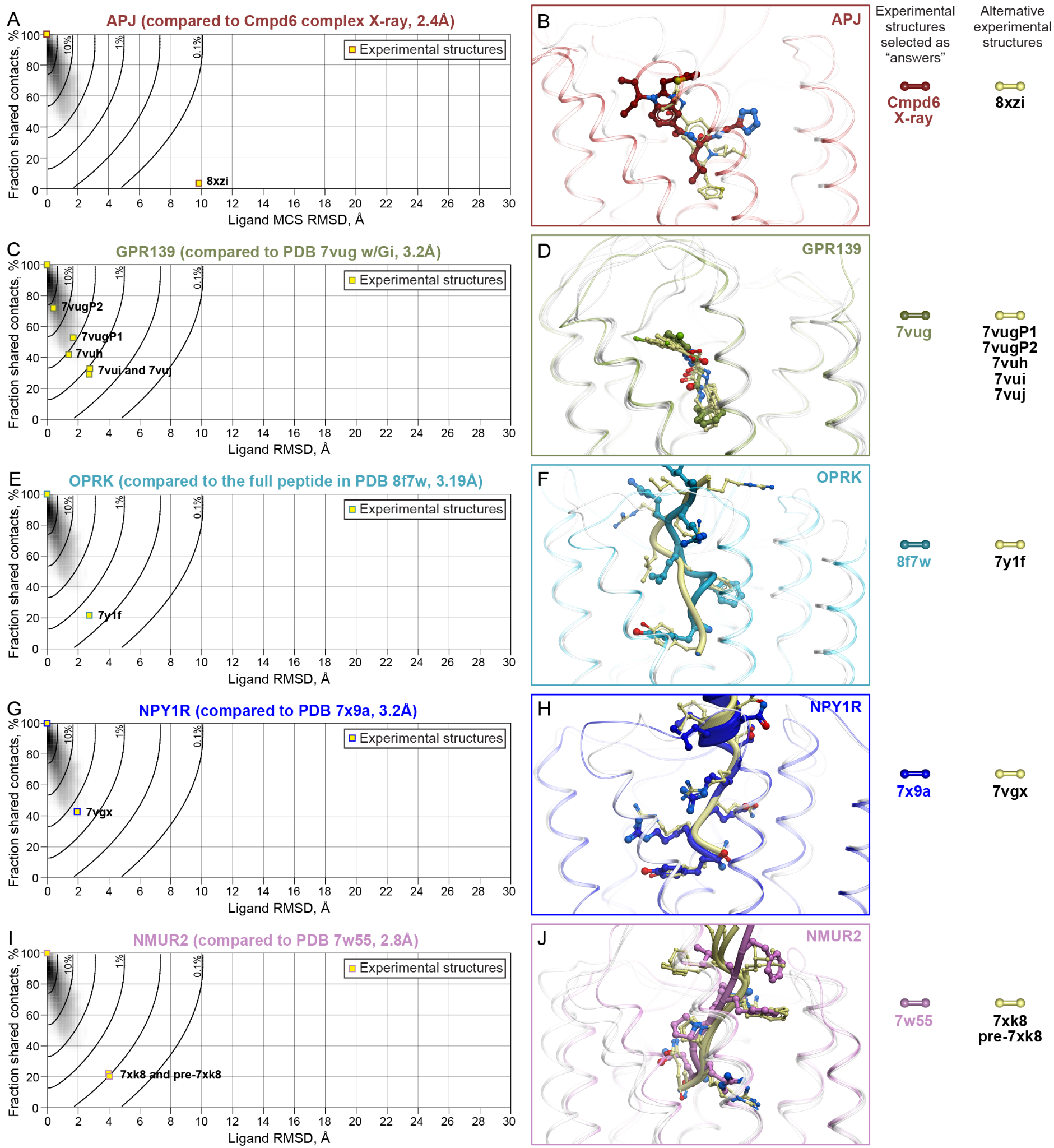

#### Supplementary Figure 1. Alternative experimental structures are available for 5/5 GPCR Dock 2021 targets, with lots of conformational variation among them.

(**A, C, E, G, I**) Heavy-atom RMSD and shared ligand-pocket contacts between the experimental structure selected as the “gold standard” answer and other experimental structures of the same (or, in the case of APJ, closely related) complex are shown on a scatter plot for APJ (**A**), GPR139 (**C**), OPRK (**E**), NPY1R (**G**), and NMUR2 (**I**).

(**B**, **D**, **F**, **H**, **J**) The ligands in the “gold standard” answer structures (colored ribbons and sticks) in comparison with the same ligands in other experimental structures of the same (or, in the case of APJ, closely related) complex (pale yellow ribbons and sticks) following optimal superimposition of the receptor TM domains.

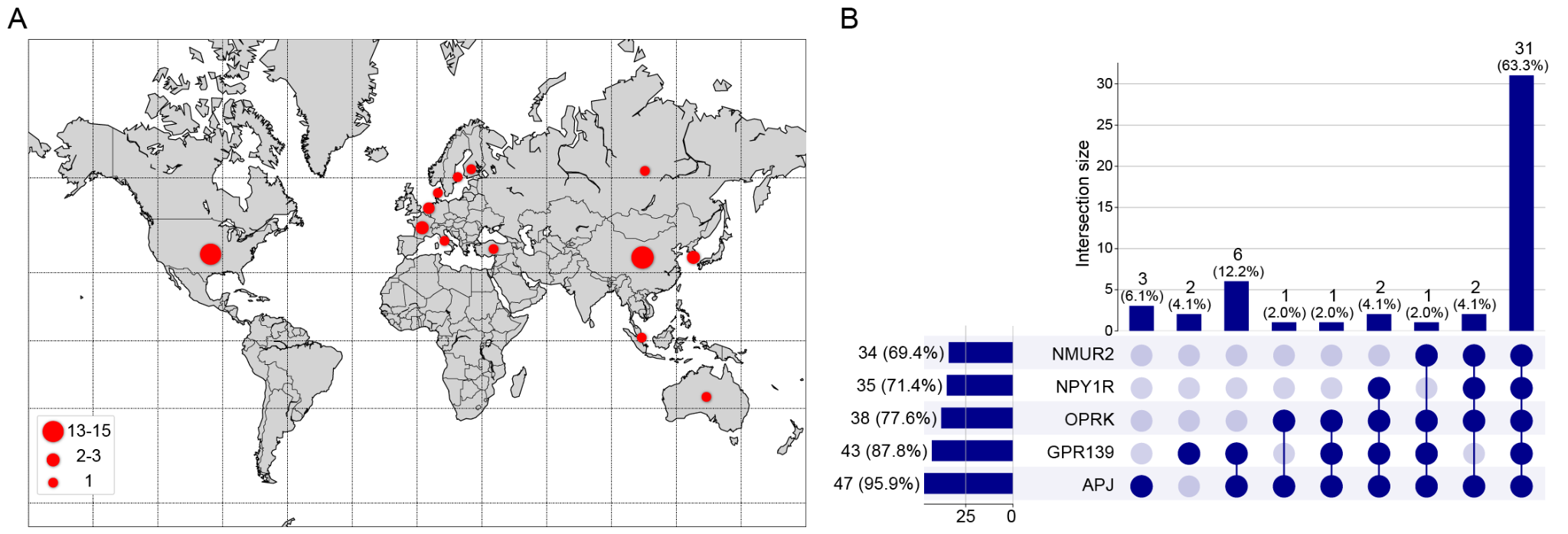

#### Supplementary Figure 2. The geography and statistics of GPCR Dock 2021 participants.

(**A**) The geographical distribution of participating groups and institutions.

(**B**) The completeness of model submissions for the five targets.

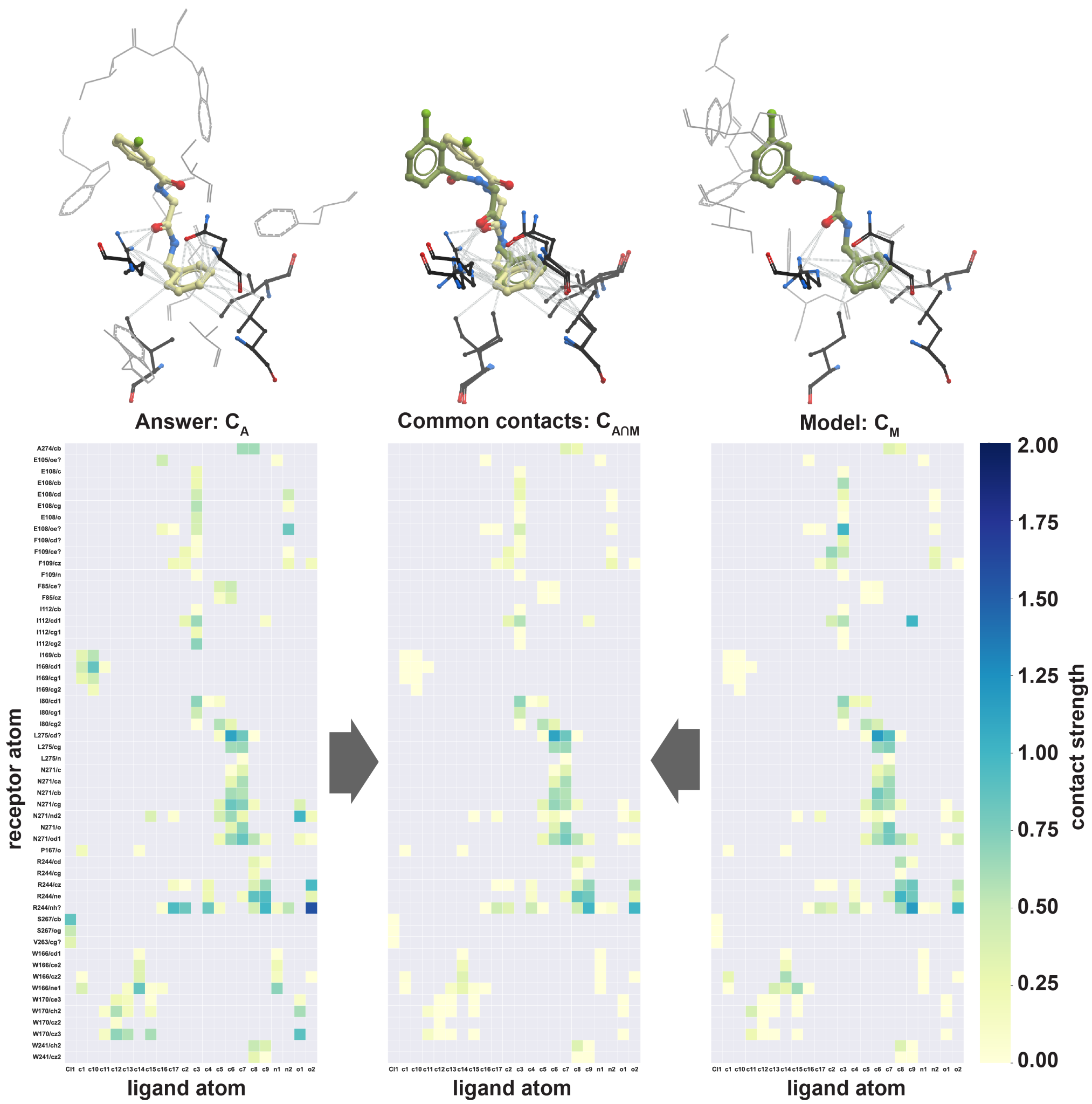

#### Supplementary Figure 3. A schematic illustrating the calculation of ligand-receptor contact prediction accuracy.

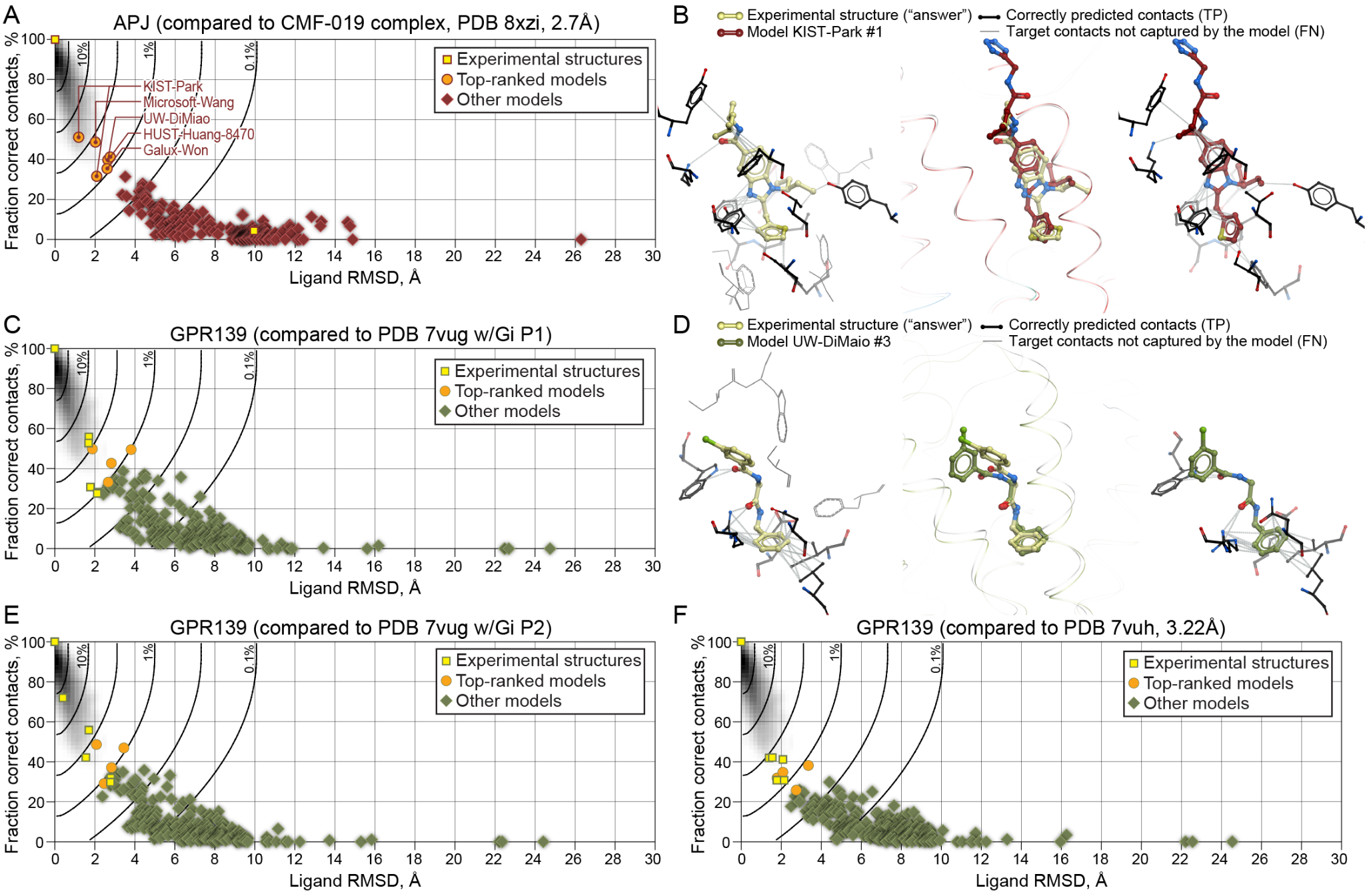

#### Supplementary Figure 4. Small molecule complex model comparisons to alternative answers.

(**A**) Prediction of ligand pose (measured as heavy-atom RMSD) and ligand-pocket contacts by the APJ/Cmpd6 models when compared against PDB XZI (APJ/CMF-019).

(**C, E-F**) Prediction of ligand pose and ligand-pocket contacts by the GPR139/JNJ complex models when compared against PDB 7VUG-P1 (**C**), 7VUG-P2 (**E**), 7VUH (**F**).

(**B, D**) Comparison of most CMF-019-like APJ/Cmpd6 prediction to PDB XZI (**B**) and of the most 7VUG-P1-like GPR139/JNJ prediction to PDB 7VUG-P1 (**D**). Left panels illustrate the target ligand-pocket contacts that are correctly predicted (black sticks) or not captured (gray wires) by the model, middle panels the overlay of compound poses following the optimal superimposition of receptor TM domains, and right panels the correctly predicted pocket contacts with the modeled ligands.

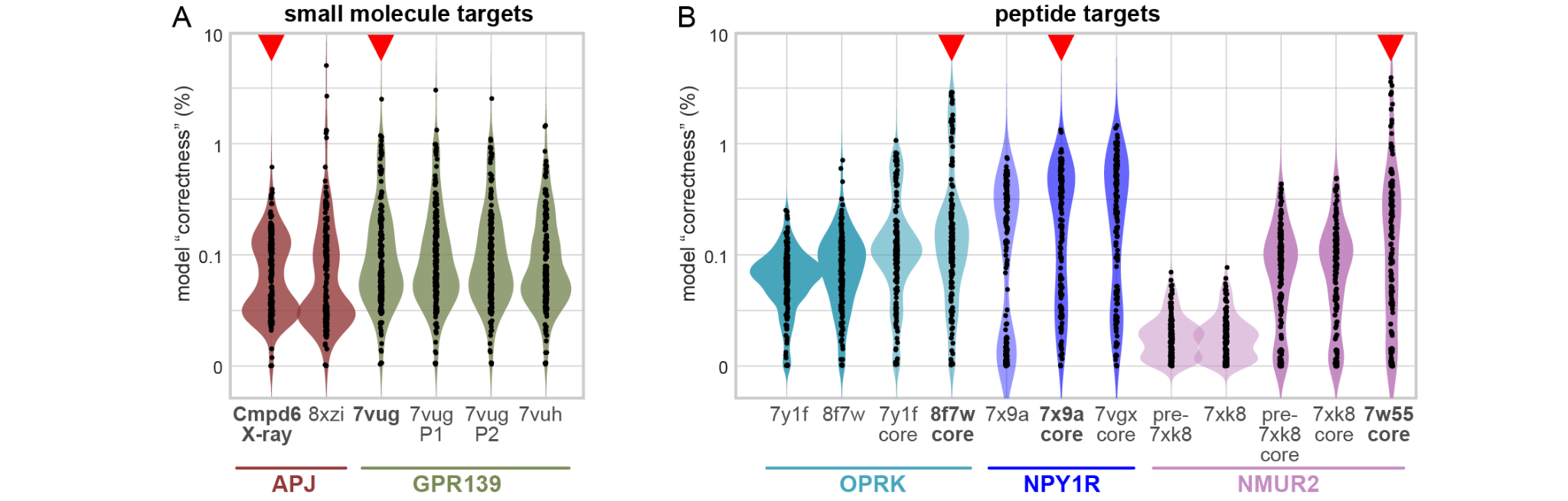

#### Supplementary Figure 5. Statistics of small-molecule (A) and peptide (B) complex model comparisons to alternative answers.

Distribution of model correctness relative to the selected “gold standard” experimental structures (red triangles) and other experimental structures of the same (or, in the case of APJ, closely related) complex for small-molecule (**A**) and for peptide (**B**) target predictions. For peptide targets, the correctness distributions when considering only the peptide ‘core’ vs the full length peptide are also shown.

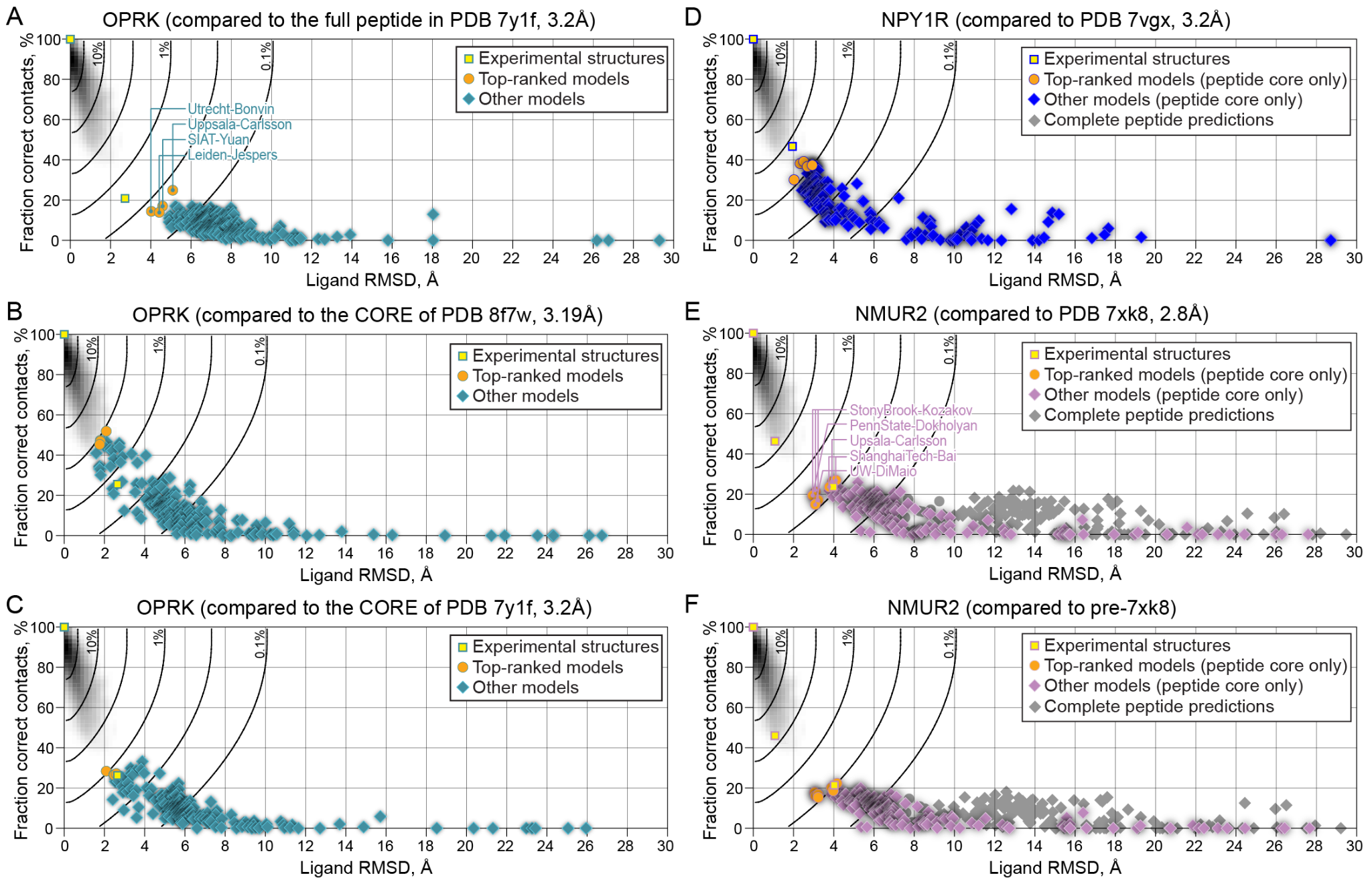

#### Supplementary Figure 6. Peptide complex model comparisons to alternative answers.

(**A-C**) Prediction of peptide pose (measured as heavy-atom RMSD) and peptide-pocket contacts by the OPRK/dynorphin complex models when compared against the full peptide in PDB 7Y1F (**A**), aa 1-5 in PDB 8F7W (**B**), or aa 1-5 in PDB 7Y1F (**C**).

(**D**) Prediction of peptide pose and peptide-pocket contacts by the NPY1R/NPY complex models when compared against PDB 7VGX.

(**E-F**) Prediction of peptide pose and peptide-pocket contacts by the NMUR2/NMU25 complex models when compared against PDB 7XK8 (**E**) or its early refinement (**F**).

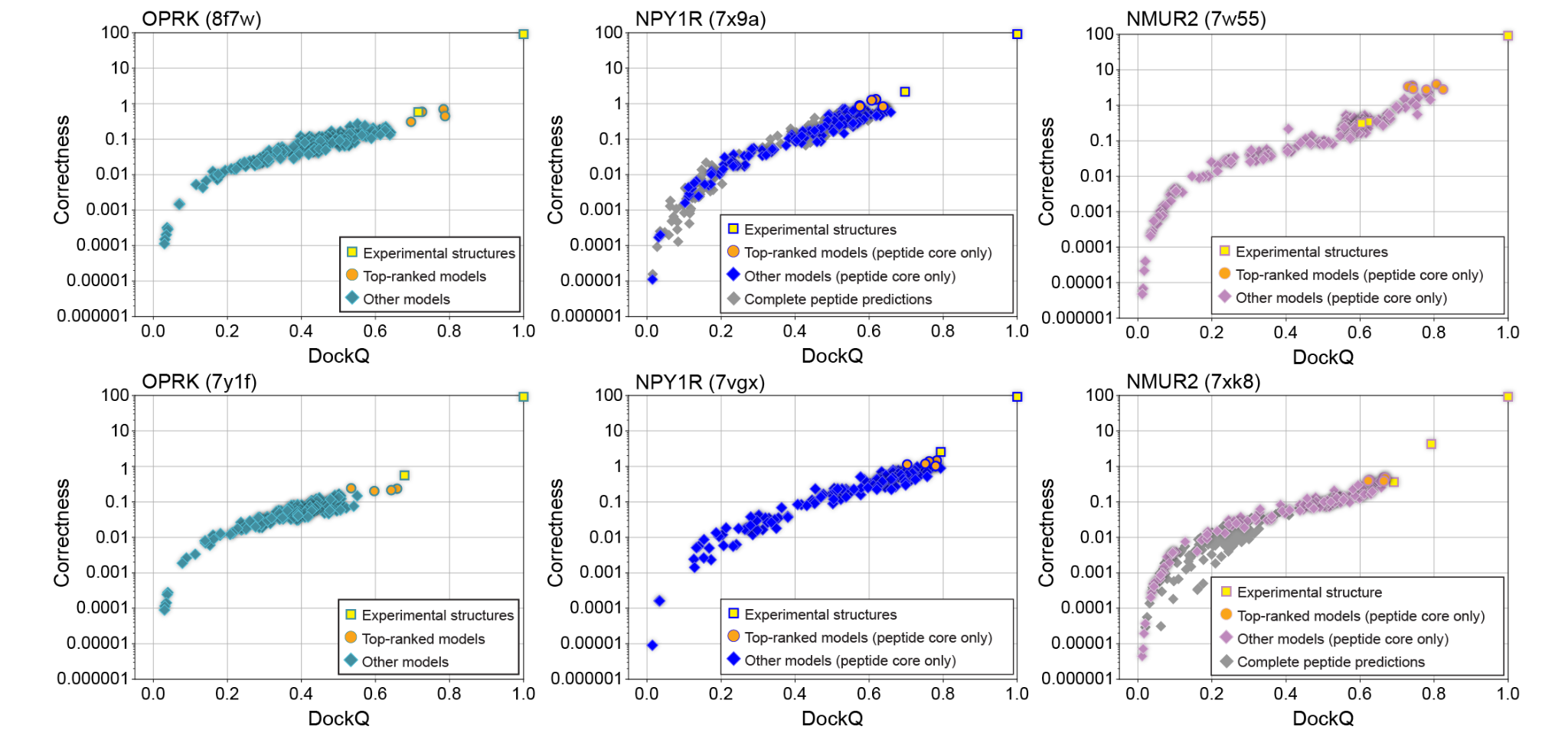

#### Supplementary Figure 7. Model “correctness” closely correlates with DockQ score.

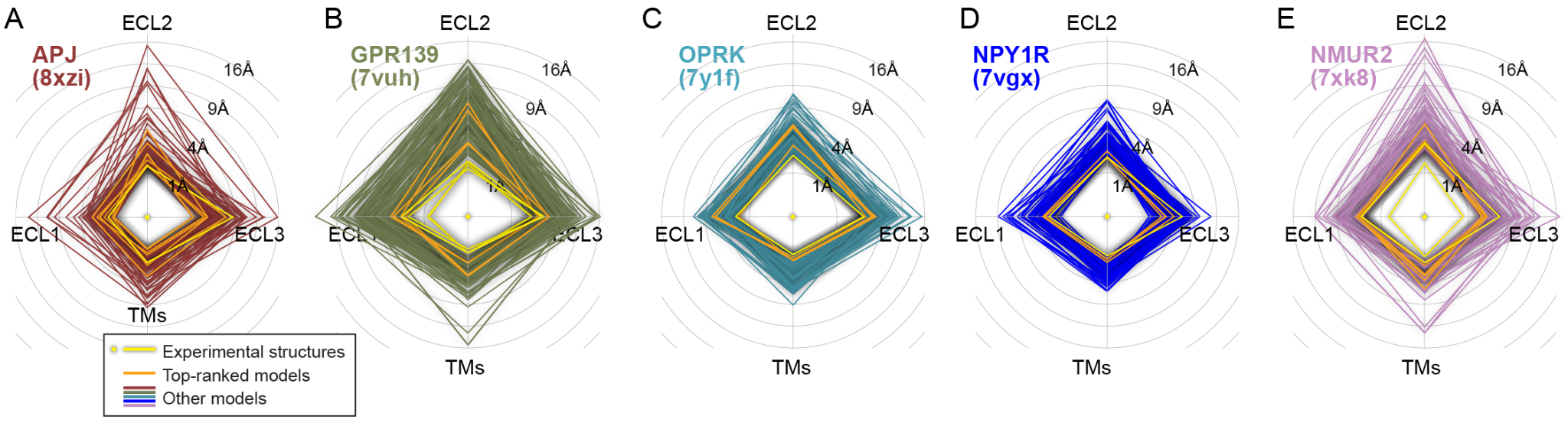

#### Supplementary Figure 8. Accuracy of receptor structure prediction when compared against the alternative experimental answers for the GPCR Dock 2021 targets.

The backbone RMSD of model transmembrane domains (TMs) and extracellular loops (ECL1, ECL2, ECL3) from the alternative structures of the GPCR Dock 2021 target complexes following the optimal superimposition of receptor TM domains. ‘Gold standard’ and other experimental structures and computational models are represented as quadrilaterals with a greater RMSD expressed as a larger radius on the corresponding axis of the radar plot.

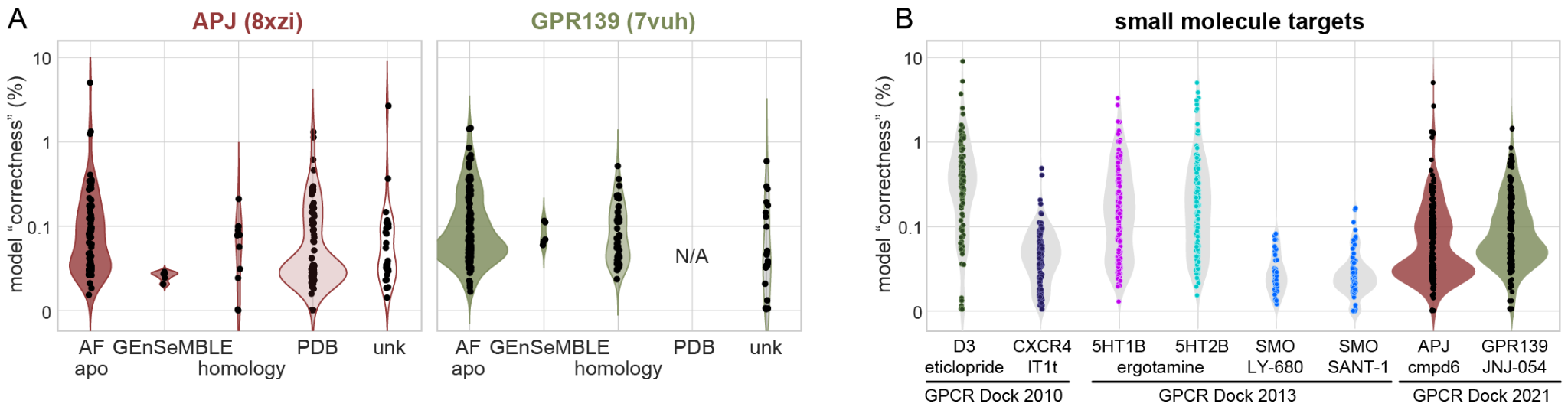

#### Supplementary Figure 9. Model correctness when compared to alternative experimental answers for the GPCR Dock 2021 small-molecule targets.

(**A**) Distribution of model correctness, relative to the alternative experimental structures, across different prediction methods employed by the participants for the small-molecule target predictions.

(**C**) Comparison of “correctness” distribution between GPCR Dock 2021 and previous GPCR Dock competitions (2010 and 2013) across small-molecule targets. GPCR Dock 2021 correctness is assessed relative to the alternative experimental structures.

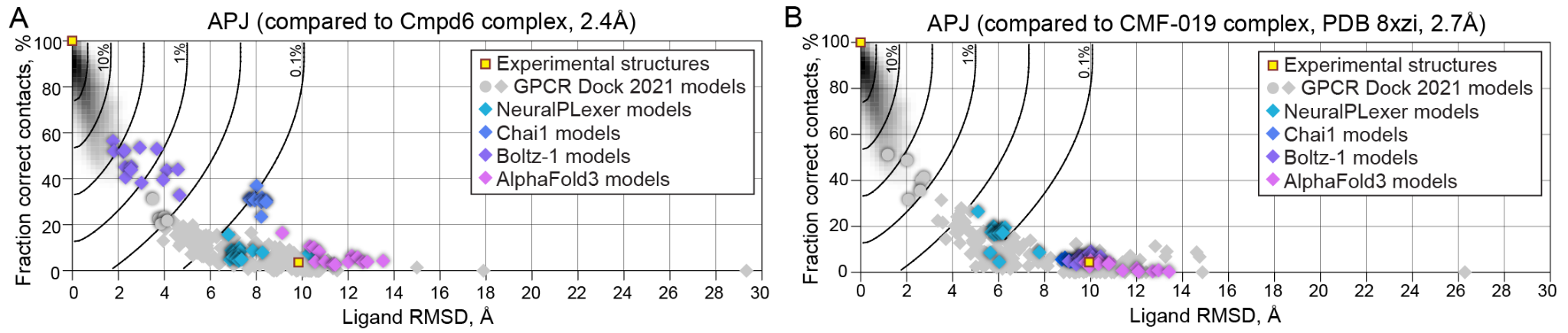

#### Supplementary Figure 10. APJ/Cmpd6 complex prediction accuracy by modern end-to-end AI methods.

(**A-B**) Prediction of ligand pose and ligand-pocket contacts by the APJ/Cmpd6 models constructed using NeuralPLexer, Chai1, Boltz-1, and AlphaFold3, and compared against APJ/Cmpd6 X-ray structure (**A**) or PDB XZI (APJ/CMF-019, **B**). The shaded background represents the distribution of the plot parameters observed across high-resolution X-ray structures in the PDB as in **Figs. 2** and **3**. Solid black curves represent isolines of the model “correctness” used for model scoring and ranking.

### Supplemental Tables

#### Supplemental Table 1.

45 groups (defined as institution/PI-associated teams with non-redundant modeling methodologies) that participated in GPCR Dock 2021

#### Supplemental Table 2.

Uninterpretable models

#### Supplemental Table 3.

Models and their corresponding correctness (for both gold standard and other answer)

#### Supplemental Table 4.

Summary of participants' modeling methods for assessment targets

### Supplemental Data

#### Supplemental Data 1.

The coordinates of APJ21 (the X-ray structure of the APJ/Cmpd-6 complex)

#### Supplemental Data 2.

The coordinates of pre-7XK8 (the preliminary refinement of the NMUR2/Neuromedin-U-25 complex structure)

#### Supplemental Data 3.

PDB coordinates of all models and all structures

#### Supplemental Data 4.

Modeling methods employed by the participants of GPCR Dock 2021
