## Supplementary material for "The 4^th^ GPCR Dock: assessment of blind predictions for GPCR-ligand complexes in the era of AlphaFold": Supp. Data 4

### GPCRdock-136: Methods

Liliana Halip, Department of Computational Chemistry, "Coriolan Dragulescu'' Institute of Chemistry, 24 Mihai Viteazu Blvd, Timişoara, Timiş 300223, România

Cristian G. Bologa, Translational Informatics Division, University of New Mexico School of Medicine, MSC09 5025, 700 Camino de Salud NE, Albuquerque, New Mexico 87131, USA.

**Building the initial models of receptors or complexes**

AlphaFold [1,2] version 2.1.1, using the default parameters and Amber relaxation was used to generate five models for each of the five receptors included in the competition. The models were generated using: APJ - the full sequence of the receptor; GPR139 - the sequence of the receptor (without the G-proteins and nanobody) in complex with an agonist peptide [3] containing the first 16 residues from the alpha-adrenocorticotropic hormone (ACTH); KOR - the truncated sequence of the receptor without the first 156 amino acid residues, which do not belong to the Kappa opioid receptor and were probably added for thermostabilization reasons only, and the sequence for the peptide ligand (Dynorphin), while ignoring the G-proteins and antibody sequences; NPY1R and NMUR2 – the sequence of the receptors in complex with their peptide ligand (NPY and respectively NMU25) and the alpha, beta and gamma G proteins. For the APJ and GPR139 receptors only the top scoring AlphaFold model was selected for docking of the small molecule ligand, while for KOR, NPY1R, and NMUR2 all five models were included in in the final submission.

**Small molecule docking**

OpenEye OMEGA [4,5] with default parameters, except for the option "-max_confs=5000" (APJ) and "-max_confs=2000" (GPR139) was used to generate 3050(APJ) and respectively 1301(GPR139) conformers for the small molecule ligand. For APJ The XRay structure of the apelin receptor in complex with an agonist peptide [6] (PDB code: 5VBL) was overlaid on top of the AlphaFold generated model, and the last 5 residues of the agonist peptide from the crystal structure were used as a template for docking the small molecule ligand using Openeye HYBRYD [7,8], for generating the models #1, #3, and #5. AutoDock[9] version 4.2.6 was used for docking the ligand into the AlphaFold model of the receptor and for generating the complex models #2 and #4.

**Model refinement**

The only minor "refinements" done were for NPY1R and NMUR2 and consisted in 1. changing the terminal COOH group to CONH2 in the peptide, and 2. for NPY1R model #5, a different rotamer of the NPY C-terminal tyrosine (Tyr 36) was selected.

**Criteria for selecting 5 best models per target**

We used a combination of docking scores and visual inspection for selecting and ranking the top models for APJ and GPR139. For KOR, NPY1R, and NMUR2, all five models generated by AlphaFold were submitted either completely unchanged (KOR) or after the minor changes described above (NPY1R, NMUR2).

**Target challenges**

While we were initially successful in receptor model generation for APJ and NMUR2 using the cloud version of AlphaFold2 Google colab [10], the notebook would run out of time or memory with the longer sequences of the other receptors, which required us to install and run AlphaFold locally on a GPU node at the University of New Mexico Center for Center for Advanced Research Computing. Also some of the receptors required removal of some artificial sequences inserted for termostabilization. As AlphaFold by default generates inactive (apo or antagonist bound) GPCR models, we did not use custom templates or multiple sequence alignments in order to force AlphaFold to generate agonist bound active states, but we plan to use this approach described in the AlphaFold Multistate paper [11] in the future.

**The role of experimental information and human expertise in generation and selection of your models**

When used appropriately, previous human experience can enhance and help computer generated models. However, human bias can also have a detrimental effect in objectively selecting the best in silico results. We tried to employ previous knowledge in the case of GPR139, for which agonist peptides were disclosed in a previously published paper. We first generated a receptor-peptide complex, and then used specific residues from the peptide to guide docking of the small molecule ligand. This approach however provided worse results than just simple docking of the ligand into the receptor.

[10] AlphaFold Colab, https://colab.research.google.com/github/deepmind/alphafold/blob/main/notebooks/AlphaFold.ipynb

[11] Heo, L., Feig, M., Multi-State Modeling of G-protein Coupled Receptors at Experimental Accuracy,

bioRxiv 2021.11.26.470086

**Group:** Xhaard-Helsinki
**ID:** GPCRdock-278

**Authors:** Henri Xhaard, Evgeni Grazhdankin, Loic Dreano

#### Target – APJ

**Strategy and hypotheses –** We used "classical" molecular modeling and docking using Modeller v10.0 (Sali & Blundell. J. Mol. Biol. 234, 779-815, 1993) together with the Schrödinger package (Schrödinger Release 2019-3: Maestro, Schrödinger, LLC, New York, NY, 2019). No AI was used to create the models.

Main hypotheses for model building and/or selection:

- Ligand occupy the same pocket as 5VBL's ligand (composed of ligand PDB codes 200, ALC, HRG, NLE).
- Key interactions with the cocrystallized ligand are likely similar to those seen in 5VBL (Figure 1)
- Ligand's tetrazole structurally mimics 5VBL ligand's carboxylate, forming an ionic interaction with K268.
- There are multiple possibilities for the structural superimposition of (either or both) of the GPCRdock 2021 ligand's two carbonyl groups vs (either or both) 5VBL's ligand's two carbonyl groups. These are indicated by arrows on Figure 2. Three possible hypotheses were devised, which correspond in the submitted models to: model 1, models 2 and 3, and models 4 and 5.
- Models 1-3 lead to a good overlap between GPCRdock ligand dimethyl group with 5VBL's peptidic ligand chloroaryl group.
- Models 4 and 5 explore the possibility that the ligand binds closer to the extracellular surface in comparison to Models 1-3.

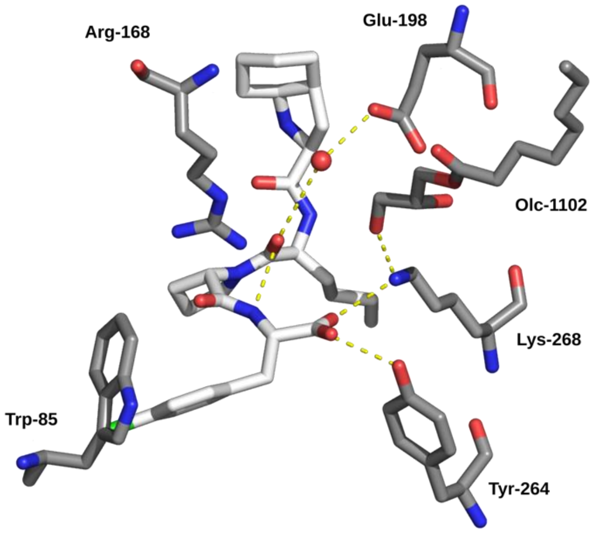

***Figure 1** – Interaction network bridging selected amino acids and water molecules of the APJ receptor (grey) in complex with peptidomimetic ligand (white) (PDB code 5VBL).*

**Receptor modeling –** The crystal structure 5VBL was used as a starting poin and corrected for missing atoms and mutated amino acids.

**Construction of the complexes –** To realize the hypotheses, we used Schrödinger's Glide SP and XP docking suite. Crystallographic waters and other cocrystallized ligands were removed prior to docking, with the exception of OLC-1102 that was thought to interact with bound ligand.

- Models 1 and 3 were created right away by the standard settings (Glide XP docking). The docking run generated a pool of 20 poses.
- Model 2was generated using Glide_XP. The docking run generated a pool of 5 poses.
- Model 4 was generated using Glide_SP and enforced by a constraint upon docking. The docking run generated a pool among of 2 poses.
- Model 5 was created by exploring a rotation of the thiophene group. This was done manually in the Schrödinger Maestro suite.

**Selection and refinement of the models –** For each of the three sets of models (27 in total), the most interesting were visually selected, leading to a pool of eight solutions. A ninth solution was created to explore different space for ligand's thiophene group (see above). Models in this pool of solutions were then scored using an in-house scoring function that will be presented elsewhere. Scores were obtained both for the binding pocket and for the ligand. The final selection was done visually and by considering optima of those scores, with aim to keep a diversity of the models. All models were subjected to an in-house water positioning protocol that takes into account ligand using Sybyl atom types. The protocol will be presented elsewhere, and was not used for scoring purpose.

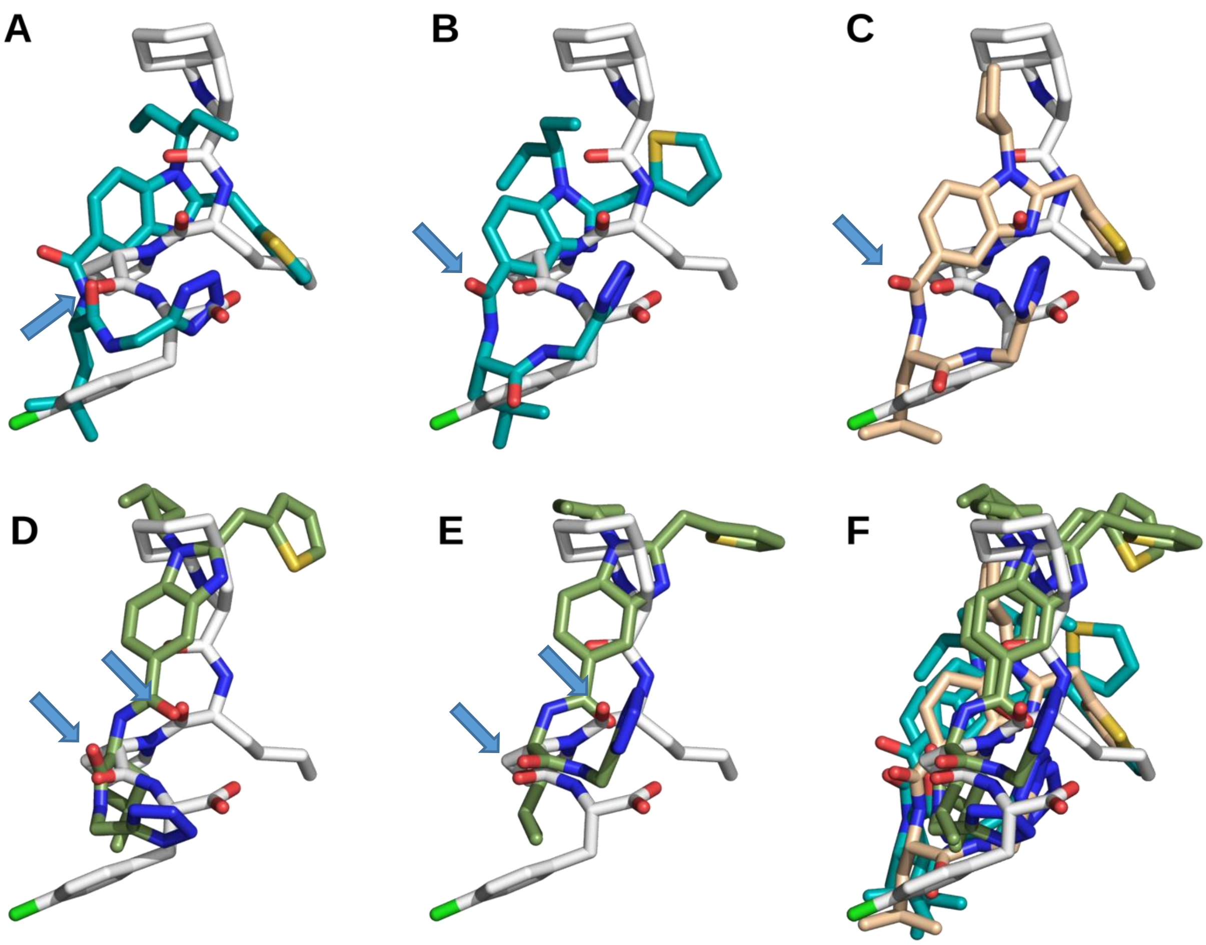

***Figure 2*** *- Superposition of the submitted ligands (cyan, pink-orange, green according to the modeling protocol used) with 5VBL's ligand structural template when the protein structures are superimposed. Panels A-E follows the same order as the submitted models 1-5. Panel A, best complex in the competition (rank 33). Panel F, superimposition of the five submitted models and of the ligand structural template (5VBL). Arrows indicate pairs of carbonyl groups that were structural matched at the model construction and/or selection stages.*

**Ranking of the models –** Ligand poses were ranked based mostly on visual overlap with 5VBL complex.

**Group:** Xhaard-Helsinki **Authors:** Henri Xhaard, Evgeni Grazhdankin, Loic Dreano
**ID:** GPCRdock-278

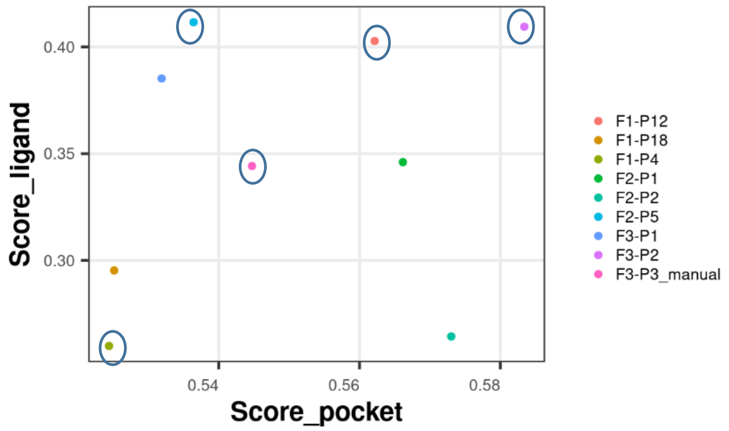

***Figure 3 –** Plot of the scores obtained for the binding pocket against the scores obtained for the ligand using an in-house scoring function. Models selected are indicated by circles: model 1, F1_P12; model 2, F1-P4; model 3, F2_P5; model 4, F3_P2; model 5, F3_P3_manual.*

**Challenges and outcome –** The receptor modeling and identification of the binding pocket were straightforward. Model 4 was the best we submitted (rank 33/201). The main challenge was that surprisingly the 5VBL's ligand carboxylate is not mimicked by the ligand's tetrazole in forming an ionic interaction with receptor. This led to a pose were ligand's tetrazole  is not extended as in the crystallographic answer. We modelled the N-terminus forming a disulphide bridge between Cys 19 and Cys 134 (Figure 4) but there is no information to confirm or infirm this disulphide bridge. Due to time constraints, no experimental data was used during the modeling.

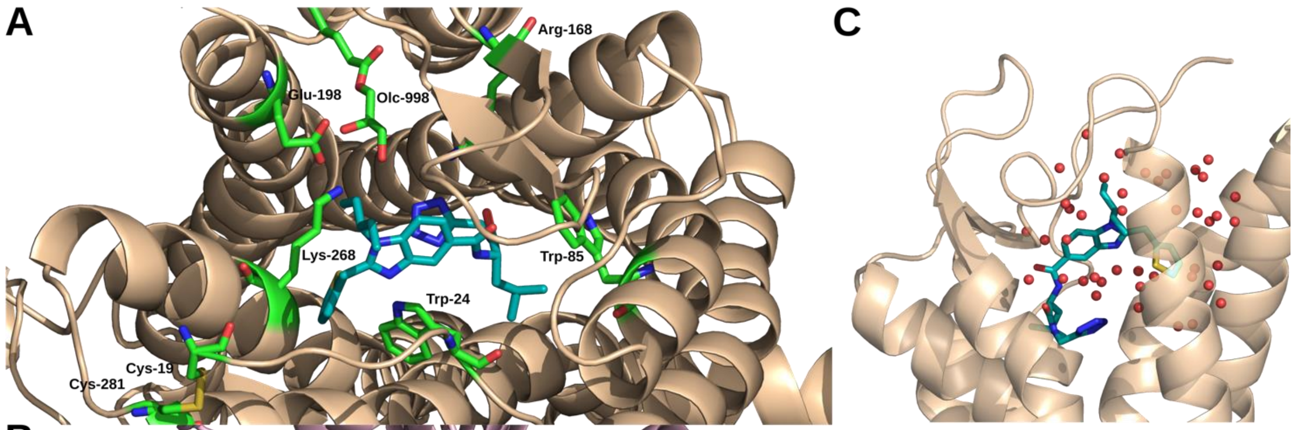

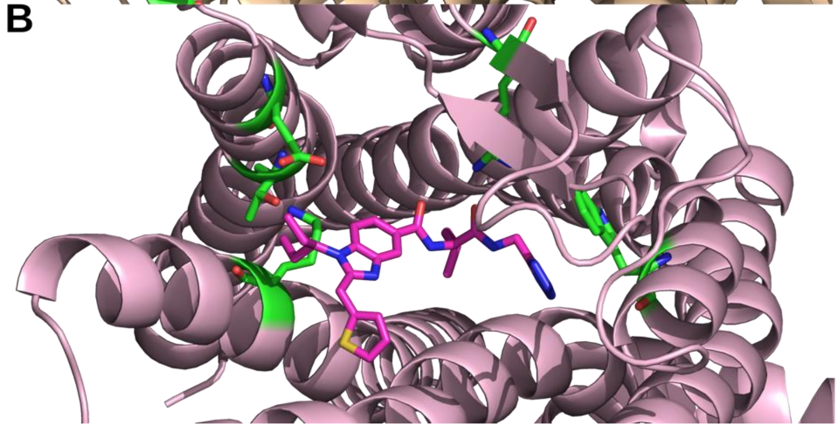

***Figure 4 –** Comparison of the best submitted model (model 1, rank 33/201) (panel A) with the solution (panel B). The ligand 's tetrazole group should have been modeled in a more extended conformation. (C), close-in view of the model 4 solvation around ligand.*

**Group:** Xhaard-Helsinki **Authors:** Henri Xhaard, Evgeni Grazhdankin, Loic Dreano
**ID:** GPCRdock-278

#### Target 2 – GPR139

**Strategy and hypotheses –** We used ”classical” molecular modeling using Modeller v10.0 (Sali & Blundell. J. Mol. Biol. 234, 779-815, 1993) and docking with Schrödinger Ldt. (Schrödinger Release 2019-3: Maestro, Schrödinger, LLC, New York, NY, 2019). The ligand was integrated in Modeller as an object recognized by constructing a topology, which allowed to explore side chain-conformation around ligand for Model 4. Two templates were selected, the neurotensin receptor type 1 (4BUO) with highest percent sequence identity, and the human adenosine A2a receptor (2YDV) to explore small variation in length/position of some transmembrane helices. No AI was used to construct the models.

Main hypotheses:

- GPR139 and the selected templates share the binding site location.
- The binding site is surrounded by a cluster of aromatic residues
- Negatively charged side chains from TM2 (E88, D84) and TM3 (D101, E105, E108) are clustered together and face the binding site. These negative charges are in part neutralized by positively charged residues.
- Extracellular loop 2 (EC2) is divided in two regions, based on the presence of large aromatic amino acid, one on top of TM4, the other at the tip of EC2. The region around W170 form the turn of the hairpin-like EC2.EC3 is key to the placement of R260 and W261, which can interact with ligand/form salt bridges.

**Receptor-ligand complex modeling –** Sequence alignments were constructed by trial-error, in particular portions of EC2 (green, ’PYYWW’, positions 162-166; cyan, ’TSVH’, 177-180) and EC3 (purple, ’LYHLYG’, 249-254; orange, ’NRWLV’, 259-263). The final models were energy-minimized.

Models 1, 2, and 3 were constructed using 4BUO as a template. A small set of distance restraints was injected into MODELLER to improve ECL and binding site side-chain positioning. The final docking study was conducted using the induced-fit protocol in Glide SP. Submitted model 1 and 2 comes from one protein model, Model 3 from anoher.

**Models 1,2 and 3**

```
>P1;T2_GPR139
sequence:T2_GPR139::::::::
---------CGLGFVPVVYYSLLLCLGLPANILTVIILSQL-----VARRQKSVY
NYLLALAAADILVLFFIVFVDFLLEDFILNMQMP----QVPDKIIEVLEFSSIHT
SIWITVPLTIDRYIAVCHPLKYHTVSYPARTRKVIVSVYITCFLTSIPYYWWPNI
WT---------------EDY---ISTSVHHVLIWIHCFTVYLVPCSIFFILNSII
VYKLRRKSN---FRLRGYSTGKTTAILFTITSIFATLWAPRIIMILY-------H
LYGAPIQNRW----------LVHIMSDIANMLALLNTAINFFLYCFISKRFRTMA
AAT*
>P1;4buo
structureX:4buo:52:A:383:A::::
NSDLDVNTDIYSKVLVTAIYLALFVVGTVGNSVTLFTLARKK/-----SLQSTVD
YYLGSLALSDLLILLLAMPVELYNFIWVHHP---WAFGDAGCRGYYFLRDACTYA
TALNVVSLSVELYLAICHPFKAKTLMSRSRTKKFISAIWLASALLAIPMLFTMG-
--LQNLSGDGTHPGGLVCTPIVD--TATLKVVIQVNTFMSFLFPMLVASILNTVI
ANKLTVMVHQ/P--GRVQALRRGVLVLRAVVIAFVVCWLPYHVRRLMFCYISDE-
----------QWTTFLFDFY—HYFYMLTNALVYVSAAINPILYNLVSANFRQVF
LST*
```

Models 4 and 5 were constructed using 2YDV human A2a receptor as a template. Model 5 was selected as the starting point for induced-fit docking. Model 5 was built starting from Model 4, loops were deleted and remodelled using Modeller. Ligand was positioned as Model 4, but a topology was constructed so that it can be recognized by the CHARMM22 force-field in Modeller as a non-rigid body. Several distance restraints were devised and applied: H264-E88, R260-D173, R244-E108, W261-L87, Ligand para-carbon(chlorobenzene substituent)-H187 and -R244, Ligand amine (proximal to chlorobenzene)-E105. An in-house optimization protocol using Modeller was then run for 300 cycles to optimize main-chain and side chain hydrogen-bonding of ligand in the regions spanning residues 88-96, 169-178 and 251-259. The procedure will be presented elsewhere. Model 4 was subjected to an algorithm that position water molecules, which will also be described elsewhere.

**Models 4 and 5**

```
>P1;T2_GPR139
sequence:T2_GPR139::::::::
CGLGFVPVVYYSLLLCLGLPANILTVIILSQLVARRQKSVYNYLLALAAADILVL
FFIVFVDFLLE------DFILNMQMPQVPDKIIEVLEFSSIHTSIWITVPLTIDR
YIAVCHPLKYHTVSYPARTRKVIVSVYITCFLTSIP----YYWWPNIWT------
--------------EDYISTSVHHVLIWIHCFTVYLVPCSIFFILNSIIVYKLRR
KSNFRLRGY/---STGKTTAILFTITSIFATLWAPRIIMILYHLYGA-----PIQ
NRWLV--HIMSDIANMLALLNTAINFFLYCFISKRFRTMAAATLKAFF-------
-----------*
>P1;2ydv
structureX:2ydv:3:A:325:A::::
IMGSSVYITVELAIAVLAILGNVLVCWAVWLN-SNLQNVTNYFVVSAAAADILVG
VLAIPFAIAISTGFCAA---------CHGCLFIACFVLVLTASSIFSLLAIAIDR
YIAIRIPLRYNGLVTGTRAKGIIAICWVLSFAIGLTPMLGW---NN---CGQPKE
GKAHSQGCGEGQVACLFEDVVPMNYMVYFNFFACVLVPLLLMLGVYLRIFLAARR
QLKQMESQS/TLQKEVHAAKSLAIIVGLFALCWLPLHIINCFTFF--CPDCS---
-H---APLWLMYLAIVLSHTNSVVNPFIYAYRIREFRQTFRKIIRSHVLRQQEPF
KAAAAENLYFQ*
```

**Group:** Xhaard-Helsinki **Authors:** Henri Xhaard, Evgeni Grazhdankin, Loic Dreano
**ID:** GPCRdock-278

**Ranking and selection of the five solutions** – The five final poses were selected both visually and based on an in-house knowledge-based score applied to atoms of the ligand and of the binding pocket. The visual selection aimed to ensure a diversity in the model EC2 and EC3 conformations as well as a proper coverage of the binding site by the poses. The five selected poses are presented at the end of this document (panel A, rank 1, and so forth).

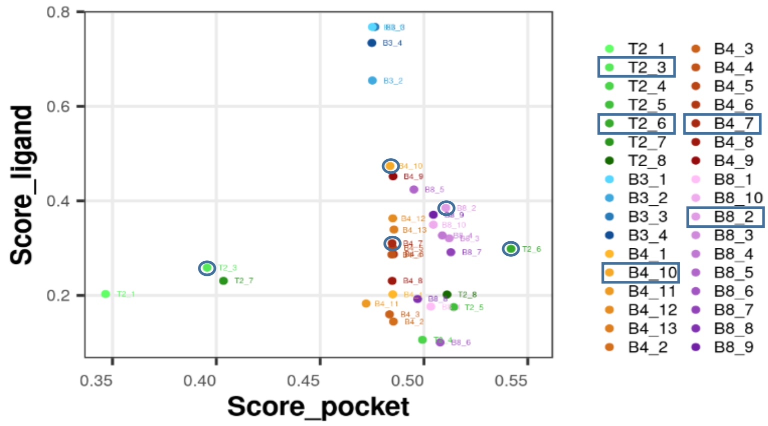

***Figure 3 –** Plot of the scores obtained for the binding pocket against the scores obtained for the ligand using an in-house scoring function. Models selected are indicated by circles.*

**Challenges –** The main challenge for GPR139 was first to model the receptor, and in particular the loop region, and then to position the ligand. In the end, our best pose has the ligand flipped along its short axis. The panel of solutions we gave (Figure 4D) are approximately in line with the solution (Figure 4E), but overall, we lack the ability to score/reconstruct proper packing. One challenge was to explore the length of the helices, in particular. Our EC2 in model 3 (rank 97/193, this is our best ranking model) is fairly close to the crystallographic answer in terms of conformation of this important loop, but the tip of the loop is shifted by ~10Å. Due to time constraints, no experimental data were used during the modeling procedure.

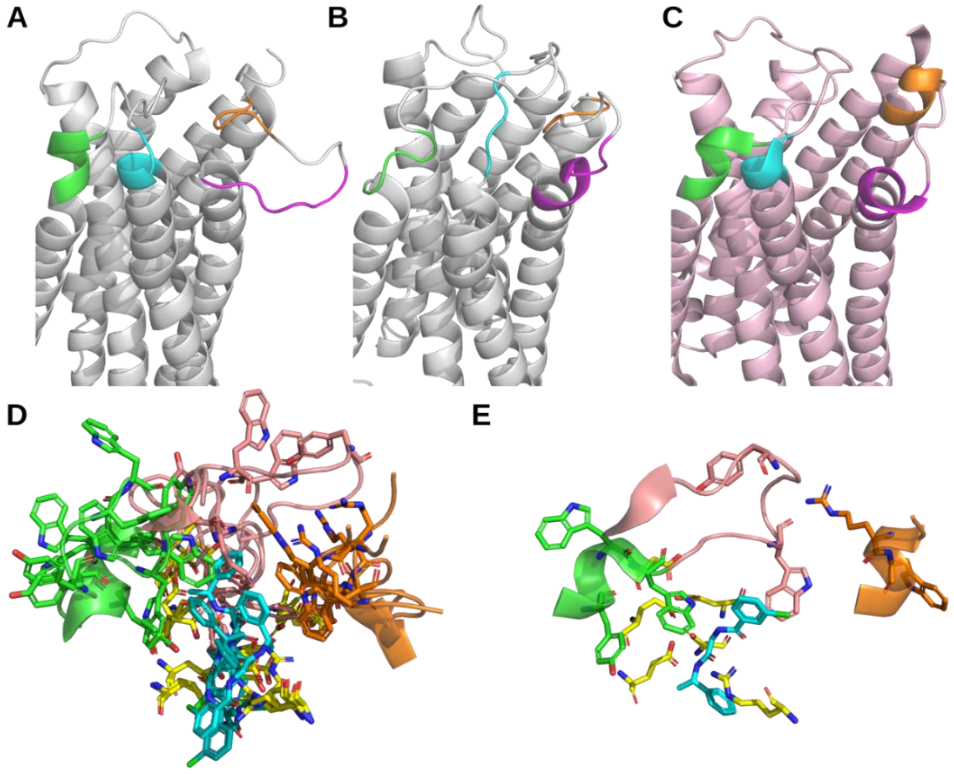

***Figure 4 –** Panels A-C Variation of the length of the transmembrane segments TM4-TM7 (green, cyan, purple and orange, respectively) in (A) model 1, (B) model 4, and (C) answer. In (D) and (E), superimposition of the five submitted model and the answer. Color coding (see alignment above) highlights the regions key for modeling.*

*Caution: the color coding in panels D and E is different that in panels A-C.*

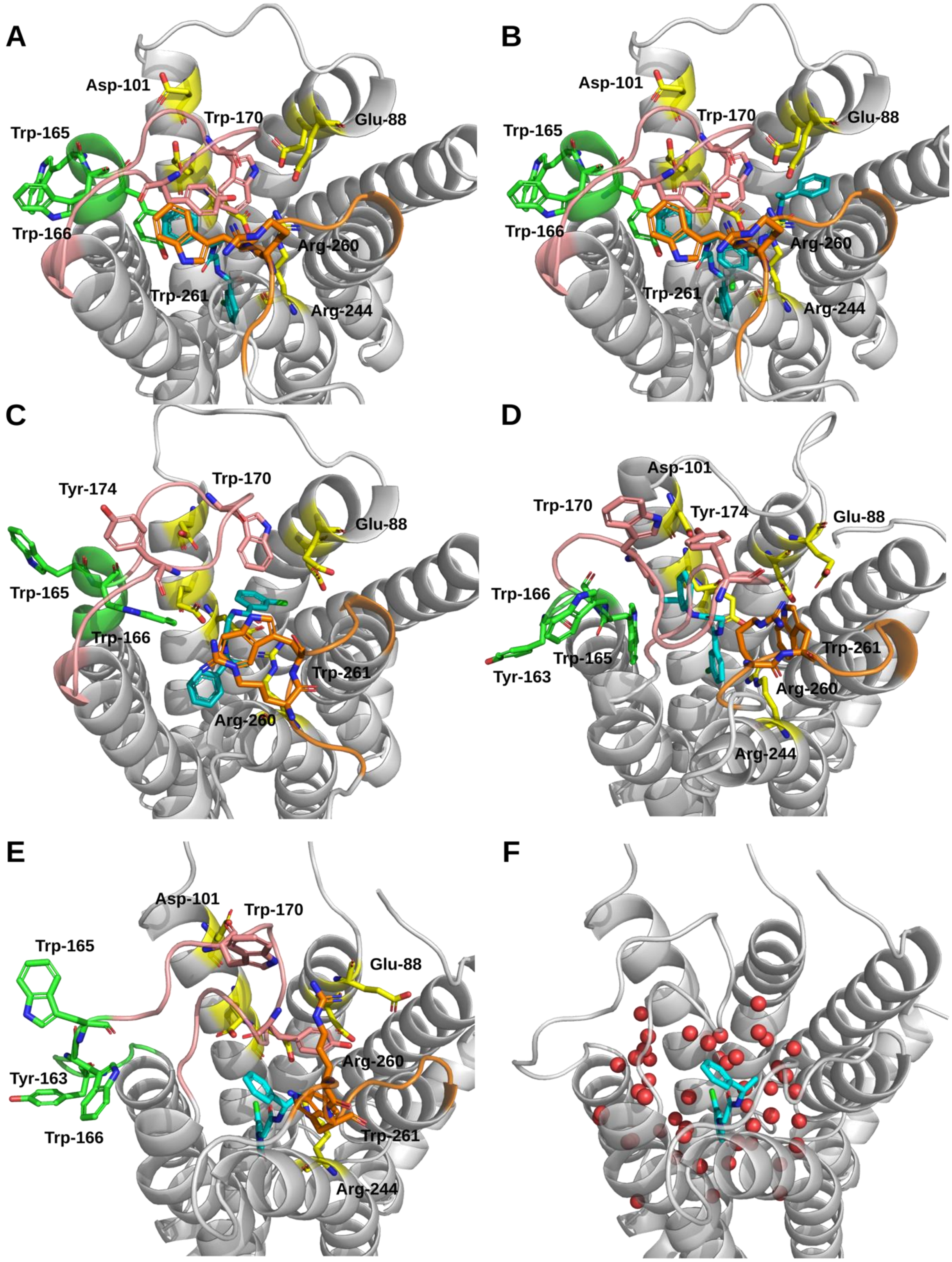

***Figure 5 –*** *Key elements of the submitted models. Panels A-E follows the same order as the submitted models 1-5. Panel C, best complex in the competition (rank 97/193). Panel F, Details of the solvation of model 5. Color indicate key positions from EC2 (green, 'PYYWW', positions 162-166; cyan, 'TSVH', 177-180) and EC3 (purple, 'LYHLYG', 249-254; orange, 'NRWLV', 259-263).*

**Group:** Xhaard-Helsinki
**ID:** GPCRdock-278

**Authors:** Henri Xhaard, Evgeni Grazhdankin, Loic Dreano

**Target 3 – KOR**

**Strategy and hypotheses** –We used hypothesis-driven peptide and protein modeling, where a "seed" of the peptide is placed on the binding site based on existing structures, and the peptide is reconstructed using Modeller (Sali & Blundell. J. Mol. Biol. 234, 779-815, 1993). Restraints were used to enforce close interactions and peptide predicted secondary structure. No AI was used to create the models.

Main hypotheses:

- The bound peptide mimics the cocrystallized ligands in KOR structures. In particular the positively charged N-terminus of the "YGGFLRRIR" peptide is forming a salt bridge with D138.
- Peptide's N-terminal residues can be modeled on top of the JDC ligand (PDB code 6VI4) or the CVV ligand (6B73).
- The peptide can form a beta-turn around its GG region, with intramolecular hydrogen bonding.

**Receptor-Ligand complex modeling** – Peptide and the missing parts from the PDB structure were reconstructed with Modeller based on the human kappa-opioid receptor (Models 2 and 3, PDB 6VI4; models 1, 4 and 5, PDB 6B73). The "seeds" for the peptide (peptide residues "YG" or "YxxF", Figure 1) were reconstructed by first changing atom denominations (from ligand HET to ATOM). Sets of restraints were used to enforce specific interactions in the models, described here only for model 4.

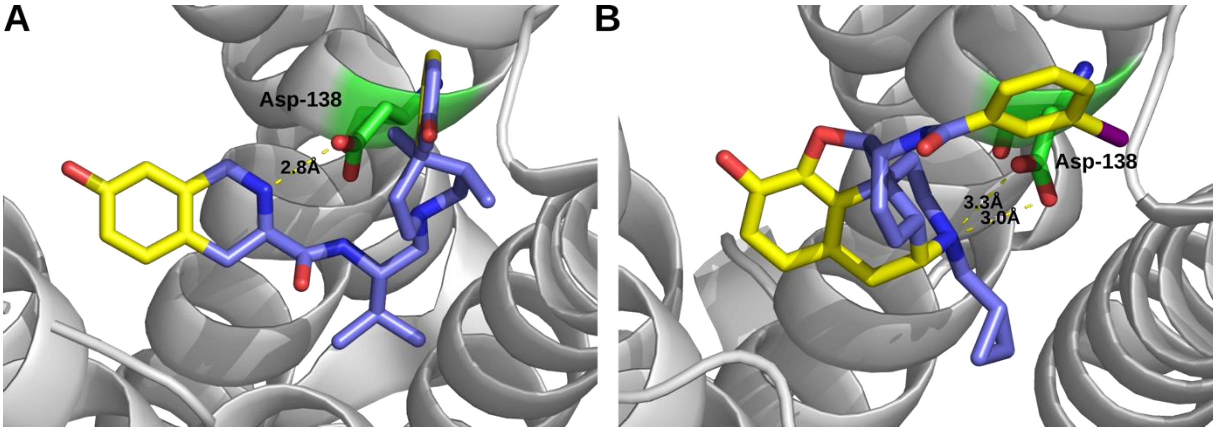

```
Models 1, 4 and 5
>P1;T3_KOR
sequence:T3_KOR::::::::
PAIPVIITAVYSVVFVVGLVGNSLVMFVIIRYTKMKTATNIYIFNLALADALVTT
TMPFQSTVYLMNSWPFGDVLCKIVLSIDYYNMFTSIFTLTMMSVDRYIAVCHPVK
ALDFRTPLKAKIINICIWLLSSSVGISAIVLGGTKVREDVDVIECSLQFPDDDYS
WWDLFMKICVFIFAFVIPVLIIIVCYTLMILRLKSVRLLSGSREKDRNLRRITRL
VLVVVAVFVVCWTPIHIFILVEALGSTSHSTAALSSYYFCIALGYTNSSLNPILY
AFLDENFKR/YGGFLRRIR*
>P1;6b73_A_seed
structureX:6b73_A_seed:56:A:343:B::::
PAIPVIITAVYSVVFVVGLVGNSLVMFVIIRYTKMKTATNIYIFNLALADALVTT
TMPFQSTVYLMNSWPFGDVLCKIVLSIDYYNMFTSIFTLTMMSVDRYIAVCHPVK
ALDFRTPLKAKIINICIWLLSSSVGISAIVLGGTKVR---DVIECSLQFPDDDYS
WWDLFMKICVFIFAFVIPVLIIIVCYTLMILRLKSVR----SREKDRNLRRITRL
VLVVVAVFVVCWTPIHIFILVEALG-TSHSTAALSSYYFCIALGYTNSSLNPILY
AFLDENFKR/Y--F-----*
```

```
Models 2 and 3
>P1;T3_KOR
sequence:T3_KOR::::::::
PAIPVIITAVYSVVFVVGLVGNSLVMFVIIRYTKMKTATNIYIFNLALADALVTT
TMPFQSTVYLMNSWPFGDVLCKIVLSIDYYNMFTSIFTLTMMSVDRYIAVCHPVK
ALDFRTPLKAKIINICIWLLSSSVGISAIVLGGTKVREDVDVIECSLQFPDDDYS
WWDLFMKICVFIFAFVIPVLIIIVCYTLMILRLKSVRLLSGSREKDRNLRRITRL
VLVVVAVFVVCWTPIHIFILVEALGSTSHSTAALSSYYFCIALGYTNSSLNPILY
AFLDENFKR ---------/YGGFLRRIR*
>P1;6vi4_A_peptide_seed2
structureX:6vi4_A_peptide_seed2:60:A:402:B::::
----VIITAVYSVVFVVGLVGNSLVMFVIIRYTKMKTATNIYIFNLALADALVTT
TMPFQSTVYLMNSWPFGDVLCKIVLSIDYYNMFTSIFTLTMMSVDRYIAVCHPVK
ALDFRTPLKAKIINICIWLLSSSVGISAIVLGGTKVREDVDVIECSLQFPDDDYS
WWDLFMKICVFIFAFVIPVLIIIVCYTLMILRLKSVRLLSGSREKDRNLRRITRL
VLVVVAVFVVCWTPIHIFILVEALG------AALSSYYFCIALGYTNSSLNPILY
AFLDENFKRCFRDFCFPL/YG-------*
```

***Figure 1 –** Structural anchors (A) ligand JDC (PDB 6VI4), used for models 2 and 3; (B) ligand CVV (PDB 6B73), used for models 1, 4 and 5. In yellow, carbon atoms used to construct the peptide "seed". Oxygen and nitrogen atoms were also included in the seed (not indicated). D138 (green) forms a salt bridge with ligand. Below, sequence alignments used for building models 1-5. Peptide seed residues (yellow) were constructed before running Modeller by changing selected ligand HET into ATOM denominations and running Modeller.*

Model 4 was built from the kappa-opioid receptor in active state, PDB code 6B73. We first modeled a structural anchor, amino acids YG, from the cocrystallized CVV ligand. The full peptide was then modeled and a set of restraints defined by trial-error to enforce packing with protein: intramolecular salt bridges networks connecting peptide R6 and receptor E297 and D223 as well as peptide R7 and receptor E203. In addition, we enforced a peptide-receptor salt bridge between D138 and peptide N-terminal; as well as aromatic/hydrophobic packing interactions: peptide F4 to receptor W124, peptide L5 to receptor I294; and peptide's I8 to receptor L212. In addition, we enforced the peptide to form a turn at its GG amino acids, by setting restraints from F4 to Y1 and G2; and between G3 and Y1. We then ran in-house distance restraints search and optimization protocol on the whole structure, 300 rounds, to increase the hydrogen bond network. The protocol will be described elsewhere.

For Model 5, we investigated flipping the peptide 180 degrees (orthogonally to the membrane plane). A 'YGGF' peptide was taken from model 4 and docked with Glide SP to identify a suitable starting pose. The pose was refined and the peptide completed using Modeller and adjusted with distance restraints.

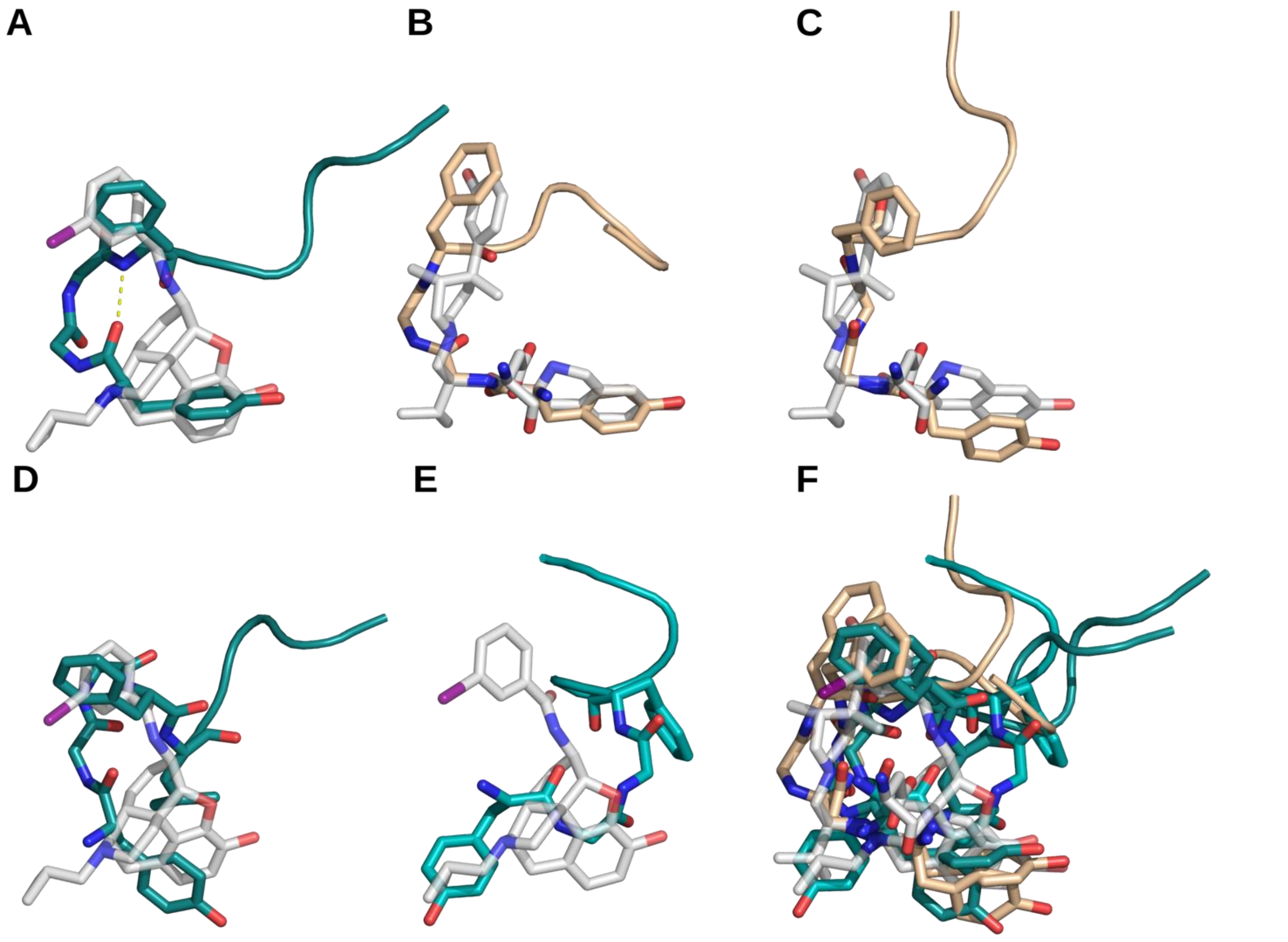

***Figure 2 –*** *Comparison of the modeled peptides with the structural anchors used from (A, D, E) PDB 6B73, or (B,C) 6VI4. Models are ordered as ranked in the Competition (panel A, model 1, etc). Panel C, best complex in the competition (rank 21). (F) Peptide space explored by the different models.*

**Ranking** – Models were ranked visually, considering their closeness to the structural templates.

**Challenges –** We correctly identified the structural anchors and hence correctly positioned the peptide N-terminus, leading to good ranks (rank 22/165 for OPKR and rank 14/165 for OPKR-core)(Figure 3).

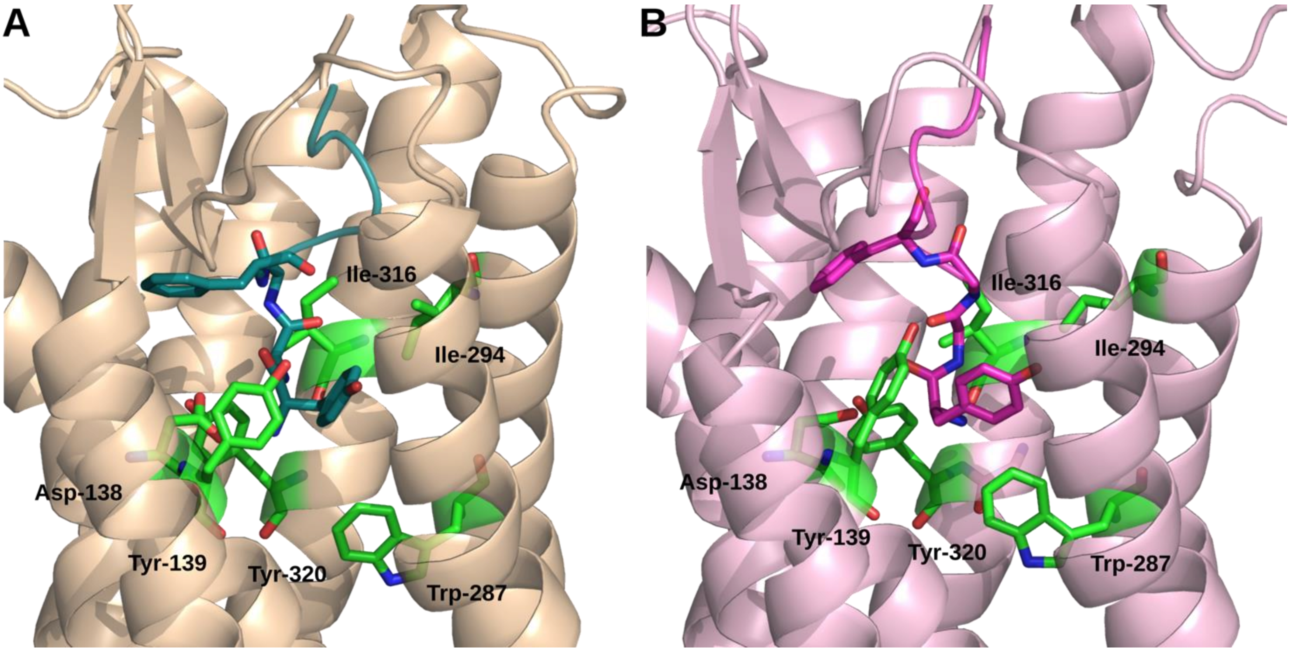

***Figure 3–** Close-in view for (A) the best submitted model, model 3, rank 21. (B) Crystal structure of the "answer".*

A challenge comes from the crystal structure itself (Figure 4), which does not bear a salt bridge between the peptide N-terminus and D138. Of note this is not in line with previous knowledge concerning opioid receptor ligands (Figure 1), as well as not parsimonious because opposite charged are located in the vicinity without neutralizing each other (Figure 4). Generally, the tools available lacked the ability to properly repack the loops; as well as to explore secondary structures including extending or shortening TMs. Due to time constraints, no experimental data was used during the modeling.

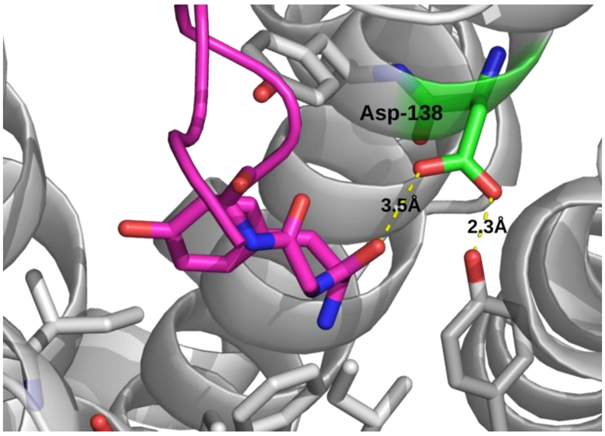

***Figure 4** – Feature of D138 in the crystallographic answer used to evaluate models. D138 appears to be neutral, since it interacts with a carbonyl and phenolic group. The likely positively charged peptide N-terminal is not neutralized by a counter-ion. A more parsimonious solution is to form a salt bridge between N-terminal and D138.*

**Group:** Xhaard-Helsinki
**ID:** GPCRdock-278
**Authors:** Henri Xhaard, Evgeni Grazhdankin, Loic Dreano

**Target 4 – NPY1**

**Strategy and hypotheses** –We used hypothesis-driven peptide and protein modeling, where a "seed" of the peptide is defined based on (i) the bound UR-MK299 ligand from the structure of the neuropeptide Y Y1 receptor (PDB code 5ZBQ) and (ii) a portion of the receptor N-terminus, and the peptide and receptor are reconstructed together using Modeller (Sali & Blundell. J. Mol. Biol. 234, 779-815, 1993). Restraints were used to enforce chosen intramolecular and intermolecular contacts. No AI was used to create the models.

Main hypotheses:

- Peptide C-terminal fits into the binding site of the human neuropeptide Y Y1 receptor, PDB 5ZBQ.
- Peptide residue R35 form a salt bridge with receptor D287.
- The 'EKNAQ' (residues 20-24) N-terminal portion of PDB ID 5ZBQ is mimicked by peptide.
- The peptide contains a long helix in its central part, as suggested by sequence-based prediction by PEPFOLD ( https://mobyle.rpbs.univ-paris-diderot.fr/ )
- The peptide binding mode has similarities with the binding mode of other peptides such as orexin B to its cognate receptor (PDB 7L1U).

**Receptor-Ligand complex modeling –** A set of five models were constructed using the PDB 5ZBQ template. Two alignments leading to different peptide position were used. A few main chain atoms were taken from PDB 7L1U (orexin B peptide bound to orexin receptor) and placed deep in the binding site to seed the most intracellular part of the peptide. The terminal NH2 was added with an awk script post-modeling.

**Models 1, 2 and 5**

```
>P1;T4_NPY1R
sequence:T4_NPY1R:::::::
FENDDCHLPLAMIFTLALAYGAVIILGVSGNLALIIIILKQKEMRNVTNILIVNL
SFSDLLVAIMCLPFTFVYTLMDHWVFGEAMCKLNPFVQCVSITVSIFSLVLIAVE
RHQLIINPRGWRPNNRHAYVGIAVIWVLAVASSLPFLIYQVMTDEPFQNVTLDAY
KDKYVCFDQFPSDSHRLSYTTLLLVLQYFGPLCFIFICYFKIYIRLKRRNNMMDK
MRDNKYRSSETKRINIMLLSIVVAFAVCWLPLTIFNTVFDWNHQIIATCNHNLLF
LLCHLTAMISTCVNPIFYGFLNKNFQRDLQFFFN/YPSKPDNPGEDAPAEDMARY
YSALRHYINLITRQRY*
>P1;5zbq_A_seed_tail
structureX:5zbq_A_seed_tail:32:A:1036:B::::
----DCHLPLAMIFTLALAYGAVIILGVSGNLALIIIILKQKEMRNVTNILIVNL
SFSDLLVAIMCLPFTFVYTLMDHWVFGEAMCKLNPFVQCVSITVSIWSLVLIAVE
RHQLIINPRGWRPNNRHAYVGIAVIWVLAVASSLPFLIYQVMTDEPFQNVTLDAY
KDKYVCFDQFPSDSHRLSYTTLLLVLQYFGPLCFIFICYFKIYIRLKRR------
--------SETKRINIMLLSIVVAFAVCWLPLTIFNTVFDWNHQIIATCNHNLLF
LLCHLTAMISTCVNPIFYGFLNKNFQRDLQFFFN/--------------------
-FSEKNAQLLA--QRY*
```

**Models 3 and 4**

```
>P1;T4_NPY1R
sequence:T4_NPY1R:::::::
FENDDCHLPLAMIFTLALAYGAVIILGVSGNLALIIIILKQKEMRNVTNILIVNL
SFSDLLVAIMCLPFTFVYTLMDHWVFGEAMCKLNPFVQCVSITVSIFSLVLIAVE
RHQLIINPRGWRPNNRHAYVGIAVIWVLAVASSLPFLIYQVMTDEPFQNVTLDAY
KDKYVCFDQFPSDSHRLSYTTLLLVLQYFGPLCFIFICYFKIYIRLKRRNNMMDK
MRDNKYRSSETKRINIMLLSIVVAFAVCWLPLTIFNTVFDWNHQIIATCNHNLLF
LLCHLTAMISTCVNPIFYGFLNKNFQRDLQFFFN/YPSKPDNPGEDAPAEDMARY
YSALRHYINLITRQRY*
>P1;5zbq_A_seed_tail
structureX:5zbq_A_seed_tail:32:A:1036:B::::
----DCHLPLAMIFTLALAYGAVIILGVSGNLALIIIILKQKEMRNVTNILIVNL
SFSDLLVAIMCLPFTFVYTLMDHWVFGEAMCKLNPFVQCVSITVSIWSLVLIAVE
RHQLIINPRGWRPNNRHAYVGIAVIWVLAVASSLPFLIYQVMTDEPFQNVTLDAY
KDKYVCFDQFPSDSHRLSYTTLLLVLQYFGPLCFIFICYFKIYIRLKRR------
--------SETKRINIMLLSIVVAFAVCWLPLTIFNTVFDWNHQIIATCNHNLLF
LLCHLTAMISTCVNPIFYGFLNKNFQRDLQFFFN/---------------FSEKN
AQLLA---------QRY*
```

***Figure 1 –*** *Sequence alignments used for modeling. The fragments of the template used to anchor the peptide are in yellow (N-terminal fragment) and in purple-green-yellow (ligand anchor). Modeller is set free to reconstruct the remaining parts of the peptide.*

Set of models constructed:

- Models 3 and 4 explore the hypothesis that the potential peptide alpha helical region is close to TM7.
- Models 1, 2, and 5 explore the hypothesis that the TM7 is farther away.
- For models 1, 3, 4, and 5, alpha-helix restraints were used on peptide residues 14 to 32.
- Model 5 was subjected to 300 rounds of optimization to increase hydrogen bonding on the whole structure. Distance restraints were set to enforce a salt bridge between receptor D287 and peptide R35 as well as interactions between receptor N299 and peptide Q343; and receptor D287 and peptide R33.

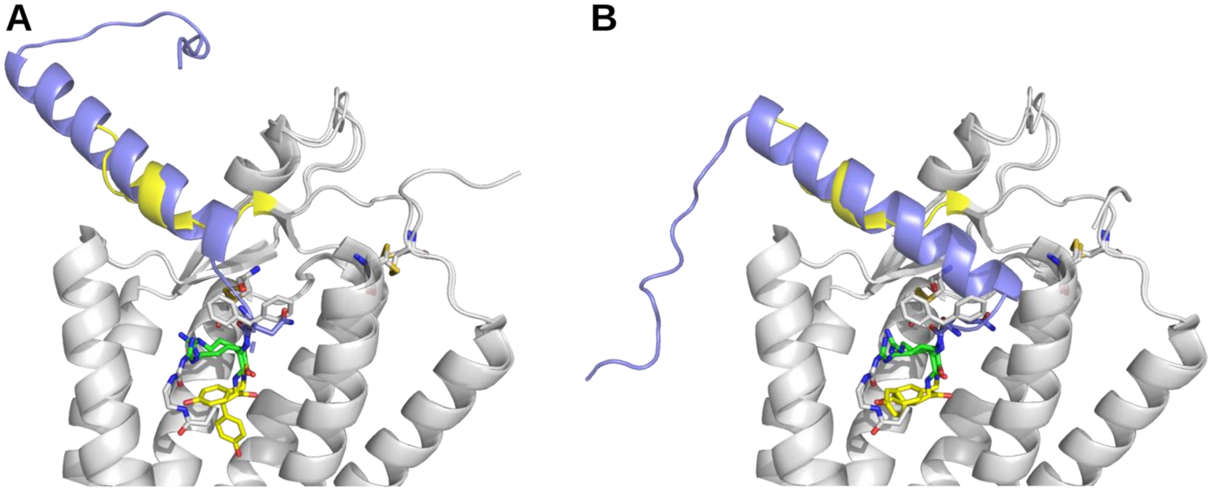

***Figure 2 –** Alignment of selected models on the structural template PDB code 5ZBQ. (A) models 1, 2, and 5. (B) models 3 and 4. Note that the purple peptide alpha helical segment is shifted towards TM1 (right of the picture) depending on the alignment used. Yellow and green, part of the structural template that were used to anchor the peptide model.*

**Ranking** – The models are relatively similar (Figure 3A). They were ranked manually.

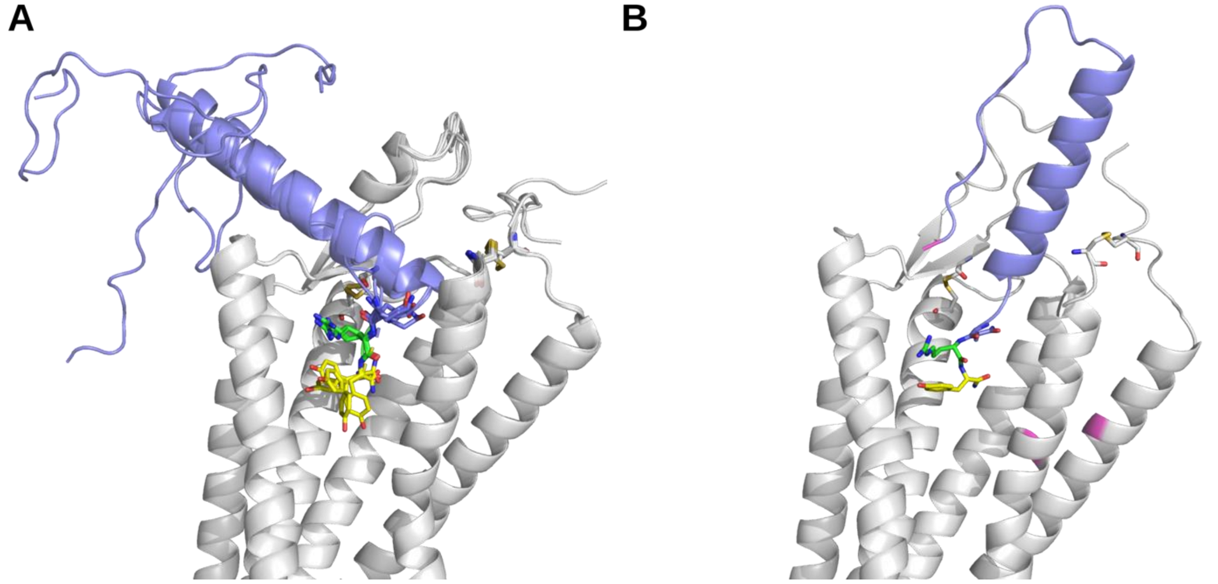

***Figure 3** – (A) Pool of models submitted. In yellow and green, the peptide Y and R anchors respectively. (B) "answer" to the competition. Magenta, poorly refined regions of the crystal structure.*

**Challenges** – We correctly identified the structural anchor deep in the binding pocket (Figure 3, Figure 4), but it was not sufficient to rank well in the competition (best rank 119/145). The $\alpha$-helical part of the peptide is there, but not properly positioned.

**Group:** Xhaard-Helsinki **Authors:** Henri Xhaard, Evgeni Grazhdankin, Loic Dreano
**ID:** GPCRdock-278

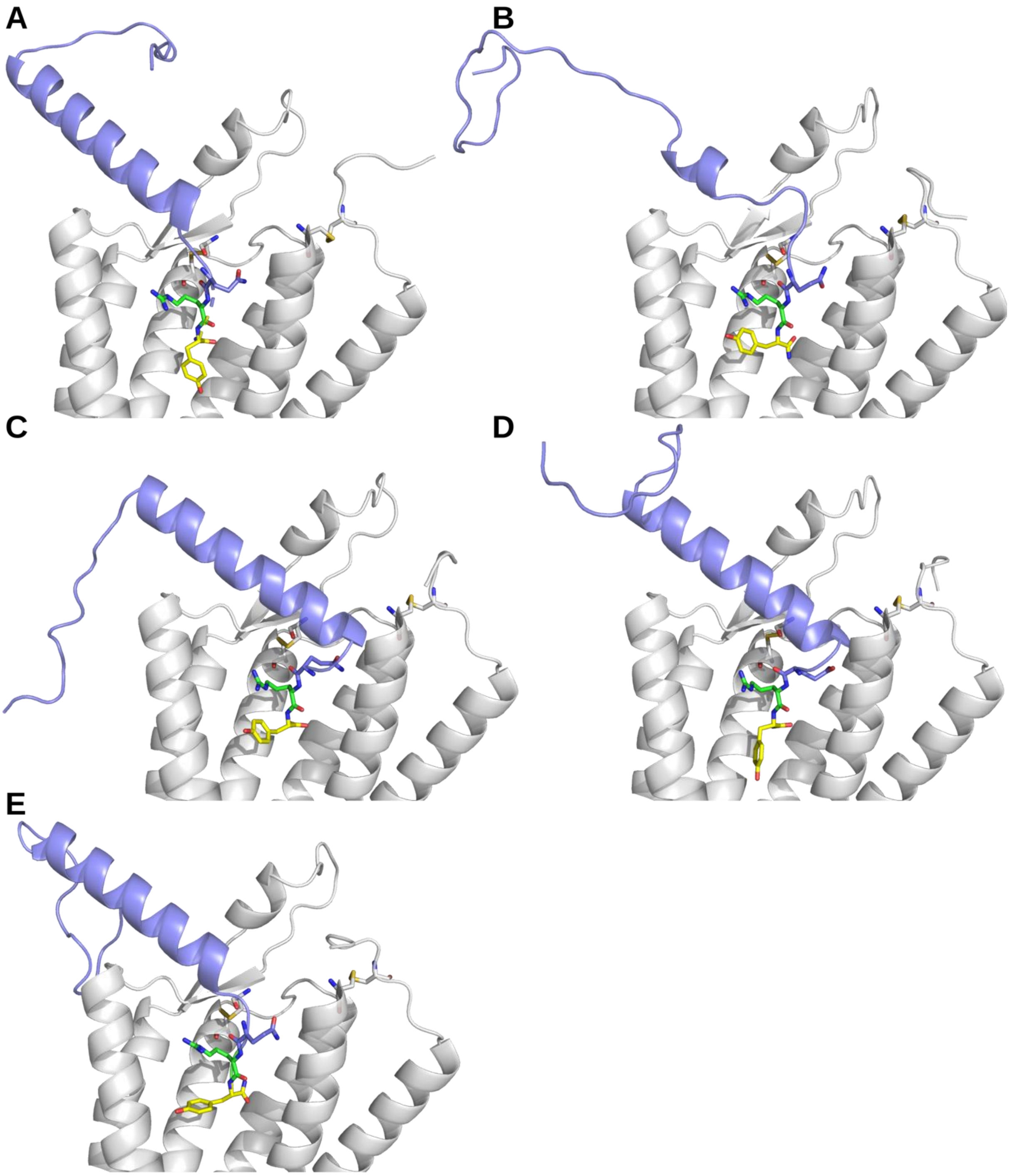

***Figure 4*** *– The five models that were submitted (rank 1, panel A, and so forth). In (D) the best model submitted, rank 119/145.*

**Group:** Xhaard-Helsinki
**ID:** GPCRdock-278

**Authors:** Henri Xhaard, Evgeni Grazhdankin, Loic Dreano

Target 5 – NMUR2

**Strategy and hypotheses** –We used hypothesis-driven peptide and protein modeling, where a "seed" of the peptide is placed on the binding site based on existing structures, and the peptide is reconstructed using Modeller (Sali & Blundell. J. Mol. Biol. 234, 779-815, 1993). Restraints to guide molecular interactions and peptide predicted secondary structure were used. No AI was used to create the submitted models.

Main hypotheses:

- The peptide binding mode has similarity to the binding mode of other peptides to their cognate GPCRs
- The C-terminal binds "deep" in the binding site. The depth of the peptide is unknown.
- The N-terminal of the peptide folds as an alpha-helix. This is by similarity to the orexin peptide and other peptides (peptide secondary structure prediction suggested a beta-strand conformation).

**Receptor-Ligand complex modeling –** The neutotensin receptor (PDB 4BUO) was selected as a template for all models, and a multiple sequence alignment constructed using ClustalW. Models were generated using MODELLER, with distance restraints set for enforcing alpha-helix in the peptide between residues 1 and 14, and a few other visually determined interactions. The terminal NH2 was added with an awk script.

**Models 1, 2, 3 and 4**

```
>P1;T5_NMUR
sequence:T5_NMUR:::::::
---------------HFFLPVSVVYVPIFVVGVIGNVLVCLVILQH-/--QAMKT
PTNYYLFSLAVSDLLVLLLGMPLEVYE-MWRNYPFLFGPVGCYFKTALFETVCFA
SILSITTVSVERYVAILHPFRAKLQSTRRRALRILGIVWGFSVLFSLPNTSIHGI
KFHYFPNGSLVPGSATCTVIKPMWIYNFIIQVTSFLFYLLPMTVISVLYYLMALR
LKKDKS-/----LEADEGNANIQRPCRKSVNKMLFVLVLVFAICWAPFHIDRLFF
SFV--EEWSESLAAVFNLVHVVSGVFFYLSSAVNPIIYNLLSRRF--------/F
RVDEEFQSPFASQSRGYFLFRPRN*
>P1;4buo_A_C
structureX:4buo_A_C:52:A:13:C::::
-----NSDLDVNTDIYSKVLVTAIYLALFVVGTVGNSVTLFTLARKK/---SLQS
TVDYYLGSLALSDLLILLLAMPVELYNFIWVHHPWAFGDAGCRGYYFLRDACTYA
TALNVVSLSVELYLAICHPFKAKTLMSRSRTKKFISAIWLASALLAIPMLFTMGL
--QNLSGDGTHPGGLVCTPIVDTATLKVVIQVNTFMSFLFPMLVASILNTVIANK
LTVMVHQ/--------PGRVQA---LRR-GVLVLRAVVIAFVVCWLPYHVRRLMF
CYISDEQWTTFLFDFYHYFYMLTNALVYYSAAINPILYNLVSANFRQVFLSTL/-
-------------RRPYIL-----*
```

**Model 5**

```
>>P1;T5_NMUR
sequence:T5_NMUR:::::::
---------------HFFLPVSVVYVPIFVVGVIGNVLVCLVILQH-/--QAMKT
PTNYYLFSLAVSDLLVLLLGMPLEVYE-MWRNYPFLFGPVGCYFKTALFETVCFA
SILSITTVSVERYVAILHPFRAKLQSTRRRALRILGIVWGFSVLFSLPNTSIHGI
KFHYFPNGSLVPGSATCTVIKPMWIYNFIIQVTSFLFYLLPMTVISVLYYLMALR
LKKDKS-/----LEADEGNANIQRPCRKSVNKMLFVLVLVFAICWAPFHIDRLFF
SFV--EEWSESLAAVFNLVHVVSGVFFYLSSAVNPIIYNLLSRRF--------/F
RVDEEFQSPFASQSRGYFLFRPRN*
>P1;4buo_A_C
structureX:4buo_A_C:52:A:13:C::::
-----NSDLDVNTDIYSKVLVTAIYLALFVVGTVGNSVTLFTLARKK/---SLQS
TVDYYLGSLALSDLLILLLAMPVELYNFIWVHHPWAFGDAGCRGYYFLRDACTYA
TALNVVSLSVELYLAICHPFKAKTLMSRSRTKKFISAIWLASALLAIPMLFTMGL
—QNLSGDGTHPGGLVCTPIVDTATLKVVIQVNTFMSFLFPMLVASILNTVIANKL
TVMVHQ/--------PGRVQA---LRR-GVLVLRAVVIAFVVCWLPYHVRRLMFC
YISDEQWTTFLFDFYHYFYMLTNALVYYSAAINPILYNLVSANFRQVFLSTL/--
------------------RRPYIL*
```

Two docking depth were tested, by varying the alignment to cognate peptide (red, alpha helical region of the orexin peptide; yellow, structural matches).

**Model 1,2,3,4 – peptide alignments**

```
orexin       -RSGPPGLQGRLQRLLQASGNHAAGILTM--
neurotensin  ---------------------RRPYIL-----
nmur_deep    ------FRVDEEFQSPFASQSRGYFLFRPRN
```

**Model 5 – peptide alignments**

```
orexin        -RSGPPGLQGRLQRLLQASGNHAAGILTM--
neurotensin   ---------------------RRPYIL-----
nmur_shallow  FRVDEEFQSPFASQSRGYFLFRPRN------
```

- Models 1,2,3, and 4 represent the deepest depth of peptide docking. They were all created using the same alignment but represent different solutions as provided by Modeller.
- Model 4 was run through a cycle of optimization (200 cycles on the whole structure) that lead to an increase of intramolecular hydrogen bonds network. Distance restraints were added to enforce salt bridges between peptide and receptor : peptide R16 to receptor V294 on the top of TM6; peptide's R24 and E102; peptide E22 to E127; as well as aromatic packing of peptide Y18 with F192.

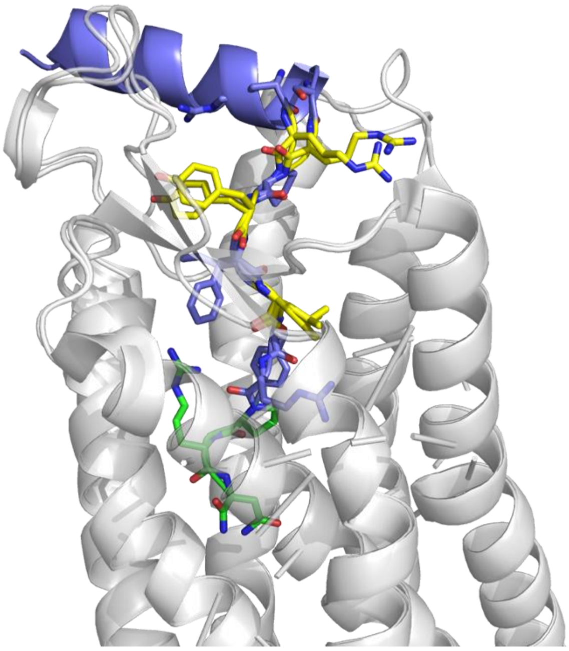

***Figure 1 –*** *Superposition of the structural template 4BUO with model 1.*

**Ranking** – The models are relatively similar. They were ranked manually, considering a deep binding mode to be more probable.

**Challenges** – We correctly identified the peptide orienting to bind with its C-terminal deep in the binding site (Figure 2). We did not explore the proper depth of the peptide (Figure 2C). The best rank achieved was 101/147. Due to time constraints, no experimental data was used during the modeling procedure.

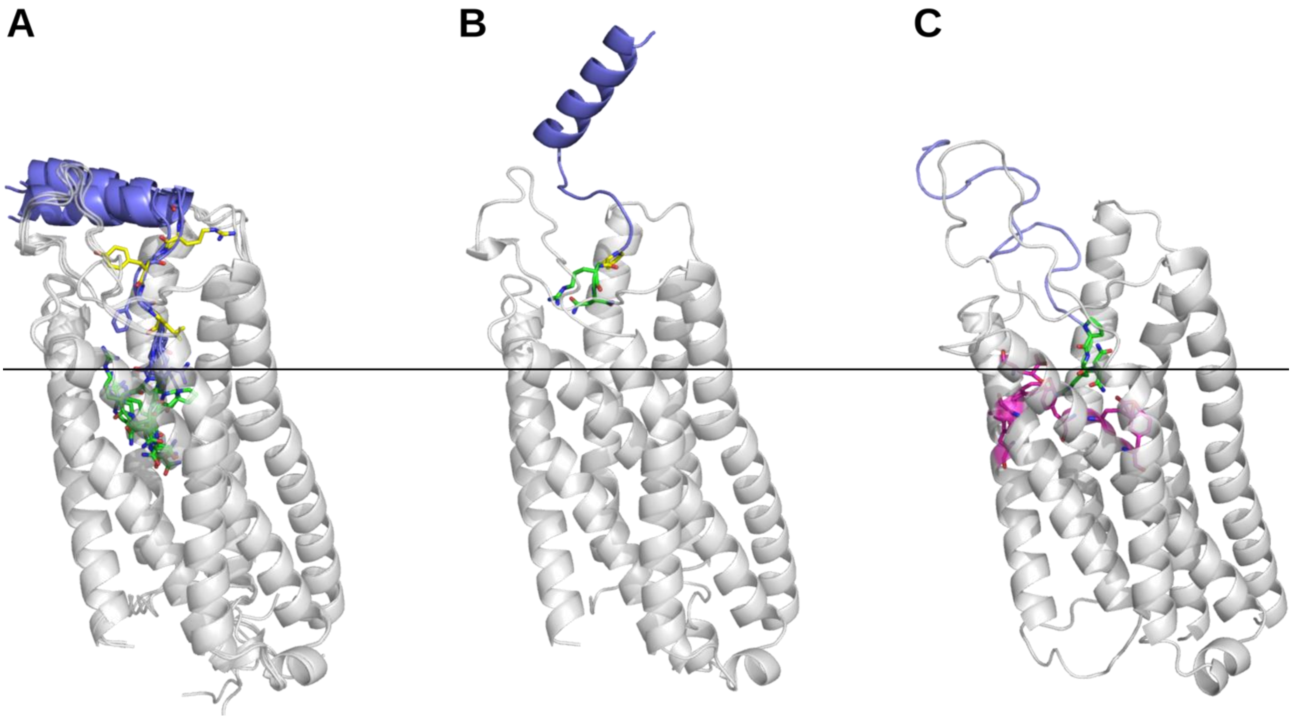

***Figure 2 –*** *Variety of poses covered by (A) Models 1-4 and (B) model 5. In (C), answer to the competition. The line indicates the peptide depth, and we set the peptide either too deep or too much closer to extracellular surface. In magenta, parts of the structure that contain clashes. The best model in the competition (panel A, model 3) has rank 101/147.*

### GPCRdock-469

Heather Noriega, Najla Alzaid, and Xiang Simon Wang*

*Molecular Modeling and Drug Discovery Core Laboratory for District of Columbia Center for AIDS Research (DC CFAR), Department of Pharmaceutical Sciences, College of Pharmacy, Howard University, Washington, DC 20059, U.S.A.*

Initial building of the receptor models can be characterized using deep learning AlphaFold 2. This network has improved the accuracy of structure prediction by incorporating neural network architectures and training procedures based on the developmental, physical, and geometric constraints of protein structures. This system jointly embeds multiple sequence alignments (MSAs) and pairwise features, a new output representation and associated loss that enables accurate end to end structure prediction. AlphaFold 2 directly predicts 3-D coordinates of all heavy atoms for a given protein using the amino acid sequence and aligned sequences of homologues as inputs and yields highly accurate structure predictions. [1] [2] It is available in multiple formats, however, in this article it was executed through ChimeraX. [3] Sequences were obtained and run through AlphaFold 2 in ChimeraX platform. Once the network finished a 3-D structure prediction appeared with models, scores, and sequence associated data in the completed file.

Models were then refined, with deletion of any unnecessary parts of the structure that were not directly related with the receptor areas of interest, and residues were renumbered. Next, each receptor with following models were evaluated. GPR139 models were superimposed to the cryo-EM structure of a class A GPCR (7VUH) from the Protein Data Bank (PDB) [4] and evaluated using the root mean standard deviation (RMSD) which ranged from 0.959-1.033 Å.

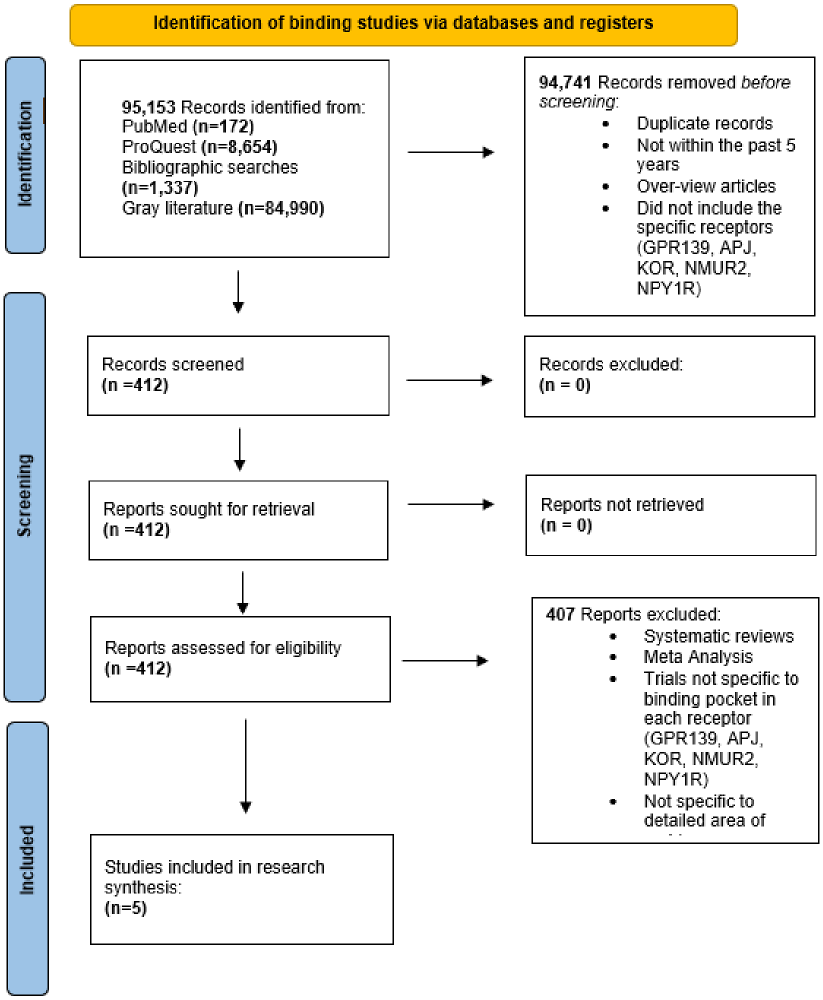

Literature searches (cf. **Figure 1**) were then employed to try and locate binding areas that may be of interest for each ligand and receptor pair.  This was challenging as there is not much information on binding areas of the receptors and decisions were made through human expertise and possible avenues of where to dock the ligands. For the receptor of GPR139, the Possible docking Residues were found to be F109, H187, W241, N271 [5] [6].

Once receptor models were predicted through AlphaFold 2, refined, and evaluated. Small molecules and peptidyl ligands were created with collaborating receptors and then docked to above possible residues, starting with GPR139. We employed flexible docking through AutoDock Vina 1.2.0. This is one of the most widely used program, with recent changes to be implemented as a means for more accurate docking. These new upgraded docking methods include, simultaneous multiple ligand docking, hydrated docking, and AutoDock4zn. [10] However, for the purpose of this project we employed the flexible docking which allowed the use to choose the residue location to dock the ligand on the receptor. This method yielded many results between the different models and a scoring method of accuracy was then assessed with docked models and used to choose the five best models per target. Our most challenging about the complex of GPR139 was to identify the potential binding residues for its ligand(s).

### GPCRdock-493 and GPCRdock-2001

This report covers both **2001** and **493** protocols

- Building the initial models of receptors (if receptors were modeled separately from ligands) or complexes. Did you use AlphaFold, RoseTTAFold, or any other deep-learning-based methodology for building the initial models? (if yes, please provide the info about the exact methodology and its implementation that you used). What other methods did you use, and how?

We used AlphaFold [1] (in ColabFold [2]) to build the initial models. To obtain more diverse protein structures, we used the standard AlphaFold, AlphaFold multimer, AF with and without templates, different MSA options among those present in ColabFold, different number of recursion cycles, AF with and without relaxation (use_amber option on / off), and also a dropout option from ColabFold-advanced.

**For APJ**, we used a mix of af2 relaxed and unrelaxed models, and added a known structure from PDB (5vbl) to the pool of protein models prepared for docking. More precisely, we used 5 unrelaxed models, 5 relaxed models, 5 relaxed models with "dropout" option turned on in ColabFold-advanced, and the 5vbl PDB structure. It was the same for both protocols 2001 and 493. After docking and rescoring, in top-5 for the protocol 2001 we've got af2-unrelaxed, 5vbl, af2-relaxed, af2-relaxed, af2-relaxed; in top-5 for the protocol 493 we've got af2-relaxed, 5vbl, af2-relaxed, af2-relaxed, af2-unrelaxed (top-1,2,3,4,5, respectively).

**For GPR139,** we generated models with and without disordered regions, for the case if those regions could somehow influence the stable part of the structure. At the docking stage we truncated them. The search over the AF parameters and options was not exhaustive. For the final TM-domain of the "chimeric" GPR139 built from two models, we used a relaxed AlphaFold model for the TM4-TM5 loop and an unrelaxed AlphaFold model for the region that we were expecting to bind with a Ca2+ ion. We then placed one / two ions into it and ran a separate MD relaxation on our side as described in the protocols file.

- Small molecule docking and/or peptide ligand docking (as applicable)

We generated the 3d ligand conformations in RDKit [3, 4], and parameterized them (along with protein models) for AutoDock Vina [5] in AutoDock Tools.

To generate docking poses, we used AutoDock Vina [5] with exhaustiveness 10 and max number of modes 25, and our in-house version of AutoDock Vina with built-in Convex-PL [6] as a scoring function that outputs 300 poses for subsequent re-scoring. We then re-scored them with Convex-PL-R [7] (**team 2001**) and KORP-PL$^{w}$ [8] (**team 493**). KORP-PL$^{w}$ does not rely on protein sidechains, which we expected to be beneficial for docking into the modeled protein structures. However, it turned out that Convex-PL-R predictions were better for GPR139.

- Any model refinement (if applicable)

**For APJ**, no

**For GPR139**, we used kpax [9] and pymol's align [10] to place the nanobody and all the domains with respect to each other. More precisely, we aligned the predicted models of the complexes of the sole TM domain, TM domain with Gα; TM domain with Gβ, and Gγ; TM domain with all G-proteins; and the nanobody with the 7f58 PDB structure, which had the same nanobody, Gβ, Gγ, and a homologous Gα.

Also, we decided to run docking with a structure that has the TM4-TM5 loop in an open conformation. However, for this model, the potential Ca ion binding site did not have any side chains suitable to coordinate the calcium ion. Thus, we cut the TM4-TM5 loop and merged it with two other TM domain structures that had ASP and GLU residues in the geometries suitable to bind the Ca ion, similarly to the PDB ids 7f58, 7piv, 6w25. We then placed 1-2 Ca ions into that site and ran a 10 ns equilibration molecular dynamics in an NPT ensemble of the TM domain with GROMACS [11].

- Criteria for selecting 5 best models per target

**For APJ**, we ran docking with all protein models for 3 intersecting binding boxes. Binding boxes were chosen based on the ligand location in the 5vbl structure. We then expanded one of the boxes so that it would cover the deeper part of the pocket. We then clustered the ligand poses with 1A RMSD cutoff. Each cluster was represented by the pose with the highest score. In the end, we selected the top-5 scored clusters.

**For GPR139**, structure preparation included more manual manipulation. By the end of the structure preparation, we had only two final GPR139 structures, in almost identical conformations. Although we performed docking to many other GPR139 models, only those from the final two structures were used in the final ranking. The 5 best models were selected with the same procedure as for APJ.

For the symmetry-aware clustering of ligand poses we used RDKit.

- Please include a section about target challenges. What did you find most challenging about each complex, and what information and approaches did you use to solve the challenge?

**For APJ,** we had an impression that the docking predictions did not look very good in terms of recovered interactions, but decided to stick to an automated pipeline without specific interventions. So this can not be called a "challenge".

**For GPR139**,

We found it quite challenging to build the correct interface between the TM domain, G-proteins, and the nanobody. It was mostly caused by the fact that nanobody is a synthetic protein with no MSA information available. As a result, AlphaFold was always placing the nanobody and sometimes the G-proteins on top of the extracellular part of the TM domain. We successfully placed the domains by manual alignment (in kpax and pymol) to the 7f58 structure with the same nanobody and its interface with G-proteins.

One of the hurdles of our GPR139 pipeline was the transmembrane region selection. As many <u>manual adjustments</u> should be done to the AlphaFold predictions for that target, we decided to choose only two TM domain conformations, to which all those adjustments were applied. These structures were then used for the final docking.

We were unsure whether the ligand docking score could be used to select the best TM domain conformation, as it often favors the structures with a higher number of protein-ligand contacts. Thus, we manually selected the structure with an open conformation of the TM4-TM5 loop, so that it would have fewer contacts and fewer steric conflicts with the ligand. Notably, that structure was actually the only “open” loop conformation we obtained with AlphaFold among many “closed” ones. It turned out that AlphaFold predictions were correct, as opposed to our intuition. The correct conformation was the “closed” one, with which we potentially could have obtained better docking predictions.

Based on the similarity between GPR139 and MC4R (PDB codes 7f58, 7piv, 6w25), we also expected a Ca ion to be present in the structure, probably having some interaction with the ligand, which might have introduced slight additional biases related to the docking binding box choice.

- Please comment on the role of experimental information and human expertise in generation and selection of your models (if applicable).

**For APJ**, we used PDB structures 5vbl, 6knm, along with Johnson et al [12], Ason et al [13] to select the binding pocket. We also used the 5vbl PDB structure for docking.

**For GPR139**, our predictions, binding pocket, and ion placement choices were influenced by Vedel et al [14]. In particular, we relied on the information about GPR139 being similar to MC4R mentioned in Vedel et al [14] and Nøhr et al [15]. Also, we checked Nehme et al [16] to better understand GA placement, and used 7f58, 7piv, 7piu, 7evw, 6w25 PDB structures for alignment to place the nanobody and improve the interdomain contacts. We used human expertise to connect the domains in GPR139 to each other, to place the CA ion, and to select the most promising model of the TM domain of GPR139.

#### GPCRdock-609 - APJ

This compound was docked into the model using SwissDock[3] webserver. Best pose was selected based on energy and relevant interactions found in literature.

The most challenging task was the selection of the conformer for such a large ligand. Moreover, we were not applying any constraints for the docking due to a lack of knowledge of specific interactions.

#### GPCRdock-609 – GPR139

Homology model of GPR139 with its $G_\alpha$ bounded was produced using Alpha Fold model multimer version[1]. Best ranked model was selected.

3D Conformation of the ligand was created using Gypsum_dl[2]. 100 conformers were produced with pH set between 6.4 and 8.4, and thoroughness set to 20. Finally, the variant with the lowest energy was selected.

The low energy conformer was further docked using SwissDock webserver[3]. Best pose was selected based on energy and relevant interactions found in literature.

The search for a low energy conformer was particularly challenging. The use of the multimer version of AlphaFold greatly improved the docking protocol, most notably to generate a GPCR in the active conformation.

### GPCRdock-609 - OPRK

The initial structure of the receptor was generated using AlphaFold multimer[1] with default settings putting the GPCR sequence with the respective Gα protein and the dynorphin. The pose with a minimum distance between the asp113 and the Nterm nitrogen atom from dynorphin was selected. The systems was then refined using molecular dynamics simulations.

Propka[2] was used to predict protonation states of the protein at a target pH 6.5. The orientation in the membrane was determined using OPM server[3]. The system was embedded into a POPC only model membrane using PACKMOL-memgen [4]. The simulation box was completed using TIP3P water molecules and neutralized using $K^+$ and $Cl^-$ ions with a final 0.15 M concentration. Molecular dynamics simulations were performed with sander.MPI and pmemd.cuda modules of AMBER18, with the ff14SB force field for the proteins, the lipid14 forcefield for the membrane.[5] ligand parameters were generated by antechamber with parameters taken from the General Amber Force Field 2 (gaff2). Bonds involving hydrogen atoms were constrained using the SHAKE algorithm and long-range electrostatic interactions were handled using Particle Mesh Ewald. The cut-off for non-bonded interaction was set to 10 Å. Each system was first minimized with AMBER sander.MPI module with 5000 steps of steepest descent algorithm then 5000 steps of conjugate gradient with a 10 $kcal \cdot mol^{-1} \cdot Å^2$ harmonic potential restraint on the protein part of the system. A second minimization run was done with restraints on the backbone only. The systems were then thermalized from 100 to 310 K for 10000 steps (restraining the protein and ligands with a 10 $kcal \cdot mol^{-1} \cdot Å^2$ harmonic potential). The system underwent 50000 steps of equilibration in the NPT ensemble and 1 bar (restraining the protein and ligands with a 10 $kcal \cdot mol^{-1} \cdot Å^2$ harmonic potential) before the production phase (100 ns). During equilibration and production phase, temperature was kept constant in the system at 310 K using a Langevin thermostat with a collision frequency of 5 $ps^{-1}$.

During all phases, a constraints between the carboxylate moiety from asp113 and the nitrogen atom from the Nterm of dynorphin was applied (5 $kcal \cdot mol^{-1} \cdot Å^2$).

<u>5 representative frames from the production phase were selected based on a k-means clustering analysis (RMSD of the ligand) using from ptraj with 100 iterations.</u>

For this target we benefited from the existence of the ionic bond between asp113 and the Nterm from the dynorphin. Moreover the analysis of the hydrophobicity revealed that the Cterm is more hydrophilic and thus subjected to be in the water phase. We then select pose with this specific orientation during the docking procedure.

### GPCRdock-609 – NMUR2

The initial structure of the receptor was generated using AlphaFold multimer[1] with default settings putting the GPCR sequence with the respective Gα protein and the NPY. The systems was then refined using molecular dynamics simulations.

Propka[2] was used to predict protonation states of the protein at a target pH 6.5. The orientation in the membrane was determined using OPM server[3]. The system was embedded into a POPC only model membrane using PACKMOL-memgen [4]. The simulation box was completed using TIP3P water molecules and neutralized using $K^+$ and $Cl^-$ ions with a final 0.15 M concentration. Molecular dynamics simulations were performed with sander.MPI and pmemd.cuda modules of AMBER18, with the ff14SB force field for the proteins, the lipid14 forcefield for the membrane.[5] ligand parameters were generated by antechamber with parameters taken from the General Amber Force Field 2 (gaff2). Bonds involving hydrogen atoms were constrained using the SHAKE algorithm and long-range electrostatic interactions were handled using Particle Mesh Ewald. The cut-off for non-bonded interaction was set to 10 Å. Each system was first minimized with AMBER sander.MPI module with 5000 steps of steepest descent algorithm then 5000 steps of conjugate gradient with a 10 $kcal \cdot mol^{-1} \cdot Å^2$ harmonic potential restraint on the protein part of the system. A second minimization run was done with restraints on the backbone only. The systems were then thermalized from 100 to 310 K for 10000 steps (restraining the protein and ligands with a 10 $kcal \cdot mol^{-1} \cdot Å^2$ harmonic potential). The system underwent 50000 steps of equilibration in the NPT ensemble and 1 bar (restraining the protein and ligands with a 10 $kcal \cdot mol^{-1} \cdot Å^2$ harmonic potential) before the production phase (200 ns). During equilibration and production phase, temperature was kept constant in the system at 310 K using a Langevin thermostat with a collision frequency of 5 $ps^{-1}$.

<u>5 representative frames from the production phase were selected based on a k-means clustering analysis (RMSD of the ligand) using from ptraj with 100 iterations.</u>

For this target we did not add a constraint on the peptide location because of the lack of knowledge. the analysis of the hydrophobicity revealed that the Cterm is more hydrophilic and thus subjected to be in the water phase. We then select pose with this specific orientation during the docking procedure.

#### GPCRdock-609 – NPY1R

The initial structure of the receptor was generated using AlphaFold multimer[1] with default settings putting the GPCR sequence with the respective Gα protein and the NPY. The systems was then refined using molecular dynamics simulations.

Propka[2] was used to predict protonation states of the protein at a target pH 6.5. The orientation in the membrane was determined using OPM server[3]. The system was embedded into a POPC only model membrane using PACKMOL-memgen [4]. The simulation box was completed using TIP3P water molecules and neutralized using $K^+$ and $Cl^-$ ions with a final 0.15 M concentration. Molecular dynamics simulations were performed with sander.MPI and pmemd.cuda modules of AMBER18, with the ff14SB force field for the proteins, the lipid14 forcefield for the membrane.[5] ligand parameters were generated by antechamber with parameters taken from the General Amber Force Field 2 (gaff2). Bonds involving hydrogen atoms were constrained using the SHAKE algorithm and long-range electrostatic interactions were handled using Particle Mesh Ewald. The cut-off for non-bonded interaction was set to 10 Å. Each system was first minimized with AMBER sander.MPI module with 5000 steps of steepest descent algorithm then 5000 steps of conjugate gradient with a 10 $kcal \cdot mol^{-1} \cdot Å^2$ harmonic potential restraint on the protein part of the system. A second minimization run was done with restraints on the backbone only. The systems were then thermalized from 100 to 310 K for 10000 steps (restraining the protein and ligands with a 10 $kcal \cdot mol^{-1} \cdot Å^2$ harmonic potential). The system underwent 50000 steps of equilibration in the NPT ensemble and 1 bar (restraining the protein and ligands with a 10 $kcal \cdot mol^{-1} \cdot Å^2$ harmonic potential) before the production phase (200 ns). During equilibration and production phase, temperature was kept constant in the system at 310 K using a Langevin thermostat with a collision frequency of 5 $ps^{-1}$.

5 representative frames from the production phase were selected based on a k-means clustering analysis (RMSD of the ligand) using from ptraj with 100 iterations.

For this target we did not add a constraint on the peptide location because of the lack of knowledge. the analysis of the hydrophobicity revealed that the Cterm is more hydrophilic and thus subjected to be in the water phase. We then select pose with this specific orientation during the docking procedure.

#### GPCRdock-861 - Methods

*Initial Receptor Model Construction*

Three protein structure prediction tools were used to generate diverse initial three-dimensional (3D) structures for the target G protein-coupled receptor (GPCR) sequences. The tools used were RosettaFold[1], RaptorX[2], and SwissModel[3], each accessed via their respective webservers. For each target, the top two predicted receptor models from RosettaFold and RaptorX were saved without alteration. Two homology models were built from different homologs when using SwissModel (Table 1). For APJ the homologs were the Apelin receptor and the C-C chemokine receptor type 5, and for GPR139 the homologs used were the Cholecystokinin 1 receptor (CCK1R) and the Cholecystokinin B receptor (CCKBR). This process resulted in six different initial receptor models per target GPCR.

**Table 1**

| Target | PDBId | GMQE | Similarity (%) | PDBId | GMQE | Similarity (%) |
|---|---|---|---|---|---|---|
| APJ | 5VBL[4] | 0.91 | 62 | 6AKY[5] | 0.68 | 40 |
| GPR139 | 7MBX[6] | 0.65 | 31 | 7F8V[7] | 0.64 | 31 |

*Preliminary Ligand Pose Generation*

Two molecular docking approaches, AutoDock Vina[8] and Glide[9,10], were used to generate candidate poses for the target receptor/ligand pairs using in-house python scripts. For Autdock Vina, ligand docking was attempted at each residue with the receptor held rigid, a 20 Å side length cubic box centered on the residue center of mass, and an exhaustiveness of 20. For Glide, each ligand structure was prepared with LigPrep[11] while each receptor model was prepared via the Protein Preparation Wizard[12]. For each receptor model, a docking grid was generated at the center of mass of each residue within the protein. This grid was constrained with an inner box of 10Å a side and an outer box of 30Å. As per standard docking protocol this inner box is used to place the ligand during the site-point search. The ligand can then drift within the confines of the outer box before assigning a score. Glide[9,10] standard precision (SP) docking was performed by the chosen ligand in each of the generated grids. The top five poses per grid were then stored and cross examined. The top two distinct ligand poses to any residue for each docking approach were saved as potential ligand-protein complexes. Docking was repeated for each of the six receptor models to result in 24 total models per receptor target.

*Building All-Atom Models in a Membrane Environment*

To run the molecular dynamics (MD) simulations described in the next section, complete structures with the same number of protein atoms were required for each of the 24 models. To achieve this, the missing segments from the SwissModel homology structures (solely residing in the termini at H1 and H7) were modeled as helixes and attached to the complexes using Schrödinger[11]. The resultant structures were then prepared for all-atom MD simulations using CHARMM-GUI[13], with membrane generation performed on only one of the models. The first model was aligned to the membrane using the Orientations of Proteins in Membranes PPM webserver[14] and the remaining models were then aligned to the membrane bound protein. The membrane used was a 3:1 molar ratio of POPC and cholesterol[15]. Explicit TIP3P water model and 100 mM NaCl resulted in initial box sizes of 95x95x112 $Å^3$ for APJ and 71x71x103 $Å^3$ for GPR139. These box sizes provided sufficient buffer zones across the periodic boundary conditions. The CHARMM-36m forcefield[16] was used for the protein and membrane while CGenFF[17] was used for the ligands.

*Temperature Replica Exchange Simulations to Assess Relative Complex Stability*

For a given target, the 24 ligand-receptor models were initially minimized and equilibrated using AMBER18[18]. For all simulations the Langevin thermostat with a coupling of 1 ps was used to keep the target temperature, initially at 310 K. The particle mesh Ewald (PME) technique was used for electrostatics and a 12 Å cut off and a switch at 10 Å was used for Lennard-Jones interactions. The semi-isotropic barostat was used to approximate sampling from the NPT ensemble during a 10 ns equilibration of the membrane and system density. After the systems equilibrated, the box for each of the 24 models was then set to the average size of the equilibrated 24 models enabling replica exchange simulations in the NVT ensemble. The systems underwent a short minimization and equilibration at the new box size before Temperature

Replica Exchange MD (TREMD) simulations were performed. For each target, three different TREMD runs were performed each with the 24 models spaced linearly from 310 K to $T_{max}$, ($T_{max}$ = 330 K for APJ, and $T_{max}$ = 340 K for GPR139). Exchanges between temperature rungs were attempted every 10 ps. The rungs were selected to achieve an acceptance frequency of 44 % for APJ and 42% for GPR139. The length of each individual simulation was 225 ns for APJ and 300 ns for GPR139 resulting in a totality of 16.2 μs for APJ and 21.6 μs for GPR139. Structures were saved every 20 ps for further analysis.

*Analysis of Simulations and Selection of Final Poses using Essential Dynamics Ensemble Docking*

The imaged protein and ligand coordinates in simulation frames at 310 K were extracted using CPPTRAJ[19]. The seven transmembrane helices (7TM), not including the more dynamic intracellular or extracellular loops, were then aligned to the first frame of the first of three replicas using VMD[20]. Next, inhouse python scripts using the MDAnalysis package[21] were used to find all residues whose side chain center of mass (COM) came within 3.0 Å of any ligand atom. Their residues were used to define the possible binding pockets of the receptor.

The xyz coordinates of the associated residue side chains were then saved to generate a protein dataset and concatenated with the atomic positions of the ligand to generate a protein-ligand dataset. Each position component was then normalized by its variance and further processed using Principal Component (PC) Analysis to extract the essential dynamics of either just the binding pocket or the binding pocket plus ligand (for the two datasets). Projection of the datasets onto the respective first two PC modes were then visualized and k-means clustering within the PYEMMA package[22] was used to partition the protein-only dataset and protein-ligand dataset. Three cluster centers were used for APJ and five were used for GPR139. For a given cluster the pairwise root mean squared deviation (RMSD) of the cartesian coordinates of the pocket (or pocket and ligand) were computed. The representative structure was selected as the structure that minimized the average RMSD from all other structures in the cluster. The top two most populated structures from the protein-ligand dataset were saved as potential final models. To generate three additional models for each target, the clustered structures obtained from only using the receptor pocket dataset were put through another round of docking. For each of the clusters, a grid was generated at the center of mass of the ligand, once again with a 10 Å inner box and 30 Å outer box. For each ligand, Glide extra precision (XP) docking[23] was performed. The top three poses with best XP glide scores were saved to complete the five final models for each GPCR target.

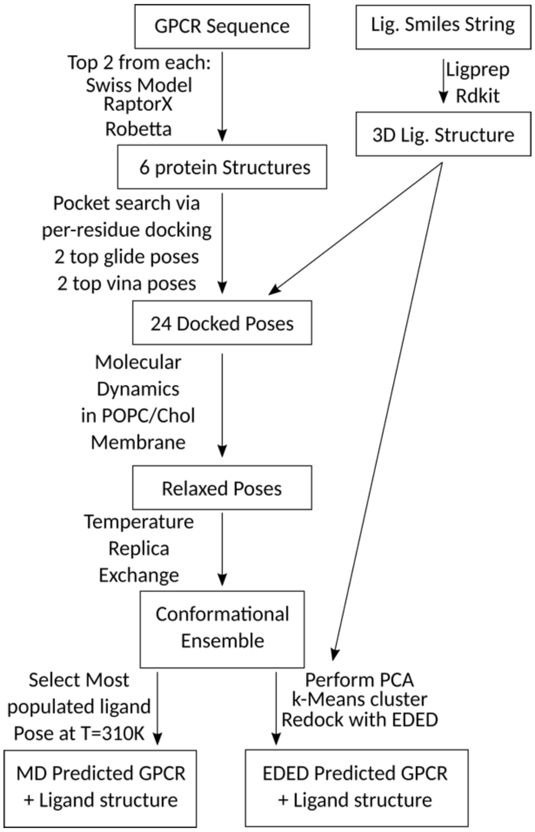

**Figure 1** Overview and workflow for our approach to GPCR and small molecule pose prediction.

#### GPCRdock-1245 - APJ

##### Molecular docking with conformer-dependent charge

For APJ, the 3D model of APJ was predicted using default parameters by AlphaFold2. And five predicted protein models were obtained according to pLDDT scores. The protein model with the highest plDDT score (plDDT=88.52) was selected for MDCC. And the protein model was prepared by using the *Protein Preparation Wizard* in Schrödinger 2018. The grid box of the protein for docking was centered at the (-2, -2, 7), to ensure that all residues in bold are within the grid box. Thirty lowest energy ligand conformers were generated for the small molecule ligand by *conformational search* in Schrödinger 2018 and then were considered in the geometry optimizations and vibrational frequencies using Gaussian 16. RESP charges were calculated for all conformers using Multiwfn3.8. These conformers with different RESP charges were docked into the prepared protein by using the *Glide* module, and the binding energies were scored and ranked by the Glide XP scoring mode. And top five models of the result using Glide XP were redocked and reordered by QM-polarized Ligand Docking. Finally, the top five models for docking scores of all results were submitted, and two of the five models were in good agreement with the experimental structure.

### GPCRdock-1245 – GPR139

#### Molecular docking with conformer-dependent charge

For GPR139, the same approach was used for model prediction as APJ. And five predicted protein models were also obtained according to pLDDT scores. The difference from APJ was the selection of the models. The protein models with the top two pLDDT scores (plDDT=80.16, plDDT=79.90) were selected for GPR139. And these protein models were prepared in the same way as APJ. The grid boxes of the proteins were centered at the (-9, 7, 4) and (-8, 0, 4) respectively for the top two models. The preparation process of the small molecule ligand was the same as APJ. All conformers with different RESP charges were docked into the two prepared proteins using Glide XP. And then, top one model of the first protein (plDDT=80.16) and the top five models of the second protein (plDDT=79.90) using Glide XP were redocked and reordered by QM-polarized Ligand Docking. Finally, five models were submitted according to the docking scores of all results, among which one model for the protein with the pLDDT score of 80.16 and four models for the protein with the pLDDT score of 79.90.

### GPCRdock-1912

#### GPCR-ligand docking with predicted structure and ligand-binding site

**Hong Wei** [1], **Jianyi Yang** [2,*]

[1]School of Mathematical Sciences, Nankai University, Tianjin 300071, China, [2]Research Center for Mathematics and Interdisciplinary Sciences, Shandong University, Qingdao 266237, China.

For the GPCR Dock 2021 experiments, we built the GPCR-ligand complex structures using predicted structure and ligand-binding site.

First, based on the rapid inter-residue distance and orientation predictions in trRosettaX [1], an optimal multiple sequence alignment (MSA) is selected from a set of MSAs generated at multiple *e*-value and coverage cutoffs in HHblits [3]. This MSA is fed into AlphaFold2 [2] to predict the GPCR structure with default settings.

For GPCR-peptide models, AlphaFold-multimer was used to build the initial complex models. The models were then refined with Rosetta FlexPepDock. The final models were selected based on consistency with the binding site predicted by COACH-D.

For GPCR-small-molecule complexes, the predicted structure model was submitted to the COACH-D [4] server to predict the ligand-binding site and initial GPCR-ligand complex structure models. COACH-D predicts ligand-binding sites and complex structures using homologous protein-ligand templates. With the predicted GPCR structure models and the ligand-binding site, the Rosetta Ligand Docking Protocol [5] is then applied to dock the ligand into the binding site region of the predicted GPCR structure models. During docking, the geometric center of the binding pocket is calculated as the mean of the coordinates of the predicted ligand-binding residues. A total of 1000 poses are generated, which are ranked based the energy scores and the consistency with the predicted binding site. Steric clashes are also taken into account. The top five poses are then submitted.

No special optimization/relaxation was applies to either the initial AlphaFold structures or the docked complexes.

### GPCRdock-2223 - APJ

#### Homology Modeling

Two different homology models were used to predict ligand binding mode. The first model (MMS_APJ_1) was retrieved from the AlphaFold[1] database, while the second one (MMS_APJ_2) was generated as stated in the following lines:

- The sequence of APJ receptor (UniProt ID: P35414) was retrieved in the fasta format from the UniProt database.
- The fasta sequence was uploaded to the PHYRE2[2] web server, and a normal mode search was performed to fetch possible templates. The structure deposited in the Protein Data Bank with accession code 5VBL[3] (Structure of Apelin receptor in complex with agonist peptide) was chosen as a template for the generation of the homology model.
- The sequence alignment between the template and the APJ receptor was imported into Molecular Operating Environment (MOE[4]) 2019.01, alongside the template structure (5VBL) retrieved from the PDB. The chimeric part of the construct (from Cys1006 to Lys1046) was removed. The rest of the structure was prepared exploiting the Structure Preparation and Protonate3D tools (removal of crystallographic water molecules, ions, and other solvent molecules, selection of the highest occupancy for each residue, assignment of the correct protonation state at pH 7.4). The Homology Model module of MOE was then used to generate the homology model, using the sequence alignment retrieved from the PHYRE2 webserver and the 5VBL structure, prepared as described before. The best model, according to MOE, was then manually refined, checking for consistency with the predicted secondary structure (according to both PHYRE2 and JPRED4[5] web server prediction). The missing intracellular loop 3 (ICL3), was then manually reconstructed as follows: residue 238 to 243 were built in α-helix conformation using the Protein Builder module of MOE, while residues 229 to 237 were modeled ab-initio using the Loop Modeler module of MOE.

MMS_APJ_2 was then refined through classical Molecular Dynamics simulations. At first, the model was aligned to the OPM file of the template (retrieved from the Orientation of Protein in Membrane database). Subsequently, system preparation for the molecular dynamics simulations was carried out using VMD[6]. The protein was embedded in a lipid bilayer consisting of phosphatidylcholine (POPC) units and explicitly solvated in a TIP3P[7] water box with the borders placed at 15 Å from any protein atom. The system charge was neutralized to a concentration of 0.154 M using $Na^+$/$Cl^-$.

Molecular Dynamics simulations were performed with the CHARMM36[8] force field using the ACEMD3[9] engine, which is based upon OpenMM7[10].

Before MD simulation, 1500 steps of energy minimization with the conjugate-gradient algorithm were performed to remove clashes and bad contacts among the system atoms. Subsequently, a three-stage system equilibration was performed. The first equilibration step consisted of 5 ns of NPT simulation with harmonic positional restraints of 5 kcal $mol^{-1}Å^{-2}$ on each atom of the protein and the lipid bilayer. The second step consisted of 10 ns of NPT simulation with harmonic positional restraints of 5 kcal $mol^{-1}Å^{-2}$ only on each protein atom and on the phosphorus atom of the POPC units, the third step consisted of 5 ns of NPT simulation with harmonic positional restraints of 5 kcal $mol^{-1}Å^{-2}$ only on the alpha carbons of the protein. During the equilibration stages, the temperature was maintained at 310 K using the Langevin thermostat, with low dumping of 1 $ps^{-1}$, while the pressure was set at 1 atm using a Monte Carlo barostat. The particle-mesh Ewald (PME[11]) method was used to calculate the electrostatic interactions with a 1 Å grid. A 9.0 Å cutoff was applied for long-term interactions. The M-SHAKE algorithm was applied to constrain the bond lengths involving hydrogen atoms.

After the equilibration stages, a productive 250 ns MD simulation was carried out in the NVT ensemble (T = 310 K), using the same setup as for the system equilibration stage except for the absence of the barostat and the restraints.

The trajectory was then clustered based on the pairwise RMSD matrix of the backbone using the TTCLUST[12] Python program, using the default parameters. The representative frame of the most populated cluster was then used for subsequent docking calculations.

#### Docking

The ligand for the APJ receptor was properly prepared for docking calculations using tools from the OpenEye suite. Starting from the SMILES of the molecule, its most probable tautomeric state was highlighted with the "Tautomers" program, then the three-dimensional coordinates were computed with "Omega", the partial charges were assigned with "MolCharge" (using MMFF as the method) and, finally, the highest relevant protomeric state was picked exploiting the "FixpKa" tool[13].

The receptors used for the calculations were MMS_APJ_1 and MMS_APJ_2. For each of these structures, all passages listed down below were carried out.

Molecular docking calculations were executed with two different docking programs, GOLD[14] (based on a genetic algorithm, developed and licensed by CCDC) and Glide[15] (a systematic algorithm released by Schrödinger).

For the docking calculations with GOLD, the center of mass for the binding site was chosen based on some of the most important residues for ligand binding, which were retrieved from literature (mainly from the article accompanying the PDB crystal 5VBL, which represents the binding event of APJ receptor with an agonist peptide[3]). The residues used to determine the binding site center were: Trp85, Tyr264, Lys268, Phe291, and Tyr299. The radius of the binding pocket was set at 20 Å. The docking was run using the "chemscore" scoring function implemented in GOLD. For the ligand of interest, 50 poses were obtained, and those were then filtered to eliminate the ones presenting clashes with the protein and/or unfavorable electrostatic contacts. For each of the remaining conformations, the difference between the energy of that specific pose and the energy of the local minimum was computed, allowing to pick the conformations with the lowest strain. Among the final solutions, the selection was executed based on visual inspection, giving higher relevance to the poses presenting the interactions described in the literature for APJ agonists. The residues considered for the visual inspection selection were: Trp85, Tyr88, Tyr93, Trp195, Tyr264, Lys268, Tyr271, Phe291, and Tyr299.

For the calculations involving Glide, the receptor grid for docking was built using as the center of mass the same coordinates adopted for the GOLD run. The docking run was executed with Glide-SP protocol, which implements GlideScore as the scoring function. In this case, also, 50 poses were produced, and those were filtered to eliminate unfavorable steric and electrostatic contacts with the target. As it was done for GOLD, the selection among the remaining conformations was executed first by internal strain energy and then following the indications given by literature about the most important residues that are involved in the ligand-receptor interaction for APJ (the same listed above for GOLD).

Among the conformations obtained with the two protocols, the final poses to submit to the post-docking phase were selected applying a consensus approach between the best solutions given by the two programs.

#### Post-docking

Each docking pose was refined through classical Molecular Dynamics simulations. At first, the model was aligned to the OPM file of the template (retrieved from the Orientation of Protein in Membrane database). Subsequently, system preparation for the molecular dynamics simulations was carried out using VMD. The complex was embedded in a lipid bilayer consisting of phosphatidylcholine (POPC) units and explicitly solvated in a TIP3P water box with the borders placed at 15 Å from any protein atom. The system charge was neutralized to a concentration of 0.154 M using $Na^+/Cl^-$.

Molecular Dynamics simulations were performed with the CHARMM36 force field using the ACEMD3 engine. Ligand parameters were retrieved from Paramchem[16], a web interface for the assignment of parameters based on the CGenFF[17] force field.

Before MD simulation, 1500 steps of energy minimization with the conjugate-gradient algorithm were performed to remove clashes and bad contacts among the system atoms. Subsequently, a three-stage system equilibration was performed. The first equilibration step consisted of 5 ns of NPT simulation with harmonic positional restraints of 5 kcal $mol^{-1}Å^{-2}$ on each atom of the protein, the ligand, and the lipid bilayer. The second step consisted of 10 ns of NPT simulation with harmonic positional restraints of 5 kcal $mol^{-1}Å^{-2}$ only on each protein and ligand atom and on the phosphorus atom of the POPC units, the third step consisted of 5 ns of NPT simulation with harmonic positional restraints of 5 kcal $mol^{-1}Å^{-2}$ only on the alpha carbons of the protein and on each ligand atom. During the equilibration stages, the temperature was maintained at 310 K using the Langevin thermostat, with low dumping of 1 $ps^{-1}$, while the pressure was set at 1 atm using a Monte Carlo barostat. The particle-mesh Ewald (PME) method was used to calculate the electrostatic interactions with a 1 Å grid. A 9.0 Å cutoff was applied for long-term interactions. The M-SHAKE algorithm was applied to constrain the bond lengths involving hydrogen atoms.

After the equilibration stages, three independent productive 25 ns MD simulations were carried out in the NVT ensemble (T = 310 K), using the same setup as for the system equilibration stage except for the absence of the barostat and the restraints.

The trajectories were then clustered based on the pairwise RMSD matrix of the ligand using the TTCLUST Python program, using the default parameters. The final pose selection was based on visual inspection of the most representative frames of each cluster.

#### Results summary

| Submitted complex | Model | Docking method |
|---|---|---|
| APJ_GPCRdock-2223-1 | MMS_APJ_2 | Glide |
| APJ_GPCRdock-2223-2 | MMS_APJ_1 | Glide |
| APJ_GPCRdock-2223-3 | MMS_APJ_2 | Glide |
| APJ_GPCRdock-2223-4 | MMS_APJ_2 | Glide |
| APJ_GPCRdock-2223-5 | MMS_APJ_1 | Glide |

### GPCRdock-2223 - GPR139

#### Homology Modeling

Two different homology models were used to predict ligand binding mode. The first model (MMS_GPR139_1) was retrieved from the AlphaFold[1] database, while the second one (MMS_GPR139_2) was generated as stated in the following lines:

- The sequence of GPR139 receptor (UniProt ID: Q6DWJ6) was retrieved in the fasta format from the UniProt database.
- The fasta sequence was uploaded to the PHYRE2[2] web server, and a normal mode search was performed to fetch possible templates. The structure deposited in the Protein Data Bank with accession code 6TPN[3] (Crystal structure of the Orexin-2 receptor in complex with HLT6641) was chosen as a template for the generation of the homology model.
- The sequence alignment between the template and the GPR139 receptor was imported into Molecular Operating Environment (MOE[4]) 2019.01, alongside the template structure (6TPN) retrieved from the PDB. The chimeric part of the construct (from Gly1001 to Lys1189) was removed. The rest of the structure was prepared exploiting the Structure Preparation and Protonate3D tools (removal of crystallographic water molecules, ions, and other solvent molecules, selection of the highest occupancy for each residue, assignment of the correct protonation state at pH 7.4). The Homology Model module of MOE was then used to generate the homology model, using the sequence alignment retrieved from the PHYRE2 webserver and the 6TPN structure, prepared as described before. The best model, according to MOE, was then manually refined, checking for consistency with the predicted secondary structure (according to both PHYRE2 and JPRED4[5] web server prediction). The missing intracellular loop 3 (ICL3), was then manually reconstructed as follows: residue 209 to 212 were built in α-helix conformation using the Protein Builder module of MOE, while residues 213 to 218 were modeled ab-initio using the Loop Modeler module of MOE.

MMS_GPR139_2 was then refined through classical Molecular Dynamics simulations. At first, the model was aligned to the OPM file of the template (retrieved from the Orientation of Protein in Membrane database). Subsequently, system preparation for the molecular dynamics simulations was carried out using VMD[6]. The protein was embedded in a lipid bilayer consisting of phosphatidylcholine (POPC) units and explicitly solvated in a TIP3P[7] water box with the borders placed at 15 Å from any protein atom. The system charge was neutralized to a concentration of 0.154 M using $Na^+/Cl^-$.

Molecular Dynamics simulations were performed with the CHARMM36[8] force field using the ACEMD3[9] engine, which is based upon OpenMM7[10].

Before MD simulation, 1500 steps of energy minimization with the conjugate-gradient algorithm were performed to remove clashes and bad contacts among the system atoms. Subsequently, a three-stage system equilibration was performed. The first equilibration step consisted of 5 ns of NPT simulation with harmonic positional restraints of 5 kcal $mol^{-1}Å^{-2}$ on each atom of the protein and the lipid bilayer. The second step consisted of 10 ns of NPT simulation with harmonic positional restraints of 5 kcal $mol^{-1}Å^{-2}$ only on each protein atom and on the phosphorus atom of the POPC units, the third step consisted of 5 ns of NPT simulation with harmonic positional restraints of 5 kcal $mol^{-1}Å^{-2}$ only on the alpha carbons of the protein. During the equilibration stages, the temperature was maintained at 310 K using the Langevin thermostat, with low dumping of 1 $ps^{-1}$, while the pressure was set at 1 atm using a Monte Carlo barostat. The particle-mesh Ewald (PME[11]) method was used to calculate the electrostatic interactions with a 1 Å grid. A 9.0 Å cutoff was applied for long-term interactions. The M-SHAKE algorithm was applied to constrain the bond lengths involving hydrogen atoms.

After the equilibration stages, a productive 250 ns MD simulation was carried out in the NVT ensemble (T = 310 K), using the same setup as for the system equilibration stage except for the absence of the barostat and the restraints.

The trajectory was then clustered based on the pairwise RMSD matrix of the backbone using the TTCLUST[12] Python program, using the default parameters. The representative frame of the most populated cluster was then used for subsequent docking calculations.

#### Docking

The ligand for the GPR139 receptor was properly prepared for docking calculations using tools from the OpenEye suite. Starting from the SMILES of the molecule, its most probable tautomeric state was highlighted with the "Tautomers" program, then the three-dimensional coordinates were computed with "Omega", the partial charges were assigned with "MolCharge" (using MMFF as the method) and, finally, the highest relevant protomeric state was picked exploiting the "FixpKa" tool[13].

The receptors used for the calculations were MMS_GPR139_1 and MMS_GPR139_2. For each of these structures, all passages listed down below were carried out.

Molecular docking calculations were executed with two different docking programs, GOLD[14] (based on a genetic algorithm, developed and licensed by CCDC) and Glide[15] (a systematic algorithm released by Schrödinger).

For the docking calculations with GOLD, the center of mass for the binding site was chosen based on some of the most important residues for ligand binding, which were retrieved from literature (mainly from the minireview from Vedel et al.[16]). The residues used for the binding site center were: Ile80, Phe109, and Trp241. The radius of the binding pocket was set at 20 Å. The docking was run using the "chemscore" scoring function implemented in GOLD. For the ligand of interest, 50 poses were obtained, and those were then filtered to eliminate the ones presenting clashes with the protein and/or unfavorable electrostatic contacts. For each of the remaining conformations, the difference between the energy of that specific pose and the energy of the local minimum was computed, allowing to pick the conformations with the lowest strain. Among the final solutions, the selection was executed based on visual inspection, giving higher relevance to the poses presenting the interactions described in the literature for GPR139 agonists. The residues considered for the visual inspection selection were: Val76, Phe79, Ile80, Val83, Ile104, Glu108, Phe109, His187, Trp241, and Arg244.

For the calculations involving Glide, the receptor grid for docking was built using as the center of mass the same coordinates adopted for the GOLD run. The docking run was executed with Glide-SP protocol, which implements GlideScore as the scoring function. In this case, also, 50 poses were produced, and those were filtered to eliminate unfavorable steric and electrostatic contacts with the target. As it was done for GOLD, the selection among the remaining conformations was executed first by internal strain energy and then following the indications given by literature about the most important residues that are involved in the ligand-receptor interaction for GPR139 (the same listed above for GOLD).

Among the conformations obtained with the two protocols, the final poses to submit to the post-docking phase were selected applying a consensus approach between the best solutions given by the two programs.

#### Post-docking

Each docking pose was refined through classical Molecular Dynamics simulations. At first, the model was aligned to the OPM file of the template (retrieved from the Orientation of Protein in Membrane database). Subsequently, system preparation for the molecular dynamics simulations was carried out using VMD. The complex was embedded in a lipid bilayer consisting of phosphatidylcholine (POPC) units and explicitly solvated in a TIP3P water box with the borders placed at 15 Å from any protein atom. The system charge was neutralized to a concentration of 0.154 M using $Na^+/Cl^-$.

Molecular Dynamics simulations were performed with the CHARMM36 force field using the ACEMD3 engine. Ligand parameters were retrieved from Paramchem[17], a web interface for the assignment of parameters based on the CGenFF[18] force field.

Before MD simulation, 1500 steps of energy minimization with the conjugate-gradient algorithm were performed to remove clashes and bad contacts among the system atoms. Subsequently, a three-stage system equilibration was performed. The first equilibration step consisted of 5 ns of NPT simulation with harmonic positional restraints of 5 kcal $mol^{-1}Å^{-2}$ on each atom of the protein, the ligand, and the lipid bilayer. The second step consisted of 10 ns of NPT simulation with harmonic positional restraints of 5 kcal $mol^{-1}Å^{-2}$ only on each protein and ligand atom and on the phosphorus atom of the POPC units, the third step consisted of 5 ns of NPT simulation with harmonic positional restraints of 5 kcal $mol^{-1}Å^{-2}$ only on the alpha carbons of the protein and on each ligand atom. During the equilibration stages, the temperature was maintained at 310 K using the Langevin thermostat, with low dumping of 1 $ps^{-1}$, while the pressure was set at 1 atm using a Monte Carlo barostat. The particle-mesh Ewald (PME) method was used to calculate the electrostatic interactions with a 1 Å grid. A 9.0 Å cutoff was applied for long-term interactions. The M-SHAKE algorithm was applied to constrain the bond lengths involving hydrogen atoms.

After the equilibration stages, three independent productive 25 ns MD simulations were carried out in the NVT ensemble (T = 310 K), using the same setup as for the system equilibration stage except for the absence of the barostat and the restraints.

The trajectories were then clustered based on the pairwise RMSD matrix of the ligand using the TTCLUST Python program, using the default parameters. The final pose selection was based on visual inspection of the most representative frames of each cluster.

##### Results summary

| Submitted complex | Model | Docking method |
|---|---|---|
| GPR139_GPCRdock-2223-1 | MMS_GPR139_1 | Gold |
| GPR139_GPCRdock-2223-2 | MMS_GPR139_1 | Glide |
| GPR139_GPCRdock-2223-3 | MMS_GPR139_1 | Gold |
| GPR139_GPCRdock-2223-4 | MMS_GPR139_1 | Gold |
| GPR139_GPCRdock-2223-5 | MMS_GPR139_2 | Glide |

### GPCRdock-2223 - KOR

#### Homology Modeling

Two different homology models were used to predict ligand binding mode. The first model (MMS_KOR_1) was retrieved from the AlphaFold[1] database, while the second one (MMS_KOR_2) was generated as stated in the following lines:

- The sequence of KOR receptor (UniProt ID: P41145) was retrieved in the fasta format from the UniProt database.
- The fasta sequence was uploaded to the PHYRE2[2] web server, and a normal mode search was performed to fetch possible templates. The structure deposited in the Protein Data Bank with accession code 6B73[3] (Crystal structure of a nanobody-stabilized active state of the kappa-opioid receptor) was chosen as a template for the generation of the homology model.
- The sequence alignment between the template and the KOR receptor was imported into Molecular Operating Environment (MOE[4]) 2019.01, alongside the template structure (6B73) retrieved from the PDB. The receptor chain of the structure was prepared exploiting the Structure Preparation and Protonate3D tools (removal of crystallographic water molecules, ions, and other solvent molecules, selection of the highest occupancy for each residue, assignment of the correct protonation state at pH 7.4). The Homology Model module of MOE was then used to generate the homology model, using the sequence alignment retrieved from the PHYRE2 webserver and the 6B73 structure, prepared as described before. The best model, according to MOE, was saved and utilized for further calculations.

MMS_KOR_2 was then refined through classical Molecular Dynamics simulations. At first, the model was aligned to the OPM file of the template (retrieved from the Orientation of Protein in Membrane database). Subsequently, system preparation for the molecular dynamics simulations was carried out using VMD[5]. The protein was embedded in a lipid bilayer consisting of phosphatidylcholine (POPC) units and explicitly solvated in a TIP3P[6] water box with the borders placed at 15 Å from any protein atom. The system charge was neutralized to a concentration of 0.154 M using $Na^+/Cl^-$.

Molecular Dynamics simulations were performed with the CHARMM36[7] force field using the ACEMD3[8] engine, which is based upon OpenMM7[9].

Before MD simulation, 1500 steps of energy minimization with the conjugate-gradient algorithm were performed to remove clashes and bad contacts among the system atoms. Subsequently, a three-stage system equilibration was performed. The first equilibration step consisted of 5 ns of NPT simulation with harmonic positional restraints of 5 kcal $mol^{-1}Å^{-2}$ on each atom of the protein and the lipid bilayer. The second step consisted of 10 ns of NPT simulation with harmonic positional restraints of 5 kcal $mol^{-1}Å^{-2}$ only on each protein atom and on the phosphorus atom of the POPC units, the third step consisted of 5 ns of NPT simulation with harmonic positional restraints of 5 kcal $mol^{-1}Å^{-2}$ only on the alpha carbons of the protein. During the equilibration stages, the temperature was maintained at 310 K using the Langevin thermostat, with low dumping of 1 $ps^{-1}$, while the pressure was set at 1 atm using a Monte Carlo barostat. The particle-mesh Ewald (PME[10]) method was used to calculate the electrostatic interactions with a 1 Å grid. A 9.0 Å cutoff was applied for long-term interactions. The M-SHAKE algorithm was applied to constrain the bond lengths involving hydrogen atoms.

After the equilibration stages, a productive 250 ns MD simulation was carried out in the NVT ensemble (T = 310 K), using the same setup as for the system equilibration stage except for the absence of the barostat and the restraints.

The trajectory was then clustered based on the pairwise RMSD matrix of the backbone using the TTCLUST[11] Python program, using the default parameters. The representative frame of the most populated cluster was then used for subsequent docking calculations.

#### Docking

For the peptide Dynorphin, a three-dimensional structure of the conformation bound to the KOR receptor is available in the Protein Data Bank (PDB code: 2N2F[12]). We chose to exploit this data, avoiding using computational methods that would distort Dynorphin in its bound form.

The receptors used for the calculations were MMS_KOR_1 and MMS_KOR_2. For each of these structures, all passages listed down below were carried out.

The method adopted was the "Ligand-based homology modelling[13]", which is based on the rearrangement of the KOR models created around Dynorphin in its bound form. This way, the three-dimensional conformation of the ligand is preserved, and the receptor adapts its internal volume to receive the ligand inside the transmembrane (TM) region.

First, the ligand was manually placed inside the TM region, ensuring that the positioning and the orientation of the ligand inside the receptor were in agreement with literature data about the KOR-Dynorphin interactions (coming both from the work of O'Connor et al.[12] and from the article of Che et al.[3]). The residues considered for the placement of the ligand were: Asp138, Met142, Lys227, Trp287, Tyr312, Gly319, Tyr320.

After the manual ligand positioning, the KOR receptor was rebuilt using the Homology modelling tool implemented in MOE, specifying to the program to consider the presence of the ligand in that specific form while creating the models. The fasta of KOR protein was modeled using as templates first the MMS_KOR_1 structure, and then the MMS_KOR_2 structure. For each template, 10 models were generated, and those were visually inspected to select the ones presenting the ligand in the most appropriate position (avoiding clashes or unfavorable contacts with the receptor) inside the TM region, also rewarding the models in which the shift of the protein structure was minimal in respect to the template.

The best Dynorphin-KOR complexes were chosen both for MMS_KOR_1 and for MMS_KOR_2. Those systems were then submitted to the post-docking phase.

#### Post-docking

Each docking pose was refined through classical Molecular Dynamics simulations. At first, the model was aligned to the OPM file of the template (retrieved from the Orientation of Protein in Membrane database). Subsequently, system preparation for the molecular dynamics simulations was carried out using VMD. The complex was embedded in a lipid bilayer consisting of phosphatidylcholine (POPC) units and explicitly solvated in a TIP3P water box with the borders placed at 15 Å from any protein atom. The system charge was neutralized to a concentration of 0.154 M using $Na^+/Cl^-$.

Molecular Dynamics simulations were performed with the CHARMM36 force field using the ACEMD3 engine.

Before MD simulation, 1500 steps of energy minimization with the conjugate-gradient algorithm were performed to remove clashes and bad contacts among the system atoms. Subsequently, a three-stage system equilibration was performed. The first equilibration step consisted of 5 ns of NPT simulation with harmonic positional restraints of 5 kcal $mol^{-1}Å^{-2}$ on each atom of the protein, the ligand, and the lipid bilayer. The second step consisted of 10 ns of NPT simulation with harmonic positional restraints of 5 kcal $mol^{-1}Å^{-2}$ only on each protein and ligand atom and on the phosphorus atom of the POPC units, the third step consisted of 5 ns of NPT simulation with harmonic positional restraints of 5 kcal $mol^{-1}Å^{-2}$ only on the alpha carbons of the protein and on each ligand atom. During the equilibration stages, the temperature was maintained at 310 K using the Langevin thermostat, with low dumping of 1 $ps^{-1}$, while the pressure was set at 1 atm using a Monte

Carlo barostat. The particle-mesh Ewald (PME) method was used to calculate the electrostatic interactions with a 1 Å grid. A 9.0 Å cutoff was applied for long-term interactions. The M-SHAKE algorithm was applied to constrain the bond lengths involving hydrogen atoms.

After the equilibration stages, three independent productive 25 ns MD simulations were carried out in the NVT ensemble (T = 310 K), using the same setup as for the system equilibration stage except for the absence of the barostat and the restraints.

The trajectories were then clustered based on the pairwise RMSD matrix of the ligand backbone using the TTCLUST Python program, using the default parameters. The final pose selection was based on visual inspection of the most representative frames of each cluster.

**Results summary**

| Submitted complex | Model | Docking method |
|---|---|---|
| KOR_GPCRdock-2223-1 | MMS_KOR_2 | Ligand-based Homology Modeling |
| KOR_GPCRdock-2223-2 | MMS_KOR_2 | Ligand-based Homology Modeling |
| KOR_GPCRdock-2223-3 | MMS_KOR_2 | Ligand-based Homology Modeling |
| KOR_GPCRdock-2223-4 | MMS_KOR_1 | Ligand-based Homology Modeling |
| KOR_GPCRdock-2223-5 | MMS_KOR_2 | Ligand-based Homology Modeling |

### GPCRdock-2223 - NMUR2

#### Homology Modeling

Two different homology models were used to predict ligand binding mode. The first model (MMS_NMUR2_1) was retrieved from the AlphaFold[1] database, while the second one (MMS_NMUR2_2) was generated as stated in the following lines:

- The sequence of NMUR2 receptor (UniProt ID: Q9GZQ4) was retrieved in the fasta format from the UniProt database.
- The fasta sequence was uploaded to the PHYRE2[2] web server, and a normal mode search was performed to fetch possible templates. The structure deposited in the Protein Data Bank with accession code 2RH1[3] (High resolution crystal structure of human B2-adrenergic G protein-coupled receptor) was chosen as a template for the generation of the homology model.
- The sequence alignment between the template and the NNMUR2 receptor was imported into Molecular Operating Environment (MOE[4]) 2019.01, alongside the template structure (2RH1) retrieved from the PDB. The chimeric part of the construct (from Asn1002 to Tyr1161) was removed. The rest of the structure was prepared exploiting the Structure Preparation and Protonate3D tools (removal of crystallographic water molecules, ions, and other solvent molecules, selection of the highest occupancy for each residue, assignment of the correct protonation state at pH 7.4). The Homology Model module of MOE was then used to generate the homology model, using the sequence alignment retrieved from the PHYRE2 webserver and the 2RH1 structure, prepared as described before. The best model, according to MOE, was saved and utilized for further calculations.

MMS_NMUR2_2 was then refined through classical Molecular Dynamics simulations. At first, the model was aligned to the OPM file of the template (retrieved from the Orientation of Protein in Membrane database). Subsequently, system preparation for the molecular dynamics simulations was carried out using VMD[5]. The protein was embedded in a lipid bilayer consisting of phosphatidylcholine (POPC) units and explicitly solvated in a TIP3P[6] water box with the borders placed at 15 Å from any protein atom. The system charge was neutralized to a concentration of 0.154 M using $Na^+/Cl^-$.

Molecular Dynamics simulations were performed with the CHARMM36[7] force field using the ACEMD3[8] engine, which is based upon OpenMM7[9].

Before MD simulation, 1500 steps of energy minimization with the conjugate-gradient algorithm were performed to remove clashes and bad contacts among the system atoms. Subsequently, a three-stage system equilibration was performed. The first equilibration step consisted of 5 ns of NPT simulation with harmonic positional restraints of 5 kcal $mol^{-1}Å^{-2}$ on each atom of the protein and the lipid bilayer. The second step consisted of 10 ns of NPT simulation with harmonic positional restraints of 5 kcal $mol^{-1}Å^{-2}$ only on each protein atom and on the phosphorus atom of the POPC units, the third step consisted of 5 ns of NPT simulation with harmonic positional restraints of 5 kcal $mol^{-1}Å^{-2}$ only on the alpha carbons of the protein. During the equilibration stages, the temperature was maintained at 310 K using the Langevin thermostat, with low dumping of 1 $ps^{-1}$, while the pressure was set at 1 atm using a Monte Carlo barostat. The particle-mesh Ewald (PME[10]) method was used to calculate the electrostatic interactions with a 1 Å grid. A 9.0 Å cutoff was applied for long-term interactions. The M-SHAKE algorithm was applied to constrain the bond lengths involving hydrogen atoms.

After the equilibration stages, a productive 250 ns MD simulation was carried out in the NVT ensemble (T = 310 K), using the same setup as for the system equilibration stage except for the absence of the barostat and the restraints.

The trajectory was then clustered based on the pairwise RMSD matrix of the backbone using the TTCLUST[11] Python program, using the default parameters. The representative frame of the most populated cluster was then used for subsequent docking calculations.

#### Docking

The peptide ligand for the NMUR2 receptor (NMU25) was properly prepared for docking calculations using the MOE suite. The fasta sequence of the peptide was passed through the "Protein builder" tool, allowing to obtain the three-dimensional structure of the ligand. Then, the "Protonate 3D" program was implemented to assign the proper protonation state to each residue at a defined pH of 7.4. Finally, all the hydrogens in the system were minimized under the AMBER10:EHT[12] force field implemented in MOE.

Considering the high molecular weight of the ligand, together with its high degree of conformational freedom, the NMU25 peptide was divided into two different parts: PEP1 (composed of residues from Phe1 to Tyr18) and PEP2 (composed of residues from Phe19 to Asn25). This was done based on literature data[13], which assesses that the residues forming PEP2 are the ones involved in the interaction with the transmembrane (TM) region of the protein, while the remaining part (PEP1) remains in the extracellular part of the receptor.

The conformations of the PEP1 sequence were created with the conformational search tool implemented in MOE, using LowModeMD as the method[14]. For the calculation, the iteration limit was set to 1000 and the other parameters were kept to the default values. For PEP1, 6 different conformations were obtained, which were visually inspected searching for the one which could be more efficiently linked with the PEP2 sequence conformations in the binding site.

The receptors used for the calculations were MMS_NMUR2_1 and MMS_NMUR2_2. For each of these structures, all passages listed down below were carried out.

The molecular docking calculations were executed with Glide[15]. The receptor grid suitable for peptide docking was created in a way that the ligand PEP2 could be located entirely within the TM region. The residues considered for the binding site definition were: Gly98, Glu127, Arg288, Val310, and Gly313. The grid was built allowing the placement of ligands up to 30 Å in length. The docking calculation was performed using the "Glide-SP-peptide" protocol, obtaining 100 poses for the PEP2 ligand. Those were filtered to eliminate the ones presenting clashes with the protein and/or unfavorable electrostatic contacts. For each of the remaining conformations, the difference between the energy of that specific pose and the energy of the local minimum was computed, allowing to pick the conformations with the lowest strain. Among the final solutions, the selection was executed based on visual inspection.

The best poses for PEP2 have then been manually linked with the best conformation of PEP1, and the sequence from Phe1 to Phe19 of the overall peptide was minimized with the AMBER10:EHT force field implemented in MOE.

The complete NMUR2 ligand conformations obtained for both MMS_NMUR2_1 and MMS_ NMUR2_2 were submitted to the post-docking phase.

#### Post-docking

Each docking pose was refined through classical Molecular Dynamics simulations. At first, the model was aligned to the OPM file of the template (retrieved from the Orientation of Protein in Membrane database). Subsequently, system preparation for the molecular dynamics simulations was carried out using VMD. The complex was embedded in a lipid bilayer consisting of phosphatidylcholine (POPC) units and explicitly solvated in a TIP3P water box with the borders placed at 15 Å from any protein atom. The system charge was neutralized to a concentration of 0.154 M using $Na^+/Cl^-$.

Molecular Dynamics simulations were performed with the CHARMM36 force field using the ACEMD3 engine.

Before MD simulation, 1500 steps of energy minimization with the conjugate-gradient algorithm were performed to remove clashes and bad contacts among the system atoms. Subsequently, a three-stage system equilibration was performed. The first equilibration step consisted of 5 ns of NPT simulation with harmonic positional restraints of 5 kcal $mol^{-1}Å^{-2}$ on each atom of the protein, the ligand, and the lipid bilayer. The second step consisted of 10 ns of NPT simulation with harmonic positional restraints of 5 kcal $mol^{-1}Å^{-2}$ only on each protein and ligand atom and on the phosphorus atom of the POPC units, the third step consisted of 5 ns of NPT simulation with harmonic positional restraints of 5 kcal $mol^{-1}Å^{-2}$ only on the alpha carbons of the protein and on each ligand atom. During the equilibration stages, the temperature was maintained at 310 K using the Langevin thermostat, with low dumping of 1 $ps^{-1}$, while the pressure was set at 1 atm using a Monte Carlo barostat. The particle-mesh Ewald (PME) method was used to calculate the electrostatic interactions with a 1 Å grid. A 9.0 Å cutoff was applied for long-term interactions. The M-SHAKE algorithm was applied to constrain the bond lengths involving hydrogen atoms.

After the equilibration stages, three independent productive 25 ns MD simulations were carried out in the NVT ensemble (T = 310 K), using the same setup as for the system equilibration stage except for the absence of the barostat and the restraints.

The trajectories were then clustered based on the pairwise RMSD matrix of the ligand backbone using the TTCLUST Python program, using the default parameters. The final pose selection was based on visual inspection of the most representative frames of each cluster.

##### Results summary

| Submitted complex | Model | Docking method |
|---|---|---|
| NMUR2_GPCRdock-2223-1 | MMS_NMUR2_1 | Glide+LowModeMD |
| NMUR2_GPCRdock-2223-2 | MMS_NMUR2_1 | Glide+LowModeMD |
| NMUR2_GPCRdock-2223-3 | MMS_NMUR2_1 | Glide+LowModeMD |
| NMUR2_GPCRdock-2223-4 | MMS_NMUR2_1 | Glide+LowModeMD |
| NMUR2_GPCRdock-2223-5 | MMS_NMUR2_2 | Glide+LowModeMD |

### GPCRdock-2223 - NPY1R

#### Homology Modeling

Two different homology models were used to predict ligand binding mode. The first model (MMS_NPY1R_1) was retrieved from the AlphaFold[1] database, while the second one (MMS_NPY1R_2) was generated as stated in the following lines:

- The sequence of NPY1R receptor (UniProt ID: P25929) was retrieved in the fasta format from the UniProt database.
- The fasta sequence was uploaded to the PHYRE2[2] web server, and a normal mode search was performed to fetch possible templates. The structure deposited in the Protein Data Bank with accession code 5ZBQ[3] (The Crystal Structure of human neuropeptide Y Y1 receptor with UR-MK299) was chosen as a template for the generation of the homology model.
- The sequence alignment between the template and the NPY1R receptor was imported into Molecular Operating Environment (MOE[4]) 2019.01, alongside the template structure (5ZBQ) retrieved from the PDB. The chimeric part of the construct (from Asn1001 to Tyr1160) was removed. The rest of the structure was prepared exploiting the Structure Preparation and Protonate3D tools (removal of crystallographic water molecules, ions, and other solvent molecules, selection of the highest occupancy for each residue, assignment of the correct protonation state at pH 7.4). The best model, according to MOE, was then manually refined, checking for consistency with the predicted secondary structure (according to both PHYRE2 and JPRED4[5] web server prediction). The missing intracellular loop 3 (ICL3), was then manually reconstructed as follows: residue 254 to 256 were built in α-helix conformation using the Protein Builder module of MOE, while residues 242 to 253 were modeled ab-initio using the Loop Modeler module of MOE.

MMS_NPY1R_2 was then refined through classical Molecular Dynamics simulations. At first, the model was aligned to the OPM file of the template (retrieved from the Orientation of Protein in Membrane database). Subsequently, system preparation for the molecular dynamics simulations was carried out using VMD[5]. The protein was embedded in a lipid bilayer consisting of phosphatidylcholine (POPC) units and explicitly solvated in a TIP3P[6] water box with the borders placed at 15 Å from any protein atom. The system charge was neutralized to a concentration of 0.154 M using $Na^+$/$Cl^-$.

Molecular Dynamics simulations were performed with the CHARMM36[7] force field using the ACEMD3[8] engine, which is based upon OpenMM7[9].

Before MD simulation, 1500 steps of energy minimization with the conjugate-gradient algorithm were performed to remove clashes and bad contacts among the system atoms. Subsequently, a three-stage system equilibration was performed. The first equilibration step consisted of 5 ns of NPT simulation with harmonic positional restraints of 5 kcal $mol^{-1}Å^{-2}$ on each atom of the protein and the lipid bilayer. The second step consisted of 10 ns of NPT simulation with harmonic positional restraints of 5 kcal $mol^{-1}Å^{-2}$ only on each protein atom and on the phosphorus atom of the POPC units, the third step consisted of 5 ns of NPT simulation with harmonic positional restraints of 5 kcal $mol^{-1}Å^{-2}$ only on the alpha carbons of the protein. During the equilibration stages, the temperature was maintained at 310 K using the Langevin thermostat, with low dumping of 1 $ps^{-1}$, while the pressure was set at 1 atm using a Monte Carlo barostat. The particle-mesh Ewald (PME[10]) method was used to calculate the electrostatic interactions with a 1 Å grid. A 9.0 Å cutoff was applied for long-term interactions. The M-SHAKE algorithm was applied to constrain the bond lengths involving hydrogen atoms.

After the equilibration stages, a productive 250 ns MD simulation was carried out in the NVT ensemble (T = 310 K), using the same setup as for the system equilibration stage except for the absence of the barostat and the restraints.

The trajectory was then clustered based on the pairwise RMSD matrix of the backbone using the TTCLUST[11] Python program, using the default parameters. The representative frame of the most populated cluster was then used for subsequent docking calculations.

#### Docking

The peptide ligand for the NPY1R receptor (NPY) was properly prepared for docking calculations using the MOE suite. The fasta sequence of the peptide was passed through the "Protein builder" tool, allowing to obtain the three-dimensional structure of the ligand. Then, the "Protonate 3D" program was implemented to assign the proper protonation state to each residue at a defined pH of 7.4. Finally, all the hydrogens in the system were minimized under the AMBER10:EHT[13] force field implemented in MOE.

Considering the high molecular weight of the ligand, together with its high degree of conformational freedom, the NPY peptide was split into two different parts: PEP1 (composed of residues from Tyr1 to Ile28) and PEP2 (composed of residues from Asn29 to Tyr36). This was done based on literature data[3], which assesses that the residues forming PEP2 are the ones involved in the interaction with the transmembrane (TM) region of the protein, while the remaining part (PEP1) remains in the extracellular part of the receptor.

The conformations of the PEP1 sequence were created with the conformational search tool implemented in MOE, using LowModeMD as the method[14]. For the calculation, the iteration limit was set to 1000 and the other parameters were kept to the default values. For PEP1, 4 different conformations were obtained, which were visually inspected searching for the one which could be more efficiently linked with the PEP2 sequence conformations in the binding site.

The receptors used for the calculations were MMS_NPY1R_1 and MMS_NPY1R_2. For each of these structures, all passages listed down below were carried out.

The molecular docking calculations were executed with Glide[15]. The receptor grid suitable for peptide docking was generated extrapolating the center of mass from the most relevant residues for NPY1R-NPY interaction, mainly retrieved from the work of Yang et al.[3]. The residues considered for the binding site definition were: Tyr100, Gln120, Cys121, Ile124, Trp276, Leu279, Thr280, Phe282, Asn283, Phe286, Asp287, and Ans299. The grid was built allowing the placement of ligands up to 30 Å in length. The docking calculation was performed using the "Glide-SP-peptide" protocol, obtaining 100 poses for the PEP2 ligand. Those were filtered to eliminate the ones presenting clashes with the protein and/or unfavorable electrostatic contacts. For each of the remaining conformations, the difference between the energy of that specific pose and the energy of the local minimum was computed, allowing to pick the conformations with the lowest strain. Among the final solutions, the selection was executed based on visual inspection, giving higher relevance to the poses presenting the contacts described in the literature for NPY1R-NPY interaction. The residues considered were the same listed above, which were considered for the receptor grid binding site definition.

The best poses for PEP2 have then been manually linked with the best conformation of PEP1, and the sequence from Tyr1 to Asn29 of the overall peptide was minimized with the AMBER10:EHT force field implemented in MOE.

The so-obtained complete NPY ligand conformations for both MMS_NPY1R_1 and MMS_NPY1R_2 were submitted to the post-docking phase.

#### Post-docking

Each docking pose was refined through classical Molecular Dynamics simulations. At first, the model was aligned to the OPM file of the template (retrieved from the Orientation of Protein in Membrane database). Subsequently, system preparation for the molecular dynamics simulations was carried out using VMD. The complex was embedded in a lipid bilayer consisting of phosphatidylcholine (POPC) units and explicitly solvated in a TIP3P water box with the borders placed at 15 Å from any protein atom. The system charge was neutralized to a concentration of 0.154 M using $Na^+/Cl^-$.

Molecular Dynamics simulations were performed with the CHARMM36 force field using the ACEMD3 engine.

Before MD simulation, 1500 steps of energy minimization with the conjugate-gradient algorithm were performed to remove clashes and bad contacts among the system atoms. Subsequently, a three-stage system equilibration was performed. The first equilibration step consisted of 5 ns of NPT simulation with harmonic positional restraints of 5 kcal $mol^{-1}Å^{-2}$ on each atom of the protein, the ligand, and the lipid bilayer. The second step consisted of 10 ns of NPT simulation with harmonic positional restraints of 5 kcal $mol^{-1}Å^{-2}$ only on each protein and ligand atom and on the phosphorus atom of the POPC units, the third step consisted of 5 ns of NPT simulation with harmonic positional restraints of 5 kcal $mol^{-1}Å^{-2}$ only on the alpha carbons of the protein and on each ligand atom. During the equilibration stages, the temperature was maintained at 310 K using the Langevin thermostat, with low dumping of 1 $ps^{-1}$, while the pressure was set at 1 atm using a Monte Carlo barostat. The particle-mesh Ewald (PME) method was used to calculate the electrostatic interactions with a 1 Å grid. A 9.0 Å cutoff was applied for long-term interactions. The M-SHAKE algorithm was applied to constrain the bond lengths involving hydrogen atoms.

After the equilibration stages, three independent productive 25 ns MD simulations were carried out in the NVT ensemble (T = 310 K), using the same setup as for the system equilibration stage except for the absence of the barostat and the restraints.

The trajectories were then clustered based on the pairwise RMSD matrix of the ligand backbone using the TTCLUST Python program, using the default parameters. The final pose selection was based on visual inspection of the most representative frames of each cluster.

#### Results summary

| Submitted complex | Model | Docking method |
|---|---|---|
| NPY1R_GPCRdock-2223-1 | MMS_NPY1R_1 | Glide+LowModeMD |
| NPY1R_GPCRdock-2223-2 | MMS_NPY1R_1 | Glide+LowModeMD |
| NPY1R_GPCRdock-2223-3 | MMS_NPY1R_1 | Glide+LowModeMD |
| NPY1R_GPCRdock-2223-4 | MMS_NPY1R_2 | Glide+LowModeMD |
| NPY1R_GPCRdock-2223-5 | MMS_NPY1R_2 | Glide+LowModeMD |

**GPCRdock-2229 - OPRK**

**Method**

Schrodinger Maestro package (2019-3) was used to perform all in silico analysis. Peptide molecule was sketched using build-module in Maestro modeling interface. The KOR crystal structure complex with the agonist (PDB: 6B73) was downloaded from RCSB PDB database. The downloaded proteins were optimized using Protein Prep Wizard in Maestro modeling interface, and we built a homology model with this PDB crystal structure as it missed some residues on loop. The grid was generated around the similar binding site as 6B73 showed using receptor grid generation wizard of Glide module. Once after grid generation, the docking was carried out using the ligand docking wizard of glide module. XP docking was the preferred mode of docking operation. The Prime minimize module was used to optimize and generate the final complex model.

### GPCRdock-2229

### Building model of GPR139/7c complexes

### for GPCR Dock 2021

#### A: TEMPLATE SELECTION AND TARGET-TEMPLATE ALIGNMENT

The sequence of the target receptors was obtained from the UniProt website (https://www.uniprot.org/). Identification of suitable templates was made using BLAST searching against Protein Data Bank proteins(pdb) database, The closest template that has the highest sequence identity and similarity and the least number of gaps was chosen for homology modeling of the target receptors. Thus, the crystal structure of the N/OFQ Opioid Receptor (PDB: 4EA3) was selected as a template to build homology models of GPR139.

#### B: PROTEIN MODEL BUILDING

The homology models of the GPR139 receptors were built using the Prime STA algorithm of the Schrodinger2019-2. The models were constructed using the default protocols.

#### C: PROTEIN MODEL REFINEMENT AND SELECTION

The Prime refinement module of Schrodinger2019-2 was used to refine the structural model of GPR139 receptors. And The protein model was further optimized by SAR analysis of the TAK-041 and 7C articles and patents.

#### D: LIGAND DOCKING

The docking experiment was conducted using the glide-v8.3 module Schrodinger2019-2.

#### E: PROTEIN/LIGAND COMPLEX REFINEMENT AND SELECTION

The bioactive conformation of the ligand was determined by the Jaguar-v10.4 module of Schrodinger2019-2, and then the correct conformation of the complex was determined by SAR analysis.

**GPCRdock-2229**

**Building models of APJ/cpd399 complex for GPCR Dock 2021**

**A: TEMPLATE SELECTION AND TARGET-TEMPLATE ALIGNMENT**
**Underlying tool(s)/platform(s).** N/A
**B: PROTEIN MODEL BUILDING**
**B1. Underlying tool(s)/platform(s).** N/A
**C: PROTEIN MODEL REFINEMENT AND SELECTION**
**C1. Underlying tool(s)/platform(s). N/A**
**D: LIGAND DOCKING**
**D1. Underlying tool(s)/platform(s).** Induced Fit Docking module of Schrodinger 2019-2

The crystal structure of the Apelin receptor (PDB ID: 5VBL) in complex with agonist peptide was used to perform ligand docking calculation. Because the target molecule structure is similar to the agonist peptide.

**E: PROTEIN/LIGAND COMPLEX REFINEMENT AND SELECTION**
**E1. Underlying tool(s)/platform(s).** Jaguar-v10.4 module of Schrodinger2019-2

All refinements were conducted in the Schrodinger program, including Conformation Search, Jaguar-Optimizations and Jaguar-Relaxed Coordinate Scan. Then the correct conformation of the complex was determined by SAR analysis.

**Methods from GPCRdock-2327** (Albert J. Kooistra, Gáspár Pándy-Szekeres, Kasper Harpsøe and David Gloriam), **Abbreviations:** AF = AlphaFold

| Model no. | GPCR | [1]Main template | PDB | [2]Other template | PDB | For regions | Modeling software | Ligand binding mode | Experimental information and human expertise | Selection criteria for top 5 | Target challenges |
|---|---|---|---|---|---|---|---|---|---|---|---|
| 1-5 | APJ | APJ | 5VBL | - | - | - | GPCRdb | Glide docking | SAR information for from patent EP2897939B1, i.e. similar binding pose across high-potency analogs, and binding site characterization were taken into account while evaluating the docking poses. The docking poses were visually inspected with the SAR and previously published APJ structures in mind to find poses with an optimal fit, i.e. similar location in the binding site of functional groups with the same characteristics. | • Similar location of similar functional groups compared to previous published structures<br>• Consistent docking pose across high-potency analogues<br>• High ranking according to docking score.<br>• Fit to binding site characterization. | Selecting the binding poses |
| 1 | KOR | KOR | 6B73 | MOR | 6DDE | Ligand, ICL1-2, IC TM ends, AlphaFold-Multimer for ECL2 | GPCRdb + MODELLER | MODELLER (template DAMGO in 6DDE) + 2N2F Dynorphin NMR 10th conformation | An iterative RMSD calculation was carried out to identify the merge locations in the intracellular ends of the 7TM helices. With this we wanted to mimic the Gi1 binding of 6DDE on our main template 6B73.<br><br>Mutagenesis data from literature were taken into account when visually evaluating the peptide-receptor complexes. | Selection was based on covering a wider variation for the ligand-ECL2 positioning and prioritizing viable ligand-receptor contacts. | • Identifying merging sites for the receptor model<br>• ECL2 modeling<br>• modeling the Dynorphin tail. |
| 2 | KOR | KOR | 6B73 | MOR | 6DDE | Ligand, ICL1-2, IC TM ends, AlphaFold-Multimer for ECL2 | GPCRdb + MODELLER | MODELLER (template DAMGO in 6DDE) + 2N2F Dynorphin NMR 1st conformation | | | |
| 3 | KOR | KOR | 6B73 | MOR KOR | 6DDE 4DJH | Ligand, ICL1-2, IC TM ends part of ECL2, ECL2-3 minimized | GPCRdb + MODELLER | MODELLER (template DAMGO in 6DDE) + 2N2F Dynorphin NMR 1st conformation minimized | | | |
| 4 | KOR | KOR | 6B73 | MOR KOR | 6DDE 4DJH | Ligand, ICL1-2, IC TM ends part of ECL2 | GPCRdb + MODELLER | MODELLER (template DAMGO in 6DDE) + Dynorphin NMR 1st conformation | | | |
| 5 | KOR | KOR | 6B73 | MOR | 6DDE | Ligand, ICL1-2, IC TM ends, AlphaFold-Multimer for ECL2 | GPCRdb + MODELLER | AlphaFold-Multimer using 48x recycling, no templates, Amber relaxation and multiple model generation (ColabFold) | | | |
| 1 | NPY1R | - | - | - | - | - | AlphaFold-Multimer + AlphaFold-multistate | Default settings AlphaFold-Multimer using no templates | Mutagenesis data from literature were taken into account when visually evaluating the peptide-receptor complexes.<br>DOIs:<br>• 10.1038/s41586-018-0046-x<br>• 10.1042/BJ20050708<br>• 10.1002/anie.201411688<br><br>When merging the main template with the other template used for the intracellular helix ends or when merging the AlphaFold-Multimer and AlphaFold-Multistate models, after a 7TM superposition, we manually identified the spatially closest residues between the two structures. These were the positions where we merged the two structures.<br><br>The position of several functional groups of the ligand in the inactive state NPY1R structure where good confirmation for our AlphaFold ligand models. | The five models were created by five different approaches.<br><br>As Neuropeptide Y was modelled quite consistently in the different models, the focus was on small variations in the placement of the peptide while evaluating AlphaFold, GPCRdb-based, and manual models for the receptor.<br><br>Model 5: Lowest energy structure from the highest occupied cluster after MD – remaining procedures only resulted in a single structure | Identifying merge positions for the receptor model, ligand-ECL2 modeling, Identifying the right tilt angle between the ligand's helical part and the receptor and modeling the interacting ECLs. |
| 2 | NPY1R | - | - | - | - | - | AlphaFold-Multimer + AlphaFold-multistate | AlphaFold-Multimer using 48x recycling, no templates, no relaxation and multiple model generation (ColabFold) – model rank 3 | | | |
| 3 | NPY1R - Gi | OX2R -Gs | 7L1U | NPY1R CCKAR - | 5ZBQ 7EZH - | ECL1-2 and TM ends TM7 end and H8 start AlphaFold-Multimer for ECL3 | GPCRdb + MODELLER + AlphaFold-Multimer | From model 1 + manual refinement | | | |
| 4 | NPY1R - Gi | CCKAR1 -Gi | 7EZH | NPY1R - | 5ZBQ - | ECL1-2 and TM ends AlphaFold-Multimer for ECL3 | GPCRdb + MODELLER + AlphaFold-Multimer | From model 1 + manual refinement | | | |
| 5 | NPY1R - Gi | NPY1R | 5ZBQ | CCKAR | - | Last helical turn of TM7 | Manual, selected TMHs moved based on 7EZH | From model 1 | | | |
| 1 | NMUR2 | - | - | - | - | - | AlphaFold-Multimer + AlphaFold-multistate + MOE (merge + minimization sidechains) | AlphaFold-Multimer using 48x recycling, no templates, Amber relaxation and multiple model generation (ColabFold) | Peptide SAR and mutagenesis data from literature were taken into account when visually evaluating the peptide-receptor complexes<br>DOIs:<br>• 10.1016/j.ejmech.2017.12.035<br>• 10.1021/acsmedchemlett.8b00105<br>• 10.1021/jm500599s<br>• 10.1016/s0196-9781(02)00097-9<br><br>When merging the main template with the other template used for the intracellular helix ends or when merging the AlphaFold-Multimer and AlphaFold-Multistate models, after iterative rounds of superpositioning with a focus on the middle of the heptahelical bundle, we manually identified the spatially closest residues (based on the backbone residues) between | As the N-terminal residues of Neuromedin U had a high predicted AlphaFold error, there was a high degree of uncertainty the placement of these residues. Therefore, the main criterion was to use different peptide conformations for each model while evaluating both AlphaFold and GPCRdb-based models for the receptor. | The secondary structure and overall fold of Neuromedin U - especially the N-terminal half. |
| 2 | NMUR2 | - | - | - | - | - | AlphaFold-Multimer + AlphaFold-multistate + MOE (merge + minimization) | Default settings AlphaFold-Multimer using no templates | | | |

| | | | | | | | | | |
|---|---|---|---|---|---|---|---|---|---|
| 3 | NMUR2 | GHSR | 7NA7 | OX2R<br>NTR1<br>AA1R | 7L1V<br>7L0R<br>5UEN | ECL2<br>TM5 end<br>TM6 start | GPCRdb +<br>MODELLER | MOE hom. modeling + manual refinement (template reversed Ghrelin peptide in 7F9Y) + sec. struct. based on model 1 (AlphaFold) | the two structures for each individual helix. These were the positions where we merged the two structures. |
| 4 | NMUR2 | GHSR | 7NA7 | OX2R<br>NTR1<br>AA1R | 7L1V<br>7L0R<br>5UEN | ECL2<br>TM5 end<br>TM6 start | GPCRdb +<br>MODELLER | MOE hom. modeling + manual refinement (template reversed Ghrelin peptide in 7F9Y) + sec. struct. based on model 1 (AlphaFold)<br>*Alternate alignment/placement* | |
| 5 | NMUR2 | - | - | - | - | - | AlphaFold-Multimer + AlphaFold-multistate + MOE (merge + minimization sidechains) | Pep-Fold3, MOE (manual refinement peptide fold based on strongest coupling residue pairs EVcouplings + placement peptide based on 7F9Y) | |

**[1]Main template modeling**
The main template was used to build models using the GPCRdb pipeline described in the publication (1) and source code deposited in GitHub (https://github.com/protwis/protwis/). This pipeline incorporates alternative templates to adjust helix lengths and corrects for helical bulges and constrictions. The pipeline also has a built-in rotamer library which models sidechains based on their specific position using GPCRdb generic residue numbers (2). The GPCRdb generic residue numbers are structure-based (based on structure superposition) instead of sequence-based as the Ballesteros-Weinstein numbers (based on sequence alignment). However, the two numbering schemes use the same number unless a transmembrane helix has bulge or constriction, which causes a structural gap which is also represented as a gap in the sequence alignment. Furthermore, the GPCRdb scheme covers ECL1-2, ICL1-2 and helix 8 which are not included in the Ballesteros-Weinstein scheme.

**[2]Secondary template modeling**
The secondary templates listed in the table were used as swap-in templates to the GPCRdb model using MODELLER (https://salilab.org/modeller/).

**AlphaFold modeling**
For the AlphaFold models several different workflow implementations have been used: the notebook accompanying the most recent AlphaFold (3)/AlphaFold-Multimer (4) release (version 2.1.1) was used as obtained from http://github.com/deepmind/alphafold, in addition the AlphaFold-Multimer notebook from ColabFold(5) was used for creating models using higher fold recycling as obtained from http://github.com/sokrypton/ColabFold. Finally, the active-state models of the receptors were obtained based on AlphaFold from https://github.com/huhlim/alphafold-multistate, which is further described in a bioRxiv preprint (6).
The AlphaFold models were evaluated using the AlphaFold predicted error as well as with mutagenesis data from literature for the receptor as well as peptide (where available). Multiple models were generated for each complex and different settings for the number of recycling iterations, the use of templates, and relaxation were explored.
Merging of AlphaFold-Multistate and AlphaFold-Multimer models was performed manually using MOE (release 2020.09) from the Chemical Computing Group and Maestro (release 2021-4) from Schrödinger by performing iterative rounds of superpositioning with a focus on the middle of the heptahelical bundle. After obtaining an optimal alignment of the backbone in the middle of the bundle, the models were fused by linking the backbone where the atomic distance was the lowest for each individual helix.

**Other modeling details**
For model 5 of the NMUR2-NMU25 complex, PEP-Fold3(6) was used for obtaining an initial alternative peptide fold, which was manually refined based on the strongest couplers as obtained from an EVcouplings (7) analysis of NMU25.

For model 5 of the NPY1R-NPY complex, an active state model was constructed from the inactive state NPY1R structure, 5ZBQ (8), by individual TM-helix super position and modification based on the active state CCKAR1 structure, 7EZH (http://dx.doi.org/10.2210/pdb7ezh/pdb). Receptor assembly of the individual TM-helices and loops was performed in MODELLER (9) (version 9.24) by using all the fragments as template for one model. The NPY peptide, ECL2, ECL3 and the N-terminus was swapped-in from the NPY1R model obtained with AlphaFold after a short MD simulation, clustering analysis in Desmond and minimization in Prime (release 2021-4, Schrödinger, LLC, NY, 2021) to sample and optimize ECL2 (100 ns at 400 K with restraints on all atoms but ECL2 and contacting residues, i.e. within 5A). The swap-in was performed using MODELLER (9) (version 9.24) by defining specific template selection from the structures to be combined. Sidechains of the peptide and receptor in the buried binding site were then optimized with the Predict Side Chains protocol in Prime (release 2021-4, Schrödinger, LLC, NY, 2021).

**Small molecule binding modes in APJ**

The small molecule ligand was docked into the homology models with Glide (release 2021-4, Schrödinger, LLC, NY, 2021) together with many analogous retrieved from a patent (WO 2014/044738). Binding poses were selected based on match of similar pharmacophore to the peptide bound to APJ in the 5VBL structure (10), binding site characteristics obtained with SiteMap (release 2021-4, Schrödinger, LLC, NY, 2021) and SAR of the ligand series. Sidechains surrounding the ligand were optimized with the Predict Side Chains protocol in Prime, (release 2021-4, Schrödinger, LLC, NY, 2021).

**GPCRdock-2646**

GPCR Dock 2021 Method
Yuguang Mu [a] and Liangzhen Zheng [bc]
[a] School of Biological Sciences, 60 Nanyang Drive, Singapore 637551
[b] Shenzhen Institute of Advanced Technology, Chinese Academy of Sciences, Shenzhen, Guangdong 518055, China
[c] Shanghai Zelixir Biotech Company Ltd., Shanghai 200030, China

For small-molecule targets (APJ and GPR139), the GPCR protein models were built through AlphaFold2 colab version (https://colab.research.google.com/github/sokrypton/ColabFold/blob/main/AlphaFold2.ipynb) without Amber relaxation. Vina 2 was then used to dock the molecules, followed by the use of our own method, SFCT method (Briefings in Bioinformatics, 2022, 1–15 https://doi.org/10.1093/bib/bbac051) to rescore. Finally the top molecules were chosen for short MD simulations.

The GPCR and peptide complex models were also constructed through the AlphaFold2 server in the "multimer" mode. The predicted GPCR-peptide complexes were taken as the initial stricture for a short run ( 10 ns ) of molecular dynamics simulations with explicit water molecules using AMBER99SB force field (Proteins, 65 (2006), pp. 712-725. https://doi.org/10.1002/prot.21123). The binding affinity between the GPCR and peptides were estimated by using MMGBSA method( AMBER package with IGB=8) for the final 2 ns MD trajectory. The ranking of the final models was based on the averaged GBSA binding affinity with entropy contribution ignored.

When the ligand is small molecule, the binding pockets of the related GPCR was predicted using the pointsite (https://www.biorxiv.org/content/10.1101/831131v1.full) and then the small molecule ligand was docked using quick vina tool (Bioinformatics (2015) 31 (13): 2214-2216. DOI:10.1093/bioinformatics/btv082) into the predicted binding pocket. The docked molecules were further rescored using the recently developed SFCT method (Briefings in Bioinformatics, 2022, 1–15 https://doi.org/10.1093/bib/bbac051). Finally the top molecules were chosen based on SFCT rescoring.

**Target challenges:** Some poses in GPCR-peptide complexes were difficult to rank, with MMGBSA predicted binding affinity very close, even manually inspection cannot provide a helpful clue. In the future, machine-leaning based protein-peptide binding affinity prediction methods are needed.

**Group code: GPCRdock-2717 ( DiMaio Lab, UW )**

**Methods:**

**Small molecule ligand preparation**: Initial ligand structures were generated from smiles string using UCSF Chimera[1]. AmberTools[2] was then used to generate am1bcc partial charges and ideal bond geometry for the small molecules. A utility script in Rosetta "mol2genparams.py" was used to generate the small molecule parameter file for GALigandDock.

**Target protein modeling**: AlphaFold [3] and RoseTTAFold [4] were used for generating the target apo structures using the sequence provided.

For Targets T01 and T02 (APJ and GPR139), we used AlphaFold and RoseTTAFold to predict the target apo structures using the target sequence with resolved density and we selected 5 models from AlphaFold predictions and 15 models from RoseTTAFold predictions. We also used RosettaCM[5] to build 25 partial threaded models based on the top 25 homologs predicted from HHpred[6]. A total of 45 putative receptor structures were used for ligand docking against each target.

For peptide Targets T03, T04 and T05, AlphaFold was used to directly predict the complex structure by appending the peptide sequence to the target sequence and changing the residue number of the peptides so that there were 100-residue gap between the receptor sequence and the peptide sequence. Two sets of target sequences were used in AlphaFold predictions, one was the raw sequence (referred as full-sequence version), the other one was only using the sequence with resolved density (referred as trimmed-sequence version).

**Rosetta GALigandDock:**
Rosetta GALigandDock[7], which combines a physically realistic energy model with genetic algorithm optimization, was used to predict the structure of protein-ligand (small molecule and peptide) complexes. Binding affinity of the docked structure is also estimated in GALigandDock using the following equation: dG = dH + TdS. The enthalpy change (dH) upon binding is estimated by dH = E(complex) - E(receptor) - E(ligand) where E(X) is the Rosetta energy of X. The estimation of entropy changes is computed by estimating the torsional entropy of rotatable bonds of the ligand using short Monte Carlo simulation. The full settings for running GALigandDock is shown below in RosettaScripts[8] XML format (see supporting information, GALigandDock XML).

For T01 and T02, the input structures of GALigandDock were manually prepared by placing the small molecule to the binding pocket of the 45 putative receptor structures. Twenty independent docking runs were performed for each of the input structures.

For the protein-peptide docking targets i.e. T03, T04 and T05, we first examined the predicted structures from AlphaFold by the predicted aligned error (PAE) and the structure with the smallest PAE from each target was used as the input to the GALigandDock. We also included the five peptide structures predicted from AlphaFold as templates in GALigandDock to bias the conformational sampling of the peptide structures during docking. One hundred independent docking runs were performed for each target.

**Criteria for selecting 5 best models:**

For small molecule targets, i.e. T01 and T02, we first sorted all docked poses by predicted binding affinities. We then removed the structures that have buried polar atoms and we only kept the structures with better predicted binding affinities if two docked poses were within 2 Å to each other. For T01, we selected top two structures from homology modeled receptors, top two structures from RoseTTAFold predicted receptors and top one structure from AlphaFold predicted receptors. For T02, we selected top two structures from homology modeled receptors, top one structure from RoseTTAFold predicted receptors and top two structures from AlphaFold predicted receptors.

Among the submitted models for T01, we failed to predict the correct docked structure (see "target challenges" below). However, among the submitted models for T02, one of them has a ligand rmsd of 1.79A compared to the crystal structure and ranked top1 in this challenge.

For peptide targets, i.e. T03, T04 and T05, we sorted all the docked structures from GALigandDock by the total energy and the predicted enthalpy of binding (dH) due to the inaccuracies of the entropy estimation on peptides. After sorting all the docked structures, we only kept the top twenty structures presented in both sorted lists. For T03, we picked two models with the lowest total energy and three models with lowest predicted dH. For T04, we picked two models with lowest total energy, two models with the lowest predicted dH and one with the lowest PAE from AlphaFold trimmed-sequence predictions. For T05, we picked two models with the lowest total energy, one model with the lowest predicted dH and one model with the lowest PAE from AlphaFold full and trimmed-sequence predictions. GALigandDock failed to predict near-native complex structures for all three cases, which is mainly due to the fact that GALigandDock is unable to fully sample the conformations of the peptides with too many degrees of freedom. One of the submitted models for T05 ranked 3rd in this challenge, which is predicted directly using AlphaFold on the full-sequence.

**Target challenges:**

The most challenging part of predicting the complex structure for small molecule docking is the receptor structure prediction in the bound complex, especially the binding site structure prediction. Accurate prediction of ligand docking is highly dependent on the accuracy of the structure forming the binding site of the receptor. The best predicted receptor structure of T01, for which we failed to provide an accurate prediction, has two binding-site surrounding helices moved considerably compared to the native structure, while the predicted binding site of T02 has a much better agreement with the crystal structure (see Figure 1). We roughly estimated the rmsd of the predicted binding site to the crystal structure using the residues within 10 Å of the ligand. The rmsd of the predicted binding site of T01 is around 7.8 Å while the rmsd of the predicted binding site of T02 is only around 3.0 Å.

For peptide ligands, the large number of degrees of freedom is very challenging for Rosetta GALigandDock protocol to accurately model the complex structure.

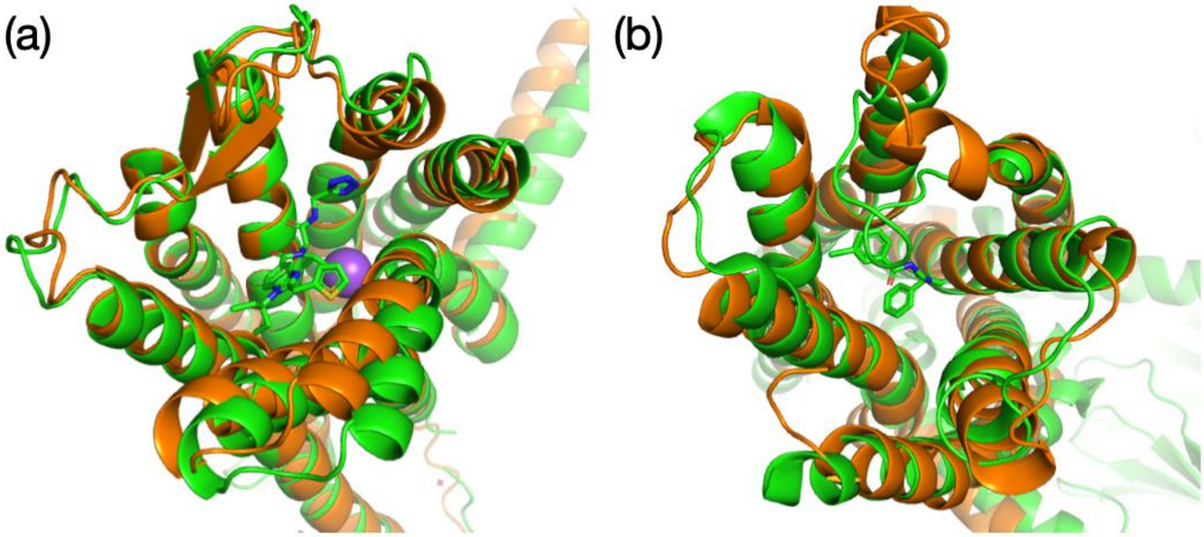

**Figure 1.** Superimpose of best predicted protein receptor structure (organe) with crystal structure (green): (a) T01 (b) T02

**Supporting Information:**

**GALigandDock XML:**

```
<ROSETTASCRIPTS>
  <SCOREFXNS>
    <ScoreFunction name="genpot_soft" weights="beta_cart">
      <Reweight scoretype="fa_rep" weight="0.2"/>
    </ScoreFunction>
    <ScoreFunction name="genpot" weights="beta_cart"/>
  </SCOREFXNS>

  <TASKOPERATIONS>
  </TASKOPERATIONS>

  <FILTERS>
  </FILTERS>

  <MOVERS>
    <GALigandDock name="dock" scorefxn="genpot_soft" scorefxn_relax="genpot"
                    runmode="dockflex" premin_ligand="1" padding="2.5" sidechains="aniso"
          final_exact_minimize="bbsc3" multiple_ligands="%%ligid%%" estimate_dG="1" >
    </GALigandDock>

  </MOVERS>

  <PROTOCOLS>
    <Add mover="dock"/>
  </PROTOCOLS>
```

```
  <OUTPUT scorefxn="genpot"/>
</ROSETTASCRIPTS>
```

[1] Pettersen, E. F. *et al.* UCSF Chimera. A visualization system for exploratory research and analysis. *J Comput Chem* **25**, 1605–1612 (2004).
[2] D.A. Case, H.M. Aktulga, K. Belfon, I.Y. Ben-Shalom, S.R. Brozell, D.S. Cerutti, T.E. Cheatham, III, G.A. Cisneros, V.W.D. Cruzeiro, T.A. Darden, R.E. Duke, G. Giambasu, M.K. Gilson, H. Gohlke, A.W. Goetz, R. Harris, S. Izadi, S.A. Izmailov, C. Jin, K. Kasavajhala, M.C. Kaymak, E. King, A. Kovalenko, T. Kurtzman, T.S. Lee, S. LeGrand, P. Li, C. Lin, J. Liu, T. Luchko, R. Luo, M. Machado, V. Man, M. Manathunga, K.M. Merz, Y. Miao, O. Mikhailovskii, G. Monard, H. Nguyen, K.A. O'Hearn, A. Onufriev, F. Pan, S. Pantano, R. Qi, A. Rahnamoun, D.R. Roe, A. Roitberg, C. Sagui, S. Schott-Verdugo, J. Shen, C.L. Simmerling, N.R. Skrynnikov, J. Smith, J. Swails, R.C. Walker, J. Wang, H. Wei, R.M. Wolf, X. Wu, Y. Xue, D.M. York, S. Zhao, and P.A. Kollman (2021), Amber 2021, University of California, San Francisco.
[3] Jumper, J. *et al.* Highly accurate protein structure prediction with AlphaFold. *Nature* **596**, 583–589 (2021).
[4] Baek, M. *et al.* Accurate prediction of protein structures and interactions using a three-track neural network. *Science* eabj8754 (2021) doi:10.1126/science.abj8754.
[5] Song, Y. *et al.* High-Resolution Comparative Modeling with RosettaCM. *Structure* **21**, 1735–1742 (2013).
[6] Söding, J., Biegert, A. & Lupas, A. N. The HHpred interactive server for protein homology detection and structure prediction. *Nucleic Acids Res* **33**, W244–W248 (2005).
[7] Park, H., Zhou, G., Baek, M., Baker, D. & DiMaio, F. Force Field Optimization Guided by Small Molecule Crystal Lattice Data Enables Consistent Sub-Angstrom Protein–Ligand Docking. *J Chem Theory Comput* **17**, 2000–2010 (2021).
[8] Fleishman, S. J. *et al.* RosettaScripts: A Scripting Language Interface to the Rosetta Macromolecular Modeling Suite. *Plos One* **6**, e20161 (2011).

**GPCRdock-2736**

Nicolas Panel‡, Israel Cabeza de Vaca‡, Flavio Ballante, Nour Aldin Kahlous, Silvana Vasile, Jens Carlsson*

Department of Cell and Molecular Biology, Science for Life Laboratory, Uppsala University, SE-75124 Uppsala, Sweden

‡ These authors contributed equally.

**Apelin receptor**

The goal was to predict the structure of the apelin receptor (APJR) in complex with a small-molecule ligand, which was described as an agonist in a patent ($EC_{50}$ = 0.099 μM).[1] The crystal structure of APJR in complex with the peptidomimetic agonist AMG3054 (PDB ID: 5VBL)[2] was selected as target for docking simulations. The receptor structure was prepared as follows: (1) solvent and buffer molecules were removed; (2) N- and C-termini were capped with acetyl and N-methyl amide groups, respectively; (3) hydrogens were added using REDUCE;[3] (4) AMG3054 was removed. ChemAxon calculator (version 21.15.0)[4] was used predict the ligand's dominant form at physiological pH. Surflex's similarity module (Version 5.125)[5] was then used to generate and superimpose 979 ligand conformers to a truncated structure of AMG3054 bound to APJR (the crystal structure conformation of the ligand): (CCCC[C@H](NC(=O)[C@@H]1C[C@@H]2CCCC[C@@H]2N1C(C)=O)C(=O)N1CCC[C@H]1C(=O)N[C@@H](Cc1ccc(Cl)cc1)C(O)=O). A total of 250 superimposed ligand conformations were generated and visually inspected. A ligand binding mode that mimicked the interactions of AMG3054 was used as input structure to perform docking simulations with AutoDock (release 4.2.6).[6] Docking input files were prepared using AutoDockTools scripts from the Utilities24 module. Molecular docking was performed using a grid centered on AMG3054, which was composed of 67x61x61 points in the *xyz* dimensions with a spacing of 0.375 Å. The Lamarckian genetic algorithm was used with a population of 300 individuals, a maximum of 5 million energy evaluations, 50000 generations and 100 runs. Five docking poses that reproduced the binding mode and key interactions of AMG3054 were then selected. The five models were subsequently relaxed by short molecular dynamics (MD) simulations. The systems were prepared with CHARMM-GUI[7] using the CHARMM36m[8] force field and ligand parameters were obtained with the CGenFF program.[9] Each complex was embedded in an equilibrated membrane bilayer composed of 142 1-palmitoyl-2-oleoyl-sn-glycero-3-phosphocholine (POPC) molecules surrounded by TIP3P water molecules with a 0.15 M concentration of neutralizing sodium and chloride ions. MD simulations were performed with GROMACS2021.3.[10] The systems were equilibrated for 1.25 ns by gradually increasing the temperature to 300 K. In this step, the receptor and ligand coordinates were maintained close to their initial coordinates by using harmonic restraints with force constants of 500 $kJ/mol/nm^2$ for the receptor backbone and ligand heavy atoms and 50 $kJ/mol/nm^2$ for the side chain heavy atoms. The final snapshot of each simulation was then minimized. Four of these minimized complexes were submitted to the GPCR Dock assessment without further refinement. For the binding mode with the best docking score, we carried out an additional 100 ns of MD simulations with harmonic restraints on the receptor backbone atoms (500 $kj/mol/nm^2$) whereas the other atoms were unrestrained. The final snapshot was minimized and submitted as the fifth model. The most challenging part of the modeling was to understand if the small-molecule ligand and AMG3054 would bind to the same receptor conformation and form similar interactions. Based on MD simulations and the similarities between the ligand and AMG3054, we decided to focus on models that had small deviations from the crystal structure with binding modes that overlapped with AMG3054.

**κ-opioid receptor**

The goal was to predict the structure of the κ-opioid receptor (KOR) in complex with a peptide agonist (dynorphin). Homology models of KOR bound to dynorphin were generated with MODELLER.[11] The structure of the μ-opioid receptor (MOP) bound the peptidomimetic DAMGO and G protein (PDB ID: 6DDE) was chosen as template.[12] DAMGO was mutated to match the first four residues of dynorphin and included in the template. Additional helical constraints were applied to the peptide between L5 and

I8 based on NMR experiments.[13] A total of 500 homology models were generated and clustered based on the Cα coordinates of the first eight residues of dynorphin with the TTclust program,[14] which led to a total of 11 clusters. MD simulations were used to assess the stability of the complexes obtained by homology modeling. One representative structure of each cluster was prepared for MD simulations using CHARMM-GUI[7] and the CHARMM36m force field.[8] The complex was embedded in an equilibrated membrane bilayer containing 145 POPC molecules surrounded by TIP3P water molecules with a 0.15 M concentration of neutralizing sodium and chloride ions. MD simulations were performed with GROMACS2021.3.[10] After an initial equilibration, five independent replicates of 20 ns were produced per model. In order to avoid large conformational changes of the receptor during the simulations, the backbone atoms of the receptor were restrained to their initial coordinates by harmonic restrains with a force constant of 50 kJ/mol/$nm^2$. Complex stability was evaluated by calculating the RMSD of the peptide Cα and monitoring receptor-peptide interactions. Five complexes were selected based on stability during the MD simulations and the diversity of receptor-ligand interactions. The homology models that were used to obtain these five snapshots were then minimized with harmonic restraints on the backbone. The five minimized complexes were submitted to the GPCR Dock assessment. The most challenging part of the modeling was to predict the structure of the peptide ligand that is not binding to the transmembrane (TM) cavity and interacts with the extracellular interface of the receptor. The addition of helical restraints derived from NMR experiments reduced the number of possible peptide binding modes in this region. Both AlphaFold2 (AF2)[15,16] and homology modeling were initially explored as tools to predict the receptor-ligand complex. Homology modeling was selected as method because the template had very high sequence identity (71%).

**Neuromedin U receptor 2 and neuropeptide $Y_1$ receptor**

The goal was to predict the structure of the neuromedin U receptor 2 (NMUR2) and neuropeptide $Y_1$ receptor (NPY1R) in complex with peptide agonists. Initial complexes were generated using the first AF2 multimer version released in GitHub.[15, 16] Receptor sequences used as inputs in AF2 were reduced by excluding unresolved residues in the N- and C-terminus. The sequences corresponding to the receptor and peptide ligand were used as input in the AF2 calculations. A total of 1000 models were generated with AF2 for each target. These models were ranked according to the number of hydrogen bonds between the receptor and peptide ligand. A frequency analysis of the hydrogen bonds was performed to identify key residues for peptide bindin. These results were compared with published mutagenesis experiments to identify models that agreed with the experimental data. For NMUR2, the interaction analysis highlighted four important interactions with the receptor (E102, E127, R288 and H309), which agreed with the results of a mutagenesis study on an insect ortholog of human NMUR2.[17] For NPY1R, we focused on the hydrogen bonds between the C-terminal fragment of the NPY peptide ($^{30}$LITRQRY$^{36}$-NH2) and the receptor. This part of the peptide is known to be crucial for high affinity binding. Based on mutagenesis data from several studies,[18-22] the following interactions were identified as important: L30-I293, R33-N299, R35-D287, Y36-$NH_2$ - Q120 and N283.

All generated AF2 models represented inactive receptor states based on the conformations of TM helices 5, 6 and 7. To obtain active conformations, we selected 100 AF2 models using interaction analysis and mutagenesis data and used these to generate homology models,. MODELLER[1] was used to generate active receptor models by combining two different templates. TM helices 5, 6 and 7 were modeled using the best template found by sequence similarity and the AF2 generated structures were used for the remaining parts of the sequence and peptide ligand. Neurotensin receptor 1 (PDB ID: 6OS9) and orexin receptor 2 (PDB ID: 7L1U) were used as templates for NMUR2 and NPY1R, respectively. The 10,000 models generated by MODELLER were filtered by molpdf (minimal value = 2200) and then minimized with Chimera using the AMBER force field. An additional filter extracted models in which the identified key residues formed intermolecular hydrogen bonds, which reduced the number of NMUR2 and NPY1R models to 215 and 33, respectively. These final models were clustered with the TTclust program[14] based on the receptor C〈 RMSD, which led to 15 clusters for each target. The 15 cluster centers were prepared for MD simulations using the protocol described for KOR and the peptide ligands were amidated by adding the C-terminal $NH_2$ group. Two simulation replicas were performed for each model with restraints in the production phase. Finally, 11 snapshots per model were clustered using the last six residues of the peptide ligand with GROMACS[2] and a representative structure was visually inspected.

Five models were selected and these were minimized with restraints on the C⟨ atoms using GROMACS and the CHARMM36m force field.[8] The resulting structures were submitted to the GPCR Dock assessment. The most challenging part of the modeling was to predict the structure of the peptide ligand that is not binding to the TM cavity and interacts with the extracellular interface of the receptor. Our strategy to solve this problem was to generate a large number of models and use interaction analysis combined with experimental mutagenesis data to identify good models. Another challenge was to both generate an accurate binding mode of the peptide and an active conformation of the receptor, which was accomplished by combining AF2 and MODELLER predictions.

### GPCRdock-2928 Methods

#### Protein Structures

Protein structures were built by homology modeling based on Swiss-Model1 and comparative modeling by Robetta server[2]. The choice of templates referred to the sequence similarity and the differences between structures, as we intended to use different input conformations for docking. We also adopted template-free modeling approach such as RoseTTAFold[3]. Finally, two structures among these candidates were selected as the 3D structures for each receptor.

#### Ligand Structures

Meanwhile, the small molecule ligands were constructed to 3D structure and the conformations of peptide ligands were generated by Monte Carlo conformation searching.

#### Docking

With the proteins and ligands, we applied flexible docking provided by Smina and glide programs to each system, which adjusts the position of sidechains of the pocket residues during docking. As for peptide ligands, HPEPDOCK Server was also used to produce possible poses[4].

All candidate complex structures were coarsely screened by a deep learning-based structure scoring algorithm, IGT[5], which is recently designed by our research. In addition, the binding affinity calculated by docking between the receptor and the ligand is also used as a reference. Hydrogens and implicit membranes were then added to the top-scored structures. The structures were energy minimized with constraints on heavy atoms. After minimization, molecular mechanics generalized Born surface area (MM/GBSA)[6] algorithm was used to evaluate the binding mode of each receptor-ligand complexes. Referring to the binding free energies and the diversity of ligand poses, 5 poses of each target were picked for the following molecular dynamics (MD) simulation.

#### MD simulation

We firstly added missed residues in homolog modeling but with density in the targets and minimized them. Then, the corresponding G proteins and antibodies were added to systems according to the alignment with homologs-G protein-antibody structure. The systems were submitted to the CHARMM-GUI server for construction with POPC membrane, 0.15 M/L KCL and OPC wars[7,8]. FF19SB, LIPID17, and GAFF2 force fields were applied to describe the interactions of protein, lipids, and small molecule ligands[7,9]. Since some systems have problems when using the FF19SB force field, CHARMM36m force field was rather applied. Amber20 was employed to produce MD simulation trajectories[10]. The minimization, heating, and equilibrium steps referred to the files provided by CHARMM-GUI. After 50-100 ns production runs, the final 5 structures for each system were picked by visual inspection considering the diversity of ligand poses and the reasonability of receptor structures.

#### GPCRdock-3183 – APJ

***Title: In Silico methods for structure prediction of Apelin receptor (APJ) drug complex***

Soo-Kyung Kim and William A. Goddard III
Materials and Process Simulation Center (MC139-74), California Institute of Technology,
1200 E. California Blvd., Pasadena, CA 91125


#### Methods

**Ensemble structure generation for Apelin receptor (APJ):** We predicted the ensemble of low energy APJ G protein-coupled receptor (GPCR) structures using the GEnSeMBLE complete sampling method. (Abrol, Bray et al. 2011, Abrol, Griffith et al. 2012, Abrol, Kim et al. 2013, Bray, Abrol et al. 2014) GEnSeMBLE identifies the transmembrane (TM) domains based on multiple sequence alignments, then selects the shape of each helix based on the force field or on homology to known structures, and then considers a complete set (~13 trillion combinations) of rigid rotations and tilting of the helices, then uses energy and diversity to select ~24 that are sufficiently stable to play a role in binding of ligands and activation and sufficiently diverse to represent structures that may be thermally accessible. Our GEnSeMBLE procedures led to successful GPCR structure predictions for C-C chemokine receptor type 5 (CCR5),(Berro, Yasmeen et al. 2013, Abrol, Trzaskowski et al. 2014) glucagon-like peptide-1 receptor (GLP1R),(Kirkpatrick A, Heo J et al. 2012) Cannabinoid receptor type 1 (CB1),(Scott, Abrol et al. 2013) Adenosine A3 receptor (AA3R),(Goddard III, Kim et al. 2010) bitter taste TAS2R38,(Tan, Abrol et al. 2012) prostaglandin D2 (PGD2R),(Shankar, Goddard et al. 2018) and κ-opioid receptor(Jeong, Kim et al. 2019). In addition, this GEnSeMBLE method has now been applied to predict 3D structures for 12 olfactory receptors out of the 400(Kim and Goddard III 2014, Malinska, Kim et al. 2021), Gi bound TAS2R4 and 5 structures out of the 25 bitter TAS2Rs(Yang, Kim et al. 2021, Yang, Mafi et al. 2021) and both the class C sweet taste TAS1R3/ 1R3 homodimer(Mafi, Kim et al. 2021b) and TAS1R2/ 1R3 heterodimer(Kim, Chen et al. 2017) receptors.

We recently studied G protein-first mechanism upon GPCR activation using ~20 μs long-scale molecular dynamics (MD) and metadynamics (metaD) simulations to 15 class A GPCRs, including opioids, adrenergics, adenosines, chemokines, muscarinics, cannabinoids, serotonins, and dopamines with Gs, Gi, Go, G11, and Gq protein (Mafi, Kim et al. 2020, Mafi, Kim et al. 2021, Mafi, Kim et al. 2022) We showed that binding of agonist to this pre-coupled GPCR-G protein complex causes the Gα protein to open into its active form, with the guanosine diphosphate exposed for signaling. We have also made dramatic advances on understanding the coupling of G protein or β-arrestin to opioid receptor using un-biased vs G protein biased agonists.(Mafi, Kim et al. 2020)

The missing side chains of E59, K60, N175, and S329 in the X-ray structure of APJ (PDB ID: 5VBL),(Ma, Yue et al. 2017) were constructed in Maestro. As shown in Fig. 1, the truncated extracellular loop 3 (EC3) between 230 and 242 was replaced by 55 residues of Rubredoxin (P00268) and then relaxed through simulated annealing with the fixed helixes (50 to 500K for 0.5 ps). Several mutations in EC2, TM3, TM5, TM6, and helix 8 (H8) in red were made. A119 (TM3), N177 (EC2), L368, and M369 (H8) are the same mutations in the X-ray structure of APJ (5VBL).

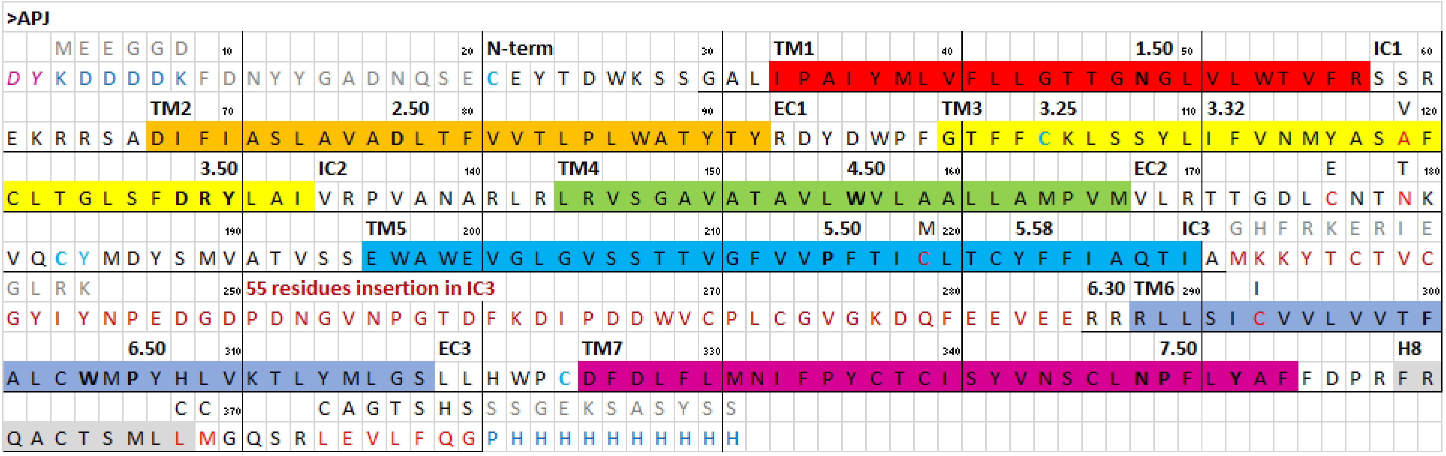

**Figure 1**. The sequence of APJ from GPCR Dock 2021. The N-terminal has two residues insertions in purple and modifications in blue. The intracellular loop 3 (IC3) has 55 residues insertions in (dark red). The C-terminal has 11 residue insertions in blue. There are mutations in EC2, TM3, TM5, TM6, and H8 in red. A119 (TM3), N177 (EC2), L368, and M369 (H8) are the same mutations in the X-ray structure of APJ (5VBL). Underlined sequences for GPCR Dock 2021 prediction. The grey residues are the original sequence from P35414 (fasta sequence of APJ).

We generated an ensemble of 24 structures for human APJ using the GEnSeMBLE method. (Abrol, Bray et al. 2011, Abrol, Griffith et al. 2012, Abrol, Kim et al. 2013, Bray, Abrol et al. 2014) For TM templates, we used the X-ray structure of APJ receptor in complex with agonist peptide (PDB ID: 5VBL). In the X-ray of APJ, two disulfide bridges were formed between C19 (N-term) and C281 (EC3) and between C102 (3.25) and C181 (EC2). The crystal structure of APJ showed the conserved TM1-2-7 networks among N1.50, D2.50 and N7.49 and TM2-3-4 networks among S2.45, T4.46, and W4.50 in the class A GPCRs, as shown in Fig. 2.

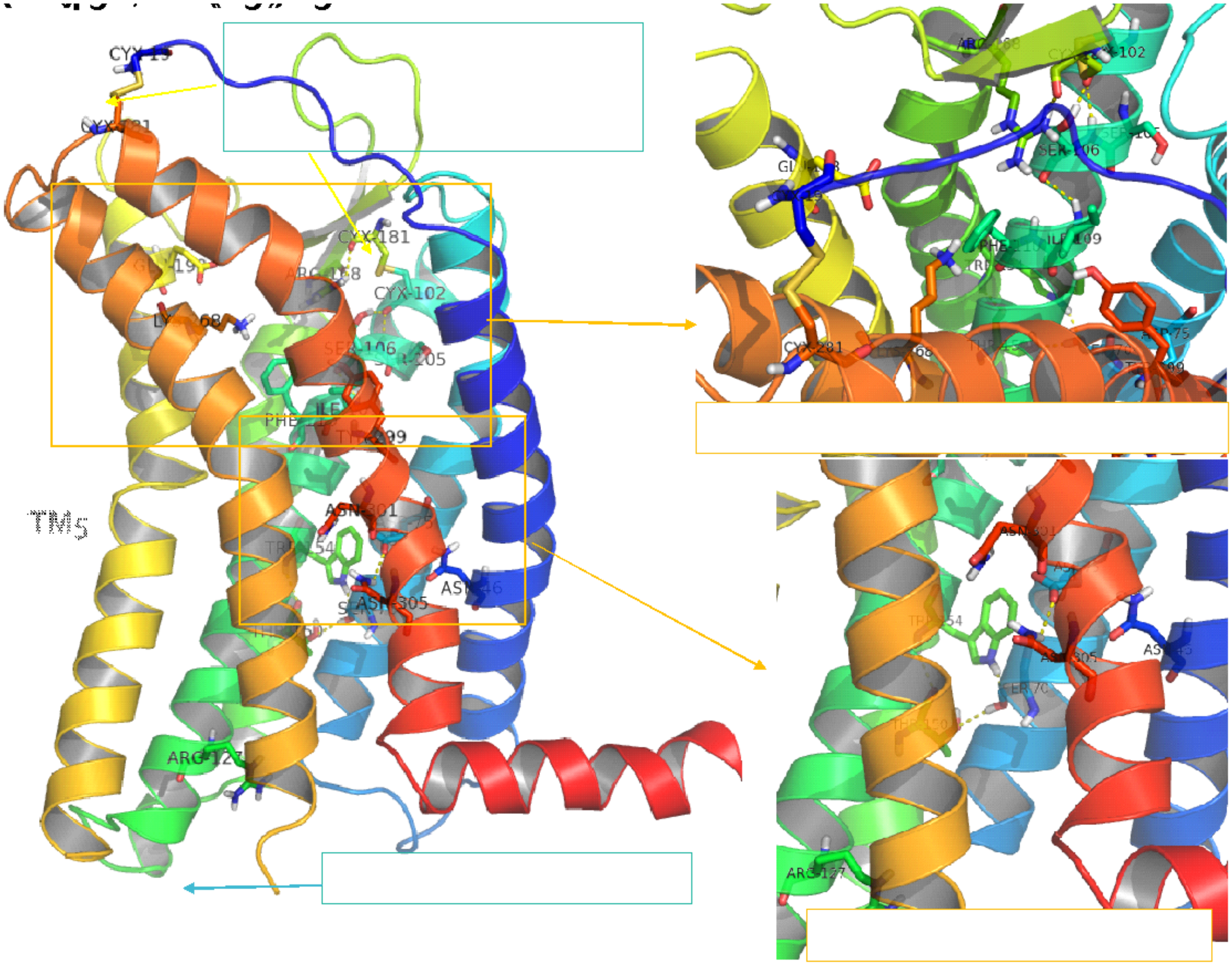

**Figure 2.** The 2.6-Å resolution crystal structure of human APJ receptor in complex with a designed 17-amino-acid apelin mimetic peptide agonist (PDB: 5VBL). In the X-ray of APJ, two disulfide bridges were formed between C19 (N-term) and C281 (EC3) and between C102 (3.25) and C181 (EC2). The crystal structure of APJ showed the conserved TM1-2-7 networks among N1.50, D2.50 and N7.49 and TM2-3-4 networks among S2.45, T4.46, and W4.50 in the class A GPCRs.

In the Top 25 of SuperBihelix (~13 trillion combinations), the all-zero structure which has the same rotation or tilting with the X-ray crystal structure of APJ ranked as Charge total (CTot), Neutral interhelical (Ni), Neutral total (Ntot), and the average rank of charge and neutral interhelical and total (CNTi) top 1. Inactive and Active contact analysis using currently available X-ray of class A GPCRs (Vaclav Cvicek, William A. Goddard III et al. 2016) shows that the highest inactive contact number is 7 from rank CNTi (Top 8, 16, 25). All top25 ensemble structures have zero active contact numbers, displaying inactive ensembles. For the docking study, we used the top 1 structure for the GPCR-Dock competition. Later we tried other 23 ensemble structures after adding all loops but we could not find better interactions.

**Table 1**. SuperBihelix CNTi rank of APJ. All-zero structure which has the same rotation or tilting with the X-ray crystal structure of APJ ranked as Charge total (CTot), Neutral interhelical (Ni), Neutral total (Ntot), and the average rank of charge and neutral interhelical and total (CNTi) top 1.

| RankCNti | H1 | H2 | H3 | H4 | H5 | H6 | H7 | H1 | H2 | H3 | H4 | H5 | H6 | H7 | H1 | H2 | H3 | H4 | H5 | H6 | H7 | CInterH | CTotal | NInterH | NTotal | RMSD |
|---|---|---|---|---|---|---|---|---|---|---|---|---|---|---|---|---|---|---|---|---|---|---|---|---|---|---|
| 1 | 0 | 0 | 0 | 0 | 0 | 0 | 0 | 0 | 0 | 0 | 0 | 0 | 0 | 0 | 0 | 0 | 0 | 0 | 0 | 0 | 0 | -643.55 | 372.76 | -488.94 | 350.78 | 0.31 |
| 2 | 0 | 0 | 0 | 0 | 0 | 0 | 0 | 0 | 0 | 0 | 0 | 0 | 0 | 0 | 0 | 0 | 0 | 0 | 0 | 0 | 0 | -643.55 | 372.76 | -488.94 | 350.78 | 0.00 |
| 3 | 0 | 0 | 0 | 0 | 0 | 0 | 0 | 0 | 0 | 0 | -15 | 0 | 0 | 0 | 0 | 0 | -15 | 15 | 0 | 0 | 0 | -633.12 | 397.35 | -470.17 | 377.33 | 0.47 |
| 4 | 0 | 0 | 0 | 0 | 0 | 0 | 0 | 0 | 0 | 0 | 0 | 0 | 0 | 0 | 0 | 0 | 0 | 15 | 0 | 0 | 15 | -659.73 | 426.69 | -488.90 | 434.25 | 0.50 |
| 5 | 0 | 0 | 0 | 0 | 0 | 0 | 0 | 0 | 0 | 0 | -15 | -15 | -15 | 0 | 0 | 0 | 0 | 0 | 0 | 0 | 15 | -636.98 | 440.84 | -468.50 | 411.20 | 0.74 |
| 6 | 0 | 0 | 0 | 0 | 0 | 0 | 0 | 0 | 0 | 0 | 0 | -30 | -15 | 0 | 0 | 0 | 0 | 15 | 15 | 0 | 0 | -646.06 | 408.32 | -461.43 | 422.88 | 1.02 |
| 7 | 0 | 0 | 0 | 0 | 0 | 0 | 0 | 0 | 0 | 0 | 0 | -15 | -15 | 0 | 0 | 0 | 0 | 0 | 15 | 0 | 15 | -632.22 | 432.64 | -461.13 | 413.86 | 0.78 |
| 8 | 0 | 0 | 0 | 0 | 0 | 0 | 0 | 0 | 0 | 0 | 0 | 0 | 0 | 15 | 0 | 0 | 0 | 0 | 0 | 0 | 15 | -637.59 | 439.96 | -466.67 | 429.97 | 0.79 |
| 9 | 0 | 0 | 0 | 0 | 0 | 0 | 0 | 0 | 0 | 0 | -15 | 0 | 0 | 15 | 0 | 0 | -15 | 15 | 0 | 0 | 0 | -624.04 | 434.18 | -464.14 | 401.36 | 0.78 |
| 10 | 0 | 0 | 0 | 0 | 0 | 0 | 0 | 0 | 0 | 0 | -15 | 0 | 0 | 15 | 0 | 0 | 0 | 0 | 0 | 0 | 15 | -648.60 | 448.97 | -472.16 | 435.47 | 0.79 |
| 11 | 0 | 0 | 0 | 0 | 0 | 0 | 0 | 0 | 0 | 0 | 15 | 0 | 0 | 15 | 0 | 0 | 0 | 0 | 0 | 0 | 15 | -642.14 | 444.86 | -466.26 | 433.60 | 0.80 |
| 12 | 0 | 0 | 0 | 0 | 0 | 0 | 0 | 0 | 0 | 0 | 0 | -15 | 0 | 0 | 0 | 0 | 0 | 15 | 0 | 0 | 15 | -660.39 | 455.36 | -487.87 | 439.40 | 0.63 |
| 13 | 0 | 0 | 0 | 0 | 0 | 0 | 0 | 0 | 0 | 0 | 0 | -15 | 0 | 15 | 15 | 0 | 0 | 15 | 0 | 0 | 0 | -627.55 | 453.78 | -459.98 | 414.41 | 0.83 |
| 14 | 0 | 0 | 0 | 0 | 0 | 0 | 0 | 0 | 0 | 0 | -15 | -15 | -15 | 0 | 0 | 0 | 0 | 0 | 15 | 0 | 15 | -625.31 | 442.64 | -457.45 | 422.81 | 0.79 |
| 15 | 0 | 0 | 0 | 0 | 0 | 0 | 0 | 0 | 0 | 0 | 0 | -15 | -15 | 0 | 0 | 0 | 0 | 0 | 0 | 0 | 15 | -628.56 | 446.12 | -456.72 | 427.14 | 0.73 |
| 16 | 0 | 0 | 0 | 0 | 0 | 0 | 0 | 0 | 0 | 0 | 0 | 0 | 0 | 0 | 0 | 0 | 0 | 15 | 15 | 0 | 0 | -619.78 | 434.88 | -458.55 | 415.15 | 0.47 |
| 17 | 0 | 0 | 0 | 0 | 0 | 0 | -10 | 0 | 0 | 0 | 0 | -15 | 0 | -15 | 0 | 0 | 0 | 15 | 0 | 0 | 0 | -647.07 | 463.14 | -472.65 | 441.36 | 1.16 |
| 18 | 0 | 0 | 0 | 0 | 10 | 0 | 0 | 0 | 0 | 0 | 0 | -30 | -30 | 0 | 0 | 0 | 0 | 15 | -30 | 0 | 0 | -626.67 | 405.83 | -448.93 | 394.71 | 1.72 |
| 19 | 0 | 0 | 0 | 0 | 0 | 0 | 0 | 0 | 0 | 0 | 0 | 0 | 0 | 15 | 0 | 0 | -15 | 15 | 0 | 0 | 0 | -619.51 | 429.88 | -456.75 | 415.46 | 0.77 |
| 20 | 0 | 0 | 0 | 0 | 0 | 0 | 0 | 0 | 0 | 0 | 15 | 0 | 0 | 15 | 0 | 0 | 0 | 0 | 0 | -15 | 15 | -631.34 | 449.60 | -456.16 | 430.18 | 0.85 |
| 21 | 0 | 0 | 0 | 0 | 0 | 0 | 0 | 0 | 0 | 0 | 0 | 0 | 0 | 15 | 0 | 0 | 0 | 15 | 0 | -15 | 15 | -633.70 | 449.26 | -458.48 | 443.06 | 0.86 |
| 22 | 0 | 0 | 0 | 0 | 0 | 0 | 0 | 0 | 0 | 0 | 15 | 0 | 0 | 0 | 0 | 0 | 0 | 0 | 15 | 0 | 0 | -622.75 | 443.72 | -456.38 | 428.96 | 0.43 |
| 23 | 0 | 0 | 0 | 0 | 0 | 0 | 0 | 0 | 0 | 0 | -15 | 0 | 0 | 15 | 0 | 0 | 0 | 15 | 0 | 0 | 15 | -627.32 | 456.41 | -457.08 | 430.76 | 0.82 |
| 24 | 0 | 0 | 0 | 0 | 0 | 0 | -10 | 0 | 0 | 0 | -15 | 0 | 0 | 0 | 0 | 0 | -15 | 15 | 0 | 0 | 0 | -625.90 | 432.62 | -451.49 | 440.97 | 1.02 |
| 25 | 0 | 0 | 0 | 0 | 0 | 0 | 0 | 0 | 0 | 0 | 0 | -30 | 0 | 0 | 0 | 0 | 0 | 15 | 0 | 0 | 0 | -625.44 | 458.48 | -455.79 | 428.53 | 0.87 |

**The DarwinDock method for predicting APJ-ligand structures:** We used the DarwinDock method based on complete sampling(Griffith 2017) to predict the ligand binding site to each of the 25 protein conformations from the GEnSeMBLE procedure. This procedure is as follows: We first alanized (replacing the 6 hydrophobic residues, I, L, V, F, Y, and W with A) the entire protein for each of the 24 protein conformations, to provide space for ligand docking. We then used DOCK4.0 to generate ~50,000 poses (but without energy calculations) sufficient to span the putative binding regions of the alanized protein. To ensure complete sampling, these poses are generated in increments of 5,000 and clustered into Voronoi families based on RMSD until <2% new families are generated. The family heads are energy scored using the Dreiding force field and the top10% by total energy are selected. Then we calculate the binding energy for all members of these top 10% families and selected the lowest energy 100 poses for further optimization. At this point we dealanize using SCREAM (Side Chain Rotamer Excitation Analysis Method) which is optimized to recognize side chains positions optimum for interactions between different chains (Victor Wai Tak Kam and William A. Goddard 2008). This leads to independent optimum side chains for each of the 100 poses. Applying this procedure to the antagonist JDTic ligand at κ-OR in the X-ray structure, leads to a structure deviating only 0.23 Å RMSD from the X-ray.(Wu, Wacker et al. 2012) This procedure led to successful GPCR bound ligand structure predictions for CCR5,(Berro, Yasmeen et al. 2013, Abrol, Trzaskowski et al. 2014) GLP1R,(Kirkpatrick A, Heo J et al. 2012) CB1,(Scott, Abrol et al. 2013) AA3R,(Goddard III, Kim et al. 2010) TAS2R38,(Tan, Abrol et al. 2012) PGD2R,(Shankar, Goddard et al. 2018) and TAS1R2/ 1R3 heterodimer(Kim, Chen et al. 2017).

We ran the conformational search of ligand in Maestro, generated a total of 525 conformations which we clustered into 15 conformations for docking. We docked each of these 15 conformations of the ligand into each of the 24 GPCR structures to select the final structures in Fig. 3 using various energies (CTot, Ni, Ntot, and CNTi) using DarwinDock method. this led to **top1.** Table 2 and 3 shows the unified cavity energy (UCav) rank (u#) and the binding energy (BE=complex E – protein E – ligand E) rank (b#), respectively. from the Total 20 structures in bod in Table 2 and 3 we did further Bindingsite Annealing. Table 4 displays the UCav rank after Bindingsite Annealing. For the final submissions to GPCR Dock 2021, we selected top 5 structures in Fig. 3, based on energy (rank1, 2: a1, a2), H-bonding numbers (rank3, 5: u1, b5), and diversity (rank4: u10).

**Table 2**. The unified cavity energy (UCav) rank (u#) of DarwinDock for APJ. Top structures in bod were selected for BindingsiteAnnealing.

| rank | filename | H-bonding | ucav | snapbe | snapbe_solv | note |
|---|---|---|---|---|---|---|
| u1.b7* | 5VBL_APJ_mut.SCHEMBL17166488.no_51.c29695.t.c.s.bgf | Y37,R170,K311,T338bb | -52.33 | -73.53 | 14.3 | overlap with X-ray, 4 HBs |
| u2 | 5VBL_APJ_mut.SCHEMBL17166488.no_24.c29106.t.c.s.bgf | Y95,S107,R170 | -51.33 | -63.61 | 5.98 | |
| u3 | 5VBL_APJ_mut.SCHEMBL17166488.no_51.c21547.t.c.s.bgf | S108,R170 | -50.96 | -65.46 | -12.69 | diversity |
| u4 | 5VBL_APJ_mut.SCHEMBL17166488.no_179.c16250.h.c.s.bgf | R170,Y314 | -50.33 | -63.12 | -7.35 | |
| u5.b10 | 5VBL_APJ_mut.SCHEMBL17166488.no_51.c22671.h.c.s.bgf | R170,T338bb | -49.49 | -72.16 | 20.02 | similar to u1 |
| u6.b11 | 5VBL_APJ_mut.SCHEMBL17166488.no_355.c3086.th.c.s.bgf | R170 | -48.62 | -71.97 | -2.27 | |
| u7.b14 | 5VBL_APJ_mut.SCHEMBL17166488.no_355.c20892.th.c.s.bgf | R170 | -47.45 | -70.13 | 20.77 | |
| u8 | 5VBL_APJ_mut.SCHEMBL17166488.no_179.c8229.th.c.s.bgf | R170,Y307 | -47.12 | -62.71 | -1.9 | |
| u9* | 5VBL_APJ_mut.SCHEMBL17166488.no_439.c29515.t.c.s.bgf | R170,K311 | -46.82 | -65.47 | 27.3 | overlap with X-ray |
| u10* | 5VBL_APJ_mut.SCHEMBL17166488.no_261.c32985.t.c.s.bgf | R170,Y342 | -46.43 | -54.56 | 14.99 | overlap with X-ray |
| u11 | 5VBL_APJ_mut.SCHEMBL17166488.no_261.c37687.t.c.s.bgf | R170,Y242 | -46.16 | -57.3 | -1.68 | |
| u12 | 5VBL_APJ_mut.SCHEMBL17166488.no_51.c2212.t.c.s.bgf | - | -46.03 | -59.2 | 24.66 | |
| u13.b12 | 5VBL_APJ_mut.SCHEMBL17166488.no_381.c25019.t.c.s.bgf | R170,Y307,R338bb | -45.84 | -71.33 | -10.5 | similar to u1 |
| u14.b13 | 5VBL_APJ_mut.SCHEMBL17166488.no_0.c35009.t.c.s.bgf | R170,Y342 | -45.63 | -70.68 | -6.87 | similar to u1 |
| u15 | 5VBL_APJ_mut.SCHEMBL17166488.no_51.c25289.t.c.s.bgf | Y37,Y184,R338bb | -45.26 | -63.06 | 25.93 | |
| u16 | 5VBL_APJ_mut.SCHEMBL17166488.no_17.c14956.t.c.s.bgf | Y307,Y342 | -45.24 | -62.02 | -3.76 | |
| u17 | 5VBL_APJ_mut.SCHEMBL17166488.no_200.c9815.t.c.s.bgf | R170,K311 | -44.77 | -57.54 | 22.23 | |

**Table 3**. The binding energy (BE=complex E – protein E – ligand E) rank (b#) of DarwinDock for APJ. Top structures in bod were selected for BindingsiteAnnealing.

| rank | filename | H-bonding | ucav | snapbe | snapbe_solv | note |
|---|---|---|---|---|---|---|
| b1* | 5VBL_APJ_mut.SCHEMBL17166488.no_51.c38430.t.c.s.bgf | R170,Y307,T338bb | -39.12 | -83.13 | -15.87 | similar to u1 |
| b2 | 5VBL_APJ_mut.SCHEMBL17166488.no_439.c22020.h.c.s.bgf | R170,C183bb,Y342 | -24.78 | -75.45 | 12.03 | |
| b3 | 5VBL_APJ_mut.SCHEMBL17166488.no_439.c28481.t.c.s.bgf | R170,K311 | -43.86 | -75.05 | 9.81 | |
| b4 | 5VBL_APJ_mut.SCHEMBL17166488.no_439.c33742.h.c.s.bgf | E200,K311 | -33.27 | -74.99 | -7.34 | |
| b5* | 5VBL_APJ_mut.SCHEMBL17166488.no_439.c43622.h.c.s.bgf | R170,E200,K311,Y307 | -28.98 | -74.28 | 9.9 | similar to u9, 4 HBs |
| b6 | 5VBL_APJ_mut.SCHEMBL17166488.no_439.c1452.t.c.s.bgf | - | -43.1 | -73.78 | 8.6 | |
| b7.u1 | 5VBL_APJ_mut.SCHEMBL17166488.no_51.c29695.t.c.s.bgf | - | -52.81 | -73.53 | 14.3 | |
| b8 | 5VBL_APJ_mut.SCHEMBL17166488.no_161.c39356.h.c.s.bgf | Y95,R170,K311 | -44.38 | -73.42 | 4.03 | diversity |
| b9 | 5VBL_APJ_mut.SCHEMBL17166488.no_161.c1512.t.c.s.bgf | R170,G204bb,K311 | -38.22 | -73.11 | 43.12 | similar to b8 |
| b10.u5 | 5VBL_APJ_mut.SCHEMBL17166488.no_51.c22671.h.c.s.bgf | Y37,R170,T338bb | -49.98 | -72.16 | 20.02 | |
| b11.u6 | 5VBL_APJ_mut.SCHEMBL17166488.no_355.c3086.th.c.s.bgf | R170,K311 | -49.82 | -71.97 | -2.27 | |
| b12.u13 | 5VBL_APJ_mut.SCHEMBL17166488.no_381.c25019.t.c.s.bgf | R170,Y307,T338bb | -45.38 | -71.33 | -10.5 | |
| b13.u14 | 5VBL_APJ_mut.SCHEMBL17166488.no_0.c35009.t.c.s.bgf | R170 | -45.94 | -70.68 | -6.87 | |
| b14.u7 | 5VBL_APJ_mut.SCHEMBL17166488.no_355.c20892.th.c.s.bgf | R170 | -49.23 | -70.13 | 20.77 | |
| b15 | 5VBL_APJ_mut.SCHEMBL17166488.no_439.c1997.h.c.s.bgf | R170,E200,S207,Y307 | -16.94 | -69.93 | -10.41 | |
| b16 | 5VBL_APJ_mut.SCHEMBL17166488.no_179.c24445.h.c.s.bgf | R170,Y307,T338bb | -44.1 | -69.07 | -2.66 | |
| b17 | 5VBL_APJ_mut.SCHEMBL17166488.no_439.c40212.h.c.s.bgf | Y95,R170,C183bb | -21.69 | -69.06 | 1.04 | |

**Table 4**. The unified cavity energy (UCav) rank after BindingsiteAnnealing (a#) from DarwinDock result for APJ

| rank | filename | H-bonding | ucav | snapbe | snapbe_solv | note |
|---|---|---|---|---|---|---|
| a1 | u3.5VBL_APJ.SCHEMBL17166488.no_51.c21547.t.c.s.f.a.bgf | W87,Y90,S108,R170 | -67.27 | -113.01 | -59.73 | Best E |
| a2 | u14.5VBL_APJ.SCHEMBL17166488.no_0.c35009.t.c.s.f.a.bgf | S108,R170,Y184,Y342 | -67.02 | -104.42 | -50.74 | 2nd best E |
| a3 | u1.5VBL_APJ.SCHEMBL17166488.no_51.c29695.t.c.s.f.a.bgf | R170 | -63.75 | -82.64 | -9.42 | |
| a4 | u13.5VBL_APJ.SCHEMBL17166488.no_381.c25019.t.c.s.f.a.bgf | I111bb,R170,K304,Y307 | -61.40 | -82.32 | -20.61 | similar to u1, 4HBs |
| a5 | b9.5VBL_APJ.SCHEMBL17166488.no_161.c1512.t.c.s.f.a.bgf | R170,S207 | -57.23 | -80.39 | -21.62 | |
| a6 | b8.5VBL_APJ.SCHEMBL17166488.no_161.c39356.h.c.s.f.a.bgf | H308 | -56.97 | -69.09 | -14.92 | |
| a7 | u5.5VBL_APJ.SCHEMBL17166488.no_51.c22671.h.c.s.f.a.bgf | R170 | -56.14 | -70.55 | 28.24 | |
| a8 | u2.5VBL_APJ.SCHEMBL17166488.no_24.c29106.t.c.s.f.a.bgf | W87,R170 | -55.79 | -89.04 | -46.50 | |
| a9 | b2.5VBL_APJ.SCHEMBL17166488.no_439.c22020.h.c.s.f.a.bgf | W87,Y90,S107,R170 | -55.63 | -83.33 | -24.89 | |
| a10 | b5.5VBL_APJ.SCHEMBL17166488.no_439.c43622.h.c.s.f.a.bgf | R170,Y184,K311 | -55.52 | -89.95 | -22.76 | |
| a11 | u8.5VBL_APJ.SCHEMBL17166488.no_179.c8229.th.c.s.f.a.bgf | R170 | -55.46 | -63.26 | -15.56 | |
| a12 | u10.5VBL_APJ.SCHEMBL17166488.no_261.c32985.t.c.s.f.a.bgf | Y42,R170 | -54.29 | -77.65 | -9.64 | |
| a13 | u6.5VBL_APJ.SCHEMBL17166488.no_355.c3086.th.c.s.f.a.bgf | R170,E200,K311 | -54.00 | -89.89 | -36.88 | |
| a14 | b6.5VBL_APJ.SCHEMBL17166488.no_439.c1452.t.c.s.f.a.bgf | R170,K311 | -53.23 | -83.70 | -16.58 | |
| a15 | u4.5VBL_APJ.SCHEMBL17166488.no_179.c16250.h.c.s.f.a.bgf | R170,Y314 | -52.92 | -81.91 | 2.28 | |
| a16 | b1.5VBL_APJ.SCHEMBL17166488.no_51.c38430.t.c.s.f.a.bgf | T338 | -52.81 | -64.12 | -0.55 | |
| a17 | b3.5VBL_APJ.SCHEMBL17166488.no_439.c28481.t.c.s.f.a.bgf | R170 | -52.22 | -80.89 | -27.41 | |
| a18 | u9.5VBL_APJ.SCHEMBL17166488.no_439.c29515.t.c.s.f.a.bgf | R170 | -51.57 | -78.49 | -21.67 | |
| a19 | u7.5VBL_APJ.SCHEMBL17166488.no_355.c20892.th.c.s.f.a.bgf | E200,Y307,K311 | -41.96 | -70.21 | -16.42 | |
| a20 | b4.5VBL_APJ.SCHEMBL17166488.no_439.c33742.h.c.s.f.a.bgf | Y307,K311 | -35.48 | -59.96 | -15.98 | |

[illegible]

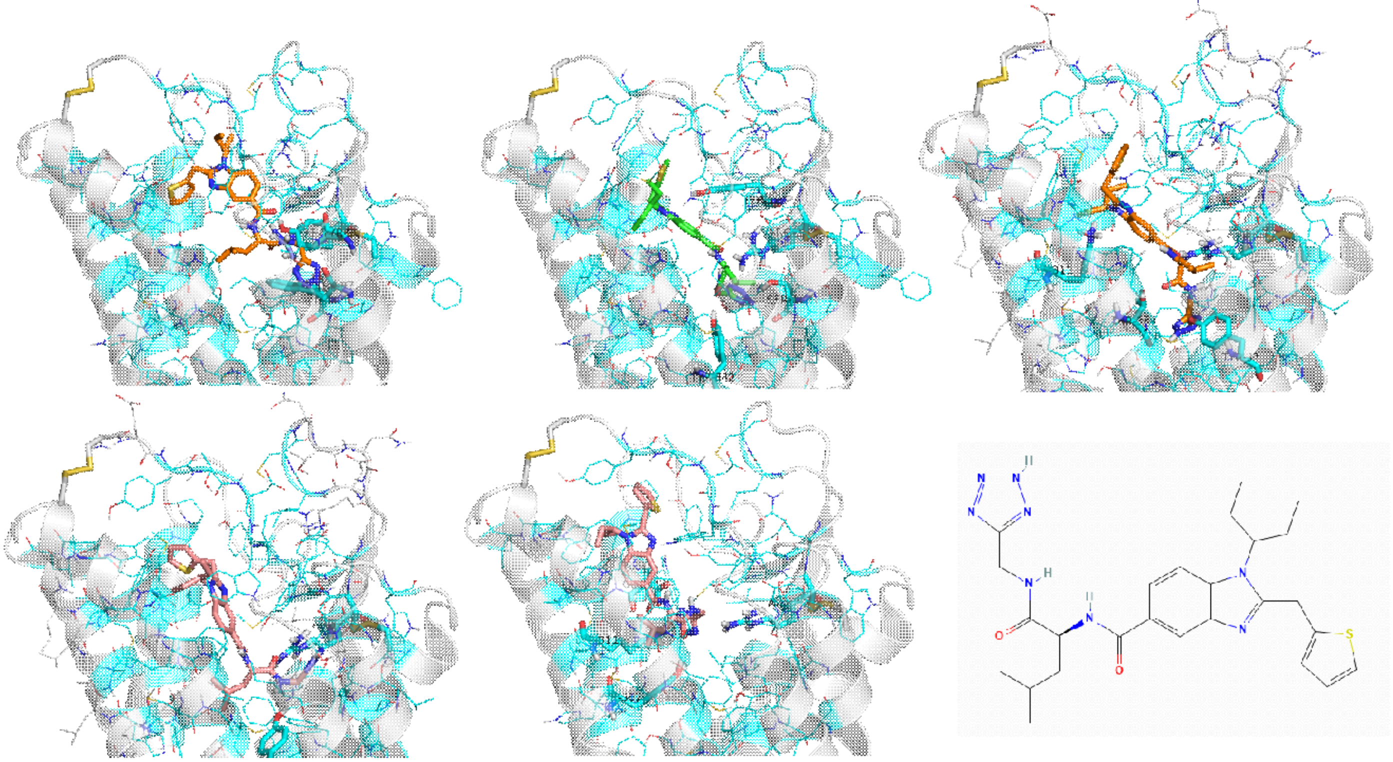

**Figure 3.** The top 5 structures of APJ-ligand complex from DarwinDock plus the ligand structure

In the final results, our model rank 4 and rank3 ranked as top3 with the ligand RMSD of 4.05 Å and top 8 with the ligand RMSD of 4.00 Å out of total 201 submitted models, respectively. Although model 3 showed less ligand RMSD, model 4 displayed better contact predictions accuracy of 23.3 %, compared to 13.0 % in model 3. In Fig. 4, our model rank 4 revealed two H-bonds at R170 with two carbonyl group. W87, Y314, and F334 formed a pi-pi stacking interaction. K311 made a pi-cation interaction. However, there is only one pi-pi interaction with F291 in the X-ray structure.

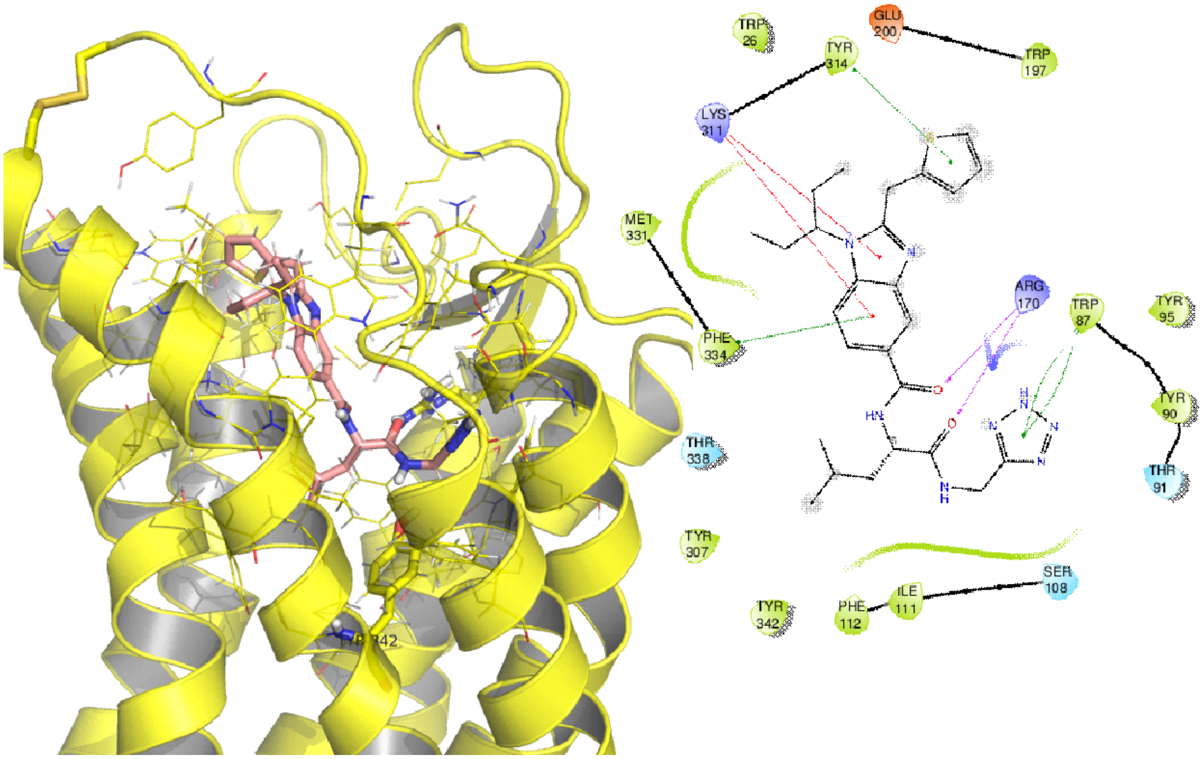

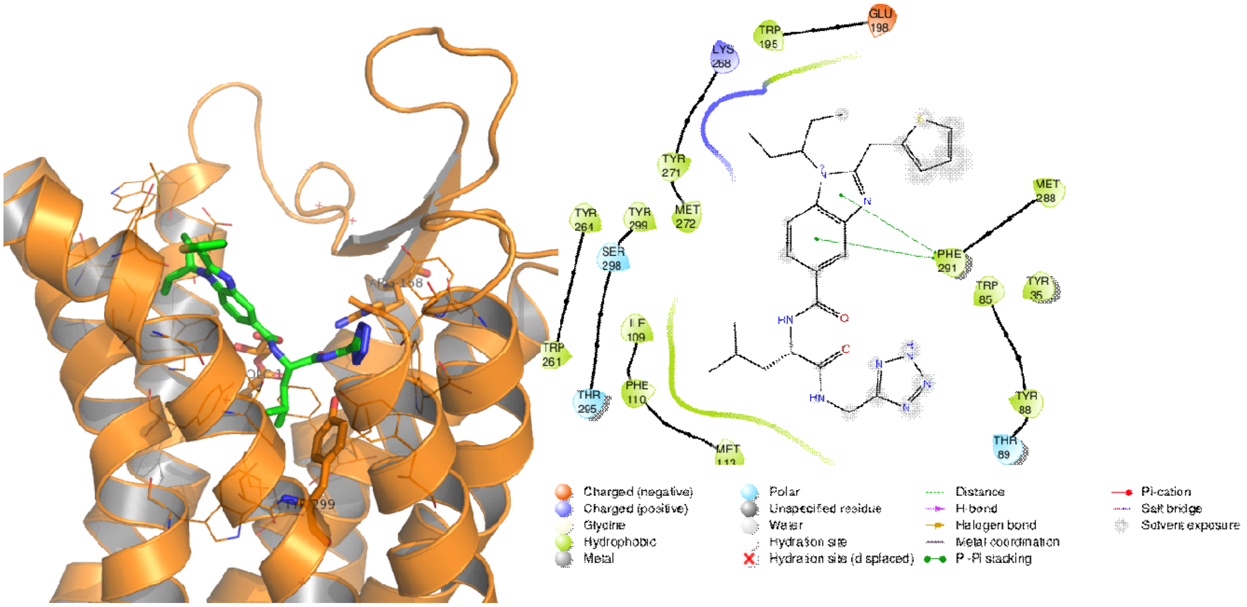

**Figure 4.** The binding site and ligand interaction diagram of (Top) our model rank 4 from the docking study (UCav 10) and (Bottom) the X-ray structure of the APJ-ligand complex

After the competition, we ran 20ns MD simulation of all five models with the backbone constraints for 10ns and then the helix backbone/ and helix constraints for another 10ns including explicit lipid and water. We inserted the GPCR-ligand bound complex into a palmitoyl-oleoyl-phosphatidylcholine (POPC) lipid membrane containing full solvent. We then added ions to neutralize the system while matching to physiological salt conditions. With the protein and ligand fixed, the solvent and membrane were first minimized with NAMD using the CHARMM36m force field (improved for folded and intrinsically disordered proteins).(Liu, Huang et al. 2014) The temperature of the system was maintained using Langevin dynamics (1 atm) with a damping coefficient of 1 (gamma) and a bath temperature of 310 K. We then carried out MD simulations until convergence to validate and optimize the ligand binding from the docking predictions. Based on the interaction energy between ligand and protein, our model rank3 (UCav 1 from DarwinDock) gave the best average and minimum total energies in Table 5.

**Table 5**. The average interaction and minimum energies between ligand and protein during MD simulation of five models

| Compound | Ave-ELECT(0-20ns) | Ave-VDW(0-20ns) | Ave-TOT(0-20ns) | Ave-ELECT(10-20ns) | Ave-VDW(10-20ns) | Ave-TOT(10-20ns) | Min-ELECT | Min-VDW | Min-TOT |
|---|---|---|---|---|---|---|---|---|---|
| rank1 | -21.24 | -60.18 | -81.42 | -20.06 | -62.08 | -82.14 | -40.45 | -70.24 | -98.16 |
| rank2 | -23.84 | -53.24 | -77.08 | -29.17 | -53.16 | -82.32 | -45.15 | -60.68 | -97.44 |
| rank3 | -22.09 | -60.05 | -82.14 | -22.91 | -59.80 | -82.71 | -37.23 | -67.99 | -99.35 |
| rank4 | -20.94 | -51.89 | -72.83 | -16.96 | -52.49 | -69.45 | -50.35 | -58.65 | -95.25 |
| rank5 | -23.87 | -53.96 | -77.84 | -23.42 | -55.34 | -78.76 | -45.09 | -60.33 | -99.24 |

From the superimposition of our model, rank 4 in pink from DarwinDock (UCav10), and the X-ray structure in green of APJ-ligand complex, they overlapped well in the same binding site, as shown in Fig. 5 left. The major deviation is at the upper TM region close to the extracellular loops. However, this deviation was minimized after 20ns MD simulation (Fig. 5 right). The average ligand and protein RMSD for 20ns MD were 1.83 Å and 0.86 Å, respectively.

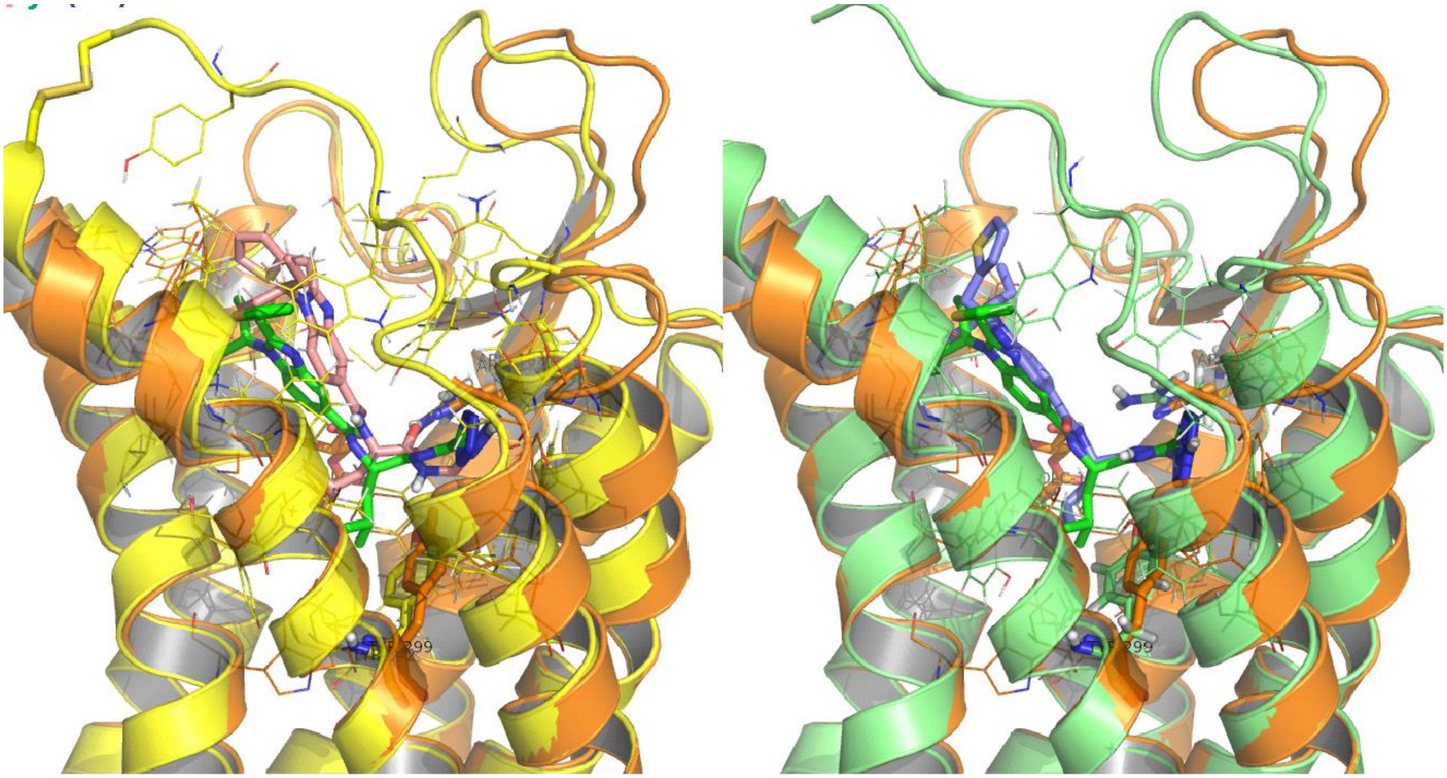

**Figure 5.** The superimposition of (Left) our model rank 4 from DarwinDock (UCav10) in pink with ligand RMSD of 4.05 Å and (Right) 20ns MD structure with the X-ray structure in green of the APJ-ligand complex.

**GPCRdock-3183 – GPR139**

#### 1. Methods

**Ensemble structure generation for orphan G protein-coupled receptor (GPCR), GPR139:** We predicted the ensemble of low energy GPR139 structures using the GEnSeMBLE complete sampling method. (Abrol, Bray et al. 2011, Abrol, Griffith et al. 2012, Abrol, Kim et al. 2013, Bray, Abrol et al. 2014) GEnSeMBLE identifies the transmembrane (TM) domains based on multiple sequence alignments, then selects the shape of each helix based on the force field or on homology to known structures, and then considers a complete set (~13 trillion combinations) of rigid rotations and tilting of the helices, then uses energy and diversity to select ~24 that are sufficiently stable to play a role in binding of ligands and activation and sufficiently diverse to represent structures that may be thermally accessible. Our GEnSeMBLE procedures led to successful GPCR structure predictions for C-C chemokine receptor type 5 (CCR5),(Berro, Yasmeen et al. 2013, Abrol, Trzaskowski et al. 2014) glucagon-like peptide-1 receptor (GLP1R),(Kirkpatrick A, Heo J et al. 2012) Cannabinoid receptor type 1 (CB1),(Scott, Abrol et al. 2013) Adenosine A3 receptor (AA3R),(Goddard III, Kim et al. 2010) bitter taste TAS2R38,(Tan, Abrol et al. 2012) prostaglandin D2 (PGD2R),(Shankar, Goddard et al. 2018) and κ-opioid receptor (κ-OR)(Mafi, Kim et al. 2020). In addition, this GEnSeMBLE method has now been applied to predict 3D structures for 12 olfactory receptors (out of the 400)(Kim and Goddard III 2014, Malinska, Kim et al. 2021), Gi bound TAS2R4 and 5 structures out of the 25 bitter TAS2Rs(Yang, Kim et al. 2021, Yang, Mafi et al. 2021) and both the class C sweet taste TAS1R3/ 1R3 homodimer(Mafi, Kim et al. 2021b) and TAS1R2/ 1R3 heterodimer(Kim, Chen et al. 2017) receptors.

**The DarwinDock method for predicting GPR139-ligand structures:** We use DarwinDock complete sampling method(Griffith 2017) to predict the ligand binding site to each of the 25 protein conformations from the GEnSeMBLE procedure. This procedure is as follows: We first alanized (replacing the 6 hydrophobic residues, I, L, V, F, Y, and W with A) the entire protein for each of the 24 ensemble protein conformations, to provide space for ligand docking. We then used DOCK4.0 to generate ~50,000 poses (but without energy calculations) sufficient to span the putative binding regions of the alanized protein. To ensure complete sampling, these poses are generated in increments of 5,000 and clustered into Voronoi families based on RMSD until <2% new families are generated. The family heads are energy scored using the Dreiding force field and the top10% by total energy are selected. Then we calculate the binding energy for all members of these top 10% families and selected the lowest 100 poses for further optimization. At this point we dealanize using SCREAM (Side Chain Rotamer Excitation Analysis Method) which is optimized to recognized side chains positions optimum for interactions between different chains (Victor Wai Tak Kam and William A. Goddard 2008). This lead to optimize the side chains independently for each of the 100 poses. Applying this procedure to the antagonist JDTic ligand at κ-OR in the X-ray structure, leads to a structure deviating only 0.23 Å RMSD from the X-ray.(Wu, Wacker et al. 2012) This procedure led to successful GPCR bound ligand structure predictions for CCR5,(Berro, Yasmeen et al. 2013, Abrol, Trzaskowski et al. 2014) GLP1R,(Kirkpatrick A, Heo J et al. 2012) CB1,(Scott, Abrol et al. 2013) AA3R,(Goddard III, Kim et al. 2010) TAS2R38,(Tan, Abrol et al. 2012) PGD2R,(Shankar, Goddard et al. 2018) and TAS1R2/ 1R3 heterodimer(Kim, Chen et al. 2017).

#### 2. Results

##### 2.1 Predictions of the ensemble structures for GPR139.

We generated an ensemble of structures for human GPR139 using the GEnSeMBLE method. (Abrol, Bray et al. 2011, Abrol, Griffith et al. 2012, Abrol, Kim et al. 2013, Bray, Abrol et al. 2014) For TM templates we used four Gq/11 coupled Cryo-electron microscopy (Cryo-EM) structures and one predicted structure from molecular dynamic (MD) simulation, 1) Cholecystokinin CCK1 (PDB ID code: 7MBY), 2) histamine H1 (PDB ID code: 7DFL), 3) 5-Hydroxytryptamine 5-HT2A (PDB ID code: 6WHA), 4) Acetylcholine M1 receptor (PDB ID code: 6OIJ), and 5) 5-HT2A MD structure for various initial conformers.
From the Top 25 of GPR139 out of ~200 million (=$6*12^7$) BiHelix structures for the active GPR139 structure, we selected by combinations of best energy, Charge total (CTot), Neutral interhelical (Ni), Neutral total (Ntot), and the average rank of charge and neutral interhelical and total (CNTi), and diversity 5 for optimization of tilts: CNTi rank Top 1, 4, 7 in Table 1 and CNi rank Top 3 and 10 in Table 2 (CNi rank Top 1 and 5 are overlapped with Table 1) were selected for SuperBihelix run to optimize the tilts. All-zero structure which has the same rotation

or tilting with the X-ray crystal structure with 5-HT2A MD structure ranked as CNTi top 7. Most of the BiHelix top 25 results were from active from the MD structure of 5-HT2A and Cholecystokinin CCK1 receptor.

**Table 1**. Bihelix CNTi rank of GPR139. Top 1, 4, 7 in yellow were selected for SuperBihelix run. All 0 structure rank Top7.

| CNTi rank | name | # active | # inactive | Active-Inactive | #Class A Contacts |
|---|---|---|---|---|---|
| 1 | **HT2AR-metaD: 0_0_0_0_0_0_0x0_0_0_0_0_0_0x0_-30_0_0_0_0_0x37.7_74.9_112.6_155.9_192.8_241.1_275.5** | 2 | 1 | 1 | 33 |
| 2 | HT2AR-metaD: 0_0_0_0_0_0_0x0_0_0_0_0_0_0x90_0_0_0_0_0_0x37.7_74.9_112.6_155.9_192.8_241.1_275.5 | 1 | 1 | 0 | 28 |
| 3 | HT2AR-metaD: 0_0_0_0_0_0_0x0_0_0_0_0_0_0x0_-30_0_0_180_0_0x37.7_74.9_112.6_155.9_192.8_241.1_275.5 | 0 | 0 | 0 | 28 |
| 4 | **HT2AR-metaD: 0_0_0_0_0_0_0x0_0_0_0_0_0_0x-60_-30_0_0_0_-30_-30x37.7_74.9_112.6_155.9_192.8_241.1_275.5** | 4 | 2 | 2 | 27 |
| 5 | HT2AR-metaD: 0_0_0_0_0_0_0x0_0_0_0_0_0_0x0_-30_0_0_-60_0_0x37.7_74.9_112.6_155.9_192.8_241.1_275.5 | 2 | 0 | 2 | 29 |
| 6 | HT2AR-metaD: 0_0_0_0_0_0_0x0_0_0_0_0_0_0x0_-30_0_0_0_-30_-30x37.7_74.9_112.6_155.9_192.8_241.1_275.5 | 4 | 2 | 2 | 31 |
| 7 | **HT2AR-metaD: 0_0_0_0_0_0_0x0_0_0_0_0_0_0x0_0_0_0_0_0_0x37.7_74.9_112.6_155.9_192.8_241.1_275.5** | 2 | 1 | 1 | 33 |
| 8 | HT2AR-metaD: 0_0_0_0_0_0_0x0_0_0_0_0_0_0x0_0_0_0_0_0_0x37.7_74.9_112.6_155.9_192.8_241.1_275.5 | 2 | 1 | 1 | 33 |
| 9 | HT2AR-metaD: 0_0_0_0_0_0_0x0_0_0_0_0_0_0x30_-30_0_0_0_-30_-30x37.7_74.9_112.6_155.9_192.8_241.1_275.5 | 4 | 2 | 2 | 27 |
| 10 | HT2AR-metaD: 0_0_0_0_0_0_0x0_0_0_0_0_0_0x0_-30_0_0_120_0_0x37.7_74.9_112.6_155.9_192.8_241.1_275.5 | 0 | 1 | -1 | 28 |
| 11 | HT2AR-metaD: 0_0_0_0_0_0_0x0_0_0_0_0_0_0x180_0_0_0_0_-30_-30x37.7_74.9_112.6_155.9_192.8_241.1_275.5 | 3 | 1 | 2 | 23 |
| 12 | HT2AR-metaD: 0_0_0_0_0_0_0x0_0_0_0_0_0_0x0_-30_0_0_90_0_0x37.7_74.9_112.6_155.9_192.8_241.1_275.5 | 0 | 0 | 0 | 28 |
| 13 | HT2AR-metaD: 0_0_0_0_0_0_0x0_0_0_0_0_0_0x0_-30_0_0_-90_0_0x37.7_74.9_112.6_155.9_192.8_241.1_275.5 | 1 | 1 | 0 | 28 |
| 14 | HT2AR-metaD: 0_0_0_0_0_0_0x0_0_0_0_0_0_0x-90_-30_0_0_0_0_0x37.7_74.9_112.6_155.9_192.8_241.1_275.5 | 0 | 1 | -1 | 26 |
| 15 | HT2AR-metaD: 0_0_0_0_0_0_0x0_0_0_0_0_0_0x30_-30_0_120_0_-30_-30x37.7_74.9_112.6_155.9_192.8_241.1_275.5 | 3 | 1 | 2 | 21 |
| 16 | HT2AR-metaD: 0_0_0_0_0_0_0x0_0_0_0_0_0_0x-60_-30_0_120_0_-30_-30x37.7_74.9_112.6_155.9_192.8_241.1_275.5 | 4 | 2 | 2 | 21 |
| 17 | HT2AR-metaD: 0_0_0_0_0_0_0x0_0_0_0_0_0_0x-90_-30_0_0_0_-30_0x37.7_74.9_112.6_155.9_192.8_241.1_275.5 | 4 | 1 | 3 | 26 |
| 18 | HT2AR-metaD: 0_0_0_0_0_0_0x0_0_0_0_0_0_0x-90_-30_0_0_180_0_0x37.7_74.9_112.6_155.9_192.8_241.1_275.5 | 0 | 0 | 0 | 21 |
| 19 | HT2AR-metaD: 0_0_0_0_0_0_0x0_0_0_0_0_0_0x90_0_0_0_180_0_0x37.7_74.9_112.6_155.9_192.8_241.1_275.5 | 0 | 1 | -1 | 23 |
| 20 | HT2AR-metaD: 0_0_0_0_0_0_0x0_0_0_0_0_0_0x0_-30_0_120_-30_0_0x37.7_74.9_112.6_155.9_192.8_241.1_275.5 | 2 | 1 | 1 | 25 |
| 21 | HT2AR-metaD: 0_0_0_0_0_0_0x0_0_0_0_0_0_0x0_-30_0_0_-150_0_0x37.7_74.9_112.6_155.9_192.8_241.1_275.5 | 0 | 0 | 0 | 32 |
| 22 | HT2AR-metaD: 0_0_0_0_0_0_0x0_0_0_0_0_0_0x60_0_0_0_0_-30_-30x37.7_74.9_112.6_155.9_192.8_241.1_275.5 | 3 | 1 | 2 | 26 |
| 23 | HT2AR-metaD: 0_0_0_0_0_0_0x0_0_0_0_0_0_0x0_-30_0_0_-120_-30_0x37.7_74.9_112.6_155.9_192.8_241.1_275.5 | 1 | 0 | 1 | 27 |
| 24 | HT2AR-metaD: 0_0_0_0_0_0_0x0_0_0_0_0_0_0x0_-30_0_0_-30_-30_0x37.7_74.9_112.6_155.9_192.8_241.1_275.5 | 0 | 0 | 0 | 32 |
| 25 | HT2AR-metaD: 0_0_0_0_0_0_0x0_0_0_0_0_0_0x0_0_0_0_-90_0_0x37.7_74.9_112.6_155.9_192.8_241.1_275.5 | 2 | 1 | 1 | 31 |

**Table 2**. Bihelix CNi rank of GPR139. Top 3 and 10 in orange were selected for SuperBihelix run, since Top 1 and 5 in yellow overlapped with CNTi rank in Table 1.

| RankCNi | H1 | H2 | H3 | H4 | H5 | H6 | H7 | Source | CintraH | CInterH | CTotal | NIntraH | NInterH | NTotal |
|---|---|---|---|---|---|---|---|---|---|---|---|---|---|---|
| 1 | **0** | **-30** | **0** | **0** | **0** | **0** | **0** | **HT2AR-metaD** | 964.00 | -548.30 | 416.17 | 585.95 | -395.78 | 190.17 |
| 2 | 0 | -30 | 0 | 0 | 180 | 0 | 0 | HT2AR-metaD | 961.64 | -546.06 | 416.30 | 581.22 | -391.98 | 189.24 |
| 3 | **0** | **0** | **0** | **0** | **-30** | **-30** | **30** | **CCK1R** | 1096.92 | -525.36 | 571.39 | 776.82 | -414.70 | 362.12 |
| 4 | 0 | -30 | 0 | 0 | -30 | -30 | 0 | HT2AR-metaD | 1022.43 | -547.17 | 476.25 | 650.20 | -388.18 | 262.01 |
| 5 | **-60** | **-30** | **0** | **0** | **0** | **-30** | **-30** | **HT2AR-metaD** | 966.36 | -562.06 | 404.37 | 592.23 | -386.86 | 205.37 |
| 6 | 90 | 0 | 0 | 0 | 0 | 0 | 0 | HT2AR-metaD | 910.77 | -538.79 | 372.76 | 545.52 | -385.44 | 160.09 |
| 7 | 180 | 0 | 0 | 0 | 180 | 0 | 120 | CCK1R | 1192.10 | -521.87 | 670.07 | 853.25 | -388.36 | 464.89 |
| 8 | 0 | -30 | 0 | 0 | -60 | 0 | 0 | HT2AR-metaD | 953.05 | -528.45 | 425.08 | 557.27 | -383.61 | 173.66 |
| 9 | 30 | -30 | 0 | 0 | 0 | -30 | -30 | HT2AR-metaD | 960.19 | -543.08 | 417.27 | 577.90 | -379.56 | 198.34 |
| 10 | **0** | **0** | **0** | **0** | **-60** | **-30** | **30** | **CCK1R** | 1109.17 | -512.15 | 597.25 | 805.45 | -392.24 | 413.21 |
| 11 | 0 | -30 | 0 | 0 | 0 | -30 | -30 | HT2AR-metaD | 935.83 | -540.03 | 395.79 | 576.29 | -378.56 | 197.73 |
| 12 | 30 | -30 | 0 | 120 | 0 | -30 | -30 | HT2AR-metaD | 993.44 | -524.53 | 469.13 | 616.76 | -381.76 | 235.00 |
| 13 | 0 | -30 | 0 | 120 | -30 | -30 | 0 | HT2AR-metaD | 1013.38 | -513.70 | 500.32 | 651.35 | -387.80 | 263.55 |
| 14 | 0 | -30 | 0 | 180 | -90 | 0 | 0 | HT2AR-metaD | 1017.68 | -531.88 | 485.50 | 647.56 | -378.15 | 269.41 |
| 15 | 0 | -30 | 0 | 0 | 120 | 0 | 0 | HT2AR-metaD | 957.43 | -543.24 | 414.49 | 569.08 | -376.74 | 192.34 |
| 16 | 0 | 0 | 0 | 0 | 0 | 0 | 0 | HT2AR-metaD | 932.35 | -527.65 | 405.54 | 566.40 | -378.18 | 188.22 |
| 17 | 0 | 0 | 0 | 0 | 0 | 0 | 0 | HT2AR-metaD | 932.35 | -527.65 | 405.54 | 566.40 | -378.18 | 188.22 |
| 18 | -30 | 0 | 0 | 0 | -30 | -30 | 120 | CCK1R | 1180.09 | -507.01 | 673.02 | 864.92 | -411.03 | 453.89 |
| 19 | 180 | 0 | 0 | 0 | 0 | -30 | -30 | HT2AR-metaD | 945.49 | -518.26 | 427.39 | 597.39 | -379.64 | 217.75 |
| 20 | -60 | -30 | 0 | 120 | 0 | -30 | -30 | HT2AR-metaD | 979.94 | -525.02 | 455.02 | 623.87 | -377.81 | 246.06 |
| 21 | -30 | -90 | 30 | -90 | 0 | 0 | 60 | HT2AR | 1142.53 | -527.09 | 615.37 | 821.25 | -375.40 | 445.85 |
| 22 | 0 | 0 | 0 | 0 | -30 | -30 | -30 | CCK1R | 1150.81 | -503.71 | 647.17 | 811.67 | -398.74 | 412.93 |
| 23 | -90 | -30 | 0 | 0 | 0 | 0 | 0 | HT2AR-metaD | 972.19 | -515.22 | 457.60 | 564.48 | -376.52 | 187.96 |
| 24 | 0 | 0 | 0 | 0 | -30 | -30 | 0 | CCK1R | 1153.58 | -507.30 | 646.08 | 818.20 | -382.49 | 435.72 |
| 25 | 180 | 0 | 0 | 0 | -30 | -30 | 30 | CCK1R | 1153.32 | -504.57 | 648.76 | 832.59 | -387.85 | 444.74 |

In SuperBihelix (~13 trillion combinations), total 18 structures (rank CNTi (5) + Rank Ni (5) + Rank Ci (4) + Rank CNi (4) ) in bond in Table 3 were selected for docking studies. There are two overlapped structures in italics in Table 3. Active contact analysis using currently available X-ray of Class A GPCRs (Vaclav Cvicek, William A. Goddard III et al. 2016) shows that the highest active contact numbers is 7 from rank Ni (Top1, 4) and CNi (Top6, 11).

**Table 3**. SuperBihelix CNTi, Ni, Ci, CNi rank of GPR139. Total 18 structures (rank CNTi (5) + Rank Ni (5) + Rank Ci (4) + Rank CNi (4) ) in bold were selected for docking studies. Overlapped structures are displayed in italics.

| RankCNTi | name |
|---|---|
| 1 | 01_HT2AR-metaD: -10_-10_-10_10_0_0_-10x15_-30_30_-30_0_-30_15x-15_30_0_0_-15_-15_15x37.7_ |
| 5 | 01_HT2AR-metaD: -10_-10_-10_10_0_-10_-10x15_-30_30_-30_-30_-30_15x-15_30_0_0_15_15_15x37 |
| 8 | 01_HT2AR-metaD: -10_-10_-10_10_0_0_-10x15_-30_30_-30_-30_-30_15x-15_30_0_0_0_-15_15x37.7_ |
| 12 | 01_HT2AR-metaD: -10_-10_-10_10_0_-10_-10x15_-30_30_-30_-30_-15_15x-15_30_0_0_0_-15_15x37 |
| 24 | 01_HT2AR-metaD: -10_-10_-10_10_0_0_-10x0_-30_30_-30_0_-30_15x-15_30_0_0_30_-15_15x37.7_7 |

| RankNi | name |
|---|---|
| 1 | 03_CCK1R: -10_0_0_0_0_0_0x15_0_0_15_0_-15_30x0_0_0_0_0_-15x37.0_75.5_112.3_155.7_191.8_ |
| 4 | 03_CCK1R: -10_0_0_0_0_0_0x0_0_0_15_0_-15_15x0_0_0_0_0_0x37.0_75.5_112.3_155.7_191.8_240 |
| 6 | 61_CCK1R: -10_-10_0_-10_10_0_-10x0_0_15_30_-15_0_-15x15_-15_0_0_15_0_0x37.0_75.5_112.3_15 |
| 13 | 03_CCK1R: -10_0_0_0_0_0_-10x15_0_0_15_0_0_0x0_0_0_0_-15_-30x37.0_75.5_112.3_155.7_191.8 |
| 15 | 03_CCK1R: -10_0_0_0_0_0_-10x15_0_0_30_0_-15_-15x0_0_0_0_0_-15x37.0_75.5_112.3_155.7_191 |

| RankCi | name |
|---|---|
| 1 | *01_HT2AR-metaD: -10_-10_-10_10_0_0_-10x15_-30_30_-30_0_-30_15x-15_30_0_0_-15_-15_15x37.7_* |
| 7 | 61_CCK1R: -10_0_0_0_0_-10_0x0_-15_0_-30_-30_15_15x0_0_0_-15_15_0_30x37.0_75.5_112.3_155.7_ |
| 13 | 01_HT2AR-metaD: -10_-10_-10_10_0_-10_-10x15_-30_30_-30_-30_0_15x-15_30_0_0_0_-30_15x37.7_ |
| 19 | 03_CCK1R: -10_0_0_0_10_0_-10x15_0_0_-15_-15_-30_15x0_0_-15_0_30_30_0x37.0_75.5_112.3_155. |
| 25 | 61_CCK1R: -10_0_0_0_0_-10_0x0_-15_0_15_-30_15_15x0_0_0_-15_15_0_30x37.0_75.5_112.3_155.7_ |

| RankCNi | name |
|---|---|
| 1 | *61_CCK1R:-10_0_0_0_0_0-10x0_0_0_0_0_0_-15x15_-15_0_0_0_0_0x37.0_75.5_112.3_155.7_191.8_* |
| 6 | 03_CCK1R:-10_0_0_0_0_0_-10x0_0_0_15_0_-15_15x15_-15_0_0_0_0_-30x37.0_75.5_112.3_155.7_19 |
| 10 | 03_CCK1R:-10_0_0_0_0_0_-10x15_0_0_15_0_15_0x0_0_0_0_30_30_-30x37.0_75.5_112.3_155.7_191. |
| 11 | 10_CCK1R:-10_0_0_0_0_0_0x0_0_0_15_0_0_15x0_0_0_0_30_0_0x37.0_75.5_112.3_155.7_191.8_240. |
| 14 | 61_CCK1R:-10_0_0_0_10_0_-10x15_0_0_30_-15_-15_-15x0_0_0_0_15_0_15x37.0_75.5_112.3_155.7_ |

Conserved hydrogen bond networks among TM in Class A GPCRs (Venkatakrishnan 2013) are also observed in GPR139. We observed TM1-2- 7 works among N1.50, D2.50 and N7, 49 in Fig. 1. Instead of TM2-3-4 network at W450 in class A GPCR, GPR139 has TM2-4 network between N2.41 and R4.43 in Fig. 1. Since there is no conservation at E6.30 (T6.30 in GPR139) and all templates have active configurations, there is no salt-bridge constraint between R3.50 and E6.30. Consistently with traditional observations of class A GPCR activations, active ensemble structures of GPR139 also showed separation of TM 3-6 in the cytoplasmic side for all top 25 structures. In putative binding sites, hot spot residues of D2.61, E3.29, E3.32, R6.51, and N7.39 were directing toward the inside of TM bundles in Fig. 1.

We added all loops and helix-8 based on the MD structure of 5-HT2A receptor in Fig. 1. After homology modeling, we ran 10 cycles of simulated annealing 600K with fixed backbone and side chains of TMs and helix 8 to relax loops, followed by minimization with terminal gradient of 0.05 using Dreiding FF. Unlike class A GPCRs, there is no disulfide constraint between TM3 (C3.25) and extracellular loop 2 (EC2). Instead, there is a H-bond between E3.29 and T177 in EC2 in Fig. 1.

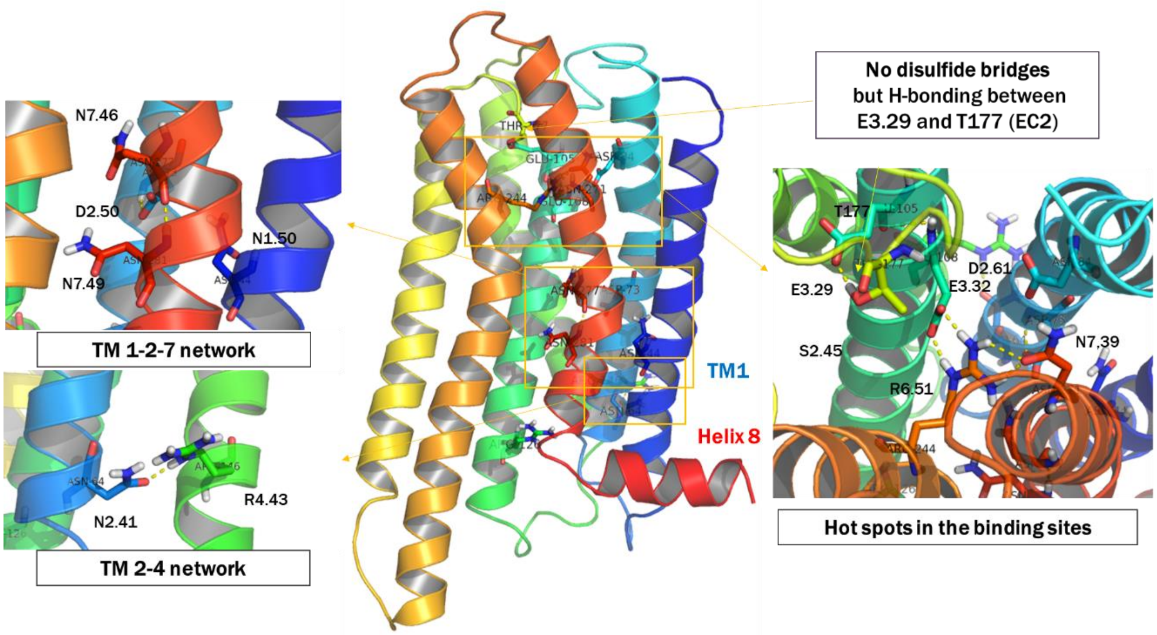

**Figure 1.** Predicted structure of GPR139 using the GEnSeMBLE complete sampling method

#### 2.2 Predicted binding site of ligand at GPR139

We predicted binding site of ligand 7c at the predicted GPR139 structure using DarwinDock complete sampling method, as shown in Fig. 2. Table 4 shows the rank of unified cavity energy (UCav) and snap binding energy (BE) from docking study. For final submissions to GPCR Dock 2021, we selected top 5 structures in Fig. 2, based on energy (rank1-4: u1-4) and H-bonding numbers (rank5: u11).

**Table 4**. The rank of unified cavity energy (UCav) and snap binding energy (BE) of ligand 7c binding poses at GPR139.

| filename | HBs | ucav | snapbe | snapbe_solv |
|---|---|---|---|---|
| GPR139.01HT2AR-metaD.CNTi12.ala.r38r39.7c.no_1.c11129.ft.c.s.a.bgf | H187,R244 | -40.93 | -55.49 | -40.32 |
| GPR139.25CCK1R.Ci25.ala.r38r39.7c.no_7.c21340.t.c.s.a.bgf | H187,R244 | -40.54 | -54.53 | -30.51 |
| GPR139.25CCK1R.Ci25.ala.r38r39.7c.no_5.c7172.ft.c.s.a.bgf | E108,R244 | -38.81 | -63.88 | -12.92 |
| GPR139.61CCK1R.Ci07.ala.r33r34.7c.no_7.c32297.ft.c.s.a.bgf | H187,R244 | -36.71 | -56.31 | -38.25 |
| GPR139.61CCK1R.Ci07.ala.r33r34.7c.no_5.c6710.ft.c.s.a.bgf | R244 | -34.00 | -45.14 | 13.35 |
| GPR139.61CCK1R.Ni06.ala.r41r42.7c.no_11.c36239.th.c.s.a.bgf | - | -29.76 | -36.19 | 18.50 |
| GPR139.01HT2AR-metaD.CNTi08.ala.r39r40.7c.no_10.c6875.h.c.s.a.bgf | E108,R244 | -29.20 | -35.81 | 43.90 |
| GPR139.01HT2AR-metaD.CNTi05.ala.r31r37r38.7c.no_4.c13127.f.c.s.a.bgf | R244,N271 | -25.95 | -31.84 | 26.01 |
| GPR139.61CCK1R.CNi14.ala.r43r44.7c.no_7.c4404.t.c.s.a.bgf | H187 | -25.64 | -51.31 | -29.33 |
| GPR139.25CCK1R.Ci25.ala.r38r39.7c.no_11.c13915.t.c.s.a.bgf | - | 51.49 | -46.75 | 37.50 |
| GPR139.61CCK1R.Ni06.ala.r41r42.7c.no_11.c5358.th.c.s.a.bgf | E108,Y163,H187,R244 | 387.80 | -61.50 | -14.69 |

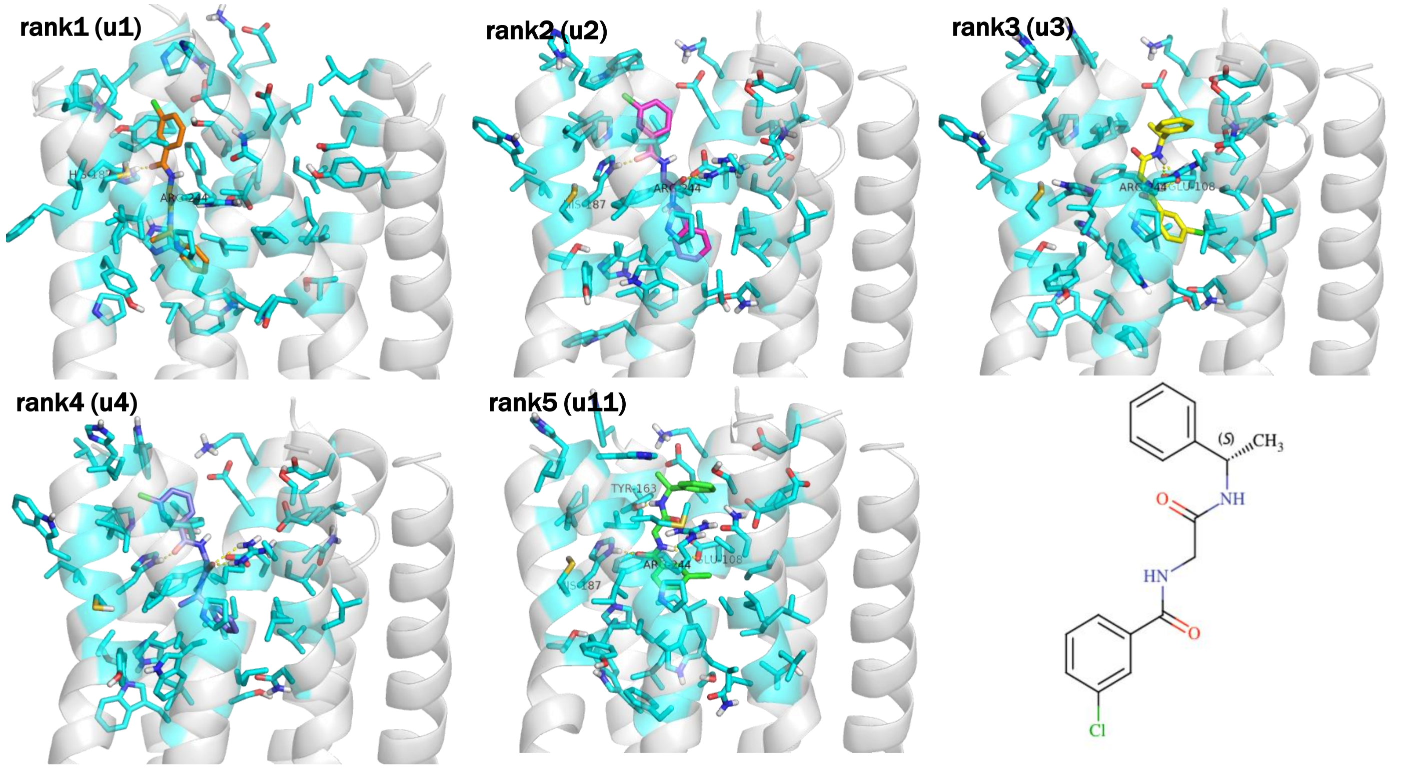

**Figure 2.** The top 5 structures of GPR139-7c complex structure using DarwinDock complete sampling method

**GPCRdock-3183 – OPRK**

#### Methods

To model the full activated κ-opioid receptor (κOR) coupled to Gi protein, we used the complex that was optimized with long scale molecular dynamics simulations described elsewhere[1]. To model dynorphin conformation, we used AlphaFold Protein Structure Database[2], in which by providing the protein sequence, it gives 10 best scored predicated conformation using deep learning methods.

To build κOR-bound dynorphin, we used the cryo-EM structure of μOR-DAMGO (a peptide ligand)-Gi protein as a template to separately superimpose dynorphin to DAMGO and κOR to μOR, respectively using the Needleman-Wunsch alignment algorithm[3] with the BLOSUM-62 matrix, which is incorporated in UCSF CHIMERA[4].

Among 10 different conformations of dynorphin, we picked the conformations that 1) there were no bad clashes between the heavy atoms of dynorphin and the backbone atoms of κOR 2) the N-terminus of dynorphin can form a salt bridge with D138$^{3.32}$. This results in only one conformation for later optimization.

To locate and optimize the interactions between dynorphin and κOR, we performed 20ns (400 cycles) of simulated annealing, in which the system was first heated from 25 to 600 K over 20ps with a sequence of 25, 100, 310, 450, 600 K and then sharply cooled back to 310 K over 30 ps. In this calculation, we placed harmonic restraints on backbone atoms with force constant of ~96 kcal.mol$^{-1}$Å$^{-2}$, while the side chains were free to find the optimum conformation. we ranked this structure as #1 since it is more consistent with docking procedure,

Subsequently, we embedded this pre-minimized κOR-dynorphin-Gi complex into a membrane bilayer composed of 333 POPC molecules in a simulation box of 113·113·149Å$^3$. We solvated this simulation box with water and ions to neutralize the system and added 0.15M NaCl, leading to ~170,000 atoms in the calculations. We then equilibrated the complex by 50ns molecular dynamics simulations, the procedure we described previously for κOR-Gi complex. The resulting complex was ranked #2. During these calculations, the proteins, peptide, POPC, and ions were described using the Charmm36m[5] parameter set. Water was described using the TIP3P model.

**GPCRdock-3183 – NPY1R**

**Computational details**

The neuropeptide Y Y1 receptor (NPY1R)-Gi protein-NPY complex structure was constructed starting from the inactive NPY1R structure (PDB code: 5ZBQ) with NPY from its NMR structure (PDB code: 1RON). Using the CHARMM-GUI (*1*, *2*), the inactive NPY1R was immersed into a 1-palmitoyl-2-oleoyl-sn-glycero-3-phosphocholine (POPC) lipid bilayer, where a total of 155 POPC molecules. The proteins and the lipid membrane were placed in a ~80 × 80 × 100 $Å^3$ box, with water molecules and 150 mM concentration of ions (sodium and chloride). The system was first relaxed by steepest-descent energy minimization. We then equilibrated the system by performing NVT (constant particles, constant volume, and 310K temperature) for 200 ps followed by NPT (constant number of particles, 1 bar pressure, and 310K temperature) for 10 ns of MD simulation, where positional restraints were placed on the heavy atoms with an initial force constant of 9.6 $kcal{\cdot}mol^{-1}{\cdot}Å^{-2}$ for proteins and 2.4 $kcal{\cdot}mol^{-1}{\cdot}Å^{-2}$ for lipids. These force constants were gradually reduced during the NPT simulation. Subsequently, further 20 ns metadynamics simulation was performed to induce the outward movement of TM6, because the intracellular end of TM6 shows a significant outward movement to create a cavity on the cytoplasmic region of the receptor that accommodates insertion of the Gα5 helix more deeply into the inside of the receptor upon activation.

Then, it was complexed with Gi protein from cholecystokinin A receptor (CCKAR)-Gi protein complex (PDB code: 7EZH). We then immersed the constructed proteins into a lipid bilayer with 271 POPC. The proteins and the lipid membrane were placed in a ~100 × 100 × 140 $Å^3$ box, with water molecules and 150 mM concentration of ions (sodium and chloride). We performed same minimization and equilibration steps as for the inactive NPY1R system mentioned above. After equilibration, we carried out 10 ns MD simulation.

All simulations were carried out using GROMACS (*3*) with PLUMED (*4*). we used the CHARMM36m force field (*5*), and the water molecules were described using the TIP3P model (*6*). The temperature was maintained at 310 K while the pressure was controlled at 1 bar. The time step was 2 fs.

#### GPCRdock-3580 (Zou)

#### Building models of protein-small molecule/peptide complexes for GPCR Dock 2021

Xianjin Xu, Rui Duan, Xiaoqin Zou*

Dalton Cardiovascular Research Center, Department of Physics and Astronomy, Department of Biochemistry, Institute for Data Science and Informatics, University of Missouri, Columbia, Missouri 65211, United States

##### A: Building the initial models of receptors

T01: The protein receptor (APJ) structure was extracted from PDB 5vbl, chain A.
T02: The protein receptor (GPR139) structure was built by AlphaFold and was downloaded from UniProt database (Access Number: Q6DWJ6).
T03: The protein receptor (OPRK) structure was built based on a homology structure PDB 6dde chain R by using the program MODELLER.
T04: The protein receptor (NPY1R) structure was built by AlphaFold and was downloaded from UniProt database (Access Number: P25929).
T05: The protein receptor (NMUR2) structure was built by AlphaFold and was downloaded from UniProt database (Access Number: Q9GZQ4).

##### B1: Small molecule docking

A recently-developed template-guiding method [1] was used for protein-ligand complex structure prediction. Briefly, for a query target, protein crystal structures (or protein structures from homology modeling) containing co-bound ligands were downloaded from the Protein Data Bank. These protein-ligand complex structures were superimposed onto the target receptor structure (see Section A), and the binding poses of co-bound ligands were used as templates as described later. For a query ligand, up to 200 conformers were generated from the SMILES string by using the OMEGA2 program (Version 2.5.1.4, OpenEye Scientific Software, Santa Fe, NM, USA. http://www.eyesopen.com) [2,3]. Then, the ligand conformers were superimposed onto the template ligands using a 3D molecular similarity measurement program, SHAFTS [4]. The best superimposed mode was used as the initial structure for further local refinement, which was performed by AutoDock Vina [5]. Finally, the generated protein-ligand complex models were ranked using a hybrid scoring function that combines both the binding energy score and the 3D similarity score. The details of the method are presented in Reference 1.

For T01 (APJ/cpd6), the template structures of the top 5 submitted models were 4zud, 6me8, 6gpx, 4yay, and 6akx, respectively.

For T02 (GPR139/JNJ), the template structures of the top 5 submitted models were 6b73, 6tot, 6ko5, 6tp4, and 4ea3, respectively.

##### B2: Peptide ligand docking

Our recently-developed method, MDockPeP2, [6] was used for the protein-peptide complex structure predictions. Briefly, the strategy of MDockPeP2 for structure prediction was divided into three stages. In Stage 1, for a query peptide, a protein structure database was searched to find fragments sharing a similar sequence. These fragments were used as templates to build initial peptide conformers. In Stage 2, peptide conformers were independently docked to a target protein surface using Fast Fourier Transform (FFT)-

based protein−protein docking. The generated models were further refined using protein-small molecule flexible docking (local flexible minimization). In Stage 3, peptide conformers were merged and evaluated using hybrid scoring which combines both energy binding scores and interface similarities. The details of the method are available in Reference 6.

**C: Model refinement**

The models for submission were optimized with a simple force-field minimization of the energy in Maestro Version 12.9.137 (Schrödinger, LLC) [7].

**D: Criteria for selecting 5 best models per target**

Predicted binding modes were ranked by their scores which were calculated by scoring functions implemented in the docking programs. The top 30 binding modes for each target were manually examined and 5 models were selected for submission.

**E: Target challenges**

Large conformational changes were observed in the ligand binding regions of the GPCRs. Protein flexibility was not fully considered in our current prediction strategies.

**F: Comments on experimental information and human expertise**

Experimental information is important to the binding mode predictions. In our protein-small molecule prediction method, co-bound ligands in the experimentally determined complex structures were used as templates to model the binding mode of a query ligand.

Human expertise is also helpful for the binding mode selection. In our prediction process, top 30 binding modes were kept for manual inspection for each target, and 5 models were selected for submission.

[7] Maestro, Schrödinger, LLC, New York, NY, 2021.

### GPCRdock-3601

### Multi-State modeling of G-protein coupled receptors with AlphaFold and followed modeling of bound ligands

Lim Heo[1] and Michael Feig[1, *]

[1] Department of Biochemistry and Molecular Biology, Michigan State University, East Lansing, MI 48824, USA

GPCR-ligand complex structures were modeled with a two-stage modeling protocol. First, receptor structures were predicted using a multi-state GPCR modeling protocol with AlphaFold2.[1, 2] Secondly, bound ligands were modeled using AutoDock Vina[3] for organic molecules (T01 and T02) or using AlphaFold-Multimer[4] for peptides (T03, T04, and T05).

##### *Receptor structure modeling*

Receptor structures were predicted using a modified protocol of AlphaFold that aims for modeling multiple conformational states of the GPCR.[2] We altered two input features from the original AlphaFold protocol[1] to enable guided modeling of the active and inactive states of the GPCR. First, homologous template structures used as initial input were selected among activation state-annotated GPCR databases instead of the HH-suite pdb70 database.[5] For example, by selecting templates from GPCR structures in the active state, structure prediction was guided towards model structures in the active state. Second, input features based on multiple sequence alignment (MSA) were modified to focus the AlphaFold neural network model more on input features from template structures. Since AlphaFold neural network model generally relies more on the MSA-based input features when there are enough homologous sequences, aligned residues in MSAs were masked by gaps at sequence positions where template structures were available.

The activation states of receptors of the GPCR Dock 2021 targets were inferred based on provided information. We considered a target receptor in the active state if it forms a complex with G-proteins. Otherwise, we modeled a target receptor in the inactive state As a consequence, target T01 was modeled in the inactive state, while targets T02–5 were modeled in the active state.

Scripts for the multi-state modeling protocol is available at https://github.com/huhlim/alphafold-multistate. The activation-state annotated GPCR databases are deposited at https://zenodo.org/record/5745217. A Google Colab notebook for the modeling protocol is available at https://colab.research.google.com/github/huhlim/alphafold-multistate/blob/main/AlphaFold_multistate.ipynb.

##### *Protein-ligand docking for organic molecules (T01 and T02)*

Organic molecules were docked to the predicted receptor structures using AutoDock Vina.[3] Three-dimensional ligand conformations were generated from SMILES strings using UCSF Chimera.[6] Molecular dynamics (MD) simulations were carried out to diversify ring conformations, bond lengths, and bond angles of the three-dimensional structures; these degrees of freedom are not sampled during protein-ligand docking using AutoDock Vina. For MD simulations, ligands were parametrized using CGenFF.[7, 8] For each ligand, one ligand molecule was solvated in a cubic TIP3P water box with a width of 40 Å. Using Langevin dynamics simulations, the solvated system was gradually heated up to 298.15 K and equilibrated at 1 bar for 1 ns. Temperature and pressures were maintained using Langevin thermostat and Monte Carlo barostat. Alternative conformations were sampled by ten replicas of 500 ns-long Langevin dynamics simulations. The sampling resulted in 50,000 conformations and clustered to four conformations using ligand heavy-atom RMSD values as the distance metric. Five ligand structures - one conformation generated by UCSF Chimera and four conformations from the MD simulations - were docked to their corresponding receptor structures using AutoDock Vina at the canonical binding site of class A GPCRs. The receptor structure was kept rigid or one of the binding site residues was set to be flexible. Docked ligand poses were clustered using ligand heavy-atom RMSD as the distance metric with a cutoff radius of 2 Å.

Five submitted models were manually selected by considering various features that are related to binding affinity. We first considered the docking score from AutoDock Vina and the cluster size to narrow down the candidates to about 10 models. Those models were further assessed by applying MD simulations. Each complex structure was solvated in a POPC lipid bilayer and a TIP3P water box using CHARMM-GUI.[9] We used the CHARMM 36m force field[10] and CHARMM 36 lipid force field[11] for modeling the protein and the POPC molecules, respectively. For ligands, we re-used the CGenFF parameters. Two replicas of Langevin dynamics simulations were performed at 298.15 K and 1 bar for 50 ns. We considered a complex structure unstable if the ligand moved significantly throughout the simulations. We also compared binding free energies using MMGB/SA analysis with HDGBvdw.[12] By considering the aforementioned criteria, we selected five models to be submitted.

***Protein-peptide docking for peptides (T03, T04, and T05)***

The complex structures of a target receptor and a peptide were modeled using a modified AlphaFold-Multimer protocol. The protocol was based on the original AlphaFold-Multimer protocol[4] but modified to take any structure as a template structure rather than searching against PDB. This modification allowed us to provide a receptor structure modeled in the active state. For target T03, when the protocol was applied, there was no paired sequence in MSA because the peptide sequence is too short. Considering that coevolutionary information is the most important factor of successful prediction by AlphaFold, the predicted model may be not accurate.[1] As an alternative, fragments of the peptide were docked to the receptor using AutoDock Vina. We obtained a converged docking result for the first five residues of the peptide. By using this binding pose as an additional input template of the modeling protocol, we obtained alternative models for the receptor-peptide complex, however, they were comparable to the models that were predicted without the docked pose. We selected five maximally diverse complex models for submission. For targets T04 and T05, there were enough paired sequences in MSA when we applied the modified AlphaFold-Multimer protocol. For both targets, five predicted models had high confidence scores, and therefore we submitted those models.

### GPCRdock-3983 and GPCRdock-8470

#### Methods for Huang group

Huanyu Tao, Keqiong Zhang, Peicong Lin, Qilong Wu, Xuejun Zhao, Sheng-You Huang*

*

*School of Physics, Huazhong University of Science and Technology, Wuhan, Hubei 430074, P. R. China*

Among the five targets in GPCRDOCK2021, T01 and T02 are the complexes between proteins and small molecules. For the two targets, we first performed a sequence search to find their templates in the protein data bank (PDB)[1]. For target T02, we also searched using the nanobody sequence because we noticed that class A GPCRs bound with Nanobody share some structural similarities. Then we ranked all the templates by their sequence similarities and selected the higher-resolution structures among the top-ranked results. The structures of the targets were built by homology modeling with Modeller[2]. For small-molecule ligands, we adopted Conformator[3] to generate 100 conformations for each ligand, and then used MDock[4] to dock the ligand conformations against the GPCR target. We visually inspected the top 200 docking poses for each ligand based on available biological information from the literature[57]. After that, 300ps molecular dynamics (MD) simulations were performed to refine each docked protein-ligand complex. Finally, we conducted the second round of visual inspection and selected 5 models as the final results of Group 3983.

Compared with Group 3983, Group 8470 skips the molecular docking step. Instead, we directly superimposed the ligand with the cocrystallized molecule of the template and adjusted the side-chain orientation in the pocket. The subsequent operations were similar to that for Group 3983. Namely, 300ps MD simulations were performed and five optimal models were finally selected.

For three peptide-protein targets (T03-T05), AlphaFold-Multimer[8] was first used to build the initial models of the complexes for OPRK, NPY1R, and NMUR2. All the default parameters were used during the modeling with AlphaFold-Multimer. The top-ranked model predicted by AlphaFold-Multimer was used for docking for each target. MDock[4] was then used to dock an ensemble of peptide conformations generated by MODPEP[9] against the ligand binding site on the modeled GPCR protein target. The peptide structures modeled by AlphaFold-Multimer were also docked against the protein. Finally, the top 100 binding modes[10] were manually inspected based on the knowledge from the literature, and the top 5 protein-peptide models were selected after MD refinement.

It is noted that for NPY1R, the experimentally derived restraints from ref [11] were used to select the binding modes. For NMU25, the conserved residues in the C-terminal region of the peptide are considered to be the key interacting residues for the peptide binding[12]. Unfortunately, three of the five experimentally derived contact pairs (Y36-Y100$^{2.64}$, R33-N299$^{7.32}$, L30-I293$^{ECL3}$) in ref [11] do not appear in the native structure of NPY1R. As such, the experimental information would have misled the selection of our binding modes, resulting in a wrong prediction of binding modes on the target.

#### GPCRdock-4190

#### Methods

##### Homology modeling

Swissmodel server was applied for the homology modeling for each target(1). In Table 1, we listed the targets with the PDB ID for the corresponding model. The choice of modeling considered the sequence similarity, the diversity of secondary structures, and the matching degree of downstream protein (Gi/Gs/None).

**Table 1.** Structure template for homology modeling

| *Target* | *Template for model 1* | *Template for model 2* | *Template for model 3* | *Template for model 4* | *Template for model 5* |
|---|---|---|---|---|---|
| APJ | 5VBL | 5VBL | 6KNM | 6KNM | 7FIT |
| GPR139 | 6UP7 | 6UP7 | 7L0P | 7L0P | 7L0P |
| KOR | 6B73 | 6B73 | 6B73 | 6B73 | 6B73 |
| NPY1R | 7MBX | 7MBX | 7MBX | 7EZH | 7EZH |
| NMUR2 | 6OS9 | 6OS9 | 6OS9 | 6OS9 | 7EZH |

##### Small molecule docking

Induced fit docking (IFD) was applied to small-molecule ligands of APJ and GPR139. Firstly, the ligands were prepared by Maestro, Schrödinger. Then, we set the classical orthosteric pockets of class A GPCRs as the pocket of APJ and GPR139 and docked ligands into it via the Induced Fit module of Schrödinger suite. Extended sampling was applied. After docking, the cluster module of CPPTRAJ from Amber was used to pick distinct binding poses of ligands for each template(2).

##### Peptide docking

Protein-protein docking was applied to peptide ligands of KOR, NMUR2, and NPY1R. Conformations of ligands were firstly generated by Schrödinger and MOE, then the protein-protein dock module of Maestro, Schrödinger was applied to generate the poses. Meanwhile, the HPEPDOCK server was also used to produce the poses using ligand sequence and structure models(3). The classical orthosteric site for class A GPCRs was set as the peptide site. The specific method for each model was listed in Table 2.

**Table 2.** Methods for the generation of binding pose.

| *Target* | *Method for model 1* | *Method for model 2* | *Method for model 3* | *Method for model 4* | *Method for model 5* |
|---|---|---|---|---|---|
| APJ | IFD | IFD | IFD | IFD | IFD |
| GPR139 | IFD | IFD | IFD | IFD | IFD |
| KOR | Maestro | Maestro | HPEPDOCK | HPEPDOCK | HPEPDOCK |
| NPY1R | Maestro | HPEPDOCK | HPEPDOCK | HPEPDOCK | HPEPDOCK |
| NMUR2 | Maestro | Maestro | Maestro | HPEPDOCK | Maestro |

### GPCRdock-4528 and GPCRdock-8669

#### Predicting GPCR-ligand and GPCR-peptide complex structures using GALAXY

The initial GPCR structures were obtained from PDB (PDB ID 5VBL and 6KNM for T01 and 6B73 for T03) or modelled by using AlphaFold2[1] and 'activation state annotated GP CR database'[2]. AlphaFold2 was run in house for various combinations ranging from single G PCR chain to full complex involving all protein chains used for experimental structure determ ination. The initial GPCR structures were further refined by sidechain rebuilding and loops m odelling with GalaxyLoop[3].

For targets with small organic ligands, protein-ligand docking was performed using ri gid-body docking methods GalaxyDock2[4] and GalaxyDock3[5], a flexible-receptor docking me thod Galaxy7TM[6], and a template-aided docking method GalaxyCSAlign [paper in preparatio n]. For peptide docking, the peptide terminus predicted to interact in the receptor core was mo deled first by docking with GalaxyDock2, refining AlphaFold2 models, or template-based mo deling with GalaxyTBM[7] (PDB ID 6DDF was used as a template for T03). The remaining par t of the peptide was built by rigid-body docking with GalaxyTongDock[8] and terminal and loo p modeling with GalaxyLoop[3]. The complex structures were refined through simple relaxatio n or by using GalaxyRefineComplex[9] with GPCR-specific Galaxy7TM score.

Final models were selected by human expert guided by scoring functions and experim ental information. Human intuition and experimental information were also used to bias struct ure sampling. For T02, receptor structure sampling was enriched by perturbing 222ARG whic h was predicted to interact with the ligand. Our Model 5 ranked 5th among all models submitt ed by all groups. While the interaction with 222ARG was predicted successfully, ECL2 loop c onformation was not for this target. For T03, helical peptide terminus was kept using experim ental information. Our Model 3 for this target was ranked 1st among all models from all partic ipants.

[We certify that Hyeonuk Woo and Chaok Seok, who are co-authors of a paper related to T04 from the same university[10] were not involved in modeling T04]

The sequence of APJ was used in AlphaFold Colab [1] with relaxation stage. Obtained structure was used in Rosetta to get all possible conformations and their corresponding total energies. [2] 100 models (i.e., conformers) were constructed with Rosetta and their corresponding energies are calculated. Total energy for each model was calculated, and based on total energy scores, lowest energy model is selected. This model was used in molecular dynamics (MD) simulations. Target protein structure was prepared using with the Protein Preparation module of the Maestro molecular modeling suite before the docking and MD simulations. Hydrogen atoms were added, bond orders are assigned and the protonation states are generated at physiological pH (7.4). Finally, a restrained minimization was applied to the optimized structure with OPLS3e forcefield. [3] A membrane bilayer medium is used for simulations. Prepared protein structure was embedded into 1-palmitoyl-2-oleoyl-sn-glycero-3-phosphocholine (POPC) lipid bilayer, the coordinates of the transmembrane domain oriented in the membrane bilayer are obtained from OPM database. [4] The system was solvated with the TIP3P water model with 15 Å thickness, and 0.15 M of NaCl ions are added to neutralize the systems. Model systems have ~200.000 atoms and were relaxed before simulations. 100 ns of production simulations with 2 fs time steps were run for the prepared system, with NPT ensemble in physiological conditions of 310 K and 1 bar. 2000 trajectory frames were collected for the system throughout the simulations. A clustering analysis was the conducted and representative structure was used in docking studied. Provided .smi code of ligand structure was converted to 3D structure by the LigPrep module of the Maestro molecular modeling suite [5]. OPLS3e force field with neutral pH is used in the ligand preparation. In order to use more accurate partial charges of the compound, a quantum mechanics (QM) optimization (B3LYP method with 6-31G** basis set) was conducted with Jaguar for the ligand and its partial charges were used in the docking. [6] In docking, Glide/SP module was used with expanding sampling options. [7] Top-docking poses were then used in MD simulations and 100 ns MD simulations were performed with Desmond. Based on clustering of the trajectories, 5 poses were selected.

**GPCRdock-5072 - APJ**

**Receptor model**

A homology model for APJ was generated in Molsoft ICM-Pro v3.9-2c using the Full Model Builder option with the PDB structure 5VBL as template. The Zero End-gap Global Alignment (ZEGA) alignment (https://doi.org/10.1006/jmbi.1997.1287) of our query sequence and 5VBL yielded 99% sequence identity and probability of log 75.3. The modelling was then submitted using full refinement, insert margin of 2, deletion margin of 3 and a MC effor to 1.00. Finally, the N and C-terminus not visible in the final structure were removed from homology model prior to docking.

**Docking (model 04)**

The receptor model was prepared in Molsoft ICM-Pro v3.9-2c by adding and optimizing the positions of hydrogen atoms, as well as the orientation and protonation states of histidine and cysteine residues and the orientation of glutamine and asparagine residues.

For the first trial of docking, the binding pocket was defined around the residues of the co-cristallized peptide buried (5VBL). The molecule to dock was manually created and charged.

The docking was performed with a docking effort of 10 and no initial constraints. From the 50 saved conformers, no one was selected because of very high scores.

Then Oliver proposed an interesting pose obtained with Autodock that I loaded and Minimized using ICM 'Minimize Ligand is soft grid' a few times.

I also obtained some pose with 'redocking ligand in current receptor' and selected the one with the best scores, manually validate them by visualization and again 'Minimize Ligand is soft grid'.

**Docking (model 03)**

The receptor model (homology model with terminals) was prepared in Molsoft ICM-Pro v3.9-2c by adding and optimizing the positions of hydrogen atoms, as well as the orientation and protonation states of histidine and cysteine residues and the orientation of glutamine and asparagine residues.

The binding pocket was defined important residues identified by Ma et al. (https://doi.org/10.1016/j.str.2017.04.008): W24, Y35, W85, Y88, Y93, R168, L173, Y182, Y185, Y264, K268, Y271, F291 and Y299; and neighboring residues within 3 (Å? ICM unit?).

The docking was performed with docking effort of 10 and no initial constrains with the negatively charged ligand. From the ten saved conformers, pose 1 and 8 (DF1_SL1, DF1_SL8) were selected after visual inspection considering the available experimental data.

**Docking (model 02 and model 01)**

The crystal structure of the apelin receptor (PDBID: 5VBL) was prepared with Free Maestro 2020-1 by adding, optimizing the orientation and protonation states of histidine and cysteine residues and the orientation of glutamine and asparagine residues. The ligand was sketched in Free Maestro, making sure to set the negative charge on its tetrazole. PDBQT files were then generated using AutoDockTools 1.5.7 both for the receptor and the ligand. The docking grid was centered on x=187.611, y=7.615 and z=54.112 and spacing of 0.254 Å. The number of points in each direction was set to the maximum available value of 126. Default parameters of Lamarckian genetic algorithm were used, random seed was set to 1234 and docking was performed with AutoDock 4.2.6. From the ten saved conformers, pose 1, with docking score of –9.88, was selected after visual inspection considering the available experimental data.

The model was subsequently used in a 50ns MD simulation of the system in a membrane embedded environment using Desmond. To this end the membrane was automatically placed to helices, the water model set to SPC with forcefield OPLS 2005, the box type to orthorhombic with padding distance of 10 Å in each direction. The trajectory was recorded every 100 ps in an NPγT ensemble at 300 K and 1.01325 bar and default parameters.

**Minimization (model 05)**

For the best pose (from Sohvi, Olivier and David), the side chains of the protein were minimized and the ligand relaxed in ICM-Pro. First, local torsions were optimized during 100 steps and 300 steps ligand relaxation followed by global energy optimization including 200 global moves and 100 additional local minimization calls and another 300 steps ligand relaxation.

**GPCRdock-5072 – GPR139**

**Receptor model**

The model for GPR139 in the active state was generated with the GPCR multi-state implementation of AlphaFold by Heo et al. (https://www.biorxiv.org/content/10.1101/2021.11.26.470086v1). In the model, the conformation of TM6 did not fully align with that of an active GPCR state, therefore we created a homology model for GPR139 with MolSoft ICM-Pro v3.9-2c with the active conformation of adenosine receptor A2A (PDB 5G53) as template in the absence of a template with a high sequence similarity.

The top predicted active AlphaFold model was integrated with the generated active homology model in the following fashion in Schrodinger Maestro (academic version 12.3): residues* 417-469 from the homology model (corresponding to the intracellular halves of TM5 and TM6) were replaced in the AlphaFold model upon superimposition of both models. Peptide backbone bonds were manually created between residues C416-F417 and W469-A470. C416 and F417 were subsequently minimized to reduced strains in the backbone. The resulting model was minimized in UCSF Chimera with 100 steps of steepest descent followed by 10 steps of gradient conjugate with restraints on Cα carbon atoms. Finally, hydrogen atoms were added. The minimized model was exported for docking as model A. A second model (model B) was exported after nine additional rounds of minimization in UCSF Chimera.

**Docking**

The receptor models A and B were prepared in Molsoft ICM-Pro v3.9-2c by adding and optimizing the positions of hydrogen atoms, as well as the orientation and protonation states of histidine and cysteine residues and the orientation of glutamine and asparagine residues.

The binding pocket was defined around the residues of interest identified by mutagenesis experiments by Nøhr et al. (https://doi.org/10.1038/s41598-017-01049-z) and Wang et al. (https://doi.org/10.1002/prp2.466). These residues included F109, H187, W241, N271; and Y33, V76, V191, and L249, respectively from both publications.

The small molecule of interest was prepared by adding hydrogen atoms and assigning atomic charges and subsequently docked independently in both receptor models using a docking effort of 10 and no constraints. The docking results were analyzed considering the available experimental data and the interaction networks generated, leading to the selection of the top 1 and top 5 poses in terms of docking score from model A and top 1 pose from model B.

The selected docked poses in model A were transferred to the receptor model B, where the side chains of the receptor were further minimized in the defined binding pocket in ICM-Pro without restraints. Minimization was performed in 200 steps of global minimization followed by 100 steps of local minimization.

| |
|---|
| * Residue numbers from reference Uniprot sequence Q6DWJ6 |

**GPCRdock-5072 - OPRK**

**Receptor model**

A homology model for KOR was generated in Molsoft ICM-Pro v3.9-2c using the Full Model Builder option. The primary structure of the chains B and A from the PDB structure 6B73 was extracted based on the residues shown in the crystal structure. We then selected chain A as template structure for our model due to higher electron density coverage of the amino acid sequence. The Zero End-gap Global Alignment (ZEGA) alignment (https://doi.org/10.1006/jmbi.1997.1287) of our query sequence and the chain A of 6B73 yielded 98% sequence identity and probability of log 57.8. The sequence was then submitted for homology modelling using Full Refinement, insert margin of 2, deletion margin of 3 and a MC effort of 1.00.

**Docking GPCR-DOCK-2021-T03_GPCRdock-5072_rank01**

The receptor model was prepared in Molsoft ICM-Pro v3.9-2c by adding and optimizing the positions of hydrogen atoms, as well as the orientation and protonation states of histidine and cysteine residues and the orientation of glutamine and asparagine residues.

The binding pocket was defined around the residues of interest identified by mutagenesis experiments and summarized by Ferré et al. (https://doi.org/10.1016/bs.vh.2019.05.006), and by residues of interest (my MD/docking) by O'Connor et al. (https://doi.org/10.1073/pnas.1510117112): Q115, Y119, D138, N141, M142, E209, D223, M226, V230, H291, I294, Y313, I316, Y320 and N322; and neighboring residues within 3 (Å? ICM?).

Peptide docking was performed with docking effort of 10 and no initial constrains, starting with AA sequence. From the ten saved conformers, pose 5 (DF2_SL5) were selected after visual inspection considering the available experimental data. The docked structure (peptide and protein side chain atoms) where optimized with Chimera (DF2_SL5min).

**Docking GPCR-DOCK-2021-T03_GPCRdock-5072_rank02-04**

The receptor model was prepared in Molsoft ICM-Pro v3.9-2c by adding and optimizing the positions of hydrogen atoms, as well as the orientation and protonation states of histidine and cysteine residues and the orientation of glutamine and asparagine residues.

The binding pocket was defined around the residues of interest identified by mutagenesis experiments and summarized by Ferré et al. (https://doi.org/10.1016/bs.vh.2019.05.006), these are Q115, Y119, D138, H291, I294, Y313, I316, and Y320. The peptide of interest was obtained from the Protein Data Banck (PDB 2N2F) and subsequently prepared in the same fashion as the receptor model.

Peptide docking was performed with a docking effort of 10 and no initial constraints. From the ten saved conformers, pose 5 was selected after visual inspection in light of the available experimental data. This pose was used as a base for re-docking, where the N-terminus was positionally constraint to the coordinates in pose 5. Two additional poses were generated in this step, which were also analyzed considering the interaction networks displayed and the experimental data available.

For the best pose, the side chains and hydrogens of the peptide and the binding pocket (residues at 5A of the peptide) were minimized in ICM-Pro. First, local torsions were optimized during 100 steps followed by global energy optimization including 200 global moves and 100 additional local minimization calls. The resulting model was selected for submission.

**GPCRdock-5072**

**Homology modelling**

The sequence of NMUR2 given by the organizers, omitting unresolved amino-acids, was submitted to SwissModel for homology modelling. The crystal structure of Neurotensin receptor type 1 (PDBID: 6OS9 chain A) with 32.75% sequence identity and co-crystalized G protein partner was used as template. The obtained model had QMEANDisCo score of 0.61 ± 0.05 and MolProbibity score of 1.64. In details 95.21% of phi-psi angles of its side-chains were in favoured and allowed Ramachandran regions; clashes between Phe89 and Phe182 and between Tyr171 and Ile175 were observed; two rotamer outliers, Phe89 and Leu231, were present; and a total of 36 bad angles were reported throughout the homology model.The homology model was then subjected to minimization with UCSF Chimera after AM1-BCC partial charges had been calculated. First 1000 steepest gradient descent and 100 conjugate gradient steps, both with step size 0.02 Å, were carried out with restraints on the protein backbone. Then five repeats of 100 steepest gradient descent and 100 conjugate gradient steps, both with step size 0.02 Å, were employed.

A similar homology model was constructed using the same sequence but forcing SwissModel to use of the active Cryo-EM structure of the human adenosine A1 receptor-Gi2-protein complex (PDBID: 7LD4). The obtained model had QMEANDisCo score of 0.61 ± 0.05 and MolProbibity score of 1.64.

The former homology model was then morphed into the latter using Chimera through 100 corkscrew interpolation steps with linear rate and 0.5 core fraction.Residues Gln268 to Val281 included from the morphed conformation were then grafted into the homology model obtained with neurotensin receptor 1 template. Stretched peptide bonds were created between Ile267 and Gln268 and between Val281 and Leu282 using Free Maestro. The obtained chimera was minimized using Chimera (100 steepest gradient descent and 100 conjugate gradient steps). Rotamers of residues of TM5 clashing with those of TM6 were selected using Free Maestro to reduce clashes (table below). Finally, the structure was minimized once again using Chimera (100 steepest gradient descent and 100 conjugate gradient steps).

| Residue | Rotamer | | | | |
|---|---|---|---|---|---|
| | Abundance | Chi1 | Chi2 | Chi3 | Chi4 |
| Lys256 | 7.4% | -62° | -66° | -177° | -178 |
| Glu259 | 15.6% | -68° | 178° | -2° | - |
| Gln268 | 30.8% | -37° | 98° | - | - |
| Phe280 | 3.0% | -60° | -60° | -30° | - |

**Docking**

The receptor model was prepared in Molsoft ICM-Pro v3.9-2c by adding and optimizing the positions of hydrogen atoms, as well as the orientation and protonation states of histidine and cysteine residues and the orientation of glutamine and asparagine residues. Peptides consisting of the last 4 and 8 residues, as well as the original peptide sequence were generated in ICM.

The binding pocket was searched for using the ICMPocketFinder. Residues surrounding the pocket were selected and a box surrounding the binding site was initialized. First, the last four residues (DVRF) of the peptide were docked as ligand using an ICM docking effort of 10. After visual inspection, the best pose was selected. Selection was based on entering of the C-terminus into the binding pocket and for the selected pose the C-terminus entered the binding pocket. Next, the last 8 residues (QFEEDVRF) of the peptide were docked, using the selected pose of the previous docking run as template with Atomic Property Fields (APF) matching. The docking effort used here was also 10. The results of these poses were also visually inspected, after which the best-energy ranked pose was chosen to further build the template. Finally, the whole peptide was docked using the best-energy ranked pose of the previous dock as template. The docking effort for this docking was set to 0.1 in order to speed up the process. The final docking step of the complete peptide was repeated multiple times in order to generate multiple poses.

**GPCRDock-5792 Methods Overview**
iMolecule (Petr Popov)

1. Building the initial models of receptors (if receptors were modeled separately from ligands) or complexes. Did you use AlphaFold, RoseTTAFold, or any other deep-learning-based methodology for building the initial models? (*if yes, please provide the info about the exact methodology and its implementation that you used*). What other methods did you use, and how?

To build starting models of the receptors first we retrieve the target conformation of the receptor (i.e. active or inactive) based on the given information about the target. Then we used both AlphaFold2.0 [1] and homology modelling (molsoft.com) to generate a set of 10 putative conformations. This was followed by selecting conformation with the target state by analysing pair-wise distances between the residues of the signalling activation pathway [2]. Top 5 starting models were used as starting conformations of 100ns full-atomic molecular dynamics simulations using Gromacs [3]. We analyzed the obtained molecular trajectories with BiteNet[4] for small molecules and BiteNet_p[5] for peptide molecules to identify conformations corresponding to the most druggable binding site. We clustered BiteNet's predictions and <u>selected representatives of Top 5 clusters as the final models for docking.</u>

The final submissions were a mixture of models built by homology (HM) and by AlphaFold2 (AF), as follows:

GPCR-DOCK-2021-T01_GPCRdock-5792-1.pdb - HM
GPCR-DOCK-2021-T01_GPCRdock-5792-2.pdb - HM
GPCR-DOCK-2021-T01_GPCRdock-5792-3.pdb - AF
GPCR-DOCK-2021-T01_GPCRdock-5792-4.pdb - HM
GPCR-DOCK-2021-T01_GPCRdock-5792-5.pdb - HM

GPCR-DOCK-2021-T02_GPCRdock-5792-1.pdb - HM
GPCR-DOCK-2021-T02_GPCRdock-5792-2.pdb - AF
GPCR-DOCK-2021-T02_GPCRdock-5792-3.pdb - HM
GPCR-DOCK-2021-T02_GPCRdock-5792-4.pdb - HM
GPCR-DOCK-2021-T02_GPCRdock-5792-5.pdb - AF

GPCR-DOCK-2021-T03_GPCRdock-5792-1.pdb - HM
GPCR-DOCK-2021-T03_GPCRdock-5792-2.pdb - HM
GPCR-DOCK-2021-T03_GPCRdock-5792-3.pdb - AF
GPCR-DOCK-2021-T03_GPCRdock-5792-4.pdb - HM

GPCR-DOCK-2021-T03_GPCRdock-5792-5.pdb - AF

GPCR-DOCK-2021-T04_GPCRdock-5792-1.pdb - AF
GPCR-DOCK-2021-T04_GPCRdock-5792-2.pdb - AF
GPCR-DOCK-2021-T04_GPCRdock-5792-3.pdb - HM
GPCR-DOCK-2021-T04_GPCRdock-5792-4.pdb - AF
GPCR-DOCK-2021-T04_GPCRdock-5792-5.pdb - HM

GPCR-DOCK-2021-T05_GPCRdock-5792-1.pdb - AF
GPCR-DOCK-2021-T05_GPCRdock-5792-2.pdb - HM
GPCR-DOCK-2021-T05_GPCRdock-5792-3.pdb - AF
GPCR-DOCK-2021-T05_GPCRdock-5792-4.pdb - HM
GPCR-DOCK-2021-T05_GPCRdock-5792-5.pdb - AF

2. Small molecule docking and/or peptide ligand docking (as applicable)

We used ICM-Pro to dock small molecules into the selected conformations with the default parameters, except for the higher value for the docking effort (30 instead of 1).

In case of peptide ligands, its starting orientations were generated using AlphaFold2.0.

Additionally, for the KOR target, we used NMR experimental data from [6].

3. Any model refinement (if applicable)

No refinement was used for small molecules, apart from the 100ns MD simulations on the initial structures described above.

In case of peptide ligands, the final orientation corresponded to the molecular dynamics' frame selected on the first step.

4. Criteria for selecting 5 best models per target

We sorted the obtained binding poses by the docking score and selected the top 5 models.

5. Please include a section about target challenges. What did you find most challenging about each complex, and what information and approaches did you use to solve the challenge?

One of the long-standing problems is how to choose right conformation for molecular docking, given that flexibility of the receptor is very limited in traditional methods. Our idea was to investigate, if binding site analysis can be helpful in selecting starting conformations for molecular docking from the molecular dynamics simulations. We therefore generated conformational ensembles with MD and filtered out conformations with low binding site score. However, there were still many conformations with high score, likely, because of the apparent cavity (orthosteric binding site) in GPCRs. Thus, from the one hand we generated the most promising starting conformations for docking, but from the other hand could not explore them all. It would be interesting to compare the true binding conformations with the generated conformational ensembles to see if i) the true conformations were captured, ii) the true conformations were missed in the clusterization step, or iii) the true conformations were not captured at all.

6. Please comment on the role of experimental information and human expertise in generation and selection of your models (if applicable).

N.A.

### GPCRdock-6153

#### Methods

*R. Guixà-González[1,2], Matthew Rodigues[2], Spencer Bliven[3], and X. Deupí[1,2]*

*[1]Condensed Matter Theory Group, Division Scientific Computing, Theory and Data, Paul Scherrer Institute, 5232 Villigen PSI, Switzerland.*

*[2]Laboratory of Biomolecular Research, Division of Biology and Chemistry, Paul Scherrer Institute (PSI), 5232 Villigen PSI, Switzerland.*

*[3]High Performance Computing and Emerging Technologies Group, Division Scientific Computing, Theory and Data, Paul Scherrer Institute, 5232 Villigen PSI, Switzerland.*

T01: APJ and T02: GPR139 (small molecule ligands)

The 3D structure of the G protein-coupled receptor (GPCR) was predicted by using the full version of the implementation of the inference pipeline of AlphaFold v2.0 [1] provided in https://github.com/deepmind/alphafold and installed on a local GPU cluster. MarvinSketch [2] was used for reconstructing, displaying, and characterizing the chemical structure of the small molecule ligands. Ligands were docked to the predicted 3D model of the receptor using Autodock Vina v1.1.2 [3]. Two models of APJ and four models of GPR139 receptor complexes were manually selected by visual inspection out of the top 10 ranked poses provided by Autodock Vina. These models were finally subjected to a brief energy minimization step using the molecular modeling toolkit [4] available in Chimera [5].

T03: KOR and T04: NPY1R (peptide ligands)

3D structures of KOR and NPY1R in complex with NPY were predicted using a full local version of the implementation of AlphaFold v2.0 [1] or AlphaFold-Multimer [6], respectively, provided in https://github.com/deepmind/alphafold. Dynorphin was docked to the predicted 3D model of KOR using the HADDOCK2.4 Web Server [7]. The top-ranked NPY1R-NPY complex and the four top-ranked KOR-Dynorphin complexes were subsequently refined using molecular dynamics (MD) simulations. The CHARMM-GUI builder [8,9] was used to embed each complex into a 80 x 80 Å$^2$ POPC membrane. Each protein-membrane system was placed into a water box made of explicit water molecules, their charge was neutralized, and the ionic strength of the system adjusted, throughout CHARMM-GUI builder's pipeline. All titratable residues of the protein were left in their dominant protonation state at pH 7.0, except for KOR Asp105 and NPY1R. Disulfide bridges were created between Cys131-Cys210 in KOR, and Cys42-Cys305 and Cys122-Cys207 in NPY1R. The geometry of the systems was first optimized by energy minimization and then equilibrated for 10 ns at constant pressure (NPT ensemble). The harmonic positional restraints initially applied to all Cα atoms of the protein were gradually released throughout the equilibration phase. Production simulations for three replicas were run for 50 ns each in the NPT ensemble at 1,013 bar and 310 K. All simulations were run using GROMACS v2021.4 [10] in combination with the CHARMM36m force field [11]. Finally, four KOR and three NPYR complexes were manually selected by visual inspection of the simulation trajectories.

### GPCRdock-6760 and GPCRdock-6867

#### Building Models of APJ, GPR139, KOR, NPY1R and NMUR2 for GPCR Dock2021

*Jiahan Li[13+], Ruidong Wu[1+], Rui Shen[1+], Zuofan Wu[1+], Rui Wang[1], Chaoran Cheng[2], Shitong Luo[1], Jiaqi Guan[2], Fan Ding[1], Jian Peng[12*]*

*[1]HelixonAI*
*[2]University of Illinois at Urbana-Champaign*
*[3]Peking University*
*+Equal Contribution*

A: PROTEIN MODEL BUILDING

As all five tasks are class-A GPCR based on UniRef[12], one doesn't bind (APJ, inactive state) and four bind to the G protein (active state), We collect a class-A GPCR-state-specific database from GPCRdb[1], similar to GPCR100[2]. Then we finetune our protein folding model on this database. Finally, we fold each task using our finetuned model with carefully selected MSAs or no MSAs and state-specific GPCR templates searching from our collected database. We also develop a no-MSAs-only-template strategy for folding each protein.

After folding, each protein is refined by AMBER Force Field[11].

B: LIGAND DOCKING AND COMPLEX REFINEMENT

We use different methods for small molecule docking (APJ, GPR139) and peptide docking (KOR, NPY1R, NMUR2).

For small-molecule docking, we first dock the small molecule to the GPCR using qvina2[3] with a human-annotated binding box around ECL2 and TMs regions, generating 10000 poses each. To evaluate different poses for a given receptor, we train an equivariant graph neural network on ReDock2020[4] for predicting ligand RMSD compared to its crystal-state pose. We rank and select different poses using the trained model. Then the protein/ligand complexes are refined by the UFF force field[13] in RDKit[9].

For peptide docking, we train a binding-site prediction model on Pepdb[5], we finetune this model on our collected GPCR dataset. We predict the binding site using receptor structure and peptide sequence. We combine our predicted binding site with peptimap[6]. Then we make use of the predicted binding constraints (binding site, contact map) and dock the peptide using CABS-dock[8] for 50 simulation cycles and HDOCK[8] with the specific binding site. We also refine the protein/peptide complexes using MMFF forcefield[14] in OpenMM[10].

**GPCRdock-6947**

#### Methods

##### Protein-small molecule docking

Our protein-small molecule docking protocol starts with the preparation of receptor structures. For two target complexes, APJ and GPR139, we chose chain B of known 5VBL PDB structure[1] and AlphaFold2[2] modeled structure of UniProt[3] ID Q6DWJ6[4], respectively. The protein structures were truncated and mutated with PyMOL[5] to match the provided sequences with experimentally resolved density. To locate the small molecule binding site, we identified hot spots using the computational mapping server FTMap[6]. FTMap uses the Fast Fourier Transform (FFT) correlation approach to sample the billions of small-molecular probe positions, and the clusters of multiple poses can be considered as hotspots. In both APJ and GPR139 cases, the majority of highly populated hot spots were located in the binding site within transmembrane helices, thus naturally defining the docking box.

For small-molecule docking, we used Glide[7,8,9] and AutoDock Vina[10] programs.

For Glide, the structures of all receptors were prepared using the Schrödinger protein preparation wizard[11,12], which provides minor structure optimization, ensuring the best performance of the following Glide docking runs. We used the standard setting of the wizard: add and optimize the positions of the hydrogen atoms and alter residue ionization/tautomer states using PROPKA[13,14] at pH 7, remove water molecules further than 3 Å away from the HET atoms, and perform restrained structure minimization with the OPLS4 force field[15] converging the heavy atoms to RMSD of 0.3 Å. The prepared structure was used to generate the grid for Glide docking. The center of the grid box was placed in the center of mass of the FTMap fragments, and the size of the box was set to 20 Å to ensure full coverage of the FTMap hotspots. We used a single-precision SP Glide docking protocol with a flexible ligand sampling: sampling nitrogen inversions and ring conformations. Epik[16,17,18] state penalties were used to adjust the scores. The initial 3D structures for the ligands were generated using Schrödinger LigPrep[19] with the following settings: use OPLS4 force field, generate possible ionization states at pH 7, use Epik, desalt, generate tautomers, retain specific input chiralities, and generate at most 32 structures per ligand.

For AutoDock Vina, the receptor and ligand structures were preprocessed and parameterized using AutoDockTools v1.5.7[20]. The ligand structures were constructed from the given SMILES strings[21] using RDKit[22] and manually checked to ensure the correct protonation state at pH 7. All AutoDock Vina calculations were carried out with default parameter values, except for the exhaustiveness, which was increased from the default value of 8 to 40 for a thorough search. The docking search space was confined to a 20 Å box large enough to cover the FTMap hotspots for both target cases.

Some manually selected Glide and AutoDock Vina docking structures were further refined using molecular dynamics (MD) simulations. Multiple independent simulations were run for each protein-ligand pair. Protein and ligand molecules were parameterized with Amber's ff14SB[23] and GAFF[24] force fields, respectively. Partial charges of the ligands were calculated using the AM1-BCC[25] charge method distributed with the AmberTools20 Antechamber program[26]. Structures were solvated in TIP3P[27] water in an octahedral box with a 10 Å buffer region around the protein and neutralized by adding the required amounts of sodium or chloride ions. Each system was minimized for 2 ns, followed by a production run of 50 ns at 300 K and 1 atm pressure; all simulations were carried out using OpenMM 7.6.0[28]. Restraints were applied to all heavy protein atoms throughout each run to retain the original shape of the protein. Clustering of the simulation results was performed on a combined trajectory of the last 25 ns of each simulation. Each trajectory was first aligned to the alpha-carbon atoms of the protein, and the L-RMSD matrix was then calculated for heavy atoms of the ligand. The result in each case was then clustered using DBSCAN[29] from scikit-learn 1.0.1[30], with a cutoff of 1 Å L-RMSD used to generate distinct ligand poses.

All the receptor-ligand binding structures by AutoDock Vina and Glide and MD cluster centroids were manually assessed and ranked to produce the final list of 5 models per target.

**Protein-peptide docking**

The three peptidic ligands were handled using a combination of ClusPro[31] peptide docking and AlphaFold[22,32] rescoring/refinement following an approach described previously[33]. Briefly, the modeling procedure was as follows. We started with generating protein-peptide complex models using default AlphaFold Multimer modeling capabilities. Then, we performed ClusPro runs, during which exhaustive rigid-body docking of peptide fragments to the receptor was carried out, followed by clustering, rescoring by cluster size, and flexible minimization of cluster centers. Additionally, we. Finally, we mixed the ClusPro and default AF2 models and re-scored/refined them with AF2 model_1_multimer by taking advantage of the AF2 template input. The resulting models were re-scored by predicted aligned error of the interface residues and submitted as our predictions.

8. Halgren, T. A. *et al.* Glide: A New Approach for Rapid, Accurate Docking and Scoring. 2. Enrichment Factors in Database Screening. *J. Med. Chem.* **47**, 1750–1759 (2004).
9. Friesner, R. A. *et al.* Extra Precision Glide: Docking and Scoring Incorporating a Model of Hydrophobic Enclosure for Protein−Ligand Complexes. *J. Med. Chem.* **49**, 6177–6196 (2006).
10. Trott, O. & Olson, A. J. AutoDock Vina: improving the speed and accuracy of docking with a new scoring function, efficient optimization and multithreading. *J. Comput. Chem.* **31**, 455–461 (2010).
11. Madhavi Sastry, G., Adzhigirey, M., Day, T., Annabhimoju, R. & Sherman, W. Protein and ligand preparation: parameters, protocols, and influence on virtual screening enrichments. *J. Comput. Aided Mol. Des.* **27**, 221–234 (2013).
12. Schrödinger Release 2022-2: Protein Preparation Wizard; Epik, Schrödinger, LLC, New York, NY, 2021; Impact, Schrödinger, LLC, New York, NY; Prime, Schrödinger, LLC, New York, NY, 2021.
13. Søndergaard, C. R., Olsson, M. H. M., Rostkowski, M. & Jensen, J. H. Improved Treatment of Ligands and Coupling Effects in Empirical Calculation and Rationalization of p $K_a$ Values. *J. Chem. Theory Comput.* **7**, 2284–2295 (2011).
14. Olsson, M. H. M., Søndergaard, C. R., Rostkowski, M. & Jensen, J. H. PROPKA3: Consistent Treatment of Internal and Surface Residues in Empirical p $K_a$ Predictions. *J. Chem. Theory Comput.* **7**, 525–537 (2011).
15. Lu, C. *et al.* OPLS4: Improving Force Field Accuracy on Challenging Regimes of Chemical Space. *J. Chem. Theory Comput.* **17**, 4291–4300 (2021).
16. Schrödinger Release 2022-2: Epik, Schrödinger, LLC, New York, NY, 2021.
17. Greenwood, J. R., Calkins, D., Sullivan, A. P. & Shelley, J. C. Towards the comprehensive, rapid, and accurate prediction of the favorable tautomeric states of drug-like molecules in aqueous solution. *J. Comput. Aided Mol. Des.* **24**, 591–604 (2010).
18. Shelley, J. C. *et al.* Epik: a software program for pK a prediction and protonation state generation for drug-like molecules. *J. Comput. Aided Mol. Des.* **21**, 681–691 (2007).
19. Schrödinger Release 2022-2: LigPrep, Schrödinger, LLC, New York, NY, 2021.
20. Morris, G. M. *et al.* AutoDock4 and AutoDockTools4: Automated docking with selective receptor flexibility. *J. Comput. Chem.* **30**, 2785–2791 (2009).
21. Weininger, D. SMILES, a chemical language and information system. 1. Introduction to methodology and encoding rules. *J. Chem. Inf. Model.* **28**, 31–36 (1988).
22. RDKit: Open-source cheminformatics. https://www.rdkit.org.
23. Maier, J. A. *et al.* ff14SB: Improving the Accuracy of Protein Side Chain and Backbone Parameters from ff99SB. *J. Chem. Theory Comput.* **11**, 3696–3713 (2015).
24. Wang, J., Wolf, R. M., Caldwell, J. W., Kollman, P. A. & Case, D. A. Development and testing of a general amber force field. *J. Comput. Chem.* **25**, 1157–1174 (2004).
25. Jakalian, A., Bush, B. L., Jack, D. B. & Bayly, C. I. Fast, efficient generation of high-quality atomic charges. AM1-BCC model: I. Method. *J. Comput. Chem.* **21**, 132–146 (2000).
26. Case, D. A. *et al. AMBER 20*.
27. Jorgensen, W. L., Chandrasekhar, J., Madura, J. D., Impey, R. W. & Klein, M. L. Comparison of simple potential functions for simulating liquid water. *J. Chem. Phys.* **79**, 926–935 (1983).

#### GPCRdock-7302

Numan Yousaf[a], Muhammad Muddassar*[a]

a) Department of Biosciences, COMSATS University Islamabad, Park Road, Islamabad, Pakistan.

#### Homology Modeling

The homology model of protein was predicted by I-TASSER web server that is a template-based method for protein structure prediction. I-TASSER first searches the PDB library (Berman et al., 2000) to identify structural templates by using LOMETS (Wu & Zhang, 2007), a threading program containing several threading algorithms. The full-length topology of models is then produced by reassembling the continuously-aligned fragment structures extracted from the templates, while the unaligned regions are built from scratch by implying Monte Carlo simulation (Y. Zhang, Kolinski, & Skolnick, 2003) based ab initio folding. To identify the low energy states, the trajectories of structures are clustered by SPICKER (Y. Zhang & Skolnick, 2004). Based on the SPICKER clusters, a second round of structure reassembly is conducted to refine the models. The structures with low energy states are further refined by full atomic simulations through ModRefiner (Xu & Zhang, 2011) and FG-MD (J. Zhang, Liang, & Zhang, 2011). Finally, the predicted model is matched with the proteins in BioLiP function library (Yang, Roy, & Zhang, 2012) to find the functional insights of query protein by sequence and structure profile comparisons (Roy, Yang, & Zhang, 2012; Yang, Roy, & Zhang, 2013).

#### Molecular Docking

The molecular docking studies was conducted by using Maestro tool of Schrodinger suite. The model generated by I-TASSER(Yang et al., 2015) was prepared by protein preparation wizard. The structure was preprocessed for structural integrity by adding hydrogen atoms and assigning bond orders for amino acids. In the refinement step, the tautomeric states and protonation were adjusted at 7.4 pH. The geometry of structure was optimized and OPLS_2005 force field (Shivakumar et al., 2012) was used to minimize the structure. The next step was to generate the grid at specific sites to conduct site specific docking. The prepared structure was superimposed on the reference protein structure (PDB ID: 5A8E) to find the binding sites of the model. The co-crystal ligand was used to generate the grid around specific binding site residues. Two positional constraints were also added. After grid generation, the GPR inhibitor was docked at the predicted binding site using the SP mode of Glide tool. The five best binding poses of GPR inhibitor in the binding site of model were selected based on the binding interactions with side chain of protein and their Glide scores.

#### GPCRdock-7350 (ComMedX)

#### Building models of APJ/Small molecule, GPR139/Small molecule, KOR/Dynorphin, NPY1R/NPY peptide and NMUR2/NMU25 complexes for GPCR Dock 2021

He Huang, Shuo Gu, Xiaobo Wan*

Computational Medicine Beijing Co., Ltd
Building 16, Beilun Industrial Park, 9 North Yongteng Road, Haidian District Beijing, China 100094

##### APJ

The protein structure we used are 5VBL [1] and 6KNM [2]. We used Glide in this docking study. The protein was prepared with a default protocol from Protein Preparation Wizard and OPLS3 force field. We applied Epik method in the ligand preparation with the range of pH from 6.4 to 8.4. Standard precision (SP) and extra precision (XP) docking protocols [3] were used to generate binding poses within a 30x30x30 Å box. We sampled up to 20 docking conformations and selected one with the best interaction with the protein.

##### GPR139

The homology model of GPR139 was from AlphaFold Protein Structure Database [4] (https://alphafold.ebi.ac.uk/entry/Q6DWJ6). We used Glide in this docking study. The detailed method was similar to APJ.

##### KOR

The 1st submitted model was the best ranking model built by the Colab version of AlphaFold Multimer [5]. The 2nd and 3rd models were built by redocking the 1st model using Rosetta FlexPepDock [6]. The 4$^{th}$ and 5$^{th}$ models were generated by Schrodinger's Glide. We used the XP and SP docking methods to sample up to 20 docking conformations and selected one with the best interaction with the protein.

##### NPY1R

All submitted models were built by the Colab version of AlphaFold Multimer.

##### NMUR2

The first three submitted models were the best ranking models built by the Colab version of AlphaFold Multimer. The 4th and 5th models were built by redocking the 1st model using Rosetta FlexPepDock.

### Criteria for selecting 5 best models per target

We manually select the poses for APJ and GPR139. Since both targets bind small molecules, we specifically focus on how the ligand can interact with the protein with less stranded hydrogens and less strain energy.

Alphafold rank, Rosetta generic full-atom energy score and conformation diversity. For the selection of docking model in KOR, we focused on the hydrogen bonding and hydrophobic interactions between the ligand the protein. In addition, we examined the internal strain of the ligand to make sure that the ligand itself should be in the low energy as well.

### Role of experimental information

#### APJ

From the APJ crystal structure, we predict the triazole group of ligand in the binding site forms electrostatic and hydrogen bonds with $R168^{4.64}$.

#### GPR139

Receptor mutants studied in a Ca2+ assay demonstrated that residues $F109^{3\times33}$, $H187^{5\times43}$, $W241^{6\times48}$ and $N271^{7\times38}$, but not $E108^{3\times32}$, are highly important for the activation of GPR139 as predicted by the receptor model [7]**.**.

#### NMUR2

From our literature survey, we gathered the following information: The 5 residues on the C terminus of Nmu-25 are highly conserved. Arg7 of Nmu-8 (corresponding to Arg24 of Nmu-25) was most important for the activity. Replacement of Arg5 of porcine Nmu-8 greatly reduced both potency and efficacy at murine NmU-R1 compared with murine NmU-R2. We were also aware of the active-like structure of neurotensin receptor NTSR1, which shares 32% sequence identity with NMUR2, in complex with neurotensin, a peptide ligand related to NmU. The information implied that the peptide must bind in a C-terminus inward mode, and the two arginines, especially Arg24, must be making important interactions in the binding pose. Other than Alphafold Multimer and FlexPepDock, we tried a few other peptide docking applications including HADDOCK, HPEPDOCK and MDockPep. The experimental information helped us make the judgement that models generated by Alphafold Multimer were more likely to be closer to reality.

### Target challenges

For NMUR2, not much experimental information was available to discern the conformation of the N-terminus of the peptide. We mostly relied on the scoring of the docking applications, namely Alphafold Multimer and Rosetta FlexPepDock, and were aware that the uncertainty of the predictions was high. For KOR and NPY1R, we didn't manage to carry out a thorough literature survey. It turned out that existing information is crucial for making accurate predictions, especially in the case of KOR.

### GPCRdock-7750 methods summary

| | APJ | GPR139 | KOR | NPY1R | NMUR2 |
|---|---|---|---|---|---|
| Model 1 | Homology Modeling (PDB: 5VBL) + Molecular Dynamics(MD) | AlphaFold + XDocK + MD | Homology Modeling (PDB: 4DJH) + FlexPepDock + MD | AlphaFold + ZDOCK + MD | AlphaFold-Multimer + MD |
| Model 2 | Homology Modeling (PDB: 5VBL) + MD | AlphaFold + XDocK + MD | AlphaFold-Multimer + MD | AlphaFold-Multimer + MD | AlphaFold-Multimer + MD |
| Model 3 | Homology Modeling (PDB: 5VBL) + MD | AlphaFold + XDocK + MD | AlphaFold-Multimer + MD | AlphaFold-Multimer + MD | AlphaFold-Multimer + MD |
| Model 4 | Homology Modeling (PDB: 5VBL) + MD | AlphaFold + XDocK + MD | AlphaFold-Multimer + MD | AlphaFold-Multimer + MD | AlphaFold-Multimer + MD |
| Model 5 | Homology Modeling (PDB: 5VBL) + MD | AlphaFold + XDocK + MD | Homology Modeling (PDB: 4DJH) + FlexPepDock + MD | AlphaFold-Multimer + MD | AlphaFold-Multimer + MD |

### GPCRdock-7750 T01 APJ

Lin Wang[a,b,#], Siyuan Tian [a,b,#], Shihang Wang [a,b,#], and Fang Bai [a,b,c,*]
*[a]Shanghai Institute for Advanced Immunochemical Studies, [b]School of Life Science and Technology, and [c]School of Information Science and Technology, ShanghaiTech University, 393 Middle Huaxia Road, Shanghai, 201210, China*

[#] These authors contribute equally

- Building the initial models of receptors (if receptors were modeled separately from ligands) or complexes. Did you use AlphaFold, RoseTTAFold, or any other deep-learning-based methodology for building the initial models? (if yes, please provide the info about the exact methodology and its implementation that you used). What other methods did you use, and how ?

APJ: APJ was molded by using the homology modeling module in Schrödinger2021-3 based on a template structure from PDB (PDB ID: 5VBL). The gap and point mutation were fixed.

- Small molecule docking and/or peptide ligand docking (as applicable)

APJ: According to the molecular interaction of ligand-APJ from the complex structure of 5VBL, the hydrophobic contacts formed between the ligand and Y271 and F291 were thought to be critical to the ligand-protein binding. Hence, hydrophobic interactions were set as constraints with these two residues, and induced fit docking from Schrödinger2021-3 was adopted to perform the complex prediction.

- Any model refinement (if applicable)

All those selected binding complexes were optimized by using the Prime module in Schrödinger2021-3. Then we used OPM server to predict the membrane position, and added cell membrane to the structure. After that, we run two 100ns' MD simulations with different random seeds for each complex. In the simulations, S-OPLS force field was taken. The trajectories from MD were clustered to get the average conformations. Then these conformations were further optimized with energy minimizations.

- Criteria for selecting 5 best models per target

The models were scored by using MM-GBSA module in Schrödinger2021-3. Both the MM-GBSA binding energies and visual inspection were considered in our ranking.

- Please include a section about target challenges. What did you find most challenging about each complex, and what information and approaches did you use to solve the challenge?

The present methods are very limited in predicting the induce fit effect occurring the binding between the ligands and proteins. Meanwhile, these end-to-end AI methods are also limited in reflecting the conformational changes, i.e., capturing different local minimum conformations.

- Please comment on the role of experimental information and human expertise in generation and selection of your models (if applicable).

In our cases, we did refer to types of experimental information, e.g., the 3D conformation of the peptide of NPY, the binding patterns between the know compounds with APJ, this information can help us to select simulation tools or reasonable binding poses.

### GPCRdock-7750 T02 GPR139

Lin Wang[a,b,#], Siyuan Tian [a,b,#], Shihang Wang [a,b,#], and Fang Bai [a,b,c,*]

*[a]Shanghai Institute for Advanced Immunochemical Studies, [b]School of Life Science and Technology, and [c]School of Information Science and Technology, ShanghaiTech University, 393 Middle Huaxia Road, Shanghai, 201210, China*

[#] These authors contribute equally

- Building the initial models of receptors (if receptors were modeled separately from ligands) or complexes. Did you use AlphaFold, RoseTTAFold, or any other deep-learning-based methodology for building the initial models? (if yes, please provide the info about the exact methodology and its implementation that you used). What other methods did you use, and how ?

GPR139: Based on the given sequence of GPR139, we used AlphaFold and Robetta webserver to do *de novo* protein modeling. Each method generated 5 predicted structures. In addition, three models constructed from homology modeling method were downloaded from GPCRdb. All these structures were sent to the Prime module in Schrödinger to optimize these predicted structures by using energy minimization and side-chain conformational prediction.

- Small molecule docking and/or peptide ligand docking (as applicable)

GPR139: JNJ-63533054 was docked against GPR139 by using our in-house docking protocol XDock (https://github.com/Wang-Lin-boop/Schrodinger-Script). As reported in the previous study, the residues L249 and V76 of GPR139 are essential to the ligand binding. The binding poses, which include such interactions, are paid pariticular attention to select the candidate binding poses. In addition, five candidate structures were further refined by running molecular dynamic simulations.

- Any model refinement (if applicable)

All those selected binding complexes were optimized by using the Prime module in Schrödinger2021-3. Then we used OPM server to predict the membrane position, and added cell membrane to the structure. After that, we run two 100ns' MD simulations with different random seeds for each complex. In the simulations, S-OPLS force field was taken. The trajectories from MD were clustered to get the average conformations. Then these conformations were further optimized with energy minimizations.

- Criteria for selecting 5 best models per target

The models were scored by using MM-GBSA module in Schrödinger2021-3. Both the MM-GBSA binding energies and visual inspection were considered in our ranking.

- Please include a section about target challenges. What did you find most challenging about each complex, and what information and approaches did you use to solve the challenge?

The present methods are very limited in predicting the induce fit effect occurring the binding between the ligands and proteins. Meanwhile, these end-to-end AI methods are also limited in reflecting the conformational changes, i.e., capturing different local minimum conformations.

- Please comment on the role of experimental information and human expertise in generation and selection of your models (if applicable).

In our cases, we did refer to types of experimental information, e.g., the 3D conformation of the peptide of NPY, the binding patterns between the know compounds with APJ, this information can help us to select simulation tools or reasonable binding poses.

### GPCRdock-7750 T03 OPRK

Lin Wang[a,b,#], Siyuan Tian [a,b,#], Shihang Wang [a,b,#], and Fang Bai [a,b,c,*]
[a]*Shanghai Institute for Advanced Immunochemical Studies,* [b]*School of Life Science and Technology, and* [c]*School of Information Science and Technology, ShanghaiTech University, 393 Middle Huaxia Road, Shanghai, 201210, China*

[#] These authors contribute equally

- Building the initial models of receptors (if receptors were modeled separately from ligands) or complexes. Did you use AlphaFold, RoseTTAFold, or any other deep-learning-based methodology for building the initial models? (if yes, please provide the info about the exact methodology and its implementation that you used). What other methods did you use, and how ?

KOR: Based on the given sequence of KOR, we used AlphaFold and Robetta webserver to do *de novo* protein modeling. Each method generated 5 structures. Meanwhile, we used homology modeling of Schrödinger2021-3 to build up three structures from different templates (PDB IDs: 4DJH, 6B73, and 6VI4). Finally, all these structures were sent to the Prime module in Schrödinger to optimize these predicted structures by using energy minimization and side chain prediction.

- Small molecule docking and/or peptide ligand docking (as applicable)

KOR: We used XDock protocol for docking and obtaining the protein-ligand complexes. Then, the obtained complex structures were further optimized by using FlexPepDock webserver.

- Any model refinement (if applicable)

All those selected binding complexes were optimized by using the Prime module in Schrödinger2021-3. Then we used OPM server to predict the membrane position, and added cell membrane to the structure. After that, we run two 100ns' MD simulations with different random seeds for each complex. In the simulations, S-OPLS force field was taken. The trajectories from MD were clustered to get the average conformations. Then these conformations were further optimized with energy minimizations.

- Criteria for selecting 5 best models per target

The models were scored by using MM-GBSA module in Schrödinger2021-3. Both the MM-GBSA binding energies and visual inspection were considered in our ranking.

- Please include a section about target challenges. What did you find most challenging about each complex, and what information and approaches did you use to solve the challenge?

The present methods are very limited in predicting the induce fit effect occurring the binding between the ligands and proteins. Meanwhile, these end-to-end AI methods are also limited in reflecting the conformational changes, i.e., capturing different local minimum conformations.

- Please comment on the role of experimental information and human expertise in generation and selection of your models (if applicable).

In our cases, we did refer to types of experimental information, e.g., the 3D conformation of the peptide of NPY, the binding patterns between the know compounds with APJ, this information can help us to select simulation tools or reasonable binding poses.

### GPCRdock-7750 T04 NPY1R

Lin Wang[a,b,#], Siyuan Tian [a,b,#], Shihang Wang [a,b,#], and Fang Bai [a,b,c,*]
[a]*Shanghai Institute for Advanced Immunochemical Studies,* [b]*School of Life Science and Technology, and* [c]*School of Information Science and Technology, ShanghaiTech University, 393 Middle Huaxia Road, Shanghai, 201210, China*

[#] These authors contribute equally

- Building the initial models of receptors (if receptors were modeled separately from ligands) or complexes. Did you use AlphaFold, RoseTTAFold, or any other deep-learning-based methodology for building the initial models? (if yes, please provide the info about the exact methodology and its implementation that you used). What other methods did you use, and how ?

NPY1R: Based on the given sequence of NPY1R, we used AlphaFold and Robetta webserver to do *de novo* protein modeling. Each server generated 5 structures. Two homology modeling structures were generated from two different templates (PDB ID: 5ZBH and 5ZBQ). These structures were sent to the Prime module in Schrӧdinger to optimize these predicted structures by using energy minimization and side-chain conformational prediction.

- Small molecule docking and/or peptide ligand docking (as applicable)

NPY1R: The peptide is too large to be perfectly docked to the receptor. Hence, we divided this peptide into two parts according to the secondary structure from an NMR structure(PDB ID: 1F8P). The first part is a loop region that covers the residues from 1 to 14 second part is the helix which consists of the residues from 15 to 36. We firstly dock the first part to the receptor by using peptide docking module in Schrödinger2021-3. Compared to the pocket shape and size of modeled structures of NPY1R, two models, one from AlphaFold and the other from Robetta webserver, which have bigger pockets were selected as receptor structures. Then, the center of the binding site was determined by using the coordinates of residues 73, 252, 94, 253, 185, 173, 275, 178, and 181. The grid for docking was estimated according to the length of the input conformation of the first part of the peptide. Glide SP-peptide scoring function was used to rank the candidate poses. Top candidate poses were visually inspected, and 9 of them were selected to be further optimized by prime module in Maestro of Schrödinger2021-3. The second part, whose conformation has been experimentally determined as the helix, was docked to the binding tunnel of NPY1R. Interestingly, we found the binding poses of the second part are very conserved. The conformation of the first part was very diverse. It indicated that the first part may present diverse conformations while interacting with the receptor. Hence, we selected five conformations of the first part which show different orientations to merge with the second part and eventually got the complete binding poeses.

- Any model refinement (if applicable)

All those selected binding complexes were optimized by using the Prime module in Schrödinger2021-3. Then we used OPM server to predict the membrane position, and added cell membrane to the structure. After that, we run two 100ns' MD simulations with different random seeds for each complex. In the simulations, S-OPLS force field was taken. The trajectories from MD were clustered to get the average conformations. Then these conformations were further optimized with energy minimizations.

- Criteria for selecting 5 best models per target

The models were scored by using MM-GBSA module in Schrödinger2021-3. Both the MM-GBSA binding energies and visual inspection were considered in our ranking.

- Please include a section about target challenges. What did you find most challenging about each complex, and what information and approaches did you use to solve the challenge?

The present methods are very limited in predicting the induce fit effect occurring the binding between the ligands and proteins. Meanwhile, these end-to-end AI methods are also limited in reflecting the conformational changes, i.e., capturing different local minimum conformations.

- Please comment on the role of experimental information and human expertise in generation and selection of your models (if applicable).

In our cases, we did refer to types of experimental information, e.g., the 3D conformation of the peptide of NPY, the binding patterns between the know compounds with APJ, this information can help us to select simulation tools or reasonable binding poses.

### GPCRdock-7750 T05 NMUR2

Lin Wang[a,b,#], Siyuan Tian [a,b,#], Shihang Wang [a,b,#], and Fang Bai [a,b,c,*]
*[a]Shanghai Institute for Advanced Immunochemical Studies, [b]School of Life Science and Technology, and [c]School of Information Science and Technology, ShanghaiTech University, 393 Middle Huaxia Road, Shanghai, 201210, China*
[#] These authors contribute equally

- Building the initial models of receptors (if receptors were modeled separately from ligands) or complexes. Did you use AlphaFold, RoseTTAFold, or any other deep-learning-based methodology for building the initial models? (if yes, please provide the info about the exact methodology and its implementation that you used). What other methods did you use, and how ?

NMUR2: Based on the given sequences of NMUR2 and NMU25 (peptide), we used AlphaFold multimer to build the complex model. Then we added the "-NH2" group to the C terminal of peptide NMU25. It is reported that the NMUR2 agonist peptide CPN-116 shared a similar sequence with the region of C terminal of NMU25. This indicates that NMU25 may bind to NMUR2 in its extracellular pocket is like CPN-116. It is further confirmed the prediction from AlphaFold Multimer.

- Small molecule docking and/or peptide ligand docking (as applicable)

NMUR2: The NMUR2 initial structure was built without using the docking-based approach.

- Any model refinement (if applicable)

All those selected binding complexes were optimized by using the Prime module in Schrödinger2021-3. Then we used OPM server to predict the membrane position, and added cell membrane to the structure. After that, we run two 100ns' MD simulations with different random seeds for each complex. In the simulations, S-OPLS force field was taken. The trajectories from MD were clustered to get the average conformations. Then these conformations were further optimized with energy minimizations.

- Criteria for selecting 5 best models per target

The models were scored by using MM-GBSA module in Schrödinger2021-3. Both the MM-GBSA binding energies and visual inspection were considered in our ranking.

- Please include a section about target challenges. What did you find most challenging about each complex, and what information and approaches did you use to solve the challenge?

The present methods are very limited in predicting the induce fit effect occurring the binding between the ligands and proteins. Meanwhile, these end-to-end AI methods are also limited in reflecting the conformational changes, i.e., capturing different local minimum conformations.

- Please comment on the role of experimental information and human expertise in generation and selection of your models (if applicable).

In our cases, we did refer to types of experimental information, e.g., the 3D conformation of the peptide of NPY, the binding patterns between the know compounds with APJ, this information can help us to select simulation tools or reasonable binding poses.

#### GPCRdock-7913

Author list: Xinqi Gong, He Huang, Daiwen Sun, Chengshi Zeng, Jiashan Li, Tong Wu, Nan Zhao, Ranxi Zhao
Address: Institute for Mathematical Sciences, Renmin University of China, Beijing, China
Contact person:

### Our methods:

Generally, we build the initial models of receptors based on AlphaFold-multimer and we used models generated by AlphaFold-monomer to replace the chain of the complex prediction structures with low prediction accuracy. Here we added some manual intervention when extracting MSA features, we used BLAST to do multiple sequence alignment for template search of monomers. Then we sorted the sequences by e-value from small to large, and selected the first or two as templates. For some sequences, the required submission starts from the middle of the sequence, so we selected the best alignment of the parts requested to be submitted as templates. Then the complex structures of ligand and receptor were generated by molecular docking such as AutoDock and MDockPP. Finally, the literature information about available binding site information for receptors and ligands were used as an experimental information to guide final model selection.

For target T01 and T02, We first modeled G protein and nanobody complexes by AlphaFold-multimer, and get the small molecule ligand structure through Openbabel. Then we used AutoDock to generated the complex structure of the receptor and the small molecule ligand. Finally, the submitted results were selected among the top 10 predictions based on binding site information.

For target T03, Alphafold2 and Alphafold-multimer was used to predict the structure of the monomer and complex separately, monomer prediction structures were used to replace the chain of the complex prediction structures with low prediction accuracy. Through multiple sequence alignment, the structure template of each chain was searched. 6VI4, 6OSA and 7DB6 were highly homologous to sequences of T03, and the B, C and D chains of 7DB6 were identical to the G_beta, G_gamma and Antibody sequences of T03, which could not only be used to refine the monomer structure, but also guide the formation of complex structure. We replaced the Alphafold-multimer prediction structure with part of the structure in 7DB6 as a template. Peptide_ligand corresponds to the nature structure of 2N2F, which has many different conformations. The different conformations of T03 and 2N2F were docked by AutoDock, and the docking results were sorted according to the binding site of 2N2F in Uniprot, and the best five candidate structures were selected.

For target T04 and T05, we firstly build complexes model based on AlphaFold-Complex, we find that chain A and chain E were not modeling well, so we generate models for them separately. We then generated 10 models for chain A, E with chain B, C, D based on MDockPP. By comparing the docking results of the two methods, it was shown that the binding conformation of chain A and chain E in the top 1 model of MDockPP and the AlphaFold2 docking results were almost the same, and this, so we choose this conformation as the first candidate conformation, and the other four candidate conformations were chosen by combining the site information about NPY1R in Uniprot. Finally, we analyzed the relative position of chain A and chain E, model 2, 3, 5, and 8 in the top 10 candidates were selected. Target T05 was modeled similar to T04, we build models of the chain A and chain E separately, and then use MDockPP for two body docking. In combination with the special sites of NMUR2 in uniprot, we selected models 1, 3, 6, and 9 in top10 as the last four candidate models.

##### Below is our methods summary:

APJ:
*AF multimer (apo), AutoDock
GPR139:
*AF multimer (apo), AutoDock
OPRK:
*AF monomer/multimer (apo), AutoDock (000[12345])
NPY1R:
*AF multimer (w/Dyn) (0001)
*AF monomer (apo), MDockPP (000[2345])
NMUR2:
*AF multimer (w/Dyn) (0001)
*AF monomer (apo), MDockPP (000[2345])

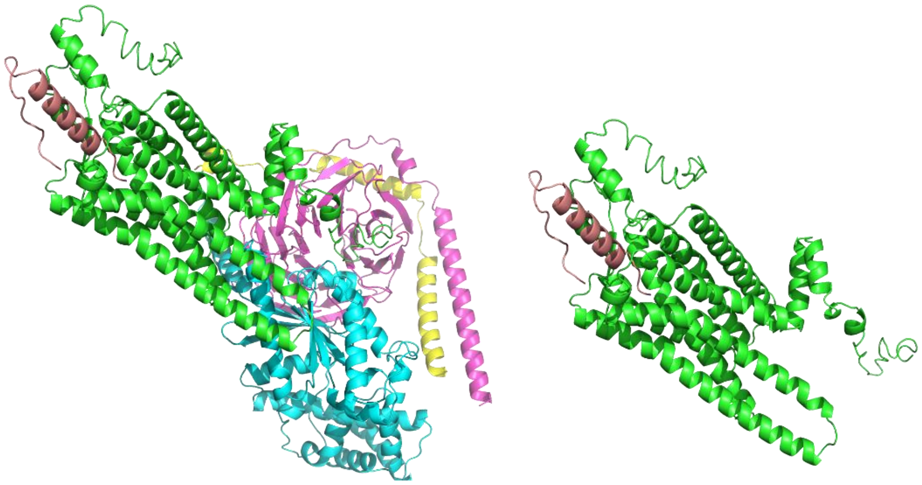

Fig 1. Example of our model (T04): the complex model of T04 and monomer model of chain A and E.

**GPCRdock-8156**

**Methods**

**Protein Model Building**

SWISS-MODEL[1] was used to build initial models of the target receptors. The 5VBL, 7L0P, 6DDF, 7LLU and 7L0P PDB experimental structures were used as templates to build the initial models of APJ, GPR139, kOR, NPY1R and NMUR2, respectively.

**Ligand/Peptide Docking**

For the ligand-bound systems (APJ and GPR139), Glide[2] was used to dock the small-molecule ligands to target receptors. The highest scored docking poses were selected to for refinement using Ligand Gaussian accelerated Molecular Dynamics (LiGaMD)[3] simulations.

For the peptide-bound systems (kOR, NMUR2 and NPY1R), the 2N2F and 1RON NMR structures were used for the Dynorphin and NPY peptides, respectively. The NMU25 peptide structure was predicted using AlphaFold2[4] as no experimental structure was available. The HPEPDOCK server[5] was then used to dock the Dynorphin peptide to the kOR and the NMU25 peptide to NMUR2. The HDOCK server[6] was used to dock the NPY peptide to the NPY1R receptor.

**Protein/Ligand complex Refinement and Selection**

Ligand Gaussian accelerated Molecular Dynamics (LiGaMD)[3] and Peptide Gaussian accelerated Molecular Dynamics (Pep-GaMD)[7] implemented in AMBER20[8] were applied to refine the GPCR-ligand and GPCR-peptide complexes, respectively. The AMBER14SB force field were used for the protein and lipid, and GAFF2 for ligand.

Simulations snapshots were aligned to the initial structure of each system regarding backbone atoms of the receptor transmembrane (TM) domain. They were then clustered using the hierarchical agglomerative clustering algorithm in CPPTRAJ[9] for the following atom selections: (1) heavy atoms in ligand/peptide, (2) backbone atoms in receptor TM domain, (3) backbone atoms in ECL2 and (4) heavy atoms within 5 angstroms of the ligand/peptide (i.e., binding pocket). The final models were selected from the top-ranked representative structural clusters after examining ligand/peptide-protein interactions (especially salt bridges, hydrogen bonds and aromatic interactions), secondary structures in peptides, ECL2 conformations and TM helices.

**Target Challenge**

One major challenge was to obtain accurate model structures of the target receptors. Homology modeling was mainly applied with SWISS-MODEL[1] to generate the receptor structures. However, the quality of these models could be poor especially in the binding pocket. This could result in inaccurate docking poses of the ligand/peptide. Moreover, it remained challenging to refine low-quality docking models using LiGaMD/Pep-GaMD simulations, due to limited competition time and short simulation lengths. We could not observe converged ligand binding pose in simulations of the APJ and GPR139 systems. For the peptide-bound systems, an additional challenge was to predict accurate receptor-bound conformations of the peptides, which were significantly different from those in the peptide alone states. Since the latter was used in our calculations, low-quality models were predicted for docking structures of GPCR-peptide systems. Significantly longer Pep-GaMD simulations would be needed to optimize the GPCR-peptide complex structures.

### GPCRdock-8321 T01 APJ

**Protocol:**
We used our recently published shape restrained High Ambiguity Driven DOCKing protocol (https://doi-org.proxy.library.uu.nl/10.1021/acs.jcim.1c00796).

The protocol consists of 4 steps:
- The identification structure of the receptor (=template receptor) bound to a ligand (=template ligand) that shares similarity with the ligand of interest
- The conversion of the template ligands' atoms into dummy beads or pharmacophore beads
- The definition of distance restraints between the atoms of the ligands of interest and the beads
- The distance driven HADDOCKing (rigid body docking + simulating annealing)

**- Structure selection**
Herein, we selected the 5VBL structure as our template structure and the residues 10:17 from its bound peptide as the template ligand.

**- Shape definition:**
The 5VBL ligand was converted into a shape of dummy beads and distances restraints of 1.0A were defined from the ligand (to dock) to the shape.

- **Docking**
Molprobity was use to handle the protonation state of Histidines. We generated a total of 46 conformers using the RDKIT ETKDG algorithm. The ligand topology as required by HADDOCK was generated with PRODRUG. We used small molecule default HADDOCK settings, generated 920 models in the rigid body docking step of HADDOCK, 200 of which were transferred to the simulated annealing step. All mutated residues of 5VBL were mutated back to native residues.

### GPCRdock-8321 T02 GPR139

We used our recently published shape restrained High Ambiguity Driven DOCKing protocol (https://doi-org.proxy.library.uu.nl/10.1021/acs.jcim.1c00796).

The protocol consists of 4 steps:
- The identification structure of the receptor (=template receptor) bound to a ligand (=template ligand) that shares similarity with the ligand of interest
- The conversion of the template ligands' atoms into dummy beads or pharmacophore beads
- The definition of distance restraints between the atoms of the ligands of interest and the beads
- The distance driven HADDOCKing (rigid body docking + simulating annealing)

We designed 2 different protocols that use different starting conformations of the GPR139. For both protocols, we derived shapes of dummy beads from 5DHG ligand, Fpocket output and 7L0P peptide, but we only describe the protocol that was used to generate the models that we selected for submission based on the HADDOCK score (all selected models are ranked within the top20) and visual inspection.

>>> protocol 1 (T02_1, T02_2)
-----------------------------

- Modeling of the receptor with AlphaFold:
The Jupyter Colab default settings were used to model GPR139 (https://colab.research.google.com/github/sokrypton/ColabFold/blob/main/AlphaFold2.ipynb).
Provided that JNJ-63533054 is an agonist of GPR139, we modeled GPR139 in the presence of the provided G-protein alpha subunit to obtain an agonist conformation of GPR139.

- Shape definition:
Due to the lack of template ligand in the PDB, we used the pocket predictor Fpocket (https://bioserv.rpbs.univ-paris-diderot.fr/services/fpocket/) to characterize the ligand binding site. Fpocket generates ensembles of beads that describe cavities. We selected the one corresponding to the GPR139 binding site and we clustered the beads using the Butina clustering algorithm with a 2.0A cutoff. We defined distance restraints of 2.0A from the shape (smallest) to the ligand (largest). Of note, the distance restraints threshold of 2.0A was favored over the usual 1.0A because the beads derived from fpocket output do not take into account the structure of the ligand.

>>> protocol 2 (T02_3, T02_4, T02_5)
-----------------------------

- Modeling of the receptor with modeler:
XXXXX

- Shape definition:
We derived a shape of dummy beads from 5DHG ligand and defined distance restraints of 2.0A from the shape (smallest) to the ligand (largest).
Of note, the distance restraints threshold of 2.0A was favored over the usual 1.0A because the beads are derived from 5DHG (antagonist) ligand whose receptor conformation differs from the one we perform the docking on, and because 5DHG ligand is antagonist and we are seeking for an agonist conformation.

Common to both protocols:
-----------------------------

Molprobity was used to handle the protonation state of Histidines. We generated a total of 20 conformers of JNJ-63533054 using the RDKIT ETKDG algorithm. The ligand topology as required by HADDOCK was generated with PRODRUG. Residues 392:406 of the GPR139 model were considered fully flexible during the simulating annealing step of HADDOCK.

We used small molecule default HADDOCK and buried (inter_rigid=0.001 and w_vdw_0=0) settings, generated 1000 models in the rigid body docking step of HADDOCK, 200 of which were transferred to the simulated annealing step.

### GPCRdock-8321 T03 OPRK

We used our recently published shape restrained High Ambiguity Driven DOCKing protocol (https://doi-org.proxy.library.uu.nl/10.1021/acs.jcim.1c00796) in combination with receptor and complex modelling as well as a final refinement with HADDOCK 2.4.

The protocol consists of 6 steps:
- The identification structure of the receptor (=template receptor) bound to a ligand (=template ligand) that shares similarity with the ligand of interest
- The conversion of the template ligands' atoms into dummy beads or pharmacophore beads
- The definition of distance restraints between the atoms of the ligands of interest and the beads
- The distance driven HADDOCKing (rigid body docking + simulating annealing)
- Modelling the complete complex with Modeller12.9
- Complex refinement with HADDOCK2.4

**Receptor modeling**

The receptor was modelled with Modeller 9.12 and by using PDB ID 6B73 as a template, a constitutively active form of the kappa-opioid receptor, to complete the missing loops. 30 models were generated in total which were ranked according to their DOPE score.

**Shape definition**

The 6B73 structure was superimposed to our receptor model, the 6B73 ligand was converted into a shape of dummy beads and we defined distance restraints of 2.0A from the shape (sm\
allest) to the ligand (largest). Of note, the distance restraints threshold of 2.0A was favored over the usual 1.0A because the beads are derived from 6B73 ligand which is bound to a receptor conformation that slightly differs from our model.

**Initial peptide conformers**

Residues 1:9 (YGGFLRRIR) from 2N2F were used.
Residues 1:4 were set fully flexible during the simulating annealing step of the docking.

**Docking**

We used peptide default HADDOCK settings and generated 1000 models in the rigid body docking step of HADDOCK, 200 of which were transferred to the simulated annealing step. Through HADDOCK score analysis and visual inspection, we selected a model showing a good anchorage of the 1:9 peptide into the binding site

**Modelling the complete complex**

This docked pose was used as a base for modelling the complete complex. The receptor conformation used in Modeller was based on the structure from the selected docked model. The NMR data of the free peptide (PDB ID 2N2F) were aligned with the peptide fragment present in the docked pose. Peptide residues 1-5 of the docked pose were maintained and residues 6-8 were extracted from the NMR data and merged with the 1-5 residue fragment. The N-terminal tail of the peptide, RPKLK, was modelled into the setup structure with Modeller 9.12 (100 models were generated). In a second modelling approach the initial receptor model (before docking) and the pose of the docked fragment were used. Both the initial receptor model and docked fragment were merged and residues 6-8 were taken from the available NMR data. 100 models were generated for this modelling setup which were ranked according to their DOPE score.

**Complex refinement**

We refined five selected models using the Energy Minimization tool of the HADDOCK2.4 Webserver, keeping the termini of the peptide charged.

### GPCRdock-8321 T04 NPY1R

We used our recently published shape restrained High Ambiguity Driven DOCKing protocol (https://doi-org.proxy.library.uu.nl/10.1021/acs.jcim.1c00796) in combination with Modeller and structure refinement with HADDOCK2.4.

The protocol consists of 6 steps:
- The identification structure of the receptor (=template receptor) bound to a ligand (=template ligand) that shares similarity with the ligand of interest
- The conversion of the template ligands' atoms into dummy beads or pharmacophore beads
- The definition of distance restraints between the atoms of the ligands of interest and the beads
- The distance driven HADDOCKing (rigid body docking + simulating annealing)
- Modelling the complete complex with Modeller 12.9
- Complex refinement with HADDOCK2.4

**Modeling of the receptor with AlphaFold**
The Jupyter Colab default settings were used to model NPY1R (https://colab.research.google.com/github/sokrypton/ColabFold/blob/main/AlphaFold2.ipynb). Provided that the peptide is an agonist of NPY1R, we modeled NPY1R in the presence of the provided G-protein alpha subunit to obtain an agonist conformation of NPY1R.

**Shape definition**
Due to the lack of template ligand in the PDB, we used the pocket predictor Fpocket (https://bioserv.rpbs.univ-paris-diderot.fr/services/fpocket/) to characterize the ligand binding site of NPY1R. Fpocket generates an ensemble of beads that describe cavities. We selected the one corresponding to the NPY1R binding site and we clustered the beads using the Butina clustering algorithm with a 2.0A cutoff. We defined distance restraints of 2.0A from the shape (smallest) to the ligand (largest).
Of note, the distance restraints threshold of 2.0A was favored over the usual 1.0A because the beads derived from fpocket output do not take into account the structure of the ligand.

**Initial peptide conformers**
The peptide conformation was predicted with Apptest. Provided that most study report an almost absolute conservation of the C-terminal part of NMU25, and that NPY shares a LXXRXRX C-terminal motif with NMU25, we followed the same logic as for T05: we docked the C-terminal (31:37) part of the NPY peptide into the NPY1R binding site.

**Docking**
We used peptide default HADDOCK settings and generated 1000 models in the rigid body docking step of HADDOCK, 200 of which were transferred to the simulated annealing step. Through HADDOCK score analysis and visual inspection, we selected a model showing a good anchorage of the 31:37 peptide (ITRQRY-NH2) into the binding site.

**Modelling the complete complex**
Starting from this model, we aligned the apptest peptide prediction with the peptide fragment (INLITRQRY) in the docked pose. The orientation of the predicted peptide part (PGEDAPAEDMARYYSALRHY) was adjusted by reducing the residue clashes between receptor and peptide. This orientation was used as a starting position for the peptide during modelling with Modeller 12.9. The receptor conformation was based on the structure from the selected docked model. 80 models were generated for this modelling step. The flexible N-terminal tail of the peptide (Y) was optimized with a second round of modelling in which 100 models were obtained.

**Complex refinement**

We refined five generated models using the Energy Minimization tool of the HADDOCK2.4 Webserver, keeping the N-terminal charged and the C-terminal neutral, including a $NH_2$ cap.

### GPCRdock-8321 T05 NMUR2

We used our recently published shape restrained High Ambiguity Driven DOCKing protocol (https://doi-org.proxy.library.uu.nl/10.1021/acs.jcim.1c00796) in combination with Modeller and structure refinement with HADDOCK2.4.

The protocol consists of 6 steps:
- The identification structure of the receptor (=template receptor) bound to a ligand (=template ligand) that shares similarity with the ligand of interest
- The conversion of the template ligands' atoms into dummy beads or pharmacophore beads
- The definition of distance restraints between the atoms of the ligands of interest and the beads
- The distance driven HADDOCKing (rigid body docking + simulating annealing)
- Modelling the complete complex with Modeller 12.9
- Complex refinement with HADDOCK2.4

**Modeling of the receptor with AlphaFold**
The Jupyter Colab default settings were used to model NMUR2 (https://colab.research.google.com/github/sokrypton/ColabFold/blob/main/AlphaFold2.ipynb).
Provided that the peptide is an agonist of NMUR2, we modeled NMUR2 in the presence of the provided G-protein alpha subunit to obtain an agonist conformation of NMUR2.

**Shape definition**
Due to the lack of template ligand in the PDB, we used the pocket predictor Fpocket (https://bioserv.rpbs.univ-paris-diderot.fr/services/fpocket/) to characterize the ligand binding site of NMUR2. Fpocket generates an ensemble of beads that describe cavities. We selected the one corresponding to the NMUR2 binding site and we clustered the beads using the Butina clustering algorithm with a 2.0A cutoff. We defined distance restraints of 2.0A from the shape (smallest) to the ligand (largest).
Of note, the distance restraints threshold of 2.0A was favored over the usual 1.0A because the beads derived from fpocket output do not take into account the structure of the ligand.

**Initial peptide conformers**
The peptide conformation was predicted with Apptest. Provided that most studies report an almost absolute conservation of the C-terminal part of NMU25, we docked the C-terminal (20:26) part of the NMU25 peptide into the NMUR2 binding site.

**Docking**
We used the shape based protocol with peptide default HADDOCK settings and generated 1000 models in the rigid body docking step of HADDOCK, 200 of which were transferred to the simulated annealing step. We defined backbone dihedral angle restraints from structure (ssdihed=alpha) to maintain the helices conformation. Through HADDOCK score analysis and visual inspection, we selected a model showing a good anchorage of the 20:26 peptide (LFRPRN-NH2) into the binding site.

**Modelling the complete complex**
Starting from this model, we aligned the Apptest predicted peptide conformation with the peptide fragment in the docked pose. In the modelling protocol performed with Modeller 12.9, the C-terminal part of the peptide originated from the docked pose (RPRN), while the predicted peptide conformation was used for the rest of the peptide (FRVDEEFQSPFASQSRGYFLF). The receptor conformation was based on the structure from the selected docked model. 200 models were generated which were ranked according to their DOPE score.

**Complex refinement**

We refined five generated models using the Energy Minimization tool of the HADDOCK2.4 Webserver, keeping the N-terminus of the peptide positively charged and the C-terminus neutral with an NH2 cap.

#### GPCRdock-8407

##### Model building

The computational models of kOR-Gi were constructed in Modeller[1] using PDBs 6B73 and 6DDE as templates. The multiple-template sequence alignment and build generation strategy were applied in the model building. A total of 20'000 initial models were generated. The best model was chosen for the model refinement according to the DOPE scores.

##### Model Refinement

All-atom molecular dynamics (MD) simulation was introduced for model refinement. The simulation system of kOR-Gi was constructed in CHARMM-GUI[2] under CHARMM36m[3] forcefield. The kOR-Gi complex model was embedded in 326 POPC lipid bilayers. The whole system was solvated with 42'000 water molecules and 0.15 M NaCl was added to bulk environment to mimic the physiological condition. A total of 200 ns MD simulation was conducted in Gromacs[4] for the kOR-Gi membrane system. To preserve the correct secondary structure conformation, the backbone of both kOR TM region as well as that of Gi protein were retrained with 10 kcal/(mol.Å$^2$) force constant. The lowest energy snapshot was extracted for further study.

The initial peptide was constructed in Maestro[5], followed by Ligprep to generate the optimal 3D conformation. Then, it was solvated with 2200 water molecules, of which 0.15 M NaCl was added. After that, a total of 1000 ns MD simulation was conducted in Gromacs. More than 5000 conformations were yielded, which were clustered into 5 different groups. The representative conformation of each cluster was extracted as an input for peptide docking.

##### Peptide docking

To obtain a better docking result, a deep learning model and score was constructed to evaluate the docking results. All the GPCR experimental structures of protein-ligand complex were downloaded from PDB database. The Convolutional Neural Networks (CNN)[6] algorithm was introduced to analyze all protein-ligand interaction fingerprint. Properties such as interaction distance, interaction types, atom types were included in the model training. An AI model was built which could be used to predict optimal binding mode of GPCR ligand. The initial binding conformations of peptide were generated in AutoDock CrankPep[7]. During the peptide docking, 200 independent searches were performed and each search using up to 10 million evaluations of the scoring function. Residues close to the ECL2 loop were kept flexible during the docking. The top 50 yielded conformations were submitted to AI model for further evaluation. The top 10 ranked models were checked manually and 5 models were selected as the final results.

##### Brief Discussion

The most challenging parts for modelling of kOR-Gi-Peptide model include the following (1) The model of kOR. Since all the resolved kOR structures only bound to small molecules, the binding pocket of kOR is too small to bind to a peptide molecule. If we dock the peptide molecule into kOR experimental structure directly, the peptide cannot accommodate to the peptide at all. Many atom clashes will be found. To overcome this problem, we introduced all-atom restrained MD simulations. After MD, the ECL2 loop opened a large space for ligand docking. (2) The relative position between kOR and Gi protein. When we analyzed all resolved GPCR-G protein complex structures, we found that GPCR and G protein had differently relative position. In this case, we took the one from μOR followed by MD simulations. (3) Peptide docking. Since the peptide molecule is flexible and large, it is very difficult to capture reliable pose for kOR. In this case, we used AutoDock CrankPep to generate the initial binding pose, then we introduced GPCR deep learning model to rescore the binding pose.

In the whole modelling processes, human expertise is also important for the judgement such as: choosing the output conformation of kOR-Gi complex from MD simulation; which residues should be kept flexible during docking; how to judge the final model according to AI predictions.

### GPCRdock-8503

1. Protein models building

The Apelin receptor in complex with an agonist peptide (PDB code: 5VBL) was selected as a template for APJ protein modeling. The full-length sequence alignment was performed on the NCBI Blast website (https://blast.ncbi.nlm.nih.gov/Blast.cgi) and mutated amino acids were corrected using UCSF Chimera. Hydrogens atoms were added via Protein Local Optimization Program (PLOP)[1].

The structure of GPR139 was predicted using AlphaFold v2.0[2], followed by side chain sampling with PLOP.

2. Protein-ligand structures generation

The ligand for APJ or GPR139 was docked into the corresponding receptor using AutoDock Vina[3]; meanwhile, rigid docking and flexible docking methods were used separately to generate induced-fit structural models. Since there are reported APJ structures, we also used RDKit for ligand sampling based on the maximum common substructure shared with the peptide in crystal structure (PDB code: 5VBL).

3. Protein/ligand complex refinement and selection

PLOP was used to perform energy minimization and rescoring. Protein-ligand complex models were selected based on the binding pose and total free energy. These selected models were then refined via a 10 ns full-atom MD simulation in POPC bilayer at 300 K using Gromacs 2021.2 package with CHARMM36m force-field parameters[45]. Ligand parameters were generated via CGenFF[6]. Input parameters for energy minimization, equilibration and production were generated by CHARMM-GUI (https://charmm-gui.org/). Snapshots in the last 4 ns MD trajectory with more than two hydrogen bonds between the receptor and the ligand were extracted and manually selected as the final solution.

---

#### GPCRdock-8824

- Building the initial models of receptors (if receptors were modeled separately from li gands) or complexes. Did you use AlphaFold, RoseTTAFold, or any other deep-learnin g-based methodology for building the initial models? (*if yes, please provide the info a bout the exact methodology and its implementation that you used*). What other met hods did you use, and how?

1. Receptor modeling was performed using AlphaFold-multimer in the full complex context p rovided by organizers. All 5 models are run, and then one with the highest pLDDT was chose n. If the target had peptide ligand, ligand was modeled altogether as well.

- Small molecule docking and/or peptide ligand docking (as applicable)
- Any model refinement (if applicable)

2,3. Small-molecule docking was performed using GALigandDock. Peptides-ligands are mode led by AlphaFold-multimer originally, but their poses were diversified by in-house tools and/ or by running **Rosetta flexpepdock protocol.**

- Criteria for selecting 5 best models per target

4. Was not systematic. Docked poses were initially filtered by deep-learned score (say ~50). Of these, higher priority (model 1~3) chosen by human inspection, followed by best deep-le arned scored ones for lower priority models (model 4~5). Small-molecule binders are ranked by DeepAccNet-ligand, which is small-molecule version of DeepAccNet (Hiranuma et al 2021 ). Protein-peptide binders are ranked by the original DeepAccNet.

- Please include a section about target challenges. What did you find most challenging about each complex, and what information and approaches did you use to solve the challenge? Please comment on the role of experimental information and human exp ertise in generation and selection of your models (if applicable).

5.
Target 1: Ligand size made it hard, also the role of the lipid molecule inserted in the TM helic es (also found in related expt PDB structure) to the ligand binding mode remains to be addre ssed. If so, the lipid molecule poses a challenge whether lipid molecules should also be expli citly treated in the docking scenario.
Target 2. Receptor modeling. Pocket varied quite a lot, so that modeling strategy largely infl uenced the docking results. It was hard to judge what modeling strategy will be the best *a pr iori*.
Target 3~5: Peptide modeling was most challenging, because methods I utilized were not we ll benchmarked or studied.

6. Activity of the ligand (whether it is antagonist or agonist) including the full sequence of th e target complex structure will be useful. Mutation studies were not very helpful because it was rather ambiguous to decipher whether the mutation changes function (usually deactiva

tes) due to interaction change or receptor stability change or even something else. Overall, model selection by human-intuition was not very successful in my experience, which can be different for other participants.

GPCRDock 2021 Methods

Congzhou Sha, Jiaxing Chen, Jian Wang, Nikolay Dokholyan

Group: GPCRdock-9077

#### Determination of G-protein-coupled receptor structure (T01 through T05)

We used AlphaFold to predict the structure of all five GPCR targets, using the default settings in the Google Colab notebook provided by the authors of AlphaFold[1]. For the three GPCR targets with peptide ligands (T03, T04, T05), we used the hetero-oligomer modelling feature of the AlphaFold notebook to predict an initial pose for the peptide ligands. The default settings of AlphaFold provide five possible structures for each of the GPCR targets, and we chose the most highly ranked GPCR structure for further evaluation. We ignored the C-terminal amine group modifications for T04 and T05, as the discrete molecular dynamics package we used for further evaluation does not support such groups.

#### Docking of small molecules (T01, T02)

We converted the SMILES format of the two small molecule ligands into mol2 files for use with MedusaDock[2,3]. MedusaDock randomly samples conformations of all rotatable bonds in the ligand along with protein receptor sidechains and evaluates the resulting poses using the MedusaScore force field[4]. We ran MedusaDock for thousands of iterations (seven days on a single core of a 2.5 GHz Intel Xeon E5-2680 v3 processor, with 2 GB RAM) and clustered the results by root mean square deviation (RMSD).

Clusters were determined using the Clustering.jl package in Julia 1.6. The clustering coefficient was chosen manually to provide a diverse range of structures. The clustering coefficient for T01 was 2.0 angstroms, and the clustering coefficient for T02 was 1.9 angstroms. We then chose the five lowest energy structures as scored by MedusaScore as our final docked poses.

*Table 1: Submitted structure IDs, the number of structures in the same cluster as the submitted structure, and the corresponding energies are shown for T01 and T02.*

| **T01** | | |
|---|---|---|
| *Structure ID* | *Number in cluster* | *Energy (kcal/mol)* |
| T01-1 | 520 | -54 |
| T01-2 | 111 | -44 |
| T01-3 | 335 | -49 |
| T01-4 | 119 | -45 |
| T01-5 | 1014 | --79 |
| **T02** | | |
| *Structure ID* | *Number in cluster* | *Energy (kcal/mol)* |
| T02-1 | 21 | -32 |
| T02-2 | 71 | -29 |
| T02-3 | 372 | -31 |
| T02-4 | 67 | -29 |
| T02-5 | 145 | -30 |

#### Docking of peptides (T03, T04, T05)

Starting with the most highly ranked pose of AlphaFold for each of the peptide-GPCR complexes, we ran discrete molecular dynamics simulations[5–8] at a variety of low temperatures, from 176 K to 231 K, to sample nearby states. Discrete molecular dynamics uses a discretized all-atom force field to calculate collisions, and is thus able to sample conformations at a faster rate than traditional molecular dynamics[5]. However, parameters do not exist for C-terminal amine groups in the software we used, so we ignored those modifications in the T04 and T05 peptide ligands. Simulations were carried out on a single core of the same processor as above, and run for a week. A step size of 1000 was chosen, and simulations were carried out for ~400,000-500,000 steps (seven days of computation), where each step corresponds to approximately 50 ps. This conversion is inexact due to the discrete nature of the simulations. The temperatures were chosen based on estimated specific heat to ensure that no major protein unfolding events occurred[9]. The simulations were then clustered using TTClust[10] with default settings, and these clusters were chosen as our final docked poses.

Since we expected that AlphaFold would provide an accurate assessment of binding site for each of the peptides, we explored only the nearby low energy states of the initial structure.

*Figure 1: The clustering dendrogram for T05 is shown, as output by TTClust. Four clusters were identified for T04 and T05, while five clusters were identified for T03.*

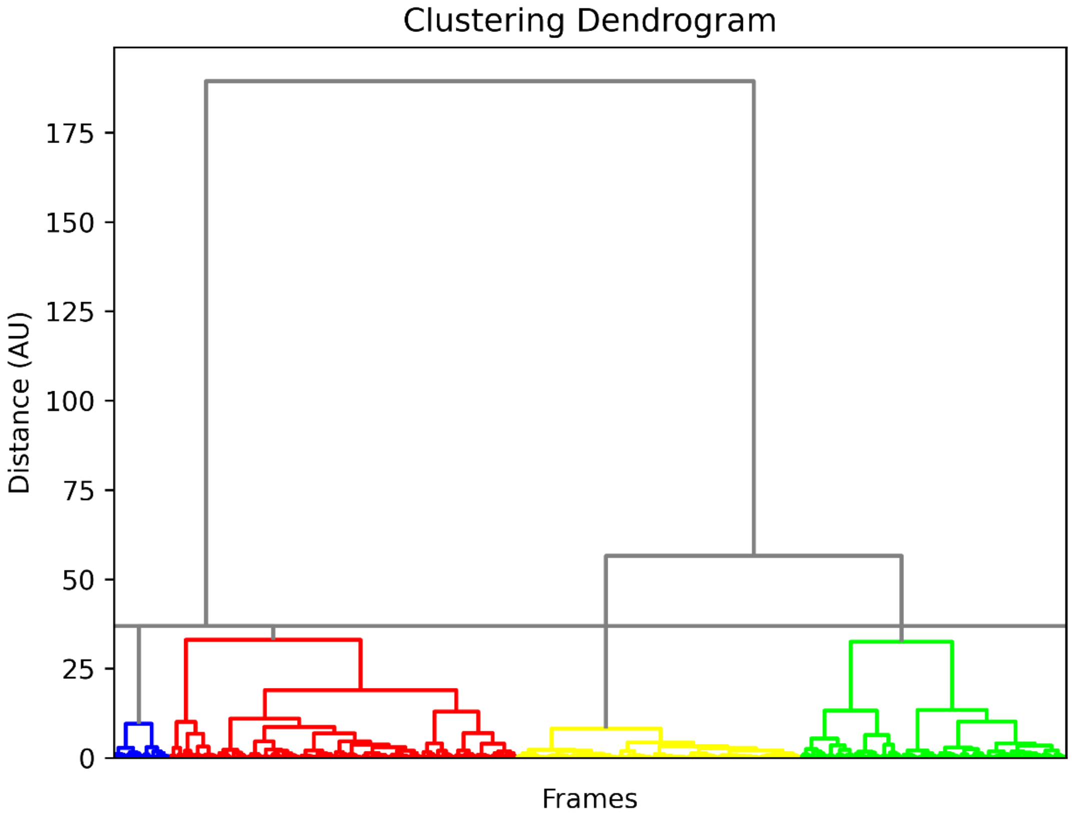

**Author contributions**

CMS performed the simulations and performed clustering. JC and JW provided technical support and various scripts for the work performed. All participants contributed to the design of our methods and discussed the data.

### GPCRdock-9686 Target 1 APJ

#### Receptor Model Generation

A model of the receptor was generated by a local installation of AlphaFold [1], in which all 5 models were modeled in what appeared to be an "open" conformation. The top-ranked model was selected and extracted for docking.

#### Docking

The receptor model was prepared using the standard protein preparation wizard in the Maestro interface. This process includes a hydrogen bond network optimization and a restrained minimization. The SiteMap application [2], through the Maestro interface was used to identify the binding site. The top-ranked binding site identified by SiteMap was the expected binding site (within the helical bundle on the extracellular side). A Glide-XP [3] receptor grid was generated at this site with a box size of 14Å. The ligand SMILES sequence was converted to an SDF file with the JMol SMILES parser [4], imported into Maestro, and prepared with the default settings in the LigPrep application. The ligand was docked into the receptor grid using Glide-XP, and the top 5 docking poses were extracted for further processing. All ligand poses were very precise, with only small variations in extended moieties (such as ring flips).

#### Refinement and Submission

Each of the 5 docked poses were put through molecular dynamics simulations using NAMD [5] for confirmation sampling. Models were simulated at 300k for 100 ns, with the transmembrane helices' alpha carbons harmonically restrained. The ligand was parameterized through CHARMM-GUI [6].

Because of the high precision in the docking results, our top 5 submissions are largely populated with these. The selected models are as follows: 1) Top ranked Glide pose, 2) Top ranked Glide pose + minimization, 3) Second ranked Glide pose, 4) Lowest Energy from MD refinement, and 5) Third ranked Glide pose. Files were cleaned in PyMol to match standard PDB format conventions before submission.

### GPCRdock-9686 Target 2 GPR139

#### Receptor Model Generation

A model of the receptor was generated by a local installation of AlphaFold [1], However, the models all appeared to be in a "closed" conformation. This was fixed by subjecting the top ranked model to a 10ns MD simulation. This model was then used for Docking.

#### Docking

The receptor model was prepared using the standard protein preparation wizard in the Maestro interface. This process includes a hydrogen bond network optimization and a restrained minimization. The SiteMap application [2], through the Maestro interface was used to identify the binding site. The second-ranked binding site identified by SiteMap was the expected binding site (within the helical bundle on the extracellular side). A Glide-XP [3] receptor grid was generated at this site with a box size of 14Å. The ligand SMILES sequence was converted to an SDF file with the JMol SMILES parser [4], imported into Maestro, and prepared with the default settings in the LigPrep application. The ligand was docked into the receptor grid using Glide-XP, and the top 5 docking poses were extracted for further processing.

#### Refinement and Submission

Just as for target 1, the 5 docked poses were put through molecular dynamics simulations using NAMD [5]. Models were simulated at 300k for 100 ns, with the transmembrane helices' alpha carbons harmonically restrained. The ligand was parameterized through CHARMM-GUI [6].

Literature Information tells us that the Halogen-containing ring is likely to be facing out of the binding pocket, so any trajectories where this is not the case were removed. The selected models are as follows: 1) Top ranked Glide pose, 2) Top ranked Glide pose + minimization, 3-5) Frames from refinement MD at local energy minima. Files were cleaned in PyMol to match standard PDB format conventions before submission.

### GPCRdock-9686 Target 3 OPRK

#### Complex Model Generation

Complex models were generated by a local installation of AlphaFold-Multimer [1]. All peptide ligands were modeled at the correct position (within the helical bundle on the extracellular side).

#### Refinement and Submission

Models were simulated in NAMD [2] for 100ns, with the peptide, ECLs, and N-terminal free, and the everything else under harmonic restraint.  From the refinement MD trajectories, we extracted frames of the simulation that exist in potential energy minima. The selected models are as follows: 1) Top ranked AlphaFold model, 2) Top ranked AlphaFold model + minimization, 3-5) Frames from refinement MD at local energy minima. Files were cleaned in PyMol to match standard PDB format conventions before submission.

[1] R. Evans *et al.*, “Protein complex prediction with AlphaFold-Multimer,” Oct. 2021. doi: 10.1101/2021.10.04.463034.

[2] J. C. Phillips *et al.*, “Scalable molecular dynamics on CPU and GPU architectures with NAMD,” *J. Chem. Phys.*, vol. 153, no. 4, p. 044130, Jul. 2020, doi: 10.1063/5.0014475.

### GPCRdock-9686 Target 4 NPY1R

#### Complex Model Generation

Complex models were generated by a local installation of AlphaFold-Multimer [1]. All peptide ligands were modeled at the correct position (within the helical bundle on the extracellular side).

#### Refinement and Submission

Models were simulated in NAMD [2] for 100ns, with the peptide, ECLs, and N-terminal free, and the everything else under harmonic restraint.  From the refinement MD trajectories, we extracted frames of the simulation that exist in potential energy minima. The selected models are as follows: 1) Top ranked AlphaFold model, 2) Top ranked AlphaFold model + minimization, 3-5) Frames from refinement MD at local energy minima. Files were cleaned in PyMol to match standard PDB format conventions before submission.

[1] R. Evans *et al.*, "Protein complex prediction with AlphaFold-Multimer," Oct. 2021. doi: 10.1101/2021.10.04.463034.

[2] J. C. Phillips *et al.*, "Scalable molecular dynamics on CPU and GPU architectures with NAMD," *J. Chem. Phys.*, vol. 153, no. 4, p. 044130, Jul. 2020, doi: 10.1063/5.0014475.

### GPCRdock-9686 Target 5 NMUR2

#### Complex Model Generation

Complex models were generated by a local installation of AlphaFold-Multimer [1]. All peptide ligands were modeled at the correct position (within the helical bundle on the extracellular side).

#### Refinement and Submission

Models were simulated in NAMD [2] for 100ns, with the peptide, ECLs, and N-terminal free, and the everything else under harmonic restraint. From the refinement MD trajectories, we extracted frames of the simulation that exist in potential energy minima. The selected models are as follows: 1) Top ranked AlphaFold model, 2) Top ranked AlphaFold model + minimization, 3-5) Frames from refinement MD at local energy minima. Files were cleaned in PyMol to match standard PDB format conventions before submission.

[1] R. Evans *et al.*, “Protein complex prediction with AlphaFold-Multimer,” Oct. 2021. doi: 10.1101/2021.10.04.463034.

[2] J. C. Phillips *et al.*, “Scalable molecular dynamics on CPU and GPU architectures with NAMD,” *J. Chem. Phys.*, vol. 153, no. 4, p. 044130, Jul. 2020, doi: 10.1063/5.0014475.
